# Lineage-Specific Venom Gene Expression Shapes Chemical Diversity in Cephalopods

**DOI:** 10.64898/2026.04.09.716377

**Authors:** Praveena Naidu, José Ramón Pardos-Blas, Saurabh Attarde, Favour Achimba, Benjamin-Florian Hempel, Ioana Clotea, Belkes Stambouli, Kim N. Kirchhoff, Melvin Williams, Jennifer McCarthy-Taylor, Mariam Gelashvili, David Sharer, Afeeda Ali, Beatrix Ueberheide, Caroline B. Albertin, Mandë Holford

**Affiliations:** Graduate Center, Programs in Biology, Biochemistry, Chemistry, City University of New York, New York, New York, USA; Department of Chemistry and Biochemistry, City University of New York, Hunter College, Belfer Research Building, New York, New York, USA; The Department of Organismic and Evolutionary Biology, Harvard University, Cambridge, Massachusetts, USA; Veterinary Centre for Resistance Research, Freie Universität Berlin, Berlin, Germany; NYU Langone Health, New York, New York, USA; Marine Biological Laboratory, Woods Hole, MA 02543, USA; Harvard Museum of Comparative Zoology, Cambridge, Massachusetts, USA

**Author notes:** These authors contributed equally (Praveena Naidu, José Ramón Pardos-Blas).

## Abstract

Animal venoms represent a major source of chemical novelty, yet how venom compounds originate, diversify, and are maintained across deep evolutionary timescales remains poorly understood. This gap is especially pronounced in cephalopods, which evolved venom systems used in predation, defense, and sexual competition, but whose venom genetic architectures, secretory cell types, and venom-producing glands remain largely unexplored. To date, only a single cephalopod venom compound with confirmed paralytic activity and a known primary sequence, SE-CTX from the golden cuttlefish *Acanthosepion esculentum*, has been described. Here, we reconstruct the evolutionary history, molecular diversity, and glandular localization of SE-CTX–like proteins using a multimodal approach. We identify 29 homologs across 20 squid and cuttlefish species and define a previously unrecognized venom gene family, which we name *deca-ctx*, specific to decapodiform cephalopods (squids and cuttlefish). Phylogenetic analyses reveal a single origin of *deca-ctx* followed by gene duplication and lineage-specific diversification, indicating long-term retention of this venom gene. Predicted DECA-CTX protein structures were separated into two clusters and 20 singletons highlighting potentially extensive structural diversity within a single cephalopod venom gene family. Proteomic analysis confirms expression of five DECA-CTX proteins across three species. Our imaging and histological analyses localize *deca-ctx* expression to specialized secretory cells within squid and cuttlefish venom glands. Together, these findings reposition SE-CTX as part of an evolutionarily and chemically diverse venom system, rather than an isolated venom protein, and establish cephalopods as a key lineage for investigating how new venom genes arise, diversify, and are integrated into functional venom arsenals.

## Main Text

Venom is a successful biochemical and ecological adaptation that has evolved independently >100 times, across all major animal lineages, where it is predominately used for defense, predation, or sexual competition ^1,2^. Venom arsenals in individual species range from a few compounds to thousands ^1^. The diversity of venom has transformed our understanding of physiology and inspired therapeutic development ^3–7^. For example, bungarotoxin, found largely in venomous snakes, has been used extensively in neuroscience to characterize different subtypes of nicotinic acetylcholine receptors (nAChRs)^8^. Additionally, several blockbuster drugs for treating a variety of human diseases and disorders have been developed from venom components, such as Captopril® for hypertension from the Brazilian pit viper, Extendin® for diabetes and Ozempic® for diabetes and weight loss, both derived from the Gila monster, and Prialt® for chronic pain, from the cone snail *Conus magus* ^9–12^. Along with this incredible chemical diversity, venom delivery systems vary greatly in structure and physiology ^1^. However, the shared chemical, molecular, and cellular features across the diverse landscape of venom systems remain largely unknown. Despite advances in genomics and transcriptomics, our understanding of venom gene regulation and the development of venom-producing tissues remains limited. To begin to address this knowledge gap, we leveraged recent technological breakthroughs in cephalopod genome sequencing and husbandry to examine the evolution of venom genes across several cephalopod taxa ^13^.

Coleoid cephalopods comprise over 800 species of soft bodied octopus, cuttlefish, and squid (**Figure 1A**) ^14^. Coleoids are remarkably successful marine predators by virtue of an array of specialized traits including adaptive camouflage behaviour, highly complex nervous systems, and use of venom to subdue and capture prey ^14^. The toxic properties of cephalopod saliva were first discovered in the late 19^th^ century when Lo Bianco injected secretions from octopus venom glands into crabs, resulting in hindered locomotion and then death ^15^. Since then, venoms from over 20 species of cephalopods have been described using biological assays and -omic technologies ^16–19^. The main effect of cephalopod venom appears to be paralysis, as demonstrated in crabs, mice and locusts ^20,21^.

**Figure 1.**
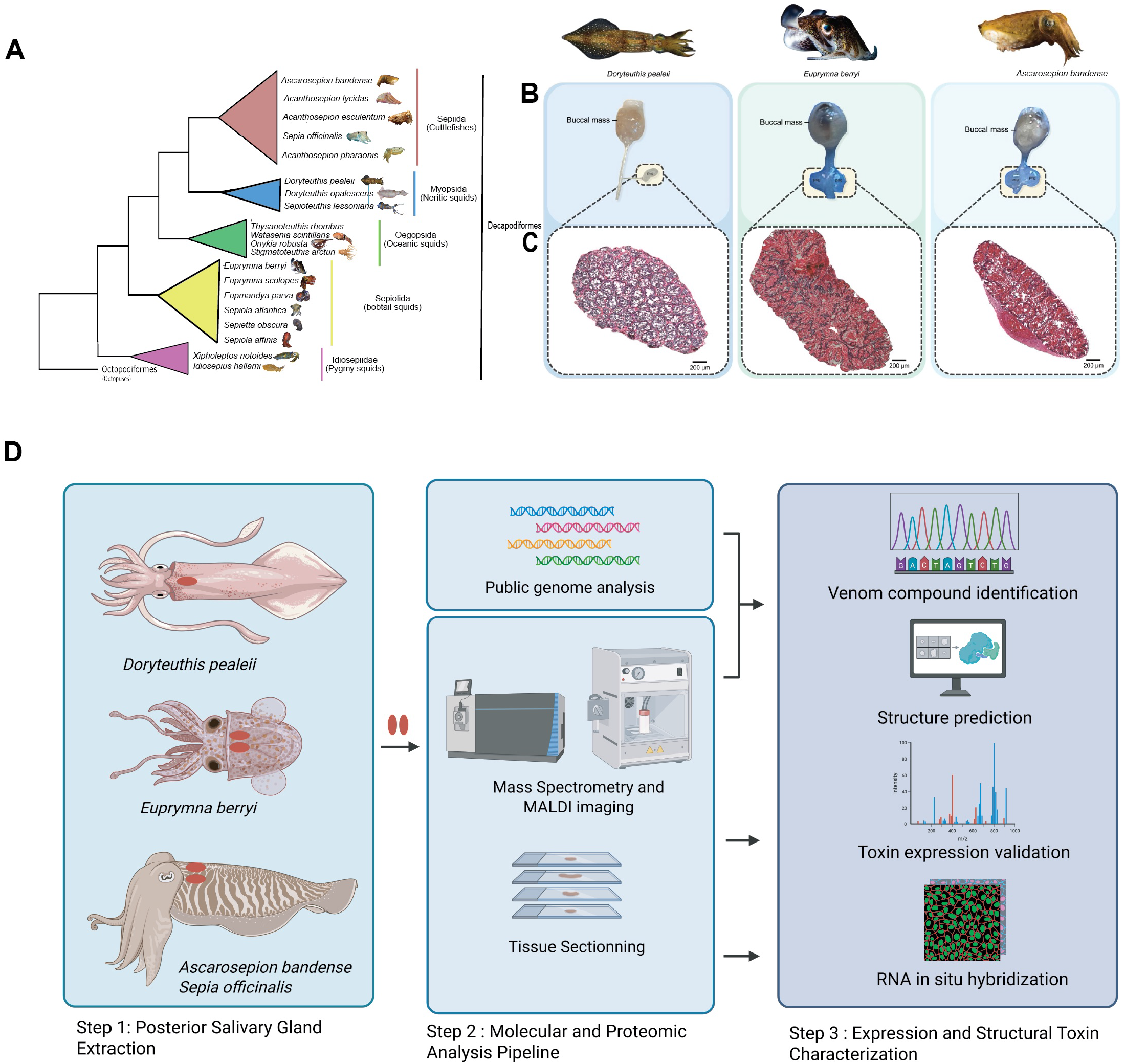
Decapodiformes are squids and cuttlefish with single or paired and posterior salivary glands. **(A)** Coleoid cephalopods can be broadly divided into decapodiformes (squids, cuttlefish) and octopodiformes. (**B)** Posterior salivary glands (PSG) were extracted from adult D*oryteuthis pealeii, Euprymna berryi* and *Ascarosepion bandense*. Positions of the intact salivary gland relative to esophagus and buccal mass for each of the three species are shown in cartoons and as actual extracted tissue. **(C)** H&E staining of paraffin-embeddings of these PSG sections reveal tissues with similar tubular morphologies. *E. berryi* and *A. bandense* appear to have more similar tubular structures in their PSG as compared to D. *pealeii*. **(D)** Pipeline used to identify, visualize and characterize putative cephalotoxin homologs across squid and cuttlefish species. In Step 1: Posterior salivary glands were extracted from *Doryteuthis pealeii, Euprymna berryi, Ascarosepion bandense* and *Sepia officinalis* species. In Step 2: Extracted posterior salivary glands were processed for RNA sequencing, Mass spectrometry and tissue sectioning. In Step 3: Publicly available genomic data along with generated transcriptomic data was used to identify the sequences of putative homologs to the previously identified *se-ctx* gene for available species. Mass spectrometry was used to validate venom gene expression and RNA-hybridization chain reaction was used to visualized expression of venom gene transcripts in adult and hatchling tissue sections. Protein structure prediction of the putative sequences was performed using AlphaFold2.0.

The posterior salivary gland (PSG) is the primary organ of cephalopod venom production ^15^. Sitting at the base of the mantle in close association with the digestive glands, they are typically found as pairs in octopus (Octopoda), cuttlefish (Sepiida), and the bobtail squid (Sepiolida) **(Figure 1B**). However, the PSG is a single gland in most species of myopsid and oegopsid squid. Several proteins with potential paralytic activity, termed cephalotoxins (CTX), are found in the PSGs of cephalopods. The nomenclature “CTX” refers to any protein expressed by the cephalopod PSG that exhibits paralytic activity ^20^. CTX has been identified in several cephalopod species including: *Acanthosepion esculentum, Acanthosepion pharaonis, Octopus vulgaris, Octopus macropus, Sepia officinalis, Eledone cirrhosa, Enteroctopus dofleini* ^22–26^. The first and only CTX with a known primary amino acid sequence, SE-cephalotoxin (SE-CTX), has been isolated from a paralytic fraction of the PSG of the golden cuttlefish (*A. esculentum; prior name: Sepia esculenta*, Uniprot: B2DCR8)^20^. SE-CTX shows low similarity to any other known venom compounds ^20^. However, homologs of *se-ctx* genes have been found in other decapodiformes (squids and cuttlefish, Figure 1) like the pharaoh cuttlefish (*A. pharaonis)* and in the common cuttlefish (*Sepia officinalis)* ^20,26^.

Here, we identify and characterize venom gene *se-ctx* homologs in 20 decapodiform species and visualize their expression in the PSGs of both hatchlings and adults (**Figure 1D**). Using comparative multimodal -omic, phylogenetic, and imaging datasets, we systematically characterize *se-ctx* homologs across several squid and cuttlefish species that share a common evolutionary lineage. We refer to this family of venom genes as *decapodiform-ctx* (*deca-ctx*) and the proteins they encode as Decapodiform-CTX (DECA-CTX). Specifically, we map *deca-ctx* expression and assess interspecific differential expression pattern in squid and cuttlefish PSGs. We describe the predicted 3D structure of identified DECA-CTX proteins, which form distinct structural clusters, suggesting that these proteins have multiple molecular targets. Our work provides evidence for widespread distribution of *se-ctx* homologs among squids and cuttlefish, suggesting that it is a common component of cephalopod venom arsenals and potentially evolved through gene duplication. Together, our findings establish the conservation of *se-ctx* expression and structural characterization across diverse squid and cuttlefish taxa, transforming SE-CTX from a single-species toxin into a deeply conserved and diversified element of the cephalopod venom system.

## Results and Discussion

### Posterior salivary gland morphology is highly conserved across adult cephalopods

To begin to examine cephalopod venoms, we conducted a comparative histological study of the PSG of the longfin inshore squid (*D. pealeii)*, hummingbird bobtail squid *(E. berryi)*, and dwarf cuttlefish (*A. bandense)*. Our PSG studies closely align with previous literature describing a variety of PSG cell types present ^26–28^. Specifically, the PSG tissue has a glandular appearance, with putative secretory cells surrounding a central lumen consistent with secretory glands (**Figure 1C**). We observed a relatively homogenous branched tubular arrangement, with *E. berryi* and *A. bandense* showing a greater similarity to each other compared to *D. pealeii* (**Figure 1C**). The cells present in each “tubule” of the *E. berryi* and *A. bandense* sections show an abundance of granules in the apical region of the cells towards the lumen of the branched tubules, compared to *D. pealeii*, whose granules appear less densely packed. These findings confirm that PSGs from diverse cephalopod taxa display conserved cellular and histological features.

### The family *deca-ctx* is duplicated across decapodiform species

To reconstruct the phylogenetic relationships of *se-ctx-like* genes that comprise the newly defined *deca-ctx* family, we surveyed available genomic data ^29–34^. We identified open reading frames (ORFs) with high similarity to originally identified *se-ctx* sequences in 18 species **(Figure 2A, Supplementary Figure 1, Supplementary Table 1**). We also found partial sequences with high similarity to *se-ctx* in two additional species, clubhook squid (*Onykia robusta)* and the firefly squid (*Watasenia scintillans)*. In the golden cuttlefish (*A. esculentum)*, the cephalopod in which the original SE-CTX protein was isolated and identified, we found an additional putative *se-ctx* gene on the same chromosome sequences which has nucleotide sequence similarity of 51.9% to the previously identified *se-ctx* gene. In species with genomic data available, we found two *deca-ctx* genes, *deca-ctx1* and *deca-ctx2*, in most decapodiforms **(Figure 2A**). However, all of the bobtails we examined seem to possess a single paralog of *se-ctx, deca-ctx1*. These results suggest that the last common ancestor of extant decapodiformes possessed two *se-ctx*-like genes, one of which was independently lost in different lineages. For example, in the four species of bobtails examined, all possess a single *deca-ctx* gene that branches with *deca-ctx1* in other lineages, suggesting a loss of *deca-ctx2* specific to a common ancestor of bobtail lineage. In contrast, the pygmy squid (*Xipholeptos notoides*) possesses both *deca-ctx1* and *deca-ctx2*, but Hallam’s pygmy squid (*Idiosepius hallami*) only has *deca-ctx1*, suggesting independent loss of *deca-ctx2* in this lineage (**Figure 2A**). The loss of venom function is not uncommon in mollusks. The terebrid marine snail lost their venom apparatus at least eight times in their evolution^35^. The evolutionary and ecological drivers for these events are not resolved.

**Figure 2:**
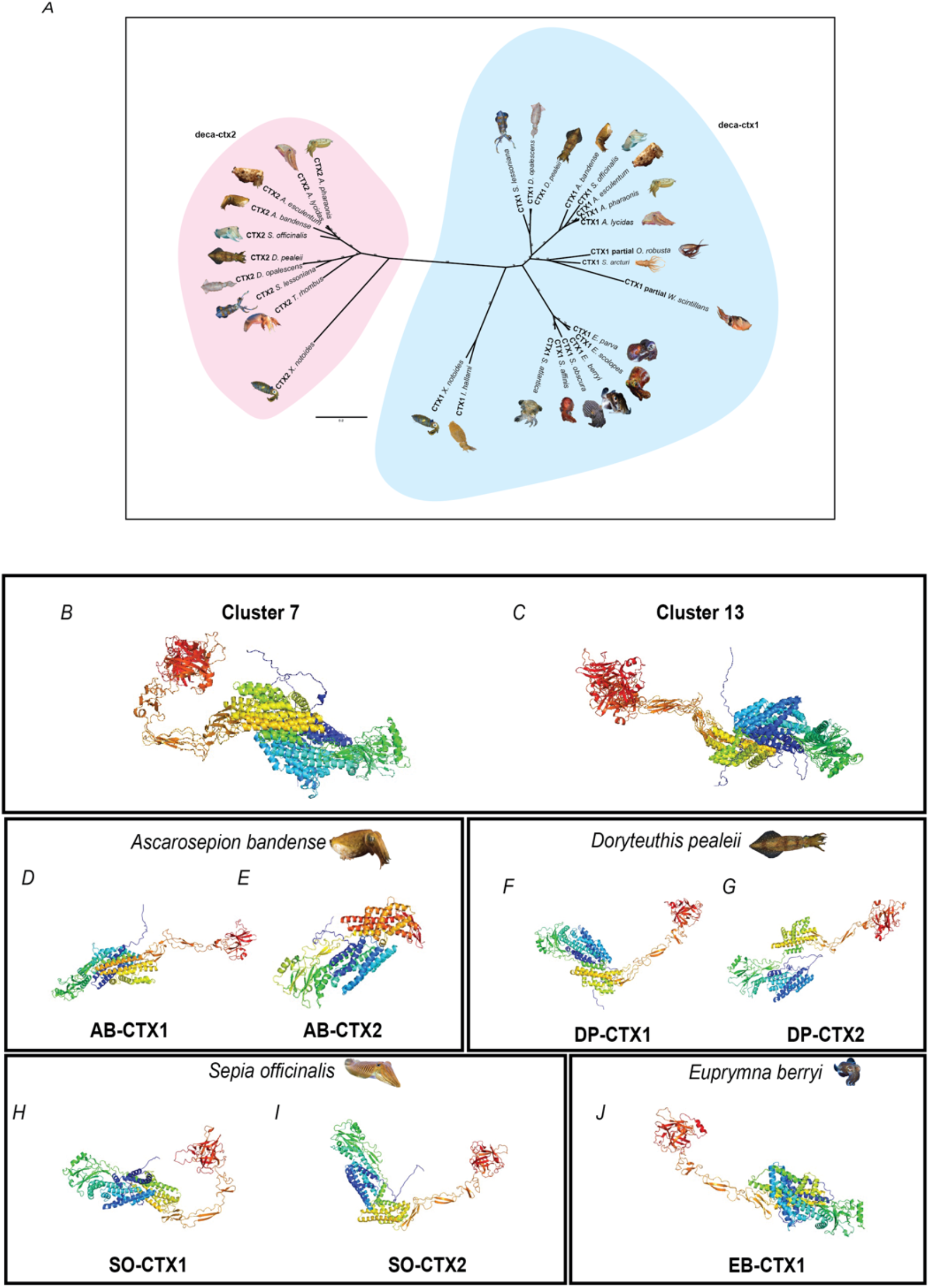
Duplication of cephalotoxin *se-ctx* gene present in squid and cuttlefish lineages display diversity of chemical structures. **(A)** Unrooted Maximum likelihood inference of the *se*-*ctx* homolog sequences of cephalopods. Two distinct paralogous se-*ctx genes, deca-ctx1*and *deca-ctx2*, are revealed in the tree reconstruction. The blue clade indicates *deca-ctx1* and includes genus *Euprymna, Eumandya, Sepiola, and Idiosepius genera* species exclusively. The pink clade indicates *deca-ctx2* and includes genera *Ascarosepion, Acanthosepion, Sepia, Doryteuthis, and Sepioteuthis genera*, but no Bobtail squid (*Euprymna)* species. The values represent the node support in percentage (1000 bootstrap). Cephalopoda species curated in this database version (Accessed July 2025) are included in the present tree. **(B-J)** AlphaFold2.0 predicted structures and clustering of Deca-CTX homologs. All predicted structures were clustered with qTMclust from TM-align and fell into two major clusters: (B) Cluster7 made up of 2 structures, and (C) Cluster13, also made up 3 structures. Shown also are representative singleton predicted structures of species-specific Deca-CTX from (D,E) *Ascarosepion bandense*, (F,G) *Doryteuthis pealeii*, (H,I) *Sepia officinalis*, and *(J) Euprymna berryi*. All structures are colored by chainbow using PyMol.

Our phylogenetic analyses of *deca-ctx* genes identified 29 genes across 20 species, revealing an evolutionary origin that predates the origin of Decapodiformes, followed by lineage-specific gene losses (**Figure 2A, Supplementary Figure S2**). Similarity searches (TBLASTN) searches of the *se-ctx* sequence against the core_nt and refseq_genomes at NCBI did not identify *se-ctx*-like sequences in non-decapodiform mollusks, including in octopuses. Therefore, the duplication of *deca-ctx* genes that we found in Decapodiformes is likely exclusive to squids and cuttlefish.

### DECA-CTX structures group into two distinct clusters and 20 structurally divergent singletons

Determining the significance of the sequence variations of the *deca-ctx* gene homologs on paralytic activity requires an understanding of the mechanism of action at the protein level. To date, the exact mechanism of any cephalotoxin is not well understood. Additionally, whether the mechanisms of action for SE-CTX and/or the DECA-CTX paralogs identified here are similar to those of other cephalotoxins is unknown. Prior functional experiments with octopus were performed with crude extracts, and there are no reported sequences for the active components ^15,21,24,36^. To determine the potential molecular activity of the DECA-CTX paralogs we identified, we predicted their structures using AlphaFold2.0 ^37^ (**Figure 2B-J**). We used TM-align to evaluate structural homology between DECA-CTX proteins ^38^. Notably, two structural clusters containing five DECA-CTX proteins were identified: A cluster of two DECA-CTX proteins IHAL-CTX1 and XNOT2-CTX1 (**Cluster7, Figure 2B**) and a cluster of three homologs SLES-CTX1, AB-CTX1 and SLYC-CTX1(**Cluster13, Figure 2C**). Twenty DECA-CTX proteins were highly structurally divergent and due to the large overall protein size and low pairwise root mean square deviation (RMSD) and template modelling (TM) scores, they did not form clusters, and are herein denoted as “singletons”. Members of clusters 13 and 7 exhibited a characteristic “mini beta barrel” feature described by 4-7 alpha helices linked by a flexible beta-sheet-loop-helix connector forming a “hockey stick-type” topology. This “hockey stick” architecture is represented with slight variations across all DECA-CTX proteins except AB-CTX1 which presents a largely alpha helical topology. It is not clear how these characteristic features correlate to functional activity, and to address that we performed an analysis of the conserved domains found in these proteins.

Analyses of conserved domains present in several of DECA-CTX sequences highlight potential functional activity (see **Supplementary Figure 3-4** and **Supplementary File 1** for comparison of conserved domains across these DECA-CTX proteins). For example, DP-CTX2, SP-CTX2, SO-CTX2 and XN-CTX2 have regions that match Low-Density Lipoprotein (LDL) Receptor class A, Sushi also known as Complement Control Protein (CCP) module or Short Consensus Repeat (SCR), Thrombospondin type 1 repeats (TSRs or TSP-1), and Epidermal Growth Factor (EGF) domains. (**Supplementary Figure 3-4, Supplementary Table 2**). Each of these domains are cysteine-rich motifs that are stabilized by multiple disulfide bonds and are usually important in binding interactions, such as ligand or receptor binding in the case of LDL-class A and EGF, respectively. This suggests that DECA-CTX proteins may modulate their paralytic activity through high-affinity interactions with extracellular ligands or cell-surface receptors involved in neuromuscular signaling. Given the diversity of the predicted structures, functional assays using these putative DECA-CTX homologs either isolated from PSG tissue or recombinantly synthesized are required to elucidate their mechanisms of action and whether the presence or absence of the conserved domains in most of the identified DECA-CTX sequences affect their venom activity.

### *deca-ctx* transcripts are expressed in PSGs of adult, hatchlings, and late-stage embryos

To test for *deca-ctx* gene expression in squids and cuttlefish, we used mRNA *in situ* hybridization by chain reaction (HCR) to visualize transcript expression in dissected PSGs of four species: *D. pealeii, E. berryi, S. officinalis* and *A. bandense*. In all four species, we found *deca-ctx* expression to be specific to cells surrounding the lumens of secretory granules in the PSG (**Figure 3)**, and we observed no expression in surrounding oesophagus or buccal mass tissue (**Supplementary Figure 5**). In *D. pealeii*, both *deca-ctx* paralogs (*dp-ctx1* and *dp-ctx2*) were found to be highly co-expressed **(Figure 3Ai-vi**, 20x vs 63x magnification, respectively). In contrast, in A. *bandense*, paralogs (*ab-ctx1* and *ab-ctx2*) were expressed in completely distinct cells within the PSG **(Figure 3Bi-vi)**. Similarly, in *S. officinalis*, paralogs (*so-ctx1* and *so-ctx2*) were expressed predominately in distinct cells within the PSG **(Figure 3Ci-vi)**. In *E. berryi*, we saw clear expression of their single *eb-ctx1* throughout their PSG as well **(Figure 3Di-ii)**. These results are the first to visualize the conservation of venom gene expression in cephalopod venom glands. Moreover, these experiments highlight the evolution of deca-*ctx* gene expression within the cell types of the PSG and indicate that while some species, such as *D. pealeii*, appear to co-express deca-*ctx* paralogues, others, like A. *bandense* express distinct deca-*ctx* paralogues in a non-overlapping manner (**Figure 3A-B**). The broad expression of *deca-ctx* genes across squid and cuttlefish suggests it is an important component of decapodiform venom arsenals.

**Figure 3:**
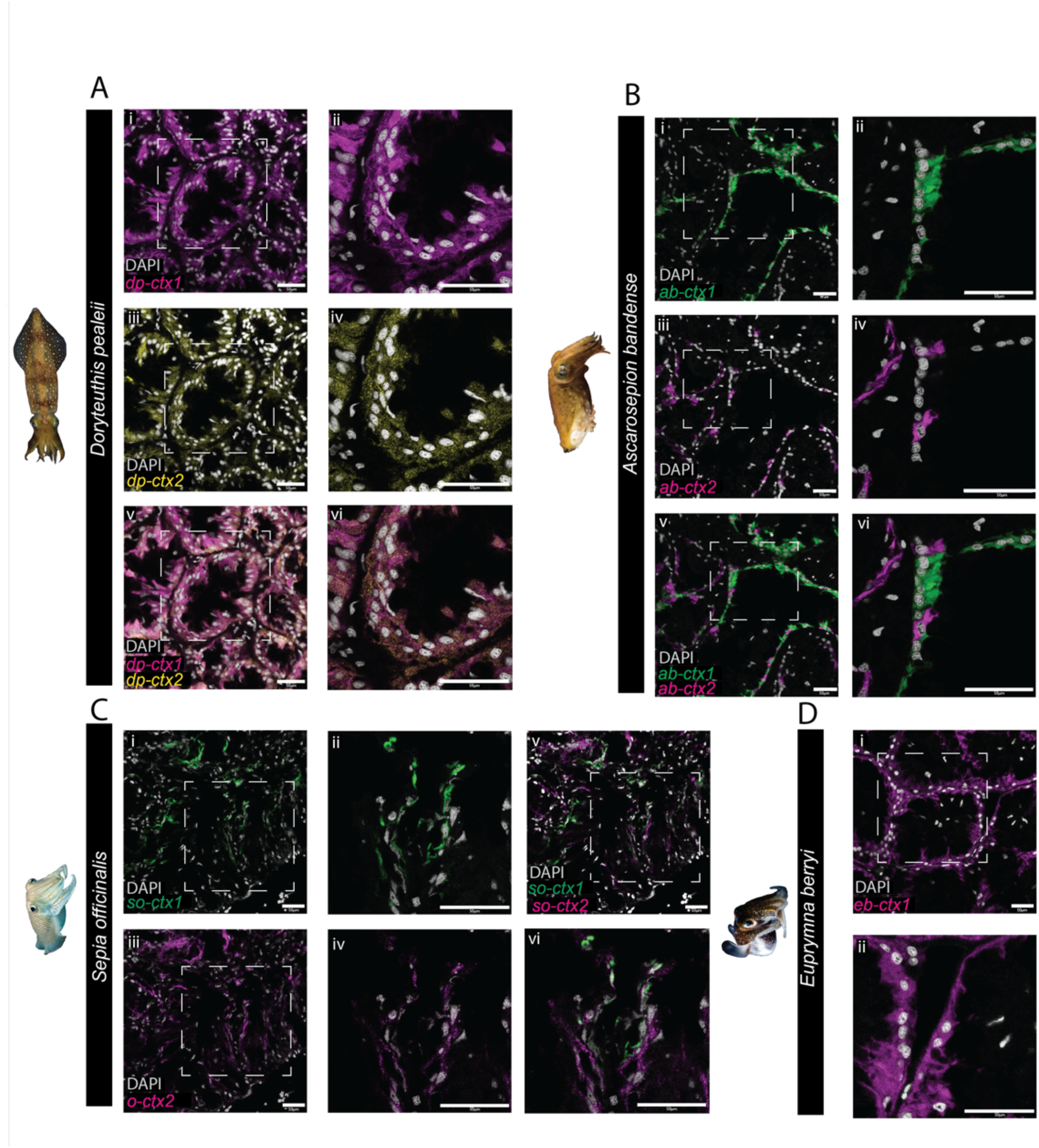
Visualisation of *dual deca-ctx* expression in adult posterior salivary gland tissue indicate heterologous tissue localization. Hybridization Chain Reaction fluorescence microscopy images of PSG sections in adult specimens showing the expression of **A)** *Doryteuthis pealeii* dp-*ctx1* mRNA (magenta) at **(i)** 20x and **(ii)** 63x magnification; *Doryteuthis pealeii dp-ctx2* mRNA(yellow) at **(iii)** 20x and **(iv)** 63x magnification; dp-*ctx1* mRNA (magenta) and *dp-ctx2* mRNA(yellow) at **(v)** 20x and **(vi)** 63x magnification. **B)** *Ascarosepion bandense* ab-*ctx1* mRNA (green) at **(i)** 20x and **(ii)** 63x magnification; *Ascarosepion bandense ab-ctx2* mRNA(magenta) at **(iii)** 20x and **(iv)** 63x magnification; ab-*ctx1* mRNA (green) and *ab-ctx2* mRNA(magenta) at **(v)** 20x and **(vi)** 63x magnification. **C)** *Sepia officinalis so-ctx1* mRNA (green) at **(i)** 20x and **(ii)** 63x magnification; *Sepia officinalis so-ctx2* mRNA(magenta) at **(iii)** 20x and **(iv)** 63x magnification; *so-ctx1* mRNA (green) and *so-ctx2* mRNA(magenta) at **(v)** 20x and **(vi)** 63x magnification. **D)** *Euprymna berryi eb-ctx1* mRNA (magenta) at **(i)** 20x and **(ii)** 63x magnification. Sections are counterstained with DAPI (grey) to visualize nuclei.

To determine when during squid and cuttlefish development the *deca-ctx* genes are expressed, we applied *in situ* HCR for *deca-ctx* transcripts in hatchlings and embryos of *E. berryi* as well as in the hatchlings *of D. pealeii*, and *A. bandense (***Figure 4**). We observed no *eb-ctx1* staining in before late embryonic stages in *E. berryi* embryos (**Figure 4A**). However, *eb-ctx1* is expressed at the base of the mantle in the shape and the region where we would expect to see the two PSGs near hatching (stage 29/30) (**Figure 4B**). Additionally, in *E. berryi* hatchlings (∼30 days old), we observe clear and specific expression of *eb-ctx1* in the PSGs and no expression in the oesophagus and surrounding brain tissue (**Figure 4C**). These findings suggest that the expression of *deca-ctx* begins within the PSG shortly before hatching in *E. berryi*. In *D. pealeii* hatchlings, we observed *dp-ctx1* expression but no *dp-ctx2* expression **(Figure 4D-E)**. Since *dp-ctx1* and *dp-ctx2* are co-expressed in the PSGs of adult *D. pealeii* (**Figure 3A**), differences in expression in the hatchlings suggests these genes are differentially regulated across life stages in this species. Similar to *E. berryi* hatchlings, *ab-ctx1* expression is present in left and right PSGs at the hatchling stage in *A. bandense* (**Figure 4F**). Taken together, this suggests *deca-ctx1* venom transcript expression is present in the hatchling stage in these three species, potentially allowing them to use their venom at this early stage of development.

**Figure 4:**
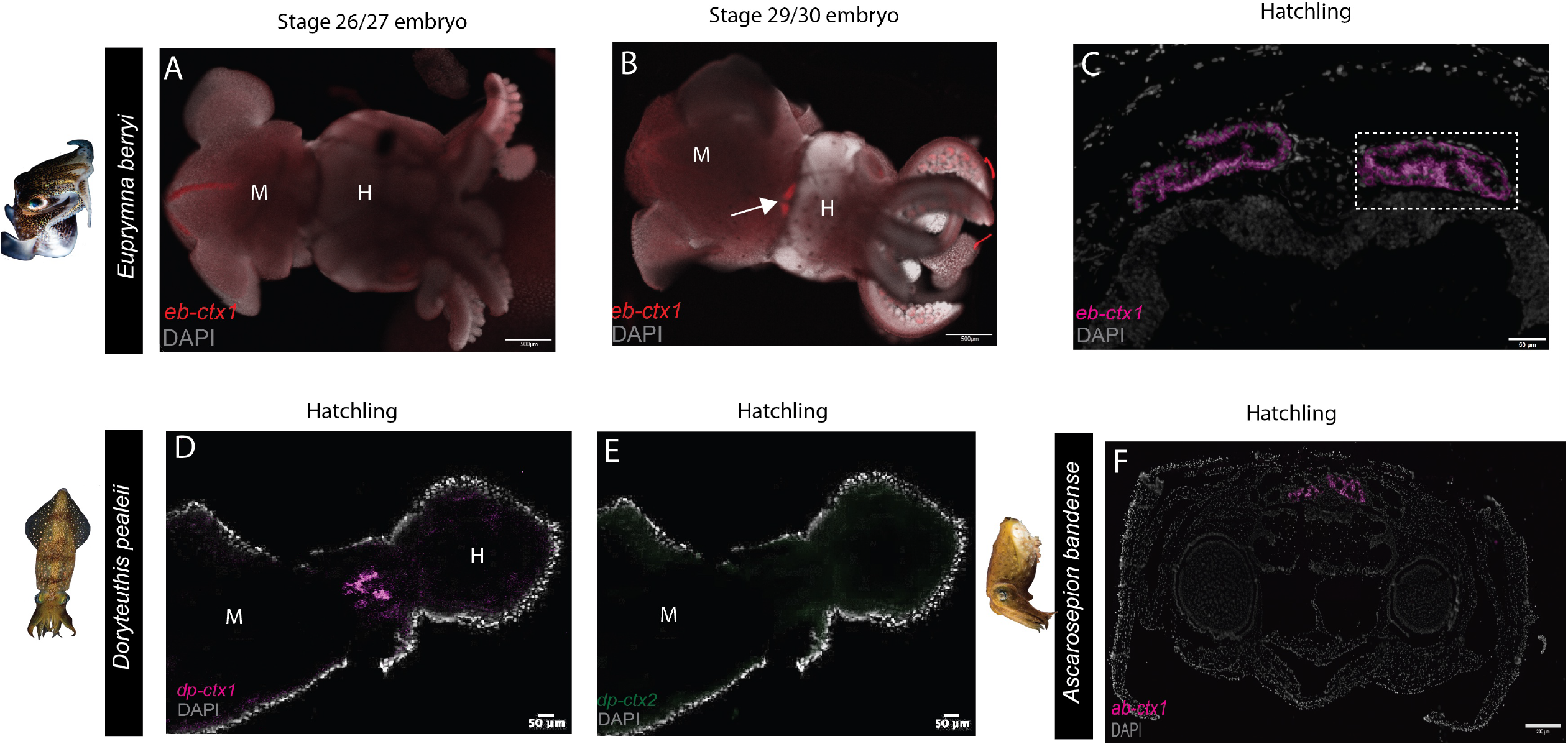
Differential developmental expression of deca*-ctx* homologs in squid and cuttlefish hatchlings and embryos. Hybridization Chain Reaction fluorescence microscopy images. **(A)** Whole mounts of *E*.*berryi* embryos were stained with *eb-ctx1*-specific (red) probe and DAPI revealed no staining in day 26/27 embryos. The redline shown on the mantle opposite the tenetacles is distinctive nonspecific staining in *E. berryi* embryos. (**B) C**lear staining was seen in day 29/30 embryos/hatchlings. White arrow indicates PSG-specific expression. (**C)** PSG-specific expression of *eb-ctx1* mRNA in *E*.*berryi* hatchling sections. The two E. berryi PSGs both express *eb-ctx1*. Enlarged view highlighted in the white hashed-box. Whole mounts of *D*.*pealeii* hatchlings were stained with *dp-ctx1-*specific (magenta) probe, *dp-ctx2-*specific (green) probe and DAPI, **(D)** Indicates *dp-ctx1* staining in the region of the two PSGs. **(E)** No *dp-ctx2* staining is present in the same whole mount.. **(F)** Similar to *E. berryi*,, we observe PSG-specific expression in both glands of *Ab-ctx1* mRNA in *A* .*bandense* hatchling sections. Sections are counterstained with DAPI (grey) to visualize nuclei.

### Mass spectrometry confirms DECA-CTX protein expression in squid and cuttlefish venom glands

To validate expression of DECA-CTX proteins, we used a multi-platform approach of bottom-up mass spectrometric analysis and matrix-assisted laser desorption/ionization (MALDI) mass spectrometry imaging (MSI) to characterize and visualize DECA-CTX proteins in the PSGs of *A. bandense, D. pealeii and E. berryi* (**Figure 5**). Bottom-up analysis of the trypsin-digested crude venom extracts from the PSGs of *A. bandense, D. pealeii*, and *E. berryi*, identified several key venom protein families, including DECA-CTX, cysteine-rich secretory proteins, phospholipase A2 and B, and serine proteases (**Figure 5A**). We found *ab-ctx1, ab-ctx2, dp-ctx1, dp-ctx2*, and *eb-ctx1* sequences from transcriptome and genome databases, confirming the presence of DECA-CTX proteins in the venom arsenals of *A. bandense, D. pealeii*, and *E. berryi* (**Supplementary File 2**). Importantly, peptide matches were highly specific to *deca-ctx* gene sequences of the same species. For example, AB-CTX1 and AB-CTX2 peptides from *A. bandense* matched exclusively with their corresponding transcript sequences and not with those from *D. pealeii* or *E. berryi*. In specimens with two PSGs, such as *E. berryi* and *A. bandense*, base peak chromatograms of the extracted venom samples were largely identical, suggesting there is no differential expression between the left and right PSGs.

**Figure 5.**
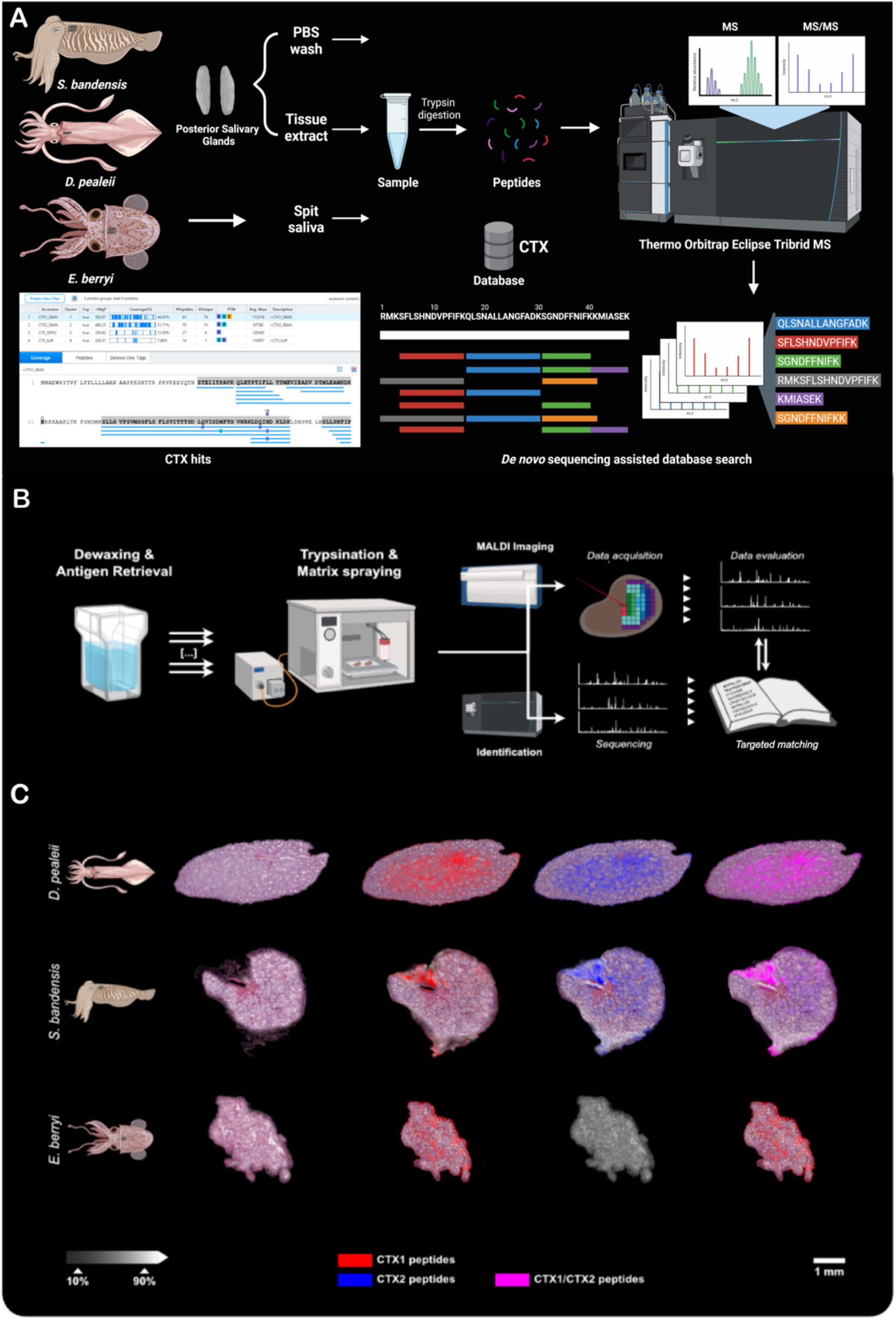
Proteomic confirmation of DECA-CTX protein expressions validates presence of transcripts in squid and cuttlefish venom glands. Shown is the bottom-up and spatial venom proteomic analyses of posterior salivary gland tissue sections for three different cephalopod venom systems. **(A)** Schematic of the mass spectrometry (LC-MS/MS) workflow used to identify cephalotoxins (CTXs) from the posterior salivary glands (PSGs) of *A*.*bandense, D. pealeii*, and *E. berryi*. PSGs from *A*.*bandense, D. pealeii*, and *E. berryi* were dissected, homogenized in PBS, lyophilized, and processed for proteomic analysis. **(B)** Preparation and analysis of spatial venomics using matrix-assisted laser desorption/ionization (MALDI) mass spectrometry imaging (MSI). Tissue sections of venom glands from *D. paeleii, E. berryi*, and *A. bandense* were deparaffinized, rehydrated, tryptic digested and loaded on to a ultrafleXtreme MALDI-ToF/ToF mass spectrometer. Raw data were processed using baseline correction, TIC normalization, and peak alignment. Peptide matches between MALDI-MSI and nLC-MS peptide library were identified using an in-house script, matching m/z values within 0.2 Da and selecting peptides based on the highest confidence score. **(C)** Visualization of DECA-CTX1, red), DECA-CTX2, blue) and both DECA-CTX1 and DECA-CTX2, purple) by characteristic m/z values (mass features) within the venom gland system (20 μm spatial resolution). The spatial venom compound distribution and internal relative intensities (white scale bar) are shown for a set of three mass features per compound and species Top row: *D. pealeii* composed of m/z 1376.79, 1697.91, 2458.13 (DECA-CTX1) and m/z 1287.74, 1725.86, 2082.08 (DECA-CTX2). Middle row: *A*.*bandense* composed of m/z 1123.60, 1564.65, 1709.61 (DECA-CTX1) and m/z 1621.85, 1974.82, 2226.06 (DECA-CTX2). Bottom row: *E. berryi* composed of m/z 1584.82, 1798.66, 2229.96 (DECA-CTX). The histological image was prepared post-MALDI-MSI by hematoxylin/eosin (H/E) staining and is shown for tissue section orientation. The spatial segmentation of the venom gland systems resulted in different numbers extracted peak ions (±0.47 Da) from normalized spectra within an m/z 600™3200 mass range (see Supplementary Tables 2).

MALDI-MSI analyses confirmed the presence of DECA-CTX proteins and allowed us to clarify differential spatial expression in the PSG by characteristic *m/z* values (mass features) within the venom gland system at 20 μm spatial resolution (**Figure 5B, C**). *D. pealeii* DECA-CTX proteins appears to be mostly in the channels in the center of the glands and along the peripheral membrane **Figure 5C, top row)**. In contrast, in *A. bandense* and *E. berryi*, there is more of a punctuated distribution in channels in the gland and along the membrane (**5C, middle and bottom rows)**. Additionally, spatial venomics corroborates transcriptomic and bottom-up findings that *E. berryi* expresses EB-CTX1 (**Figure 5C, bottom row**). The unique distributions of *deca-ctx* homologs in the PSGs could reflect how the protein is expressed and released. Differential expression of predatory versus defensive venoms has been suggested to be related to spatial distribution in various venom glands ^39^. The identification of species-specific DECA-CTX sequences by mass spectrometry aids in our understanding of evolutionary divergence in the venom arsenal of cephalopods. This divergence potentially reflects adaptations to distinct ecological niches or diets ^35^.The sequence variation present in the DECA-CTX family needs to be further investigated for their biological activity to clarify the impact of these variations on the protein or peptide’s bioactivity, target specificity or regulatory control.

In summary, we present the first systematic, comparative analysis of *ctx* genes across cephalopods, revealing their diversity in molecular structure and spatial expression. Phylogenetic analyses suggest that *deca-ctx* genes originated early in cephalopod evolution and diversified through lineage-specific expansions, resulting in pronounced variation in sequence composition and domain architecture. Notably, *deca-ctx* genes are expressed in the PSGs of hatchlings, suggesting this venom repertoire may be available to deploy from early life stages for predation or defense. Although the functional activity of the identified *deca-ctx* homologs remains to be experimentally determined, conserved domain analyses reveal the presence of key structural motifs characteristic of the paralytic SE-CTX toxin. These domains, LDL-Class A, Sushi, TSP-type1, and EGF, often appear in the same protein and are small cysteine-rich modules that play a role in binding interactions related to cell signaling, among other activity. The conservation of these domains suggests functional similarity and implicates these proteins in venom-mediated prey immobilization. The combination of conserved structural cores and variable domains is also consistent with functional diversification and suggests that *deca-ctx* proteins may span a range of biochemical properties and venom activities.

We integrated comparative genomics, phylogenetics, imaging, and mass spectrometry to validate the expression of *deca-ctx1* and *deca-ctx2* in longfin inshore squid (*Doryteuthis pealeii)*, hummingbird bobtail squid (*Euprymna berryi)*, dwarf cuttlefish (*Ascarosepion bandense)*, and common cuttlefish (*Sepia officinalis)*, and to corroborate predicted molecular heterogeneity at the protein level. This multi-modal approach enables direct linkage between gene-level diversity, structural variation, and expressed venom components; moving beyond gene prediction to experimentally grounded validation of venom arsenals in traditionally non-model organisms like cephalopods. Together, our findings provide a snapshot of the molecular and structural diversity of cephalopod venoms while establishing a foundation for future studies examining how evolution, development, sequence divergence, tissue specificity, and three-dimensional structure contribute to functional diversification of *deca-ctx* homologs across biological scales in one of the most ancient venomous animal lineages ^40^. Cephalopods are a comparative context for examining how gene duplication and structural innovation drive venom diversification across ecological and evolutionary timescales. More broadly, our study advances understanding of venom evolution and highlights cephalopods as a robust comparative model for studying underexplored structurally diverse bioactive molecules.

## Methods

### Ethics Statement

All experimental procedures involving cephalopods were conducted in accordance with relevant ethical guidelines with IACUC approval. Efforts were made to minimize animal discomfort and reduce the number of individuals used.

### Paraffin embedding and sectioning

Samples were collected at the Marine Biological Laboratory in Woods Hole, MA (USA). Adult *E. berryi, A. bandense* and *S. officinalis* dissections were performed by Dr. Lisa Abbo. Adult posterior salivary gland samples were incubated overnight in 4% paraformaldehyde (PFA) at 4°C. Samples were washed with DEPC-PBS then dehydrated stepwise into 100% ethanol. Samples were cleared in 2 changes of histosol for 30 minutes each, incubated in two 1:1 histosol:paraffin solution changes for 30 minutes each and incubated in paraffin overnight. Samples were then incubated in four 1 hour changes of paraffin before being placed in a sectioning mold for solidification. Paraffin blocks were sectioned on microtome as 10 µm sections. Water and PBS solutions used were diethylpyrocarbonate (DEPC)-treated overnight and autoclaved.

### *Se-ctx* identification, alignment and tree building

Transcripts with high similarity to *se-ctx* were identified through BLAST searches against publicly-available genomic data (**Supplementary Table 1**). Protein sequences were aligned using Muscle 5.1 and visualised with Geneious Prime® 2024.0.5 (https://www.geneious.com) ^41^. To build phylogenetic tree, nucleotide sequences were aligned using Muscle 5.1. PartitionFinder2 was used to identify best partition (Scheme_step_1 = (pos1, pos3) (pos2)) and nucleotide substitution model (GTR GAMMA I) ^42^. Maximum likelihood phylogeny of *se-ctx* genes was estimated using RAxML 8.2.11 ^43^. Phylogenetic tree was visualized using Geneious Prime® 2024.0.5.

### Structure prediction and clustering

Cephalotoxins structures were predicted using AlphaFold2.0 installed on a local computer and available on the google deepmind github page. The models were predicted with templates available in the downloaded databases listed on the github page (https://github.com/google-deepmind/alphafold). A python script was used to run an iterative prediction of all the sequences. The sequences that were not included in the list were run with template directly from the command line using the command described in the github page. The predicted structures were clustered using TMalign’s qTMclust on the command line. TMalign groups structures based on TM-scores and TM-align threshold was set at 0.7. Twenty-two clusters were obtained.

### Section *In situ* Hybridization Chain Reaction

Hybridized probes were designed by Molecular Instruments (MI; L.A., USA) based on six putative *deca-ctx* sequences from *D. pealeii, E. berryi, S. officinalis* and *A. bandense* (**Supplementary Table 1**). The *in situ* HCR protocol followed Choi *et al*. (2018) with modifications according to Criswell and Gillis (2020) ^44,45^. Briefly, slides were deparaffinized, rehydrated, and treated with 10 µg/mL proteinase K at 37 °C for 10 minutes, followed by a rinse in DEPC water. After pre-hybridization at 37 °C for 30 minutes in MI-hybridization buffer, probe sets (0.8 µL of 1 µM stock per 100 µL buffer) were applied and incubated overnight at 37°C. Slides were then washed in a series of MI-HCR wash buffer/5x SSCT solutions Hairpins (H1 and H2) were added to amplification buffer (200 µL per slide). After a 30-minute pre-amplification step, slides were incubated overnight in MI-amplification buffer with Hairpins (H1 and H2) at room temperature in the dark. On the third day, slides were washed in the dark as follows: 5x SSCT for 5 minutes, 5x SSCT for 15 minutes twice, and 5x SSCT for 5 minutes. Coverslips were applied with Fluoromount G+DAPI (thickness 1.5 mm). Slides were stored in the dark and allowed to dry overnight before being stored at 4 °C in the fridge.

### Whole mount *In situ* Hybridization Chain Reaction for *E*. *berryi*

Two *E. berryi* embryos at stages 26/27 and 29/30 of development were collected, and the chorion was gently removed under a binocular microscope. The embryos were anesthetized in a 1:1 7.5% w/v MgCl_2_ solution in seawater in a 1.5 mL Eppendorf tube and then, briefly submerged in boiling water. After cooling, embryos were fixed in 4% PFA for 2 hours, followed by washes with PBS-T and detergent. Embryos were transferred in preheated storage buffer at 37 °C and subsequently stored at -20 °C until hybridization. Whole-mount hybridization chain reaction was performed in two key steps: probe incubation and hairpin amplification as described in Ahuja e al., 2023^46^. Briefly, fluorophore-conjugated hairpins and probes were newly designed and purchased from Molecular Instruments Inc. for the most highly expressed component in cephalopod venom, cephalotoxin, as well as for the embryonic development marker hedgehog as positive control. First, embryos were pre-incubated in hybridization buffer (Cat. BPH02726, Molecular Instruments) at 37 °C for 30 minutes, followed by overnight incubation at 37° C in hybridization buffer containing the probe solutions. After probe incubation, embryos were washed with wash buffer at 37 °C in a series of steps (5, 15, and 30 minutes), and incubated in wash buffer for 1 hour at 37 °C. Second, for the hairpin incubation, hairpins (H1 and H2) were heat-shocked at 95 °C for 90 seconds and dark-incubated for 30 minutes at room temperature. Embryos underwent several wash steps 5x SSCT buffer and amplification buffer (Cat. BAM02926, Molecular Instruments) at room temperature. Then, embryos were incubated with the hairpin solution overnight at 25 °C. Final washes were performed, including one DAPI staining step (1:1000), and subsequently, the embryos were stored in 5x SSCT buffer until imaging on Zeiss LSM900 Confocal microscope.

### Microscopy and Image Processing

HCR slides were imaged with Zeiss LSM880 confocal microscope with 10x, 25x, 40x and 63x magnification with 405 nm, 488 nm and 647 nm wavelengths for detection. H&E brightfield images were taken with Zeiss Axioscope5. Whole mount imaging was conducted on a Zeiss LSM900 confocal microscope with 10x magnification, setting 488 nm and 647 nm wavelengths for detection and an image resolution of 1024 × 1024 pixels. Images were processed with Nikon elements, Zen v.3 and FIJI and overlayed in Adobe Photoshop.

### Bottom-up Liquid Chromatography - Mass Spectrometry Analysis

PSGs were dissected from *A. bandense, D. pealeii*, and *E. berryi*, and stored at -80 °C until further processing. Three types of samples were prepared from each species: spit saliva directly collected from the beak, PBS washes of the PSGs, and PSG tissue homogenates. For the homogenates, tissues were thawed on ice, homogenized in PBS, centrifuged to remove debris, and the supernatant was lyophilized. All samples were reconstituted in HPLC-grade water and subjected to reduction with 15-30 mM dithiothreitol (DTT), alkylation with 35-70 mM iodoacetamide (IAA), and trypsin (400 ng) digestion overnight at room temperature. Digested peptides were acidified with 10% trifluoroacetic acid (TFA) to a pH of 2, desalted using Pierce C18 spin columns, and sequentially eluted using 40% and 80% acetonitrile in 0.5% acetic acid. Organic solvents were removed via vacuum centrifugation, and peptides were reconstituted in 0.5% acetic acid prior to LC-MS/MS analysis.

Peptide analysis was conducted on an EASY-nLC 1200 system coupled to an Orbitrap Eclipse Tribrid mass spectrometer (ThermoFisher Scientific). A portion of each sample (varying from 1/90th to 1/10th of the starting material, depending on sample type and species) was injected and separated using a 145-minute gradient, with mobile phases consisting of 2% (solvent A) and 80% acetonitrile (solvent B) in 0.5% acetic acid. The gradient was held for 5 min at 5% solvent B, ramped in 120 min to 35% solvent B, in 10 min to 45% solvent B, and in another 10 min to 100% solvent B. “High resolution full MS spectra were obtained at 120,000 resolution for a scan range of 100-1500 m/z with an AGC target of 4×105 and a maximum ion time of 50 ms. Fragment ion scans were acquired in the orbitrap following HCD fragmentation (resolution 30,000, AGC target 2×105, maximum ion time 200ms, 1m/z ion isolation window, NCE 27%). Data were analyzed using PEAKS Studio 11 against a custom database containing species-specific transcriptomes, genomes, and NCBI-nr cephalopod sequences. Searches were conducted with precursor and fragment mass tolerances of 10 ppm and 0.02 Da, respectively, allowing semi-specific trypsin digestion with up to two missed cleavages. Carbamidomethylation (C) was set as a fixed modification, while oxidation (M) and deamidation (N/Q) were allowed as variable modifications. Protein identifications were accepted based on a -10lgP score ≥40, presence of at least one unique peptide, and a false discovery rate (FDR) threshold of 1%. All raw mass spectrometry data have been deposited in the ProteomeXchange repository under accession number PXD058861.

### MALDI-Mass Spectrometry Imaging

Tissue sections (6 µm thickness) of venom glands from *D. paeleii, E. berryi*, and *A. bandense* were mounted on a poly-L-lysine-coated (0.05% in MilliQ-water, Sigma Aldrich) conductive glass slide (Bruker Daltonik GmbH, Germany). Before deparaffinization, the slides were preheated for 1 hour at 60 °C. Then, paraffin was removed in xylene and tissue sections were processed through 100% isopropanol and successive hydration steps of 100% ethanol followed by 96%, 70%, and 50% ethanol, each for 5 minutes, as previously described ^47^. Full rehydration was achieved by immersing the sections in ultrapure water (GenPure xCAD Plus System, Thermo Fisher Scientific, USA). Heat-induced antigen retrieval was carried out by steaming tissue sections in ultrapure water for 20 minutes. After air-drying the slides for 10 minutes, tryptic digestion was performed using an automated spraying system (HTX TM-Sprayer, HTX Technologies LLC, ERC GmbH, Germany) to deliver onto each section 16 layers of tryptic solution (20 µg Promega Sequencing Grade Modified Porcine Trypsin in 800 µL digestion buffer including 20 mM ammonium bicarbonate with 0.01% glycerol) at 30 °C. The tissue sections were incubated in a humidity chamber saturated with potassium sulfate solution for 2 hour at 50 °C. Subsequently, four layers of matrix solution (7 g/L α-cyano-4-hydroxycinnamic acid in 70% acetonitrile and 1% trifluoroacetic acid) were applied at 75 °C using the same automated spraying system.

MALDI imaging was performed on an ultrafleXtreme MALDI-ToF/ToF mass spectrometer (Bruker Daltonik GmbH, Germany) operating in reflector mode with a detection range of *m/z* 600–3200, 500 laser shots per spot, 1.25 GS/s sampling rate and raster width of 20 μm. Imaging data acquisition and processing were managed using FlexImaging 5.0 and flexControl 3.4 software (Bruker Daltonik GmbH, Germany). External calibration was performed using a peptide calibration standard (Bruker Daltonik GmbH, Germany). After MALDI imaging analysis, the matrix was removed from the tissue sections with 70% ethanol, followed by hematoxylin and eosin (HE) staining for histological annotation.

The MALDI-MSI raw data were imported into the SCiLS Lab software (Version 2025a Pro, Bruker Daltonik GmbH, Germany) using settings preserving the total ion count, baseline correction, and converted into the SCiLS base data (.sbd) and simulation model (.slx) file. Peak detection and alignment were conducted across a selected dataset (interval width = 0.325 Da) using a standard segmentation pipeline in maximal interval processing mode with total ion count (TIC) normalization and weak denoising.

To match aligned *m/z* values from MALDI-MSI with the peptides identified by nLC-MS/MS, we developed an in-house script with parameter settings as previously described ^47^. Briefly, the comparison of MALDI-MSI and nLC™MS/MS *m/z* values required the identification of > 1 peptide(s) ^48^. In the first dimension, all masses from the LC-MS peptide library and the MALDI-MSI data were matched within an absolute search mass window ≤ 0.2 Da. In the second dimension, the mass from the LC-MS peptide library with the highest converted p-value (-10lgP score) was selected as the correct match (see **Supplementary File 3**).

## Supporting information

Main Figures

## Acknowledgments

The authors acknowledge Dr. Lisa Abbo, Brett Grasse, Taylor Sakmar, the Cephalopod Program, and other members of the Marine Research Center at the Marine Biological Laboratory for providing specimens and guidance in their handling and dissection. The authors also acknowledge Dr. Andrew Gillis for his guidance, training, and sharing of in-situ hybridization reagents and techniques.

## Author contributions

MH designed research; PN, JR, SA, FA, BH, IC, BS, KK, MW, JM, MG, DS, AA, BU performed research; CA, AG contributed new reagents/analytic tools and guidance; PN, JR, SA, FA, MW and KK analyzed data; and MH, PN, JR, SA and FA wrote the paper with edits and contributions from all authors.

## Supporting Information

Supporting information can be found in Supplementary files 1-4.

## Contact and Competing Interest

The authors have declared no competing interest.

## References

1. Schendel, V., Rash, L. D., Jenner, R. A. & Undheim, E. A. B. The Diversity of Venom: The Importance of Behavior and Venom System Morphology in Understanding Its Ecology and Evolution. Toxins 11, 666 (2019).

2. Chung, W.-S., Kurniawan, N. D., Marshall, N. J. & Cortesi, F. Blue-lined octopus Hapalochlaena fasciata males envenomate females to facilitate copulation. Curr. Biol. CB 35, R169–R170 (2025).

3. Holford, M., Daly, M., King, G. F. & Norton, R. S. Venoms to the rescue. Science 361, 842–844 (2018).

4. Verdes, A. et al. From Mollusks to Medicine: A Venomics Approach for the Discovery and Characterization of Therapeutics from Terebridae Peptide Toxins. Toxins 8, 117 (2016).

5. Escoubas, P. & King, G. F. Venomics as a drug discovery platform. Expert Rev. Proteomics 6, 221–224 (2009).

6. Achimba, F., Faezov, B., Cohen, B., Dunbrack, R. & Holford, M. Targeting Dysregulated Ion Channels in Liver Tumors with Venom Peptides. Mol. Cancer Ther. 23, 139–147 (2024).

7. Bordon, K. de C. F. et al. From Animal Poisons and Venoms to Medicines: Achievements, Challenges and Perspectives in Drug Discovery. Front. Pharmacol. 11, (2020).

8. Chang, C. C. Looking back on the discovery of alpha-bungarotoxin. J. Biomed. Sci. 6, 368–375 (1999).

9. D’Angelo, A. et al. Captopril in the treatment of hypertension in type I and type II diabetic patients. Postgrad. Med. J. 62 Suppl 1, 69–72 (1986).

10. Coulter-Parkhill, A., McClean, S., Gault, V. A. & Irwin, N. Therapeutic Potential of Peptides Derived from Animal Venoms: Current Views and Emerging Drugs for Diabetes. Clin. Med. Insights Endocrinol. Diabetes 14, 11795514211006071 (2021).

11. Miljanich, G. P. Ziconotide: neuronal calcium channel blocker for treating severe chronic pain. Curr. Med. Chem. 11, 3029–3040 (2004).

12. Furman, B. L. The development of Byetta (exenatide) from the venom of the Gila monster as an anti-diabetic agent. Toxicon Off. J. Int. Soc. Toxinology 59, 464–471 (2012).

13. Baden, T. et al. Cephalopod-omics: Emerging Fields and Technologies in Cephalopod Biology. Integr. Comp. Biol. 63, 1226–1239 (2023).

14. Hanlon, R. T. & Messenger, J. B. Cephalopod Behaviour. (Cambridge University Press, 2018).

15. Lo Bianco, S. Notizie biologiche riguardanti specialmente il periodo di maturità sessuale degli animali del golfo di Napoli. Mitth. Aus Zool. Stn. Zu Neapal 8, 385–440 (1888).

16. Whitelaw, B. L. et al. Combined Transcriptomic and Proteomic Analysis of the Posterior Salivary Gland from the Southern Blue-Ringed Octopus and the Southern Sand Octopus. J. Proteome Res. 15, 3284–3297 (2016).

17. Fingerhut, L. C. H. W. et al. Shotgun Proteomics Analysis of Saliva and Salivary Gland Tissue from the Common Octopus Octopus vulgaris. J. Proteome Res. 17, 3866–3876 (2018).

18. Undheim, E. a. B. et al. Venom on ice: first insights into Antarctic octopus venoms. Toxicon Off. J. Int. Soc. Toxinology 56, 897–913 (2010).

19. Fry, B. G., Roelants, K. & Norman, J. A. Tentacles of venom: toxic protein convergence in the Kingdom Animalia. J. Mol. Evol. 68, 311–321 (2009).

20. Ueda, A., Nagai, H., Ishida, M., Nagashima, Y. & Shiomi, K. Purification and molecular cloning of SE-cephalotoxin, a novel proteinaceous toxin from the posterior salivary gland of cuttlefish Sepia esculenta. Toxicon 52, 574–581 (2008).

21. Key, L. N., Boyle, P. R. & Jaspars, M. Novel activities of saliva from the octopus Eledone cirrhosa (Mollusca; Cephalopoda). Toxicon Off. J. Int. Soc. Toxinology 40, 677– 683 (2002).

22. Ghiretti, F. Cephalotoxin: the Crab-paralysing Agent of the Posterior Salivary Glands of Cephalopods. Nature 183, 1192–1193 (1959).

23. Ghiretti, F. TOXICITY OF OCTOPUS SALIVA AGAINST CRUSTACEA*. Ann. N. Y. Acad. Sci. 90, 726–741 (2006).

24. Cariello, L. & Zanetti, L. Alpha-and beta-cephalotoxin: two paralysing proteins from posterior salivary glands of Octopus vulgaris. Comp. Biochem. Physiol. C 57, 169–173 (1977).

25. Songdahl, J. H. & Shapiro, B. I. Purification and composition of a toxin from the posterior salivary gland of Octopus dofleini. Toxicon 12, 109–112 (1974).

26. Gonçalves, C., Moutinho Cabral, I., Alves de Matos, A. P., Grosso, A. R. & Costa, P. M. Transcriptome profiling of the posterior salivary glands of the cuttlefish Sepia officinalis from the Portuguese West coast. Front. Mar. Sci. Volume 11-2024, (2024).

27. Boucaud-Camou, E. & Roper, C. F. E. THE DIGESTIVE SYSTEM OF RHYNCHOTEUTI-llON PARALARV AE (CEPHALOPODA: OMMASTREPHIDAE).

28. Anadón, R. Functional Histology: The Tissues of Common Coleoid Cephalopods. in Handbook of Pathogens and Diseases in Cephalopods (eds Gestal, C., Pascual, S., Guerra, Á., Fiorito, G. & Vieites, J. M.) 39–85 (Springer International Publishing, Cham, 2019). doi:10.1007/978-3-030-11330-8_4.

29. Albertin, C. B. et al. Genome and transcriptome mechanisms driving cephalopod evolution. Nat. Commun. 13, 2427 (2022).

30. Gavriouchkina, D. et al. A single-cell atlas of the bobtail squid visual and nervous system highlights molecular principles of convergent evolution. Nat. Ecol. Evol. 9, 1245–1262 (2025).

31. Song, W. et al. Pharaoh Cuttlefish, Sepia pharaonis, Genome Reveals Unique Reflectin Camouflage Gene Set. Front. Mar. Sci. 8, (2021).

32. Rencken, S. et al. Chromosome-scale genome assembly of the European common cuttlefish Sepia officinalis. eLife 14, (2025).

33. Lorig-Roach, R. et al. Phased nanopore assembly with Shasta and modular graph phasing with GFAse. Genome Res. 34, 454–468 (2024).

34. Sanchez, G. et al. The chromosomal genome sequence of the bigfin reef squid, Sepioteuthis lessoniana d’Orbigny, 1826 and its associated microbial metagenome sequences. Wellcome Open Res. 10, 351 (2025).

35. Castelin, M. et al. Macroevolution of venom apparatus innovations in auger snails (Gastropoda; Conoidea; Terebridae). Mol. Phylogenet. Evol. 64, 21–44 (2012).

36. Krause, R. Ueber Bau und Function der hinteren Speicheldrusen der Octopoden. Acad. Wiss. 51, 1085–98 (1897).

37. Jumper, J. et al. Highly accurate protein structure prediction with AlphaFold. Nature 596, 583–589 (2021).

38. Zhang, Y. & Skolnick, J. TM-align: a protein structure alignment algorithm based on the TM-score. Nucleic Acids Res. 33, 2302–2309 (2005).

39. Dutertre, S. et al. Evolution of separate predation-and defence-evoked venoms in carnivorous cone snails. Nat. Commun. 5, 3521 (2014).

40. Allcock, A. L., Lindgren, A. & Strugnell, J. M. The contribution of molecular data to our understanding of cephalopod evolution and systematics: a review. J. Nat. Hist. 49, 1373– 1421 (2015).

41. Edgar, R. C. Muscle5: High-accuracy alignment ensembles enable unbiased assessments of sequence homology and phylogeny. Nat. Commun. 13, 6968 (2022).

42. Lanfear, R., Frandsen, P. B., Wright, A. M., Senfeld, T. & Calcott, B. PartitionFinder 2: New Methods for Selecting Partitioned Models of Evolution for Molecular and Morphological Phylogenetic Analyses. 10.1093/molbev/msw260.

43. Stamatakis, A. RAxML version 8: a tool for phylogenetic analysis and post-analysis of large phylogenies. Bioinformatics 30, 1312–1313 (2014).

44. Choi, H. M. T. et al. Third-generation in situ hybridization chain reaction: multiplexed, quantitative, sensitive, versatile, robust. Dev. Camb. Engl. 145, dev165753 (2018).

45. Criswell, K. E. & Gillis, J. A. Resegmentation is an ancestral feature of the gnathostome vertebral skeleton. eLife 9, e51696 (2020).

46. Ahuja, N. et al. Creation of an albino squid line by CRISPR-Cas9 and its application for in vivo functional imaging of neural activity. Curr. Biol. CB 33, 2774-2783.e5 (2023).

47. Hempel, B.-F. et al. Spatial Venomics-Cobra Venom System Reveals Spatial Differentiation of Snake Toxins by Mass Spectrometry Imaging. J. Proteome Res. 22, 26–35 (2023).

48. Cillero-Pastor, B. & Heeren, R. M. A. Matrix-Assisted Laser Desorption Ionization Mass Spectrometry Imaging for Peptide and Protein Analyses: A Critical Review of On-Tissue Digestion. J. Proteome Res. 13, 325–335 (2014).

