## Supplementary Figures for "Lineage-Specific Venom Gene Expression Shapes Chemical Diversity in Cephalopods"

**A**

```
                            1 ........10........20........30........40........50........60........70........80
CTX1_SEPES 3159 bp          1 ATGA------TGGGG---ACGTCGCGGTGTGTGATCCTTTTGTTTGCGCTGCTTTTGTGGGCGGCCAACGCAGCACCCCC
CTX2_SEPES 3129 bp          1 ATGG------TTCTG---------TGGCAGGTGTTGTTTTTAGTTCCGTTGCTATGGCAATGTGTTCAGGGTTTTTCACC
CTX1_DPEA 3156 bp           1 ATGA------GCGGA---ACTTGGTGGCACGTGCCGTTTCTGTTCCCGCTGCTTTTGCTGGGAGCCAATGGAGCACCTTC
CTX2_DPEA 3123 bp           1 ATGG------TTTTG---------TGGCAAGTGTTGTTCTTATTTCCATTGCTATGGTTAACTATTCGTTGTTTTCCCCT
CTX1_SBAN 2946bp            1 ATGA------TGGCG---GAGTGGCGGTATACGGTCTTTTTGTTCCCGCTGCTTTTGTTGGCGGCCAACGCAGCACCACC
CTX2_SBAN 2286 bp           1 ATGT------TTCTG---------TGGCAGGTGCTGTTTTTAGTCCCGTTGCTATGGCAACATGCTCAGGGTATTTTACC
CTX_EBER 3159 bp            1 ATGA------TGGGGTCATCATGGCCGCGGGTGCTGCTTTTGTTTGTCCTTCCATTGTTGGGGGTCAGTGGAGCCAACGT
CTX2_SOFF 3129 bp           1 ATGG------TTCTG---------TGGCAGAGGTTGTTTTTCGTTCCATTGCTATGGCAATGTGTTCAGGGTGTTTTACC
CTX1_SOFF 3141bp            1 ATGA------AGGAG---GCGTGGCGGTATGTGATCCTATTTTTGGCGGTGCTTTTGTTGGCGGCCAACGGAGCACCCCT
CTX_ESCO 3159 bp            1 ATGA------TGGGGTCATCATGGCCGTGGGTGCTGTTTTTATTTGCCCTTCCGTTGTTGGGGATCAGTGGAGCAACCGT
CTX2_SLES 3132 bp           1 ATGG------TTCTG---------TGGCAAGTGTGGTTTTTATTTTCATTGCTATGGTTGGCTGTTCGCGGTTTTCCACC
CTX1_SLES 3159 bp           1 ATGA------TAGAA---GCAAAGTGGCACGTGTCGCTTCTGTTGCTGCTCCTTTTATTGGGAGCCAATGAAGCATCTTC
CTX_TRHO 3150 bp            1 ATGG------TTCTG---------TGGCAGGCGTTGTTTTTACTTCCGATGCTATGGCTTAGCGTTCACGGTTCTTCATC
CTX_SATL 3147 bp            1 ATGG------GGTCG---TCATGGCCGCGAGTGCTGCTTTTCTTTGCGCTTCTGTCGTTGGGGATCGGTGGAGCAACCGT
CTX2_SLYC 3129 bp           1 ATGG------TTCTG---------TGGCAGGTGTTGTTTTTAGTTCCGTTGCTATGTCAATGTGTTCAGGGTTTTTCACC
CTX1_SLYC 3159 bp           1 ATGA------TGGGG---ACGTCGTTGTGTGTGATCCTTTTGTTTGCGCTGCTTTTGTGGGCAGCCAACGCAGCACCCCC
CTX_EPAR 3150 bp            1 ATGA------TGGGGTCATCATGGCCGCGGGTGCTGCTTTTGTGTGCCCTTCCGTTGTTGGTGGTCAGTGGAGCAACCGT
CTX2_SPHAR 3129 bp          1 ATGG------TTCAG---------TGGCAGGTGTTGTTTTTAGTTCCGTTGCTATGGCAATGTGTTCAGGGTTTTTCACC
CTX1_SPHAR 3159 bp          1 ATGA------TGGGG---ACGTCGCGGTGTGTGATCCTTTTGTTTACGCTGCTTTTGTGGGCGGCCAACGCAGCACCCCC
CTX_WSCI_PARTIAL 1653 bp    1 --------------------------------------------------------------------------------
CTX_IHAL 3156 bp            1 ATGG------CGACA---GAGTGGCAGCTGATATTTTTTACAGTTGCTTTACTAATTCAGGGCAACAATGGCGCCCCTAC
CTX2_XNOT 3183 bp           1 ATGA------CTTTG---------TGGCAGGCATTATTTCTACATTTATTTTTATTGATATGTGCTACAGTTGTTTCATC
CTX1_XNOT2 3150 bp          1 ATGG------TGGCA---GAGTGGCAGCTGATATTTGTTACAGTTGCTCTGGTTTTCCAAGGGACCAATGGCGCCCCTGA
CTX_OROB_PARTIAL 1431 bp    1 --------------------------------------------------------------------------------
CTX_SAFF 3147 bp            1 ATGG------GGTCG---TCATGGCCGCGAGTGCTGCTTTTCTTTGCGCTTCTGTCGTTGGGGATCGGTGGAGCAACCGT
CTX_SOBS 3156bp             1 ATGG------GGTCG---TCATGGCCGCGACTGCTGCTTTTGTTTGTCCTTCTGTCGTTGGGCATCAGTGGAGAAACCGC
CTX_Sarc 2907 bp            1 --------------------------------------------------------------------------------
CTX2_Dopa 3123 bp           1 ATGG------TTTTG---------TGGCAAGTGTTGTTCGTATTTCCATTACTATGGTTAACTATTCGTTGTTTTCCCCT
CTX1_Dopa 3123 bp           1 ATGAAAATGAGAGGA---ACTTGGTGGCACGTGCCGCTTCTGTTCCCGCTGCTTTTGCTGGGAGCCAATGGAGCACCTTC

                           81 ........90.......100.......110.......120.......130.......140.......150.......160
CTX1_SEPES 3159 bp         72 AGAGATTCATACCACGAG---ACC---------------------AAATGT------TCCTGAAGAAATAA---AAAGA-
CTX2_SEPES 3129 bp         66 AGACGACAATGCTACTTC---------------------------------------TGGTGATCACCCAG---TAGCA-
CTX1_DPEA 3156 bp          72 AGAGATTACCAC---TAA---ATT---------------------TCCGTT------ACTTGAAGAAAACA---TACAC-
CTX2_DPEA 3123 bp          66 GGAAGACAATGCTGCTGT------------------------------TGT------TTGTGATCAAGCGC---AAGTA-
CTX1_SBAN 2946bp           72 CGAGAGTCACACCACTAG---ACC---------------------AAAAGT------TCCTGAAGAA---T---ATCAA-
CTX2_SBAN 2286 bp          66 AGATGACGGTGCGACTTC---------------------------------------TCATGATCAACTCG---TAGCC-
CTX_EBER 3159 bp           75 AATGGTTAATAATACTGA---ACT---------------------CGACAT------TATTCAAATGATGA---ACACA-
CTX2_SOFF 3129 bp          66 AGACGACGGGGCTACTTC---------------------------------------TCGTGATCACCCAG---TCGCA-
CTX1_SOFF 3141bp           72 AGAGAGTACTACCTCGGA---ACC---------------------A------------------------G---AAGAA-
CTX_ESCO 3159 bp           75 AATGGTTAATAATACTGA---ACT---------------------CGACAT------TATTCAAATGATGA---ACACA-
CTX2_SLES 3132 bp          66 GGAAGAAAATACTGCTAT------------------------------TGT------TTGCAATCAATTGG---AAGCA-
CTX1_SLES 3159 bp          72 AGATATTATTGCCACTAA---ATT---------------------TCCGTT------TCTTGAAAAGAACA---CACAT-
CTX_TRHO 3150 bp           66 AGATGACAGTACCGCAGC---TGG------------------------TGT------CAGTAGTCAGATGG---TAGCA-
CTX_SATL 3147 bp           72 AGCGGTTGACAATAGTCA---AGT---------------------CGACAT------TCTTGGAATGATCA---ACACG-
CTX2_SLYC 3129 bp          66 AGACGACATTGCTATTTC---------------------------------------CCGTGATCACCCAG---TAGCA-
CTX1_SLYC 3159 bp          72 AGAGAGTCATACCACGAG---ACC---------------------AAAAGT------TCCTGAAGAAATAA---AAAGA-
CTX_EPAR 3150 bp           75 GCTCG------ATACTGA---ACT---------------------CGACAT------TTTTCAACTGATGA---ACACA-
CTX2_SPHAR 3129 bp         66 AGACGACATTACTACTTC---------------------------------------CCGTGATCACCCAG---TAGCA-
CTX1_SPHAR 3159 bp         72 AGAGAGCCATACCACGAG---ACC---------------------AAAAAT------TCCTGAAGAAATAA---AAAGA-
CTX_WSCI_PARTIAL 1653 bp    1 --------------------------------------------------------------------------------
CTX_IHAL 3156 bp           72 AGACACTAATGTCAGCCA---AGT---------------------CATCGT------ACCAAATGTCCCAA---TGGAC-
CTX2_XNOT 3183 bp          66 GCAAGATTGTGATGCGCCAAACAGTTACACTAGTGATATAGATCCTAGCATTAATCATAGTGATCATCAATTAAATGGGA
CTX1_XNOT2 3150 bp         72 AAACACTAATGTCAGCCA---AAT---------------------AGTAGT------ACCAAATGTACCTA---CGGAA-
CTX_OROB_PARTIAL 1431 bp    1 --------------------------------------------------------------------------------
CTX_SAFF 3147 bp           72 AGCGGTTGACAATAGTCA---AGT---------------------CGACAT------TCTTGGAATGATCA---ACACG-
CTX_SOBS 3156bp            72 AGAGATTGACAATAGTCA---AGT---------------------CCACAT------TTTTGGAATGACCA---ACACA-
CTX_Sarc 2907 bp            1 --------------------------------------------------------------------------------
CTX2_Dopa 3123 bp          66 GGAAGACAATGCTGCTGT------------------------------TGT------TTATGATCAAGCGC---AAGTA-
CTX1_Dopa 3123 bp          78 AGAGATTACCAC---TAA---ATT---------------------TCCGTT------ACTTGAAGAAAACA---TATAC-

                          161 .......170.......180.......190.......200.......210.......220.......230.......240
CTX1_SEPES 3159 bp        118 --------CCAAATTCAACCGAAA---TAGAAAC---TCCTGCAGTGA---------------AACAATTGGAAACGCCA
CTX2_SEPES 3129 bp        103 --------ATGGACGAAACGAAAAAAGACAATCA---AACTGAAAACC---------------AGACCCCTCAGTTTCCA
CTX1_DPEA 3156 bp         115 --------TTCAATTCAACCGAAC---CTAAGCCTACTGCTGCACTAA---------------AAGATTTAGACCCTCCA
CTX2_DPEA 3123 bp         106 --------ATGAATCAAACTGGAT---GTAAACC---TTCTGAAAGCA---------------ATACTCCTGATTTCCCA
CTX1_SBAN 2946bp          115 --------ACAAATTCAACCGAAA---TAATAAC---TCCTGCTGTGA---------------AACAGTTGGAAACTCCA
CTX2_SBAN 2286 bp         103 --------ATGAACGAAACGAAAAAAGACGACCA---AACTAAAAATC---------------AGACCCCCCAGATTCCA
CTX_EBER 3159 bp          121 --------ACGAGTTCAACGGAAG---TCGAAAC---CCCATTGGTGA---------------AACAATTTGAAACTCCG
CTX2_SOFF 3129 bp         103 --------ATGAACGAAACGAAAAAAGAGGAAGT---AACTGAAAAGC---------------AGACCCCTCAGTTTCCT
CTX1_SOFF 3141bp          100 --------ACCAACTCAACCGTAA---TTGCAAC---TCCTGCCGTGA---------------AAATAATAGAAACGCCA
CTX_ESCO 3159 bp          121 --------ACGAGTTCAACAGAAG---TCGAAAC---CCCCTTGGTTA---------------AACAATTTGAAACTCCG
CTX2_SLES 3132 bp         106 --------ATGAATCAGACTGGAT---GTAAATC---TCCTGAACACA---------------ATAATTCTAATTTACCA
CTX1_SLES 3159 bp         118 --------CTCAATTCAACTGAAA---CAAAACCTATTCCTAAAGTGA---------------AGGATTTAGAACCTCCA
CTX_TRHO 3150 bp          109 --------ATGAACGAAACAGAAAAGGTCCCTGG---AACCGAAAGTGACCGCCCAGCGAACCGTTCCTCTGAACTACCA
CTX_SATL 3147 bp          118 --------ACGGGTCCAACCGAAG---TTGAAAC---CCCCGTCGTGA---------------AACAAATGGAAACACCG
CTX2_SLYC 3129 bp         103 --------ATAGATGAAACGAAAAAAGACAATCA---AACTGAAACCC---------------AGACCCCTCAGTTTCCA
CTX1_SLYC 3159 bp         118 --------CCAAATTCAACCGAAA---TAGTAAC---TCCTGCAGTGA---------------AAACATTGGAAACGCCA
CTX_EPAR 3150 bp          115 --------ACAAGTTCAACGGAAG---TCGAAAC---CCCCTTGGTGA---------------AACAATTTGAAACTCCG
CTX2_SPHAR 3129 bp        103 --------ATGGACGAAACGAAAAAAGACAATCA---AACTGAAACCC---------------AGACGCCACAGTTTCCA
CTX1_SPHAR 3159 bp        118 --------CCAAATTCAACCGAAA---TAGTAAC---TCCTGCAGTGA---------------AACAATTGGAAACGCCA
CTX_WSCI_PARTIAL 1653 bp    1 --------------------------------------------------------------------------------
CTX_IHAL 3156 bp          118 --------AACAACTCAACAGAAA---TAAAAAC---CCCTGTCGTCA---------------AAGAAATGGAATCACCC
CTX2_XNOT 3183 bp         146 CAGATGTTGGGAATACAACTTTA---TGCAAAGA---AACCGAAATTA---------------AGAGTTCTGGTTTACCA
CTX1_XNOT2 3150 bp        118 --------TTCAACTCAACTGAAA---TAAAAAC---TCCAGTTGTGA---------------AAGAAATGGAATCCCCT
CTX_OROB_PARTIAL 1431 bp    1 --------------------------------------------------------------------------------
CTX_SAFF 3147 bp          118 --------ACGGGTCCAACCGAAG---TTGAAAC---CCCCATTGTGA---------------AACAAATGGAAACGCCG
CTX_SOBS 3156bp           118 --------ACGGGTCCAACCGAAG---TTGAAAC---CCCCACTGTGA---------------AACAACTTGAGACGCCG
CTX_Sarc 2907 bp            1 --------------------------------------------------------------------------------
CTX2_Dopa 3123 bp         106 --------ATGAATCAAACTGGAT---ATAAACC---TCCTGAAAGGA---------------ATACTCCTGAGTTCCCA
CTX1_Dopa 3123 bp         121 --------CTCAATTCAACCGAGC---CTAAGCCTACAGCAGCACTAA---------------AAGATTTAGAACCTCCA

                          241 .......250.......260.......270.......280.......290.......300.......310.......320
CTX1_SEPES 3159 bp        169 TCGATTTTCCTTCTCACCACCCTCGAAGTGGCAGAAGCGGACGTTGATAGCACATTAGAGACGATGAAGGACAGAAA---
CTX2_SEPES 3129 bp        157 ACGGTTATGATAATGACGACTCTTGAAGTTGTGAAGTCTGAAATAGAAGATGTTTTGAATTATGAAGGCGAAGCCGG---
CTX1_DPEA 3156 bp         169 CTAATTTTCCTTCAAACAAGTCTGGAAGTGGCAGAAGCGGATTTAGATAGAAATATAGAGACTATAAAAGAGAAAAG---
CTX2_DPEA 3123 bp         157 ATGACCATGATAATGACAACTCTTGAAATGGTGCAGACGGAGATTGGGGAACTTCAGAATTTTGAAACCGAACAACT---
CTX1_SBAN 2946bp          166 ACAATTTTCCTTCTCACCACCATGGAAGTGATAGAAGCGGATGTTGACACTATGTTAGAGGCGATGAAAGACAAAAA---
CTX2_SBAN 2286 bp         157 ACGGCTATGATAATGACGACGCTTGAAGTTGTGAAGTCTGAAATAGATGATGCTTTGAATTATGAAGGCGAAGCAGG---
CTX_EBER 3159 bp          172 TCAATTTTCCTGTTGACGACCCTCGAAGAGGTCGATGCGGACTTCGATAAACGCTTGGAAACGATCAAAGACAAAAA---
CTX2_SOFF 3129 bp         157 ACGGCTATGATAATGACGACGCTTGAAGTTGTGAGGTCTGAAATAGAAGATGTTTTGAATTATGAAGGCGAAGCGGT---
CTX1_SOFF 3141bp          151 TCGATATTCCTTCTCACCACTCTTGAAGTGGCGGAAGCGGAAGTAGATAAATCGTTAGAGTCGATGAAGAGCAGAAA---
CTX_ESCO 3159 bp          172 TCAATTTTCTTGTTGACGACCCTCGAAGAGGTCAATACGGACTTCGATAAACGCTTGGAAACGATCAAAGACAAAAA---
CTX2_SLES 3132 bp         157 ATGGTCATGATAATGACGACTCTTGAAGTGGTGCAGTCAGAGATTGGAGAACTTCAGAATGAGGAAACTGAAAAAGA---
CTX1_SLES 3159 bp         172 TTAATTTTCCTTCAAACGGGTCTGGAAGTGGCAGAAGCGGATTTAGATAAAAACCTTGAAACAATAAAAGAGAAAAA---
CTX_TRHO 3150 bp          178 TCGATCATTATAATGACAACTCTTGAAGTGATGCAGTCGGAGATTGAAAAAACCCAGAATGATGGAACCGAGAAAAG---
CTX_SATL 3147 bp          169 TCAATTTTCCTGCTGACGACCCTCGAAGAGGTCGACACTGATTTTGATAAACGCTTGGAAACAATCAAAGACAAAAA---
CTX2_SLYC 3129 bp         157 ACACTTATGATAATGACGACTCTTGAAGTTGTGAAATCTGAAATAGAAGATTTTTTGAATTACGAAGGCGAAGCAGG---
CTX1_SLYC 3159 bp         169 ACGATTTTCCTTCTCACCACCCTCGAAGTGACAGAAGCTGACGTTGATAGCACATTAGAGGCGATGAAGGACAAAAA---
CTX_EPAR 3150 bp          166 TCAATTTTCTTGTTGACGGCCCTCGAAGAAGTCAATACAGATTTTGATAAACGCTTAGATACGATCAAAGACAAAAA---
CTX2_SPHAR 3129 bp        157 ACAGTCATGATAATGACGACTCTTGAAGTTGTAAAATCTGAAATAGAAGATGTTTTGAATTACGAAGGCGAAGCAGG---
CTX1_SPHAR 3159 bp        169 ACGATTTTCCTTCTCACCACCCTCGAAGTGGCAGAAGCTGACGTTGATAGCACATTAGAGTCCATGAAGGAGAAAAA---
CTX_WSCI_PARTIAL 1653 bp    1 --------------------------------------------------------------------------------
CTX_IHAL 3156 bp          169 ACAATTTTTCTCATGACGAGTCTCGAAGTTGCTGAATCGGATTTTGAAAAGAAAATGGATTTAGCCAGTAACAAAGT---
CTX2_XNOT 3183 bp         205 ACTGTCGTAATTATAACCAGTTTGGAAGTGATACAAGTCGAAATTGATTCCTATCAGGATGAAGACAACAATAAAGATAG
CTX1_XNOT2 3150 bp        169 ACCATTTTTCTCATGACGAGTCTTGAAGTTGCTGAGTCAGATTTCGAAAAGAAAATGGACTCAGCAAGTAACAAAGT---
CTX_OROB_PARTIAL 1431 bp    1 --------------------------------------------------------------------------------
CTX_SAFF 3147 bp          169 TCAATTTTCCTGCTGACGACCCTCGAAGAGGTCGACACCGATTTTGATAAACGCTTGGAAACAATCAAAGACAAAAA---
CTX_SOBS 3156bp           169 TCAATTTTCCTGTTGACCAGCCTCGAAGAGATCGACACCGATTTCGATAAACGCTTGGAAACACTCAAAGACAAAAC---
CTX_Sarc 2907 bp            1 --------------------------------------------------------------------------------
CTX2_Dopa 3123 bp         157 ATGACCATGATAATGACAACTCTTGAAATGGTACAGACGGAGATTGGGGAACTTCAGAATTTTGAAACCGAACAACT---
CTX1_Dopa 3123 bp         175 CTAATTTTCCTTCAAACAAGTCTGGAAGTGACAGAAGCGGATTTAGATAAAAATATAGAGGCTATAAAAGAGAAAAG---

                          321 .......330.......340.......350.......360.......370.......380.......390.......400
CTX1_SEPES 3159 bp        246 ---TAAAAAGAATTCAGCTAAATTGTCTAAAATTGGTAATAATATGAAGTCACTTCTTAGTGTCTTTAGTGTATTTGGTG
CTX2_SEPES 3129 bp        234 ---AAACAAATTTTTCAGCATAAAGAAAAAACATGCAAACAGTCTAAAAATCTTCAACTCTCTCTTCAATGCACTTGGAG
CTX1_DPEA 3156 bp         246 ---TAAAAATCCTGCTGCTAAAGTATCCAAATTTGGTCATGGTGTGAAAAAAATTTGTTCTATGTTTAACGTTTTTGGTG
CTX2_DPEA 3123 bp         234 ---AACCAATGATTTTGGCAAAAA---------GTCAAAGACTTTAAAAGTCTTCAGTTCTGTGTTCAACGCATTAGGAG
CTX1_SBAN 2946bp          243 ---CAAAAAGAAGGCAGCTAAATTGACCAAATTTGGTAATGATATGAAGTCACTTCTCGGAGTCTTCAGTGTAATGAGTG
CTX2_SBAN 2286 bp         234 ---AACCAAGATTTTTAGAATAACGTCAAAACATGCGAAGACTTTAAAAATCTTCAACTCTTTCTTGAATGCATTTGGGG
CTX_EBER 3159 bp          249 ---CAAAAAGCCGCCTGCCAAAATGTCTTCAATTGGCAACAGTATGAAAAAGTTTTGCTCTATGTTTAGTGTAATGAGCG
CTX2_SOFF 3129 bp         234 ---AAACAATTTTTTAAGTATAAGCAAAAAACATGCCAAGACCCTAAAAATCTTCAACTCTCTCTTCACTGCATTCGGGG
CTX1_SOFF 3141bp          228 ---TAAAAAGAACGCAGCTCAATTTGACAAAATTGGCAAGAACGTCAAGTCACTTCTTAGTGTCTTTAGTGTATTTAGCG
CTX_ESCO 3159 bp          249 ---CAAAAAGTCGCCTGCCAAAATATCTTCAATTGGCAACAGTATGAAAAAGTTTTGCTCTATGTTTAGTGTAATGAGCG
CTX2_SLES 3132 bp         234 ---AAGCAATAGTTTGAGTAAAAAGACAAAAAAATCAAAGATTTTAAAGATCTTCAGTTCTTTGTTCAATGCATTCGGCG
CTX1_SLES 3159 bp         249 ---GAAAAATCCTATAGCTAAAATATCCAGATTTGGCAATGGGGTAAGACTTTTTTGTTCTATATTTAGTATTACTAGTG
CTX_TRHO 3150 bp          255 ---AATCAATGGTTTGAGCAAAGGAATTAAACACGCGAAGATTTTAAAAGTCTTCTGTTCCCTGTTCAATGCATTCGGGG
CTX_SATL 3147 bp          246 ---TAAAAAGACGCCCGCCAAAATGACCTCATTTGGCAACAGTATGAAAAAGTTTTGCTCTATGTTTAGTGTGATGAGCG
CTX2_SLYC 3129 bp         234 ---AAACTCATTTTTCAGCGTAAAGAAAAAACATGCAAAGACTCTAAAAATCTTCAACTCTCTCTTCAATGCACTTGGAG
CTX1_SLYC 3159 bp         246 ---TAAAAAGAATTCAGCTAAATTGTCTAAAATTGGTAATAATATGAAGTCGATTCTTAGTGTCTTTAGTGTAATGAGTG
CTX_EPAR 3150 bp          243 ---CAAAAAGCCGCCTGCCAAAATGTCTTCAATTGGCAACACTATGAAAAAGTTCTGCTCTATGTTTAGTGTAATGAGCG
CTX2_SPHAR 3129 bp        234 ---AAACAAATTTTTCAGCGTAAAGAAAAAACATGCAAAGACTCTAAAAATCTTCAACTCTCTCTTCAATGCACTTGGAG
CTX1_SPHAR 3159 bp        246 ---TAAAAAGAATTCAGCTAAATTGTCTAAACTTGGTAATAATATGAAGTCGCTTCTTAGTGTCTTTAGTGTAATGAGTG
CTX_WSCI_PARTIAL 1653 bp    1 --------------------------------------------------------------------------------
CTX_IHAL 3156 bp          246 ---GAAGAAGAATCCGATGAAATTAACGAAAATTGGAAACAGTCTTCGAAGATTCACTTCCATGTTTAGCATGTTGGGCG
CTX2_XNOT 3183 bp         285 CAAGAAAAATAAACCCAAGCGAACCCCGATACTTTCTAAAAGCGTCAAAATGTTCAGCTCCTTTTTGAATTCTATGGGAG
CTX1_XNOT2 3150 bp        246 ---GAAAAAGAATCCGGTGAAACTAACGAAATTTGGAAATAGTCTTAGAAGATTCACCTCAATGTTCAGTATGTTGGGTG
CTX_OROB_PARTIAL 1431 bp    1 --------------------------------------------------------------------------------
CTX_SAFF 3147 bp          246 ---TAAAAAGACGCCCGCCAAAATGACCTCATTTGGTAACAGTATGAAAAAGTTTTGCTCTATGTTTAGTGTGATGAGCG
CTX_SOBS 3156bp           246 ---TAAAAAGACGTCTTCCAAAATGACCTCATTGGGCAACGGCCTGAAAAAGTTTTGCTCCATGTTTAGTGTGATGAGCG
CTX_Sarc 2907 bp            1 --------------------------------------------------------------------------------
CTX2_Dopa 3123 bp         234 ---AACCAATGATTTTGGCAAAAA---------GTCAAAGACTTTAACTGTCTTCAGTTCTGTGTTCAACGCATTAGGAG
CTX1_Dopa 3123 bp         252 ---TAAAAATCCTGCTGCTAAAGTATCCAAATTTGGTCATGGTGTGAAAAAGATTTGTTCTATGTTTAGCGTTTTTGGTG

                          401 .......410.......420.......430.......440.......450.......460.......470.......480
CTX1_SEPES 3159 bp        323 GTTTCTTAAGCCTCTTATCTGTCGTCACAACAACCTCAGATCTGCAAGTCATTTCCGACATGTTTACCGGAGTTAACAGA
CTX2_SEPES 3129 bp        311 GATTTCTGACTGTGCTGTCTGCCTTTACGCAAACAAGCGATTTAGAAGTCATTACTGCCATGTTCAAAGAAGTGAACAAA
CTX1_DPEA 3156 bp         323 GTTTCTTTAGCTTATTAGCCACTGTCACATCATCATCAGACCTGAAGGTTATTACGGGTATGTTTGATGAAGTCAACAAA
CTX2_DPEA 3123 bp         302 GATTTTTAACTTTATTGTCTGTTTTTACAGGAACCGATGATTTAGAAGTCATTTCTTCCATGTTCAAACAAGTGAATAAA
CTX1_SBAN 2946bp          320 GCTTCTTAAGTTTTTTATCAGTCATTACAACAACCTCAGATCTGCAAGTCATTTCCGACATGTTTACCGGTGTTAACAGA
CTX2_SBAN 2286 bp         311 GATATCTATCCCTGTTGTCTGCCTTTACACAAACTAGCGATCTAGAAGCTATTACATCCATGTTCAAAGAAGTGAACAAA
CTX_EBER 3159 bp          326 GCTTTTTGAGTTTGCTGTCTGTGGTCTCAACCACATCCGATTTGCAGGCCATTACGGATATGTTCGACGGCGTGAACAAA
CTX2_SOFF 3129 bp         311 GATTTCTGTCTGTGTTGTCTACCTTTACACAAACCAGTGATTTGGAAGTCATAACGGGGATGTTCAAAGAAGTGAACAAA
CTX1_SOFF 3141bp          305 GCTTCTTGAGCCTCTTATCCGTCGTCACAACAACGTCAGATCTGCAGGTCATTTCAGACATGTTTACCGGAGTTAACAAA
CTX_ESCO 3159 bp          326 GCTTTTTGAGTTTGCTGTCTGTGGTCTCAACCACATCCGATTTGCAGGCCATTACGGATATGTTCGACGGCGTGAACAAA
CTX2_SLES 3132 bp         311 GATTTTTAAATTTGCTGTCTGTCTTTACGGGAACCGATGATTTGGAAGTCATTTCTTCCATGTTCAAAGAAGTGAACAAA
CTX1_SLES 3159 bp         326 GCTTTTTTAGTTTATTAGCTACTGTCACATCATCATCAGACTTACAGGTCATTTCCGACATGTTTACCGAAGTCAACAAA
CTX_TRHO 3150 bp          332 GATTCCTGACTGTGTTGTCTGCCTTGACGGGAACCAGTGATTTGGAGGTCATTTCGTCCATGTTCGAAGAAGTGAACAAA
CTX_SATL 3147 bp          323 GCTTTTTGAGTCTGTTGTCTGTGGTCTCAACCACATCCGATCTGCAGGCCATTACAGATATGTTCGACGGCGTGAACAAA
CTX2_SLYC 3129 bp         311 GTTTTCTAACTGTGCTGTCTGCCTTTACGCAAACTAGCGATTTAGAAGTCATTACAGCTATGTTTAAAGAAGTGAACAAA
CTX1_SLYC 3159 bp         323 GCTTCTTAAGCCTCTTATCAGTCATAACAACAACCTCAGATCTGCAAGTCATTTCCGATATGTTTACCGGTGTTAACAGA
CTX_EPAR 3150 bp          320 GCTTTTTGAGTTTGTTGTCTGTGGTCACAACCACATCCGATTTGCAGGCCATTACGGATATGTTCGATGGCGTGAACAAA
CTX2_SPHAR 3129 bp        311 GTTTTCTAACTGTGCTGTCTGCCTTTACGCAAACTAGCGATTTAGAAGTCATTACAGCCATGTTCAAAGAAGTGAACAAA
CTX1_SPHAR 3159 bp        323 GCTTCTTAAGCCTCTTATCAGTCATAACAACAACCTCAGATCTGCAAGTCATTTCCGACATGTTTACCGGAGTTAACAGA
CTX_WSCI_PARTIAL 1653 bp    1 --------------------------------------------------------------------------------
CTX_IHAL 3156 bp          323 GCTTTTTGTCCATGTTGGCCGTGGTGACCACAACGACCGACTTACAGGTCATTTCCGACATGTTTGGAGAGGTCAACAAG
CTX2_XNOT 3183 bp         365 GCTTTCTATCTTTGCTGTCTGTCTTCACAGATACTTCTGATCTGGAGGTCATTTCGGGTATGTTTAAAGAAGTGCACAAA
CTX1_XNOT2 3150 bp        323 GGTTTTTGTCCATGCTCGCCGTGATGACTACAACTACTGATTTGCAAGTCATTTCCGACATGTTTGGAGAAGTAAACAAG
CTX_OROB_PARTIAL 1431 bp    1 --------------------------------------------------------------------------------
CTX_SAFF 3147 bp          323 GCTTTTTGAGTCTGTTGTCTGTGGTCTCAACCACATCCGATCTGCAGGCCATTACAGATATGTTCGACGGCGTGAACAAA
CTX_SOBS 3156bp           323 GCTTCTTGAGTCTGTTGTCTGTGGTCTCAACCACGTCCGATCTGCAAGCCATTACGGATATGTTTCACGGCGTGAACAAA
CTX_Sarc 2907 bp            1 --------------------------------------------------------------------------------
CTX2_Dopa 3123 bp         302 GATTTTTAACTTTATTGTCTGTTTTTACAGAAACCGATGATTTAGAAGTCATTTCTTCCATGTTCAAACAAGTGAATAAA
CTX1_Dopa 3123 bp         329 GTTTCTTTAGCTTATTAGCTGCTGTCACATCATCATCAGACCTGACGGTTATTACGGGTATGTTTGATAAAGTCAACAAA

                          481 .......490.......500.......510.......520.......530.......540.......550.......560
CTX1_SEPES 3159 bp        403 AAACTGGACCAAATAAACGATAAATTAGACAAGTTAGACAACTCTGTTGAGCTTCAGGGACTGTTGACGAATTACATTCC
CTX2_SEPES 3129 bp        391 AAACTGGATAGAATTACTCGACGTATCGATAATCTGGAAAACTCAATCGAATTGCAGAGACTTTTGTCGAATTATATTCC
CTX1_DPEA 3156 bp         403 AAATTAAACAAAATAACTGATCAATTAAAAAAATTGGACAATTCGGTTCAACTAGAGGGACTCTTGTCTAATTACATTCC
CTX2_DPEA 3123 bp         382 AAGCTAGATAAAATATCACAGCAAATAGACAATGTAGAAAATACAGTAGAATTGCAGGGACTTTTGTCCAATTACATCCC
CTX1_SBAN 2946bp          400 AAACTAGACCAAATAAACGATAAATTAGACAAATTAGACAATTCTGTTGAACTTCACAGTCTTTTGTCGAATTTCATTCC
CTX2_SBAN 2286 bp         391 AAACTGGATAGAATTACTCAACGTATAGATAATCTGGGAAATACAGTCAAACTGCAGGGACTTTTGTCGAATTATATTCC
CTX_EBER 3159 bp          406 AAGCTTGACGACATAAGCGATCGGTTGGGCCGGTTGGACGATAAGGTCGAACTGCAAACACTCTTGTCGAATTATATGCC
CTX2_SOFF 3129 bp         391 AAACTGGACAGAATTAATCGACGTATAGACAGTCTGGGAAACACAGTCGAATTGCAGAGACATTTGTCCCATTATATCCC
CTX1_SOFF 3141bp          385 AAACTGGACCAAATCAACGACAAATTAGACAAGTTGGACAACTCTGTTGAGCTTCAGGGACTGTTATCGAATTACATTCC
CTX_ESCO 3159 bp          406 AAGCTTGACGACATAAGCGATCGGTTGGGCCGGTTGGATGATAAGGTCGAACTGCAAGCACTCTTGTCGAATTATATGCC
CTX2_SLES 3132 bp         391 AGACTTGACAAAATTGCACTGCAAATAGACAATGTTGAAAATACAGTAGAATTACAGGGACTTTTGTCCAATTACATACC
CTX1_SLES 3159 bp         406 AAACTAGACAAAATAACCGACCGATTAGACAAATTGGACAATTCAATTGAACTACACGGACTCTTGTCTAATTACATACC
CTX_TRHO 3150 bp          412 AAACTGGACAATATTGCCCGGCGAATAGACAATGTCGAAAGTGCAGTCGAAATGCAGGGACTTTTATCAAATTACATTCC
CTX_SATL 3147 bp          403 AAGCTGGATGACATAAGCGATCGGTTGGGCCGGTTGGATGATTCGGTCGAACTGCAAGGACTCTTGTCGAGTTATATCCC
CTX2_SLYC 3129 bp         391 AAACTGGATAGAATTACTCGACGTATAGATAATCTGGAAAACTCAGTCGAATTGCAGAGACTTTTGTCGAATTATATTCC
CTX1_SLYC 3159 bp         403 AAACTGGACCAAATAAACGATAAATTAGATAAATTAGACAACTCTGTTGAGCTTCAGGGACTGTTGTCGAATTACATTCC
CTX_EPAR 3150 bp          400 AAGCTGGACGACATAAGCGATCGGTTGGGCCGATTGGACGATAAGGTCGAACTGCAAACACTTTTATCGAATTATATGCC
CTX2_SPHAR 3129 bp        391 AAACTAGATAGAATTACTCGACGTATAGATAATCTGGAAAACTCAATCGAATTGCAGAGACTTTTGTCGAATTATATTCC
CTX1_SPHAR 3159 bp        403 AAACTGGACCAAATAAACGATAAATTAGACAAGTTAGACAACTCTGTTGAGCTTCAGGGACTGTTATCGAATTACATTCC
CTX_WSCI_PARTIAL 1653 bp    1 --------------------------------------------------------------------------------
CTX_IHAL 3156 bp          403 AAACTCGATAAAATCAACGATAGACTCGTTAAGCTCGATAACAGCGTTGAATTGCAGAGTCTATTGTCGAATTACATCCC
CTX2_XNOT 3183 bp         445 AAACTTGATAAAATATACAACCGTATTGACAACTTAAAAGGTTCGATTGAACTGCAGGGCCTTCTATCAAATTATATCCC
CTX1_XNOT2 3150 bp        403 AAACTCGATAAAATCAACGATCAACTTATTAAACTTGATGACAGTGTGGCATTGCAGAGTCTATTGTCAAATTATATCCC
CTX_OROB_PARTIAL 1431 bp    1 ------------------------------------------------------------CTGTTGTCCAATTACATTCC
CTX_SAFF 3147 bp          403 AAGCTGGATGACATAAGCGATCGGTTGGGCCGGTTGGATGATTCGGTCGAACTGCAAGGACTCTTGTCGAGTTATATCCC
CTX_SOBS 3156bp           403 AAGCTGGACGACATAAGCGATCGTCTGGGCCGGTTGGATGATTCGGTCGAACTGCAAGGACTCTTGTCGAGTTATATCCC
CTX_Sarc 2907 bp            1 --------------------------------------------------------------------------------
CTX2_Dopa 3123 bp         382 AAACTAGACAAAATTTCACAGCAAATAGACAATGTAGAAAATACAGTAGAATTGCAGGGACTTTTGTCCAATTACATCCC
CTX1_Dopa 3123 bp         409 AAATTAGACAAAATAACTCATCAATTAGAAAAATTGGACAATTCGGTTCAACTACAGGGACTCTTGTCTAATTACATTCC

                          561 .......570.......580.......590.......600.......610.......620.......630.......640
CTX1_SEPES 3159 bp        483 TTGGCAATATTCTGTGAAAAATGGAATCGAAAAACTGATTGAAACCTACAAGAAAATGGTGGAAGAAACAAATATGAATA
CTX2_SEPES 3129 bp        471 CTGGCATTTTTCTGTGATAAACGGGATGGAAAAATTAACAGAAACCTACACTTCTATGGCACAAGAAGCAGACATCAAAA
CTX1_DPEA 3156 bp         483 TTGGCAATATTCAGTGACTAATGGATTTGAAAAACTGGTAGAAACTTACACAGCAATGGTGAAAGAAACAGACATTAATA
CTX2_DPEA 3123 bp         462 GTGGCACTTTTCTGTCATAAATGGGATTGAAAAATTGTCCGAAACCTTCTCTCTTATGGCCATAGAAGCAGACATTAGGA
CTX1_SBAN 2946bp          480 TTGGCAATATTCTGTGAAAAATGGAATAGAAAAACTGGTAGAAACCTACTCGAAAATGGTGAAAGAAACGAATATCAATA
CTX2_SBAN 2286 bp         471 TTGGCATTTTTCTGTGATAAATGGAATAGAAAAATTAACAGAAACTTACACTTCTATGGCAAAAGAAGCAGACATTAGAA
CTX_EBER 3159 bp          486 CTGGCAATACTCGGTGTCAAACGGGATAGAGAAACTGGTTGAAACTTATTCAGCAATGTCGAAAGTGTCCGAAATAAATA
CTX2_SOFF 3129 bp         471 CTGGCACTTCTCCGTAATAAACGGCATGGAGAAATTAACAGAAACCTACACTTCTATGGCAAAAGAAGCAGACATCAGGA
CTX1_SOFF 3141bp          465 TTGGCAATATTCCGTGAAAAATGGAATCGAAAAACTGATCGAAACCTACAAGAAAATGGTGAGAGAAACGAATATAAATA
CTX_ESCO 3159 bp          486 CTGGCAATACTCGGTATCGAACGGGATCGAGAAACTGATTGAAACTTATTCAGCAATGTCGAAAGTGTCCGAAATAAATA
CTX2_SLES 3132 bp         471 GTGGTACTTTTCTATCATAAATGGGATTGAAAAATTGGCAGAAACCTATTCTGTTATGGCCAAAGAACCAGATATTAGGA
CTX1_SLES 3159 bp         486 TTGGCAATATTCAGTGACTAATGGAATCGAAAAACTGGTAGAAACTTACACAGCAATGGTGAAAGAACCAGACTTTAATA
CTX_TRHO 3150 bp          492 GTGGCATTTTTCTGTAATCAACGGGATTGAAAGGTTGACAGAAACCTACTCTGCCATGGCGAAAGAACCAGAAATCAGGA
CTX_SATL 3147 bp          483 CTGGCAATACTCGGTGACGAACGGGGTCGAAAAACTGACAGAAACATATTCAGCAATGTCGAAAGTGTCTGAGATAAACA
CTX2_SLYC 3129 bp         471 CTGGCATTTTTCTGTGATAAACGGAATGGAAAAATTAACAGAATCCTACACTTCTATGGCACAAGAAGCAGACGTCAGAA
CTX1_SLYC 3159 bp         483 TTGGCAATATGCTGTGAAAAATGGAATCGAAAAACTGATTGAAACCTACAAGACAATGGTAGAAGAAACGGATATTAATA
CTX_EPAR 3150 bp          480 CTGGCAATATTCGGTGTCGAACGGGATCGAGAAATTGATCGAAACTTATTCAGCAATGTCGAAAGTGTCCGAAATAAATA
CTX2_SPHAR 3129 bp        471 CTGGCATTTTTCTGTGATAAACGGGATGGAAAAATTAACAGAAACCTACACTTCTATGGCACAAGAAGCAGACGTCAGAA
CTX1_SPHAR 3159 bp        483 TTGGCAATATTCTGTGAAAAATGGAATCGAAAAACTGATTGAAACCTACAAGGCAATGGTGGAAGAAACGGATATCAATA
CTX_WSCI_PARTIAL 1653 bp    1 --------------------------------------------------------------------------------
CTX_IHAL 3156 bp          483 TTGGGAATATTCCATAACTAATGGGATTCATAAACTTGTCGAGACTTACACATCAATGGCAAAAGAATCGAACATGCATC
CTX2_XNOT 3183 bp         525 ATGGTATTTTACAGTCATTAACGGCGTGGAAATGCTAACAGAGACCTACATGTCCATGGCGAAAGAACAAGATCTCAGAA
CTX1_XNOT2 3150 bp        483 TTGGGAATATTCAATAACGAATGGGATAAACAAACTTATTGAAACTTATACAGCAATGGCAAAAGAATCTAACATGCATG
CTX_OROB_PARTIAL 1431 bp   21 TTGGCAATATTCGGTAAAACACGGAATGGAAAAACTGGTTGAAACTTACACAGCGATGGCGAAAGAAGGCGACATCAACA
CTX_SAFF 3147 bp          483 CTGGCAATACTCGGTGACGAACGGGGTCGAAAAACTGACAGAAACATATTCAGCAATGTCGAAAGTGTCTGAGATAAACA
CTX_SOBS 3156bp           483 CTGGCAATACTCGGTGGCGAACGGCGTCGAAAAACTGATAGAAACCTATTCAGCAATGTCGAAAGTGTCCGAAATAAACA
CTX_Sarc 2907 bp            1 -------------------------------------------------------ATGGCGAAAGAAGAAGATATTAATA
CTX2_Dopa 3123 bp         462 GTGGCACTTTTCTGTCATAAATGGGATTGAAAAATTGTCCGAAACCTTCTCTCATATGGCCATAGAAGAAGACATTAGGA
CTX1_Dopa 3123 bp         489 TTGGCAATATTCAGTGACTAATGGATTTGAAAAACTGGTTGAAACCTACACAGCAATGGTGAAAGAAACAGACATTAATA

                          641 .......650.......660.......670.......680.......690.......700.......710.......720
CTX1_SEPES 3159 bp        563 AACGACGACTTATGGCTGAAAATTTTATCTTATTCTTTGAAAATAACCAAATCGAATCAAACATTAACAATCTCATTAAA
CTX2_SEPES 3129 bp        551 AACGAAGAATTCAGGCCGAAAGATTCATAAAGTTTTTTGAAGACAACAATATTGAAAGTCATATTAACAATCTCATTAGG
CTX1_DPEA 3156 bp         563 AACGATCACTTATGTCAGAAAATTACATCTTATATTTTGAAAATAATCAAATCGAATCGAATATCAATAATCTCATTAAA
CTX2_DPEA 3123 bp         542 AACGACGGATACAAGCCGAAGATTTTATCAAGTTTTTTGAAGACAATCAGATTGAGAGTCATGTTAATAATCTCATTCAA
CTX1_SBAN 2946bp          560 AACGACGACTTCTAGCTGAAAATTTTATTTCATTCTTTGAAAATAATCAAATCGAAGCAAACATCAACAACCTCATTAAA
CTX2_SBAN 2286 bp         551 AAAGACGAATTCAAGCCGAAAATTTCATCAAGTTCTTTGAAGACAACAATATCGAAAGTCACATTAATAATCTCATTAGA
CTX_EBER 3159 bp          566 GGCGACGAATTATAGCTGAGAATTTTATCTTATTTTTCGAGAACAACCAAATTGAAGCAAATATCAACAATCTGGTGCGA
CTX2_SOFF 3129 bp         551 AACGACGAATACAGGCCGAAAGCTTCATCAAGTATTTTGAAGACAACAATATTGAAAGTAATGTTAACAATCTCATTAGG
CTX1_SOFF 3141bp          545 AACGACGACTTATAGCCGGGAACTTTATCTTATTCTTCGAGAATAATCAAATTGAATCCAACGTAAACAACCTCGTTAAA
CTX_ESCO 3159 bp          566 GGCGGCGAATTATAGCTGAGAATTTTATCTTATTTTTCGAGAACAACCAAATTGAAGCAAATATCAACAATCTGATGCGA
CTX2_SLES 3132 bp         551 AACGACGAAATCAGGCCGAAGATTTTATCAAATTTTTTGAAGACAACCAAATCGAAAGTCATTTTGATAATCTCATGAAG
CTX1_SLES 3159 bp         566 AACGACGACTAATAGCAGAAAATTTCATCTTATATTTTGAGAATAACCAAATCGAACTAAATATCAATAATCTCATTAAA
CTX_TRHO 3150 bp          572 AACGAAGAATCCAGGCGGAACACTTTATCAAGTTTTTTGAAGACAACCAGATCGAAAGTAATGTTAATAATCTCATTAGG
CTX_SATL 3147 bp          563 GGCGACGAATCATCGCTGAGAATTTCATCTTATTTTTCGAGAACAACCACATCGAAGCGAATATCAATAATCTGGTACGA
CTX2_SLYC 3129 bp         551 AACGACGAATTCAGGCCGAAAGATTCATTAAGTTTTTTGAAGACAACAATATCGAAAGTCATATTAACAATCTCATTAGA
CTX1_SLYC 3159 bp         563 AACGACGACTCATAGCTGAAAATTTTATCTTATTCTTTGAAAATAACCAAATCGAATCAAACATTAACAATCTCATTAAA
CTX_EPAR 3150 bp          560 GGCGGCGAATTATAGCTGAGAATTTTATCTTATTTTTCGAGAACAACCAAATTGAAGCGAATATCAACAATCTAATGAGA
CTX2_SPHAR 3129 bp        551 AACGACGAATTCAGGCTGAAAGATTCATTAAGTTTTTTGAAGAAAACAATATCGAAAGTCATATTAACAATCTCATTAGG
CTX1_SPHAR 3159 bp        563 AACGACGACTTATAGCTGAAAATTTTATCTTATTCTTTGAAAATAACCAAATCGAATCAAACATTAACAATCTCATTAAA
CTX_WSCI_PARTIAL 1653 bp    1 --------------------------------------------------------------------------------
CTX_IHAL 3156 bp          563 AAAGACGGGCTATTGCTGGAAATTTTATCAATTTTTTCGAGAACAACCAGATTGAATCGAATATCAATAATCTGATTTAT
CTX2_XNOT 3183 bp         605 AAAGACGAATCCTGGTCGAAAATTATATTAGCTATTTTGAAAGCAACCAGGTTGGTAGCAATGTTAACAATCTGCTGAGA
CTX1_XNOT2 3150 bp        563 AAAGACGGGTAATAGCCGGAGATTTCATCAATTTTTTCGAAAACAACCAGATAGAATCGAATATTAATAATCTAATTTAT
CTX_OROB_PARTIAL 1431 bp  101 AACGACGATTACTTGCGGAACATTTCATAATATTCTTTGAGAACAACCAAATCGAATCGAACATCTATAACCTGATCAAA
CTX_SAFF 3147 bp          563 GGCGACGAATCCTCGCTGAGAATTTCATCTTATTTTTCGAGAACAACCAAATCGAAGCGCATATCAATAATCTGGTGCGA
CTX_SOBS 3156bp           563 GGCGACGAATCCTCGCTGAGAATTTCATCTTATTTTTCGAGAACAACCAAATCGAAGCGAACGTCAACAACCTGATGCGA
CTX_Sarc 2907 bp           26 AACGGCGTATTATAGCCGAAGGTTTCATAATATTCTTTGAGAGCAATCAGATCGAATCGAACATCAATAACCTAATCAAA
CTX2_Dopa 3123 bp         542 AACGACGGATACAAGCCGAAGATTTTATCAAGTTTTTTGAAGACAATCAAATTGAGAGTCATGTTAATAATCTCATTCAA
CTX1_Dopa 3123 bp         569 AACGATCACTTATGTCAGAAAATTTCATCTTATATTTTGAAAATAATCAAATCGAATCGAATATCAATAATCTCATTAAA

                          721 .......730.......740.......750.......760.......770.......780.......790.......800
CTX1_SEPES 3159 bp        643 CTGACCACAACAACCGACGCGGTTCAT---CAAAACATGCTTTTTAACGAACTGCTTGACGAAGCAGGTTGTGATATTAT
CTX2_SEPES 3129 bp        631 ATCACGACGACGTCTGACTCTGCGTTT---TATAAGAACATTTACAGATTATTGATCAATAAAGCAGGCTGTAATTTGCC
CTX1_DPEA 3156 bp         643 TTGACCCTAACAACTGATGCAGTTCAT---CAAAATCAACTTTTTGATAAATTGATTGATGAAGCAGGTTGTGACATTGT
CTX2_DPEA 3123 bp         622 GTAACAACAAAGTCTGACTCGGTATTT---TATGAAAACATTTATCAATCGCTTTTCAATAAAGCTAACTGCAAGATGAA
CTX1_SBAN 2946bp          640 CTGACCACAACTGCTGATGCAGTCCAT---CGAAATGTGCTTTTTAACGAACTGATTGACGAAGCAGGTTGTGATTACAT
CTX2_SBAN 2286 bp         631 ATTACGACGACGTCTGATTCTGCGTTT---TATGAAAACGTTTACAGATTACTAATCAACAAAGCAGGTTGTAATTTACT
CTX_EBER 3159 bp          646 ATTACGACCACGTCCGATGCAATTCAA---CAAACGAACCTATTTATAGAACTGATTGAGGAATCTAAGTGTAACGTTAT
CTX2_SOFF 3129 bp         631 CTTACCACGACATCCGACTCTGCGTTT---TATAAAAACATTTACAGATTATTGCTTAATAAAGCCGGTTGTAATTTGCT
CTX1_SOFF 3141bp          625 CTGACCACAACAACCGATGCCGTCCAC---CAAAATATGCTTTTTGAAGAACTGATTGATGAAGCAGGTTGTGATATTGT
CTX_ESCO 3159 bp          646 ATTACCACCACGTCCGATTCAATTCAA---CAAACGAACCTATTTCAAGAACTGATTAAGGAATCTGAGTGTGACGTTAT
CTX2_SLES 3132 bp         631 CTAACAACGGTATCTGACTCAGTGTTT---TATAAAAATACTTTTCAATTACTTCTCAAAAAAGCTGGCTGCAAGATGCC
CTX1_SLES 3159 bp         646 TTGACTGTAACATCAAATATAATTCAG---CAAAATCAACTTTTCAATGAACTGATAGACGAAGCGGGTTGTAATATTAT
CTX_TRHO 3150 bp          652 CTAACGACAGTGTCTGACTCAGTGTTT---TATAAAAACATCTACTCATCATTGATCAATAAAGCCGGTTGTAATATGCA
CTX_SATL 3147 bp          643 ATTACGACCACCTCCGATACAATTCAA---CAAACGAACCTTTTCAAGGAACTGATTGAGAAATCCGATTGTGACATAAT
CTX2_SLYC 3129 bp         631 ATCACGACGACTTCTGACTCTGCGTTT---TATAAAAACATTTACATATTATTGATCAATAAAGCAGGTTGTAATTTGCC
CTX1_SLYC 3159 bp         643 CTGACCACAACAACCGATGCAGTTCAT---CAAAACATGCTTTTTAACGAACTGATTGACGAAGCAGGTTGTGATATTGT
CTX_EPAR 3150 bp          640 ATTACAACCACATCAGATACAATTCAA---CAAACGAACCTATTTCAAGAACTGATTGAAAAATCTGAGTGTGACGTAAT
CTX2_SPHAR 3129 bp        631 ATCACGACGACGTCTGACTCAGCGTTT---TATAAAAACATTTACAGATTATTGATCAATAAAGCAGGCTGTAATTTGCC
CTX1_SPHAR 3159 bp        643 CTGACCACAACAACCGATGCAGTTCAT---CAAAACATGTTGTTTAACGAACTGATTGACGAAGCAGGTTGTGATTTTGT
CTX_WSCI_PARTIAL 1653 bp    1 --------------------------------------------------------------------------------
CTX_IHAL 3156 bp          643 ATAACGACAACGTCGAGTGCAGTTCTCCATAAAAAGAAACTTTTTGATGAACTCATCGATAAGGCCGGTTGTAATTTCGT
CTX2_XNOT 3183 bp         685 ATGACCACTACGACCGATTCGGGAATA---CACAAAAATATTTATAAATTGATGCTTAAAAAAGCTGGCTGTAATTTATC
CTX1_XNOT2 3150 bp        643 ATAACGATAACGTCTAGCGCGAGTCTGGGTAAAAAGAAGCTTTTTGATGAACTTATTGATGAAGCGGGTTGTAATTTCAT
CTX_OROB_PARTIAL 1431 bp  181 TTGACCACCACGTCCGATATAGTTCAG---CAAAAGAACCTGTTCAACGAACTGATTGAGGAGGCGGGTTGCGACGCTAT
CTX_SAFF 3147 bp          643 ATTACGACCACCTCCGATACAATTCAA---CAAACGAACCTTTTTAAGGAACTGATTGAGAAATCCGATTGCGACATAAT
CTX_SOBS 3156bp           643 ATTACGACCACATCTGATACAATTCAA---CAAACGAACCTCTTTAAGGAACTGATTGAGAAATCCGAGTGTAACATAAT
CTX_Sarc 2907 bp          106 CTGACCACCGAATCCGATACAGTTCAG---AAAAA-AAACTGTTTAACGAACTGATTGACGAAGCGGACTGCGACATCAT
CTX2_Dopa 3123 bp         622 CTAACAACAAAGTCTGACTCGGTATTT---CATGAAAACATTTATCAATCGCTTTTCAATAAAGCTAACTGTAAGATGAA
CTX1_Dopa 3123 bp         649 TTGACCCTAACAACTGATACAATTCAT---CAAAATCAACTCTTTAATAAACTGATTGATGAAGCGGGTTGTGACATTGT

                          801 .......810.......820.......830.......840.......850.......860.......870.......880
CTX1_SEPES 3159 bp        720 TAGATTAACACGAATTTATATGCACGTCAGAAGAATTTTTTATCAAGGTACTCAATTAGTTTTAGCCTATAATTCATTTA
CTX2_SEPES 3129 bp        708 CAAATTAAAGGTCATTTATGAGAGGGTGACTCAGATCGTTACTTCCGGGTCTCAGCTTATATTGGCATATCGCAGTTTTA
CTX1_DPEA 3156 bp         720 CAGGTTAACACGATTGCATTTTCAGATCCAACAAATATTTACTCAAGGTTGTCAATTAGTTTTAGCCTACAACTCATTTA
CTX2_DPEA 3123 bp         699 CAAATTAAACTTAATTCACGAGAAAGTAACTGGGATTATTACTTCGGGGTCTCAACTTGTTTTGGCATATCGAAGTTTAA
CTX1_SBAN 2946bp          717 TAGGTTAACGCGAATTTATATGCACATCAGAAGAATTTTTTTTCAAGGTTGTCAATTAGTTTTAAGCTATCATTCATTTA
CTX2_SBAN 2286 bp         708 CAAATTGAAGGTCATTTATGAAAGGGTGACTCAGATCGTTACTTCCGGTTCTCAACTTATATTAGCATATCGCAGTTTTA
CTX_EBER 3159 bp          723 CCGATTATCGAATGTCTATCTACAGGTGAAACGGATTTTTATGCAAGGTTCTCAATTGGTTCAAGCCTATCACGCCTTCA
CTX2_SOFF 3129 bp         708 CCGATTAAAGGTTATTTATGAGAGGGTGAGTCAAATCGTCACTTCCGGTTCTCAACTTATTTTGGCCTATCGCAGTTTTA
CTX1_SOFF 3141bp          702 TAGATTGACGCGTATTTATATGCACGTCAGAAGAATTTTCTATCAGGGTTGTCAGTTGGTTTTAGCTTATAATTCATTTA
CTX_ESCO 3159 bp          723 CCGATTATCGAATGTCTATCTACATGTGAAACGGATTTTTATGCAAGGTTCTCAATTGGTTCAAGCCTATCACGCCTTCA
CTX2_SLES 3132 bp         708 GAAATTAAACTTAATTTATGAGAGAGTGACTCGAATCCTTACTTCAGGTTCTCAACTTGTTTTGGCATATCGCAGTTTTA
CTX1_SLES 3159 bp         723 CAGGTTAACTCGATTGCATATTTATATCAAACGAATCTTTACTCAAGGTGGTCTATTAATTGTAGCTTACAACTTGTTTA
CTX_TRHO 3150 bp          729 AAAATTAAATTCAATTTACGAAAGAGTGACTCGGATCATTACTTCTGGTTCTCAACTTATTTTGGCATATCGCAGTTTTA
CTX_SATL 3147 bp          720 CCGATTAACGAATGTCTATCTACATGTGAAACGGATCTTTATGCAAGGTTCACAACTGGTTCAAGCCTATCACGCCTTCA
CTX2_SLYC 3129 bp         708 CAAATTAAAGGTCATTTATGAGAGGGTGACTCAAATCGTTACTTCCGGGGCTCAGCTTATATTGGCATATCGCAGTTTTA
CTX1_SLYC 3159 bp         720 TAGATTAACACGAATTTATATGCACGTCAGAAGAATTTTTTATCAAGGTACTCAATTAGTTTTAGCCTATAATTCATTTA
CTX_EPAR 3150 bp          717 CCGATTATCGAACGTCTATCTACATGTGAAACGGACTTTTATGCAAGGTTCTCAACTGGTTCAAGCCTATCACACCTTCA
CTX2_SPHAR 3129 bp        708 CAAATTAAAGGTTATTTATGAGCGGGTGACTCAAATCGTTACTTCCGGGTCTCAGCTTATATTGGCATATCGCAGTTTTA
CTX1_SPHAR 3159 bp        720 TAGATTAACACGAATTTATATGCACGTCAGAAGAATTTTTTATCAAGGTACTCAATTAGTTTTAGCTTATAATTCATTTA
CTX_WSCI_PARTIAL 1653 bp    1 -------------------------------------------------------------------------------A
CTX_IHAL 3156 bp          723 AAGATTAACTGGCATTTATACGCACATGAAACGAATATTTACTCAAGGTTCCCAATTGATTTTAGCTTATTATACTTTTA
CTX2_XNOT 3183 bp         762 GAAACTAAAGGAGGTTCATGAAAAGGTTGCCCGGATAATTATGTCTGGTTCAGAACTTGTTTTAGCTTATCGAATTTTTA
CTX1_XNOT2 3150 bp        723 CAGATTGACCGGAATTTATACGCATATGAGAAGAATCTTTACCCAAGGATCCCAATTGGTTTTAGCTTATTATTCTTTTA
CTX_OROB_PARTIAL 1431 bp  258 AAGGTTAACACAGGTCTATTTGCATATCAAGCGAATCTTTACTGAGGGTTGTCAATTGATTTTAGCCTATCGATCCTTTA
CTX_SAFF 3147 bp          720 CCGATTAACGAATGTCTATCTACATGTGAAACGGATCTTTATGCAAGGTTCTCAACTGGTTCAAGCCTATCACGCCTTCA
CTX_SOBS 3156bp           720 TCGATTAACGAATGTCTACCTACATATGAAACGGATCTTTATGCAAGGTTCTCAACTGGTCCAAGCCTATCACGCCTTCA
CTX_Sarc 2907 bp          182 CAGGTTAGCACAAGTTCATTTGCATATCAAACGAATATTTACCCAAGGTTCTCAATTAATTCTAGCCTATCACTCCTTGA
CTX2_Dopa 3123 bp         699 CGAATTAAACTTAATTCATGAGAAAGTAACTGGGATTGTTACTTCGGGGTCTCAACTTGTTTTGGCATATCGAAGTTTTA
CTX1_Dopa 3123 bp         726 CAGGTTAACACGGTTGCATTTTCAGATCCAACAACTATTTACTAAAGGTTGTCAATTAGTTTTAGCCTACAACTCATTTA

                          881 .......890.......900.......910.......920.......930.......940.......950.......960
CTX1_SEPES 3159 bp        800 AACAAATGGATCCTCCAGAAATGAAAAAATATCTTAACGCTTTGATATTTATAAGAAATATGTATCAATCTCGCGTTTGG
CTX2_SEPES 3129 bp        788 TGCAAATAGAGATACCAAAGTTGAAAAAATACTTGGACGCTTTGCTTTTATTTCGACAAATATACGAAAAGAGTATTTGG
CTX1_DPEA 3156 bp         800 AGCGAATGGAATCCCCATCCATTCAAAAGTATATTGACGCTTTGATATACATTCGGAACATTTACGAATCTCGCGTTTGG
CTX2_DPEA 3123 bp         779 AACAATTGGAGAAACCAAAAATGACAAAGTATCTAAATGCCTTATTTTCACTTCGGAAAATGTTTGATGCACAAACTTGG
CTX1_SBAN 2946bp          797 AACAAATGGATCCTCCAGAAACTAAAAAATATATTAACGCTTTGGTATTTATCCGAAACATGTACCAATCTCGCGTTTGG
CTX2_SBAN 2286 bp         788 TGCACATGCAGGTACCAAAGTTGAAGAAATATTTCGACAGTTTATTTATATTCCGCCAAATATATGAAAAAAGTATTTGG
CTX_EBER 3159 bp          803 AGCAGATGGAGATGCCGAACATTCAAAAGTATCTCGACGCTTTGGTGTACATTCGGACAGCATACGAATCCCGCCTTTGG
CTX2_SOFF 3129 bp         788 TGCAAATGCAGATACCGAAGGCGAAAAAATATTTCGACGTCTTGTTTTTATTTCGACAAATATATGATAAGCGTATTTGG
CTX1_SOFF 3141bp          782 AACAAATGGAACCCCCAGAAATTAAAAAATATCTTAACGCTTTGATATTCATTCGAAACATGTACCAAGCTCGCATTTGG
CTX_ESCO 3159 bp          803 AGCAGATGGAGATGCCGAACATTCAAAAGTATCTCGACGCTTTGCTGTATATTCGGACAGCATACGAATCCCGCCTTTGG
CTX2_SLES 3132 bp         788 AACAATTAGAGAAACCAAAAATGACAAAATATATGAATGCCTTATTTTTACTTCGACAAATATTTGAAGCACATGATTGG
CTX1_SLES 3159 bp         803 AACAAATGGAGCCGCCAAACATCCAAAAGTTTCTTAACGCTTTGATATACATTCGGAAAATTTACGAATCCCGCGTTTGG
CTX_TRHO 3150 bp          809 AACAAATGGATAAACCAAAGTTGAAAAAATATTTGAACGCCTTGTTTTCTCTTCGGCAAATTTTCGAGGCTCGCGTTTGG
CTX_SATL 3147 bp          800 AGCAAATGGAGATGCCGAATATTCAAAAGTATCTCGACGCCTTGGTGTACATTCGGACAGTTTACGAATCCCGCGTCTGG
CTX2_SLYC 3129 bp         788 TGCAAATACAGATACCAAAGTTGAAAAAATATTTGGACGCCTTGTTTTTATTTCGACAAATATACGAAAAGCGTATTTGG
CTX1_SLYC 3159 bp         800 AACAAATGGATCCTCCAGAAATTAAAAAATATCTGAACGCTTTGATATTTATAAGAAATATGTATCAATCCCGCGTTTGG
CTX_EPAR 3150 bp          797 AACAGATGGAGATGCCGAACATTCAAAAGTATCTCGACGCTTTAGTGTATATTCGAACAGTATACGAATCTCGCCTTTGG
CTX2_SPHAR 3129 bp        788 TGCAAATACAGATGCCAAAGTTGAAAACATATTTGGACGCCTTGTTTTTATTTCGACAAATATACGAAAAGCGTATTTGG
CTX1_SPHAR 3159 bp        800 AACAAATGGATCCTCCAGAAATTAAAAAATATCTTAATGCTTTGATATTTATAAGAAATATGTATCAATCCCGCGTTTGG
CTX_WSCI_PARTIAL 1653 bp    2 AACAAATGGAACCTCCACGTATTGAGCAGTTCACTGAAGCTTTGTTAGTTATCAGGAAATTGTATGGAAATCGAATATGG
CTX_IHAL 3156 bp          803 AAGACGGCAAAGTTCCGCTCGTTCAAACATATATTGATGCTTTAACCTCCATCCGTAATACGTACGATTATCGAATCTGG
CTX2_XNOT 3183 bp         842 AACGTATGGAAAAACCAAACCAGAAGAAATATATAAACGCTTTGCTTTTTTACAGAAGACTCTACGAAAACCAAGTTTGG
CTX1_XNOT2 3150 bp        803 AGGACGGAAAAGTCCCGCTTGTTCAAACATACATTGATGCTTTAACATCAATCCGTAATACCTATGACTATCGAATCTGG
CTX_OROB_PARTIAL 1431 bp  338 AGCAAATGGAAGCTCCGCGCATAGAGGGGTTCCTCGACGCTGTGGCGTTCATTCGGAAAATGTACGACAAACGCGTCTGG
CTX_SAFF 3147 bp          800 AGCAAATGGAGATGCCGAATATTCAAAAGTATCTCGACGCCTTGGTGTACATTCGGACAGTTTACGAATCCCGCGTCTGG
CTX_SOBS 3156bp           800 AGCAAATGGAGATGCCGAGTATCCAAAAATATCTCGACGCCTTGGTGTACATTCGGACACTTTACGAATCCCGCGTCTGG
CTX_Sarc 2907 bp          262 AGCATATGGAACCTCCGCGCATTCAGAAGTTCCTTGAGGCTTTATTATTCACTCGGAAAGTGTACGGGAACCGCGCTTGG
CTX2_Dopa 3123 bp         779 AACAATTGGAGAAACCAAAACTGACAAAGTATCTAAATGCCTTATTTTCATTTCGGAAAATGTTTGATGCACAAACTTGG
CTX1_Dopa 3123 bp         806 AGCAAATGGAATCCCCATCCATTCAAAAGTATATTGACGCTTTGATAAAAATTCGGAACATTTACGAATCTCGCGTTTGG

                          961 .......970.......980.......990......1000......1010......1020......1030......1040
CTX1_SEPES 3159 bp        880 CACTGCAAAGAAACAACGATCGCCCAGTCCAAGAAAGATATCAAAGATATTGTCAAGACAAATGCTAAATTTGGA-----
CTX2_SEPES 3129 bp        868 CACTGCAAGGAAACGGCTATTGATCGTTCAAAGAAAACCATTTATAGAATGCTCATTGGGAAGAAAT-------------
CTX1_DPEA 3156 bp         880 TACTGTAAAGAAACGACAATAGTTCAGTCCAAAAAGGATGTTGTAAAAATTGTGACAGCAAATTCTCACCTTAGT-----
CTX2_DPEA 3123 bp         859 TATTGTAAAGAAACGGCTATTGATCGTTCGAAAAAGGCAATTTTTCAAATGATTAATGGAACTAAGA-------------
CTX1_SBAN 2946bp          877 CATTGTAAAGAAAACACAATCGCTCACTCCAAGAAAGATATCAAAGATGTTGTCAAGAAGGATGCTAAATTAGGA-----
CTX2_SBAN 2286 bp         868 CACTGTAAGGAAACTGCTATTTCTCGCTCGAAGAACGCCGTTTATAGAATGCTCTATGGGAAAAAGT-------------
CTX_EBER 3159 bp          883 CGCTGTAAAGAAACAGCAATCCTTCGATCAAAGAACACTATCGTGGAGATAGTCAAGCGAGACCCACACCTTGGG-----
CTX2_SOFF 3129 bp         868 CGCTGTAAGGAAACGGCTATTGATCGTTCGAAGAGAGCCGTTTATAGAATGCTCATTGGGGAGAAGT-------------
CTX1_SOFF 3141bp          862 CACTGTAAAGAAACCACGATTGCCAACTCCAAGAAAGACATCAAGGATATTGCCAAGACACATGCTAAATTTGGT-----
CTX_ESCO 3159 bp          883 CGCTGTAAAGAAACATCAATCCTTCGATCAAAGAATTCTATCATCGAGATAGCCAAGCGAGACTCACACCTTGGG-----
CTX2_SLES 3132 bp         868 CATTGTAAGGAAACGGCTATTGCCCGCTCTAAAAAGGCCGTTCTTCAAATGATCTCTGGGAATAAGA-------------
CTX1_SLES 3159 bp         883 CACTGTAAAGAAACGACAATAATCCGCTCCAAGAATGATGTTGTAAAAATTATCACGAAAAATCCTCACTTGGAA-----
CTX_TRHO 3150 bp          889 TATTGTAAGGAAACAGCAATTCCTCGTTCTAAAAGAAACATCCTTAGAATGCTTGGTGGGAGAAAGT-------------
CTX_SATL 3147 bp          880 CGCTGTAAGGAAACATCGATCGTTCGATCGAAGAATGATATCATAAAGATTGCCAAGAGAGACTCGAACGTCGGG-----
CTX2_SLYC 3129 bp         868 CACTGCAAGGAAACGGCAATTGATCGCTCAAAGAAAACCATTTATAGAATGCTCAATGGGAAGAAGT-------------
CTX1_SLYC 3159 bp         880 CACTGCAAAGAAACCACGATCGCCCACTCCAAGAAAGATATCACAGATATTGTCAAGACAAATGCTAAATTTGGA-----
CTX_EPAR 3150 bp          877 CGCTGTAAAGAAACATCGATCCTTCGATCGAAGAACGAGATCATGAAGATAGTCAAGCGAGACTCACATCTTGGG-----
CTX2_SPHAR 3129 bp        868 CACTGCAAGGAAACGGCTATTGATCGCTCAAAGAAAACCATTTATAGAATGCTCAATGGGAAGAAGT-------------
CTX1_SPHAR 3159 bp        880 CACTGCAAAGAAACTACGATCGCCCACTCCAAGAAAGATATCACAGATATTGTCAAGGCAAATGCTAAATTTGGA-----
CTX_WSCI_PARTIAL 1653 bp   82 CATTGTAAAGAAACCACAATCGCCCGATCCAAGACCGACATAGAAGAAATTGTCAAGAATAATACCGGTAATGGG-----
CTX_IHAL 3156 bp          883 CACTGTAAGGAAACAACTATCACTCGCTCGAAAAGTGATGTAGATAAACTAGTGAAGAAGAACAGTAACGAAGGAAGAGG
CTX2_XNOT 3183 bp         922 TATTGTAAAGAAATGGCCATTATTCGCTCGAAGAGTGCTGTTAAAAAAATGTTAAGTGGAGTAAAAG-------------
CTX1_XNOT2 3150 bp        883 CACTGTAAGGAAACGACTATTACCCGATCAAAAAGTGATGTTGATAAAGTAGTAACGAAAAGCAGTGAAGAAGAGAGGGG
CTX_OROB_PARTIAL 1431 bp  418 CACTGTAAAGAAACCACGATCGTCCGCTCCAAGGATGACATAGCAAAAATTGCCAAGAAATATTCTAACTTCGGA-----
CTX_SAFF 3147 bp          880 CGCTGTAAGGAAACATCGATCGTTCGATCGAAAAATGATATCATAAAGATTGCCAAGAGAGACTCGAACGTCGGG-----
CTX_SOBS 3156bp           880 CGCTGCAAAGAAACGTCGATCACTAGATCGAAGAACGATATTACAAAGATAGCCAAGCGAGACTCGCGCCTCGGG-----
CTX_Sarc 2907 bp          342 CACTGTAAAAAAAACACGATCACCCTCTCCAAGAATGACATTGTAAAAAA-----AGAATGATTCCGACCTCGGA-----
CTX2_Dopa 3123 bp         859 TATTGTAAAGAAACGGCTATTGATCGTTCGAAAAAGGCCGTTTTTCAAATGATTAATGGAACTAAGA-------------
CTX1_Dopa 3123 bp         886 TACTGTAAAGAAACGACAATAGTTAACTCCAAGAAGGATGTTGTAAAAATTGTCACGACAAATTCTCACCTTAGT-----

                         1041 ......1050......1060......1070......1080......1090......1100......1110......1120
CTX1_SEPES 3159 bp        955 -ATTACTACAGTACTAAGAAAAATTAATAGTGAACTTTCTAGAAAATATCCTTGGTACAGCTGGTCGATCGTCACTGTCA
CTX2_SEPES 3129 bp        935 --CACGTGTTCCCTTGAAACGCATCGCCAGTATGCTTTCCCGTAATTTTCCGTGGTATTCTTGGTCACTTGGCTTAACAC
CTX1_DPEA 3156 bp         955 -ATAACTCCGTTACTGAAAAATATTAATGATGACCTTTCGAGAAATTATCCTTGGTACAGTTGGTCAATTGTAAATATCA
CTX2_DPEA 3123 bp         926 --AACGATTTCCCTTAAAGCGAATCGGAAAAATGCTTTCCATTAATTATCCGTGGTATTCTTGGTCACTTGGCCTCACAA
CTX1_SBAN 2946bp          952 -ATTACTACAGTAATAAAAAATATTAATAGTGAACTCTCTAAAAAATATCCTTGGTACAGCTGGTCGATCGTAACTGTCA
CTX2_SBAN 2286 bp         935 --CACGAGTTTCATTAAAAGTTATCGCCAATATGCTTTCCTACAATTATCCGTGGTATTCTTGGTCACTTGGCTTAACAC
CTX_EBER 3159 bp          958 -ATAATAAAACTACTAGTAAACATTAAGGAAGAGCTTTCGAAGACATATCCTTGGTACAGTTGGTCAGTTGTGAATCTCC
CTX2_SOFF 3129 bp         935 --CACGGTCTTCATTGAAACGCATCGCCAATATGCTTTCCAGTAATTATCCGTGGTATTCTTGGTCACTTGGTTTAACAA
CTX1_SOFF 3141bp          937 -ATTTCTACATTATTAACAAATATTAATAAGGAGCTTTCTAAAAAATATCCTTGGTATAGCTGGTCGATCGTAACTATCA
CTX_ESCO 3159 bp          958 -ATAATAAAACTACTAGTAAACATTAAGAAAGAGCTTTCGAAGACATATCCTTGGTACAGTTGGTCGGTTGTGAATCTCC
CTX2_SLES 3132 bp         935 --AAAGAGTAGGATTAAAAAGCATCGCCAAAATGCTTTCCATTAATTATCCGTGGTATTCTTGGTCACTTGGTCTAACAA
CTX1_SLES 3159 bp         958 -ATAACAACGCTACTAAAAAAAATTATTGATAACCTTTCGAGGAATTATCCTTGGTACAGTTGGTCGATTGTGAATGTCA
CTX_TRHO 3150 bp          956 --CAAGACCGGCATTGAAACGCATCGGTAATGTCCTTTCCAAGAATTATCCGTGGTATACTTGGTCACTTGGCCTAACTC
CTX_SATL 3147 bp          955 -ATCATAAAACTACTGGGAAACATTAACAGGGATCTTTCGAAGACACATCCTTGGTACAGTTGGTCGATTGTGAATCTCC
CTX2_SLYC 3129 bp         935 --CACGGGTTCCCTTGAAACGCATCGCCAGTATGCTTTCCCGTAATTTTCCGTGGTATTCTTGGTCACTTGGTTTAACAC
CTX1_SLYC 3159 bp         955 -ATTACTACAGTACTAAGAAAAATTAATAGTGAACTTTCTAGAAAATATCCTTGGTACAGCTGGTCGATCGTAACTGTCA
CTX_EPAR 3150 bp          952 -ATAATAAAACTACTAGTAAACATTAAGAAAGAGCTTTCGAAAACACATCCATGGTACAGTTGGTCGGTTGTGAATCTCC
CTX2_SPHAR 3129 bp        935 --CACGGGTACCCTTGAAACGTATCGCCAGCATGCTTTCCCGTAATTTTCCGTGGTATTCTTGGTCACTTGGCTTAACAC
CTX1_SPHAR 3159 bp        955 -ATTACTACAGTACTAACAAAAATTAATAGTGAACTTTCTAGAAAATATCCTTGGTACAGCTGGTCGATCGTAACTGTCA
CTX_WSCI_PARTIAL 1653 bp  157 -ATAACGTCATCACTTGAAAAGATTAACGCAGAACTCTCGAAAAAATATCCTTGGTACTCCTGGTCAGTCGTGAATGTCC
CTX_IHAL 3156 bp          963 AGCAACGGTTACAGTAAAGAATGTCAATAAGGCCCTTTCGAAAAAGTACCCTTGGTACAGTTGGTCAATTATTAACAACA
CTX2_XNOT 3183 bp         989 --CTATTCCATCACTGAAGCGAATTGCGAAAATGCTTTCTGCAAATTTCCCATGGTACTCTTGGTCCCTTGGCATAACAA
CTX1_XNOT2 3150 bp        963 AGCAACTGTTACACTAAACAATGTCAATAAGGCTCTTTCGCAAAAGTATCCTTGGTACAGTTGGTCAATTGTTAACATCA
CTX_OROB_PARTIAL 1431 bp  493 -ATACCTGCACAGCTTAAGAATATCAATGTTGCTCTCTCTAGGAAATATCCCTGGTACAGTTGGTCGATCGTGAATGTCG
CTX_SAFF 3147 bp          955 -ATCATAAAACTACTGGGAAACATTAACAGGGATCTTTCGAAGACACATCCTTGGTACAGTTGGTCGATTGTGAATCTCC
CTX_SOBS 3156bp           955 -ATAATAAAATCACTTGGAAACATTAACAAGGAGCTTTCGAAGACACATCCTTGGTACAGTTGGTCGATCGTAAATCTCC
CTX_Sarc 2907 bp          412 -ATAACTACACTACTGAAAAATGTCAATGGTGCTCTCTCAAAAAAATATCCTTGGTACAGTTGGTCGATCGTAAATGTCA
CTX2_Dopa 3123 bp         926 --AAAGATTTCCCTTAAAACGAATCGGCAAAATGCTTTCCATTAATTATCCGTGGTATTCTTGGTCACTTGGCCTCACAA
CTX1_Dopa 3123 bp         961 -ATAACTCCGTTACTGAAAAATATTAATTATGGTCTTTCGAGAAAGTATCCTTGGTACAGTTGGTCAATTGTAAATGTCA

                         1121 ......1130......1140......1150......1160......1170......1180......1190......1200
CTX1_SEPES 3159 bp       1034 A---AAAAATGCTTGCAAACCAAAGAAATTCAACTTTAGGCAATCAATTTTATGAAATGGAAGCGGTTGGGCCACATGGT
CTX2_SEPES 3129 bp       1013 G---GAAATGGATAGGTTCCCGCGAAAACGTTGTATCGGGGAATCAATACTATGAACTGAAAGATCTGTGGCCGCTTGGA
CTX1_DPEA 3156 bp        1034 G---AAGAATGCTGAGGCAGGAAAAGAATTCTCCTATAGGAAACCAATATTATGAACTGGAAAAGGTCGGGTCAAATGGT
CTX2_DPEA 3123 bp        1004 AACCAAAATCGCCACATTTCCGTGATAACTTTTATTCGGGAAACCAATTCTATGAACTGAAAGGTCTATGGCCTCTTGAA
CTX1_SBAN 2946bp         1031 G---AAAAATGCTTTCAGACCAAAGAAATTCGACTATTGGCAATCAGTATTATGAAATGGAAAAGGTTGGACCATATGGC
CTX2_SBAN 2286 bp        1013 G---AAAACGGATAGGTTCCCGTGAAAACGTAGTCTCGGGAAATCAATATTATGAACTGAAAGATTTGTGGCCGTTTGGA
CTX_EBER 3159 bp         1037 A---GAGAATGATTGGAGCGGAGGTTAATGCGACAACGGGAAATCAATTTTATGAAATGGAAAAGGTTGGATCATACGGT
CTX2_SOFF 3129 bp        1013 C---AAAATGGACAGATTCCCGGGAAAACGTAGTATCGGGAAACCAATATTATGAACTGAAAGATCTGTGGCCGTTTCGA
CTX1_SOFF 3141bp         1016 G---AAAACAGCTTGCAAGCCAAAGAAATTCGACTTCAGGCAACCAGTATTACGAAATGTTATCGGTTGGAGAATATGGT
CTX_ESCO 3159 bp         1037 A---GAGAATGATTGGAGCAGAGGTTAATGCGACATCGGGAAATCAATTTTATGAAATGGAAAAAGTTGGATCATACGGT
CTX2_SLES 3132 bp        1013 AAAAAAGATCTCTACATTCCCGTCAGAACTTCGTTTCAGGAAATCAATTCTATGAGCTGAAAGGTCTGTGGCCTTTTGAA
CTX1_SLES 3159 bp        1037 G---AAGAATGCTGAGTGAAGAAAGGAATTCTACTTTAGGGAACCAATATTATGAACTGGAAAAGATTGGACCAAAGGGC
CTX_TRHO 3150 bp         1034 G---AAACCAGCTAAATTTCAACCGGGAATCAGCCTCGGGAAACCAATTCTATGAACTGAAACGGCTGTGGCCGCTTGGA
CTX_SATL 3147 bp         1034 G---GAGAATGATCGGAGCGGAGGCTAATGCGACATCAGGGAATCAGTTTTATGAGATGGAAAAGGTTGGATCATACGGT
CTX2_SLYC 3129 bp        1013 G---GAAATGGATAGGTTCCCGCGAAAACATCGTATCGGGGAATCAATACTATGAACTGAAAGATCTTTGGCCGCTTGGA
CTX1_SLYC 3159 bp        1034 A---AAAACTGCTTGCAAACCAAAGAAATTCGACTTTAGGAAATCAATTTTATGAAATGGAAGCGGTTGGGCCACATGGT
CTX_EPAR 3150 bp         1031 G---GAGAATGATTGGAGCGGAGGTTAATGCGACGTCGGGAAATCAATTTTATGAAATGGAAAAGGTTGGATCATACGGT
CTX2_SPHAR 3129 bp       1013 G---GAAATGGATAGGTTCCCGCGAAAACGTCGTATCGGGGAATCAATACTATGAACTGAAAGATCTTTGGCCGCTTGGA
CTX1_SPHAR 3159 bp       1034 A---AAAACTGCTTGCAAGCCAAAGAAATTCGACTTTAGGGAACCAATTTTATGAAATGGAAGCGGTTGGGCCACATGGT
CTX_WSCI_PARTIAL 1653 bp  236 G---GAGACTGCTAGAAGCTGAACAGGATTCCGTAATTGGAAATCAGTATTATCAAATGGAAAAGGTTGGGAAACATGGT
CTX_IHAL 3156 bp         1043 A---AAGATGGGTGGGTTCGAAGAAAAGCGCCACCTCGGGCAATCAGTATTACGAAATGTTAAAAGTAGGGAAATATGGT
CTX2_XNOT 3183 bp        1067 A---AAAATGGAATAGCGCCAGGCACAATTATGTCTCGGGTAACCAATTCTATGAAATGGAAGGTTATCTGCCTTTGGGA
CTX1_XNOT2 3150 bp       1043 A---AAGATTGACTAGTTC------GACCTCTGCATCTGGTAATCAGTACTACGAAATGATAAAAGTAGGGAAATATGGT
CTX_OROB_PARTIAL 1431 bp  572 G---GGGATTGCTTGAAGC------GGATTCTGCAATGGGGAACCAGTATTATGAAATGACAAAGGTTGGTTCACATGGT
CTX_SAFF 3147 bp         1034 G---GAGAATGATCGGAGCGGAGGCTAATGCGACATCAGGGAATCAGTTTTATGAGATGGAAAAGGTCGGATCATACGGT
CTX_SOBS 3156bp          1034 G---GAGACTGATCGGAGCGGAGGTTGATTCGACTTCCGGAAATCAGTATTACGAGATACAAAACGTTGGATCACACGGT
CTX_Sarc 2907 bp          491 G---GAGACTGCTTGATGCGAAAAGTGGTTCTACTATGGGGAACCAATATTATCAAATGGAAAAAGTTGGTTATTATGGT
CTX2_Dopa 3123 bp        1004 AACAAAAATCGCCACATTTCCGTGATAACTTTTATTCGGGAAACCAATTCTATGAACTGAAAGGTCTATGGTCTCTTGAA
CTX1_Dopa 3123 bp        1040 G---AAAATTGCTGAGGAATGAAAAGAATTCTCCAACAGGAAACCAATATTATGAACTGGAAAAGGTTGGGTCAAATGGT

                         1201 ......1210......1220......1230......1240......1250......1260......1270......1280
CTX1_SEPES 3159 bp       1111 TCGAACTTTGTCGTGATTTGGCAAGGCTTTAAAGAACATTCACAATGTGAAGATATTCAGAAAGCCAACACTGTCGCTGT
CTX2_SEPES 3129 bp       1090 ATATATTTGGTGGTAATTTGGCAGGGTGTAACCGAAACCTCTCAATGTAATCAAATGCCCAAAGCCAACACAGTCGTCTT
CTX1_DPEA 3156 bp        1111 GTGAACTTGGTTGTCACTTGGCAGGGTTCAAATGAAAAATCCCAATGTCGAGAGATTGGGAAAGCCAATACTTTTCTTTT
CTX2_DPEA 3123 bp        1084 ACATCTTTGCTGATAACATGGCAAGGTGTAAATGAAACATCTCGATGCAATCGAATGCTAAAAGCAAATACAATTATCTT
CTX1_SBAN 2946bp         1108 TCGAACTTTGTTGTGACTTGGCAAGGCTCCAAAGAAAATTCACAGTGTGAAGATATTTACAAAGCCAATACTGTTGTCGT
CTX2_SBAN 2286 bp        1090 ATACATTTGATGGTAATTTGGCAGGGTGTAACCGAGACATCTCATTGTAATCAGATGCCCAACGCCAACACAGTTGTCTT
CTX_EBER 3159 bp         1114 TGGAATCTAGCTGTCACTTGGCAGGGTACAGACGAGAAATCTCAGTGTCAAGACATCGGAAAGGCTCAGACTGTTGTCTT
CTX2_SOFF 3129 bp        1090 ACATATTTGGTCGTAATATGGCAGGGTGTAACCGAAACATCTCAATGTAATCGCATGCCCAAAGCCAACACAGTCGTCTT
CTX1_SOFF 3141bp         1093 TCGAACCTTGTCGTTACTTGGCAAGGCTCCAAAGAAGAGGCACAGTGTGAGGATATTCAGAAAGCCAACACTGTTGTGGT
CTX_ESCO 3159 bp         1114 TGGAATCTAGCTGTCACTTGGCAGGGTACAGACGAGAAATCTCAGTGTCAAGACATCGGAAAGGCTAAGACTGTTGTCTT
CTX2_SLES 3132 bp        1093 ATGACTTTGATAATAACTTGGCAAGGTGTAAACGAAATATCTCGATGTAATCACATGCTAAAAGCCAATACAATTATCTT
CTX1_SLES 3159 bp        1114 GTGAACTTAGTTGTGACTTGGCAAAGTTCCAACGAAAAATCCTCTTGTCAAGCGATTGGGAAAGTCAATACTTTTGTCTT
CTX_TRHO 3150 bp         1111 ATATCTCTCGTGATAACTTGGCAGGGTGTAAACGAAGTATCCCGATGTAATCAAATGCTAAAAGCCAACACGATAGTGTT
CTX_SATL 3147 bp         1111 TGGAATTTGGCTGTCGCTTGGCAGGGTACCGACGAGAAGTCTCAGTGTCAAGACATCGGAAAGGCCAATACTGTTGTATT
CTX2_SLYC 3129 bp        1090 ATATATTTGGTGGTAATTTGGCAGGGTGTAACCGAAACCTCTCAATGTAGTCACATGCCCAAAGCCAACACAGTCGTCTT
CTX1_SLYC 3159 bp        1111 TCGAACCTTGTCGTGATTTGGCAAGGCTCCAAAGAAAATTCACAATGTGAAGATATTCAGAAAGCCAACACTGTCACCGT
CTX_EPAR 3150 bp         1108 TGGAATCTAGCTGTCACTTGGCAGGGTACAGACGAGAAATCTCAGTGTCAAGACATCGGAAAGGCCAATACTGTTCTCTT
CTX2_SPHAR 3129 bp       1090 ATATATTTGGTGGTAATTTGGCAGGATGTAACCCAAACCTCTCAATGTAATCAAATGCCCAAAGCCAATACAGTCGTCTT
CTX1_SPHAR 3159 bp       1111 TCGAACCTTGTCGTGATTTGGCAAGGCTCCAAAGAAAATTCACAATGTGAAGATATTCAGAAAGCCAACACTGTCGCCGT
CTX_WSCI_PARTIAL 1653 bp  313 GTGAATTTGGTCGTGGCCTGGCAAGGGTCAGATGAGAAACCCCAGTGTTCAGACGTTCGTGAAGCAAACACTTTCTTGTT
CTX_IHAL 3156 bp         1120 ATGAATATCGTGGTTGCCTGGCAAGGAGTCGATGAAAAAGCTGAGTGTTATGATATCGACCGATCAAAAACCTTTCTTTT
CTX2_XNOT 3183 bp        1144 CTGTCAGTAATTGTCGTATGGCAAGGTGTGAATGAAAAGTCTAACTGTAAACTAATGCCCAAAGCTAATACTATCGTGTT
CTX1_XNOT2 3150 bp       1114 ATGAATTTAGTGGTTGCCTGGCAAAGTGCCGATGAAAAACCTGAGTGTTATGATATTGACCAATCTAAAACATTCCTTTT
CTX_OROB_PARTIAL 1431 bp  643 ATGAACTTGGTTGTGTCTTGGCAGGGATCCGACGAGAATCCCCAATGCCAAGACATTAGAAAGGCCAATACGTTAGTCTT
CTX_SAFF 3147 bp         1111 TGGAATTTGGCTGTCGCTTGGCAGGGTACCGACGAGAAGTCTCAGTGTCAAGACATCGGAAAGGCCAATACTGTTGTATT
CTX_SOBS 3156bp          1111 TGGAATTTAGCCGTCACTTGGCAGGGTACCGATGAGAAGTCTCAGTGTCAAGACATCGGAAAGGCCAATACCGTTGTCTT
CTX_Sarc 2907 bp          568 TGGAACTTGATCGTGACGTGGCAGGGTTCCGCCGAGAAGCCCCAGTGCCATAACATTCGAGAAGCCAATACTTTTGTCTT
CTX2_Dopa 3123 bp        1084 ACATCTTTGCTGATAACATGGCAAGGTGTAAATGAAACATCTCGATGTAATCGAATGCTAAAAGCAAATACAATTATCTT
CTX1_Dopa 3123 bp        1117 GTGAACTTGGTCGTCACTTGGCAGGGTTCAAATGAAAGATCCCAATGTCGAGAGATTGGGAAAGCCAATACTTTTCTTTT

                         1281 ......1290......1300......1310......1320......1330......1340......1350......1360
CTX1_SEPES 3159 bp       1191 CCTTACTATATGTAAGTCGTGTCATCAGAGTCACGTTTTTACACCTAGCAACATGTTAAACAAAAATACATGCCCTAATA
CTX2_SEPES 3129 bp       1170 CCTTGACATATGTAAAAGGTGCCCCAAAACATATATGTACTCTCCTAAAAATTCATTGTCAAGTATTAAGTGCCCGGATG
CTX1_DPEA 3156 bp        1191 CCTTAGTATATGTAAGTCGTGTGAACGGAGTCACGTTTTGATTTCCGAGAACATGCAAAGCAAAAATAAATGTCCTAAAG
CTX2_DPEA 3123 bp        1164 CCTTGATATGTGTAAAAGATGTTTGAAAACATACCTGTACGCCCCTGAAAATATGTTGACAAGTATTAAGTGTCCTAAAG
CTX1_SBAN 2946bp         1188 CCTTACTATTTGTAAGTCATGTCATAATAGTCACGTTTTCACTCCTAGTAACATGTTAGACAAAAATAAATGCCCCAAAG
CTX2_SBAN 2286 bp        1170 CCTTGACCTCTGTAAAAAGTGTCCCAAAACATATATGTACTCTTCTAAAAATATGCTTTCAAGTATTAAGTGCCCGAAAG
CTX_EBER 3159 bp         1194 CCTAAGTATCTGCAAGAGGTGCCATACCAGTCGCGTCGCCATTTCCGGAGACATGCTCAGCATGAACAAATGTCCAGCAA
CTX2_SOFF 3129 bp        1170 TTTTGATATTTGTAAAAGGTGCACTGAAACATATATATACTCTCCTAAAAATACTTTGTCAAGTATAAAATGCCCAAGCG
CTX1_SOFF 3141bp         1173 CGTTAGTATATGCAAGTCGTGTCATAATAGTCACGTTTTTACTCCTAGCAACATGTTAGACAAAAATAGTTGCCCCCAAA
CTX_ESCO 3159 bp         1194 CCTAAGTATCTGCAAGAGGTGCCATACCAGTCGCGTCGCCATTTCCGGAGACATGCTCAGCAAGAACAAATGTCCAGCAA
CTX2_SLES 3132 bp        1173 CCTTGACATTTGTAAACGGTGTGTGAAAACATACCTGTACGTTCCTCAAAATATGTTGTCAAGTATGAAGTGCCCGCAAG
CTX1_SLES 3159 bp        1194 CCTTAGTATATGTCCGTCGTGTCAAGATAGTCACGTTTTTATTTCCGACAACATGCTAAGCAAAAATAAATGCCCTAAAG
CTX_TRHO 3150 bp         1191 CATTGACATCTGTAAATCGTGCCAAAAATCATACCTGTACGCTTCCAGAAACATGTTGGCGAGTATTAAGTGCCCCAAAG
CTX_SATL 3147 bp         1191 CCTGAGTATCTGCACGGCGTGCCATACCAGTCGCGTCTCCATTTCCAAAGACATGCTCAGCAAGAACAAATGCCCAGCAG
CTX2_SLYC 3129 bp        1170 CCTTGACATGTGTAAAAGGTGCCCCCAAACATATATGTACTCTCCTAAAAAGTCATTGTCAAGTATTAAGTGCCCGAATG
CTX1_SLYC 3159 bp        1191 CCTTACTATATGTAAGTCGTGTCATCAGAGTCACGTTTTCACTCCTAGCAACATGTTAAACAAAAATAAATGCCCTAATG
CTX_EPAR 3150 bp         1188 TCTAAGTATCTGCAAGGCGTGCCATACGAGTCGCGTCGCCATTTCCAAAGACATGCTCAGCAAGAACAAATGTCCAGCAG
CTX2_SPHAR 3129 bp       1170 CCTTGACATGTGTAAAAGGTGCCCCCAAACATATATGTACTCTCCTAAAAAGTCATTGTCAAGTATTAAGTGCCCAAATG
CTX1_SPHAR 3159 bp       1191 CCTTACTATATGTAAGTCGTGTCATCAGAGTCACGTTTTCACTCCTAACAACATGTTAAACAAAAATAAATGCCGTAATG
CTX_WSCI_PARTIAL 1653 bp  393 TCTAAGTATTTGCAAGTCGTGTAAAAATAGTCACACTTTTGTTCCTAAGAATATGATAAGCAAAACTAAATGCCCTAAAA
CTX_IHAL 3156 bp         1200 TATCGGCATTTGTCGTGGCTGTCAGAACAGTCATGTTTCCATCAACGAAGACATGTTGAGTAAAAATAAATGTCCGAAGA
CTX2_XNOT 3183 bp        1224 AATTGAAACTTGTAAAAGGTGCAAAAACTCTTATTTGTACGGTTCAAAAACCATGCTGACTAGTTCAAAATGTCCGACAA
CTX1_XNOT2 3150 bp       1194 TCTTAGCATTTGTAAACGTTGTCAAAACAGTCATGTTTCCATAAACGAAAACATGTTGAATAAAAATAAATGTCCAAAAA
CTX_OROB_PARTIAL 1431 bp  723 CCTTAGTATATGTAAGTCGTGCCAAAATAGTCGCACTTATATTCCCAGCAACATGCTTAGCAACAATAAATGCCCCCCAA
CTX_SAFF 3147 bp         1191 CCTGAGTATCTGCACGGCGTGCCATACCAGTCGCGTCTCCATTTCCAAAGACATGCTCAGCAAGAACAAATGCCCAGCAG
CTX_SOBS 3156bp          1191 CCTAAGTATCTGCAAGGCGTGCCATACCAGTCGCGTCTCCATTGCCAAAGACATGCTCAGCAAGAACAAATGCCCAGCAG
CTX_Sarc 2907 bp          648 TCTTAGCATAGGTAAGTCGTGTCAAATTAGTCGCACTTATATTCTAAAAAATATGCTAAGCAAAAATAATTGCCCTAAAA
CTX2_Dopa 3123 bp        1164 CCTTGACATGTGTAAAAGGTGTTTGAAAACACACCTGTACGCCCCTGAAAATATGTTGACAAGTATTAAGTGTCCTAAAG
CTX1_Dopa 3123 bp        1197 CCTTAGTATATGTCAGTCGTGTGAACAGAGTCACGTTTTGATTTCTGAGAACATGCAAAGCAAAAATAAATGTCCTAAAG

                         1361 ......1370......1380......1390......1400......1410......1420......1430......1440
CTX1_SEPES 3159 bp       1271 ATCAATACCCACAGGTGAAAGCATTTATCGATCGACGAGAGCCTTTTCGTGATGAAATTCAGAGGAAGAAATCGGATGTC
CTX2_SEPES 3129 bp       1250 AGAAGTATCCAAGGCTGAAAAAATTCATCGACAAACGATTTCCTGATGGCACGGAAAGTC-----CGGGCC-GGAGTCTC
CTX1_DPEA 3156 bp        1271 AAAACTATCCTCAAGTGAAAGCTTTTATCGATCAAAGAGGCCCGGAAGTTGACAAAAATG---AGAAGAAACATGATGCC
CTX2_DPEA 3123 bp        1244 AGAGGAACCCAGATATTAAGAATTACATCGACAAAACTTTTCCTGGGGGTACGGAAATTC-----CAGGCA-GAAGAATC
CTX1_SBAN 2946bp         1268 ATCGCTACCCACAAGTGAAAGCTTTAATCGATCAACACGAGCCTCAGCGTAACGAAAGGGAGAGGAAGGGGGTCGATAGC
CTX2_SBAN 2286 bp        1250 AGAAGTATCCGAATCTGAAGAAATTCATCGACAAACAATTTCCTGACCGTACAGAAAGTC-----CGGACA-GAAGTCTC
CTX_EBER 3159 bp         1274 AAACTTATCCAGAAGTAAAAGCTTTTATCAATCAGAGAGGATCAAATCTTGTTAAGAATG---AAAGGAAACACGATGCC
CTX2_SOFF 3129 bp        1250 AGAGGTATCCAAAGCTGAGGAAATTTATAAACAAACGATTTCCTGATGGCACGGAAAGCC-----CGGGCA-GAATAACC
CTX1_SOFF 3141bp         1253 ATAACTACCCACAGGTGAAAGCAATTATCAATCCACGAGAACCTCAGTTTGAAGAAAGCGACAGGAAGAAACTTGATGTC
CTX_ESCO 3159 bp         1274 AAACTTATCCGGAAGTAAAAGCTTTTATCAATCAAAGAGGATCAAATCTTGTTAAGAATG---AAAGGAAACACGATACC
CTX2_SLES 3132 bp        1253 AGAGGTACCCAATTATTAAAGATTTCATCGACAAAAGTTTTCCTGATGGTACGGAAAGTC-----CAGGCA-GAAGAATC
CTX1_SLES 3159 bp        1274 AAAACTATCCACAAGTAAAAGATTTTATCGATCAAAGAGGACCGGAACTTGACAAAAACG---GGAAGAAATATGATGCC
CTX_TRHO 3150 bp         1271 AGAGGTATCCGAAAATAAAAAAATTCATAGACAAAAAATTTCCAAACGATACAAGAAGTC-----CAAGCG-GAAAAATC
CTX_SATL 3147 bp         1271 AGACTTATCCGGAAGTAAAAGCCTTTATCAATCAGAGGGGCTCACATCTTGTTAAGAATG---AAAAGGAATACGATACC
CTX2_SLYC 3129 bp        1250 AGAAGTATCCAAAACTGAAAAAATTCATCGACAAACGATTTCCTGATGGCACGGAAAGTC-----CGGGCC-GAAGTATC
CTX1_SLYC 3159 bp        1271 ATCAATACCCACAGGTGAAAGCATTTATCGATCGACGAGAGCCTATGCGTAGAGAAAGGGAGAGGAAGAAAACCGATATC
CTX_EPAR 3150 bp         1268 AAACTTATCCAGAAGTGAAAGCTTTTATCAATCAGAGAGGATCACATCTTGTTAAGAATG---AAAAAGAACACGATGCC
CTX2_SPHAR 3129 bp       1250 AGAAATATCCAAAGCTGAAAAAATTCATCGACAAACGATTTCCTGATGGCACGGAAAGTC-----CGGGCC-GAAGTATC
CTX1_SPHAR 3159 bp       1271 ATCAATACCCACAGGTGAAAGCATTTATCGATCGACGAGAGCCTATGCGTGAAGAAATAGAGAAGAAGAAAACCGATATC
CTX_WSCI_PARTIAL 1653 bp  473 AGAGATACCCTGAAGTGAAAGCTTTGATCGACGAAAGAGGGCCGCATCTTGATGAAAATG---GAGAAGGACACGACGCC
CTX_IHAL 3156 bp         1280 ATCGTTACCCACGGGTGAAGGCATTTATCGACAGGCAAGGACCGGAACTTGATAGAAATA---GTAAGGGATACGATGCT
CTX2_XNOT 3183 bp        1304 ACACTTTTCCAAATATCAAGAGTTTAATTGATGCAAGGACTCCTCATGGGACAAGAAGTT-----TAAATG-ACGAGATT
CTX1_XNOT2 3150 bp       1274 ATCTTTATCCGCGGGTGAAAGCATATATTAACAGAGAAGGGCCAGAACTTGATAAAGATG---GTAAGAAATACGATGCA
CTX_OROB_PARTIAL 1431 bp  803 AACGCAATCCTCAAGTAAAAGCTTTCATCGATCAAAAAGGTTCGCATCTTTATAGAAATA---TGAAGAAGAAGGATGTG
CTX_SAFF 3147 bp         1271 AGACTTATCCGGAAGTAAAAGCCTTTATCAATCAGAGAGGCTCACATCTTGTTAAGAATG---AAAAGGAATACGATACC
CTX_SOBS 3156bp          1271 GAACTTACCCGGAAGTGAAAGCCTTTATCAATCAGAGAGGATCACATCTCGTTAAGAATG---AAAAGGAATACGATACC
CTX_Sarc 2907 bp          728 ACGTAAACCATCAAGTGAAAGCTTTAATCGATGAAAGAGGCCCACATCCTGACAAAAATC---GGAAGGAACATGATGTC
CTX2_Dopa 3123 bp        1244 AGAGGAACCCCGATATAAAGAATTACATCGACAAAAGTTTTCCTGGGGGTACGGAAATTC-----CAGGCA-GAAGAATC
CTX1_Dopa 3123 bp        1277 AAAACTATTCTCAAGTGAAAGCTTTTATCGATCAAAGAGGCCCGGAAGTTGACAAAAATC---GGGAGAAACATGATGCA

                         1441 ......1450......1460......1470......1480......1490......1500......1510......1520
CTX1_SEPES 3159 bp       1351 TTCTGGGTAGCTGCAGGATTCAAAGCCCCAGGTAACCCATGTAATCATGGCTGTAACGGTCATGGCGAGTGTAAAGTGGT
CTX2_SEPES 3129 bp       1324 CACTGGATTGCGGCCGGTTTTAAACCTAAAAAGGAGCCTTGCAATAATGCTTGTAATAATCATGGTCAATGTAAAATAAT
CTX1_DPEA 3156 bp        1348 TTCTGGGTAGCTGCAGGATTCAAATCATCTGGTGACGCATGTAGTCATCGCTGTAACAATCATGGTGAGTGTAGAATGGT
CTX2_DPEA 3123 bp        1318 CATTGGGTTGCGTCTGGTTTTAAACAGCAAAGGAACCCTTGCAGAAATGCCTGTAATAACCAGGGTCAGTGTAAAGTAAT
CTX1_SBAN 2946bp         1348 TTCTGGGTAGCTGCGGGATTCAAATCCTATGGCAACCCATGTAATCATCGCTGTAACGATCATGGCGAGTGTAAAGTAGT
CTX2_SBAN 2286 bp        1324 CACTGGATTGTAGCTGGTTTTAAATCCCGAAAGAATCCTTGCGAAAATGCCTGTAATAATCATGGTCAGTGTAAAGTAAT
CTX_EBER 3159 bp         1351 TTCTGGGTTGCAGCAGGATTCAAATCGCCTGGAAACCCATGTGACCATCGATGCAACAACCATGGCGAGTGTAAAATAAT
CTX2_SOFF 3129 bp        1324 CACTGGATTGCGGCTGGTTTTAAAGGTCGAAAGCAGCCTTGCAAAAATGCCTGTAATAATCATGGTCAGTGTAAAGTAAT
CTX1_SOFF 3141bp         1333 TTCTGGGTAGCTGCAGGATTCAAAGCCCCCGGTAACCCATGTAGTCATCGCTGTAACGGTCATGGCGAGTGTAAAGTTGT
CTX_ESCO 3159 bp         1351 TTCTGGGTTGCAGCAGGATTCAAATCGCCTGGAAACCCATGTGACCATCGATGCAACAACCATGGCGAGTGTAAAATAAT
CTX2_SLES 3132 bp        1327 TATTGGATTGCCGCTGGTTTTAAACGGCAAAGGAACCCTTGCAAAAACGCCTGCAATAAAAATGGTCAATGTAAAGTAAT
CTX1_SLES 3159 bp        1351 TTCTGGGTAGCTGCAGGATTCAAATCACTCGGTAATTCATGTAATCATCACTGTAACAATCATGGTGAATGTAGAGTGGT
CTX_TRHO 3150 bp         1345 CACTGGATTGCAGCTGGTTTTAAACCCCGGGGTGACCCTTGCAAAAATGCCTGTAATAACAACGGTCAGTGTAAAATAAT
CTX_SATL 3147 bp         1348 TTCTGGGTTGCAGCAGGATTCAAATCGCCTGGAAACCCATGCGACCATCGATGCAACGACCATGGCGAGTGTAAGATAAT
CTX2_SLYC 3129 bp        1324 CATTGGATTGCGGCTGGTTATAAAGCTGAAAAGGAGCCTTGCAAAAATGCTTGTAATAATCATGGTCAGTGTAAAGTAAT
CTX1_SLYC 3159 bp        1351 TTCTGGGTAGCTGCAGGATTCAAAGCCCCAGGTAACCCATGTAATCATCGCTGTAACGACCATGGCGAGTGTAAAGTGGT
CTX_EPAR 3150 bp         1345 TTCTGGGTTGCAGCAGGATTCAAATCGCCCGGAAACCCATGTGACCATCGATGCAACAACCATGGCGAGTGTAAAATAAT
CTX2_SPHAR 3129 bp       1324 CAATGGATTGCGGCTGGTTTTAAAGCTGAAAAGGAGCCTTGCAAAAATGCTTGTAATAATCATGGTCAGTGTAAAGTAAT
CTX1_SPHAR 3159 bp       1351 TTCTGGGTAGCTGCAGGATTCAAAGCGTCAGGTAACCCATGTAATCATCGCTGTAACGGTCATGGCGAGTGTAAAGTGGT
CTX_WSCI_PARTIAL 1653 bp  550 TTCTGGATCGCTGCTGGATTCAAAGTCTCTAGGAATCCGTGTTCCCATCGCTGTAATAATCACGGTCAATGTAAAGTGGT
CTX_IHAL 3156 bp         1357 TTCTGGTTGGCTGCAGGTTTTAGCAGCGAAACGAACCCTTGTAGCCATCGTTGTAACAATAGGGGCGAATGCAAAGTTGT
CTX2_XNOT 3183 bp        1378 TTTTGGATAGCTGCTGGTTTTAAACCTCGTAAGAATCCTTGCAAAAATGCTTGCAATAACAATGGCCGTTGTAAAGCGAT
CTX1_XNOT2 3150 bp       1351 TTCTGGTTGGCTGCAGGCATTAGTAACGAAAGGAACCCTTGTAGCCATCGTTGCAACAATCGCGGAGAATGTAAAGTCGT
CTX_OROB_PARTIAL 1431 bp  880 TTCTGGGTCGCTGCCGGTTTCAAAGCTTCTGGGAACCCATGTTCCCATCGATGTACCAATCGTGGTAAATGTAAAGTGGT
CTX_SAFF 3147 bp         1348 TTCTGGGTTGCAGCAGGATTCAAATCACCTGGAAACCCATGCGACCATCGATGCAACGACCATGGCGAGTGTAAGATAAT
CTX_SOBS 3156bp          1348 TTCTGGGTTGCAGCCGGATTCAAATCGCCTGGAAACCCGTGCGACCATCGATGCAACAACCATGGCGAGTGCAAGATAAT
CTX_Sarc 2907 bp          805 TTCTGGGTTGCTGCAGGATTCAAGTCCTCTGGCAACCCATGTGATCATCGTTGTAACAATCATGGTAAGTGTAAAGTGGT
CTX2_Dopa 3123 bp        1318 CATTGGATTGCGTCTGGTTTTAAACAGAAAATGAAGCCTTGCAGAAATGCCTGTAATAACCAGGGTCAATGTAAAGTAAT
CTX1_Dopa 3123 bp        1354 TTCTGGGTAGCTGCAGGATTCAAATCATCTGGTGACGCATGTGGTCATCGTTGTAACAATCATGGTGAGTGTAGATTGGT

                         1521 ......1530......1540......1550......1560......1570......1580......1590......1600
CTX1_SEPES 3159 bp       1431 CCCTTACACCGATCAATTCCAATGCTTTTGTCACGGTAATTACGAGGGGAAAATGTGTCAGAAAAAGATACAAATGAAAC
CTX2_SEPES 3129 bp       1404 TCCATACACTGATCAAATTCAATGTTTTTGTTACGCGAATTATGTTGGACACAATTGCGAAACAGAAATAAGCGGAGACA
CTX1_DPEA 3156 bp        1428 TCCTTACACCGATCAATTCCAATGTTTCTGTCAAACTAATTATGAAGGAGAAAAGTGTGAAACAAAAATAGAAATAAACC
CTX2_DPEA 3123 bp        1398 TCCATACACTAGTCAAATTCAATGTTTTTGTTACGCCAATTATATAGGGAAAAACTGCGAAACAGAAATAGTTGAAGAGA
CTX1_SBAN 2946bp         1428 CCCATACACAGATCAATTTCAATGTTTTTGTCATGACAATTACGAAGGGAAAATGTGTGATAGAAAGTTACAAATGAAAC
CTX2_SBAN 2286 bp        1404 TCCTCATACTGATCAAATTCAGTGTTTTTGCGACGCAAATTATGTAGGACACAATTGCGAAACAGAAATAGTTGCAGATA
CTX_EBER 3159 bp         1431 TCCATATACCGATCAATTCCAGTGTTTTTGTCAATCGAGTTATGAAGGGAAAACATGTGAAGAAAAAATACAAATGAATA
CTX2_SOFF 3129 bp        1404 TCCATACACGGATCAAATTCAGTGTTTTTGCTACGCCAATTACGTAGGAGACAATTGCGAGACGGAAATAGCTGTAGACA
CTX1_SOFF 3141bp         1413 CCCTTACACAGACCAATTCCAATGCTTCTGTCACGATAGCTACGAAGGGAAAATGTGTGACCAAAAGATACAAATGAAAC
CTX_ESCO 3159 bp         1431 TCCATATACCGATCAATTCCAGTGTTTTTGTCAATCGGGTTATGAAGGGAAAACATGTGAAGAAAAAATACAAATGAATA
CTX2_SLES 3132 bp        1407 TCCATACACTCATCAAAGTCAATGTTTTTGCTACGCTAATTATATGGGGGAGAATTGCGAAACAGAAATAGTTCAAGAGA
CTX1_SLES 3159 bp        1431 TCCTTACACAGACCAATTCCAGTGTTTCTGCCAAGCTGATTATGAAGGAGAAAAGTGTGACAAAAAAATAGAAATCAACC
CTX_TRHO 3150 bp         1425 CCCATATACTGACCAAGTGCAGTGTTTTTGCTATGCCAGTTATGCAGGGGAGAATTGCGAAACTGAAATAGTCGAGGACA
CTX_SATL 3147 bp         1428 TCCGTATACCGATCAATTCCAGTGTTTTTGCCAATCGAATTATGAAGGCGAAAAATGTGAACGAAAAATACAAATGAATA
CTX2_SLYC 3129 bp        1404 TCCATACACTGATCAAATTCAATGTTTTTGCTACGCGAATTATGTAGGATACAACTGCGAAACAGAAATAAGCGGAGACA
CTX1_SLYC 3159 bp        1431 CCCTTACACAGATGAATTCCAATGCTTTTGTCACAATAATTACGAAGGGAAAATGTGTCACAAAAAGATACAAATGAAAC
CTX_EPAR 3150 bp         1425 TCCGTATACCGATCAATTCCAGTGTTTTTGTCAATCGAGTTATGAAGGGAAAAAATGTGAACGGAAAATACAAATGAATA
CTX2_SPHAR 3129 bp       1404 TCCATACACTGACCAAATTCAGTGTTTTTGCTACGCGAATTATGTCGGATACAATTGCGAAACAGAAATAAGCGGAGAAA
CTX1_SPHAR 3159 bp       1431 CCCTTACACAGATCAATTCCAATGCTTTTGTCACAATAATTACGAAGGGAAAATGTGTGACAAAAAGATACAAATGAAAC
CTX_WSCI_PARTIAL 1653 bp  630 TCCATACACGGATCAATTCCAATGTTTCTGTTATGCTGATTACGAAGGTCAGATGTGTGAGAGGGAAATAAAGATGAATC
CTX_IHAL 3156 bp         1437 TCCTTACACAAACCAATTCCAATGTTTCTGTCGAAATAATTACGAAGGTGAAAAATGCGAATCGAAAATTGAAATAAAAA
CTX2_XNOT 3183 bp        1458 CCCGTTCACCAACGAAATTCAATGCTTTTGTTTTGCTAATTTTGATGGAGAGCAATGCGAGATTGAAATCGTCGAAAATA
CTX1_XNOT2 3150 bp       1431 TCCTTACACAAACCAATTCCAGTGTTTTTGTCAAAATAATTACGAAGGCGAAAAATGTGAATCGAAAATAGAAGTAAAAA
CTX_OROB_PARTIAL 1431 bp  960 TCCTTACACGAACCAATTCCAATGTTTCTGCCAAGCGAGTTATGAAGGCGAAAAGTGTGAGAGTCAAATAGAGATGAACG
CTX_SAFF 3147 bp         1428 TCCGTATACCGACCAATTCCAGTGTTTTTGCCAATCGAATTATGAAGGCGAAAAATGTGAACGAAAAATACAAATGAATA
CTX_SOBS 3156bp          1428 TCCGTATACCGACCAATTCCAGTGTTTTTGTCAACCGCGTTATGAAGGAGAAAAGTGTGAACGAAAAATACAGCTGAATA
CTX_Sarc 2907 bp          885 TCCTTACACGGACCAATTCGAATGTTTCTGCCAAGCAAGCAATGAAGGGGAAATGTGTGAAAGTAAGATAGAAATGAACC
CTX2_Dopa 3123 bp        1398 TCCATACACTAGTCAAATTCAATGTTTTTGTTACGCCAATTATATAGGGAAAAACTGCGAAACTGAAATAGTTGAAGAGA
CTX1_Dopa 3123 bp        1434 TCCTTACACCGACCAATTCCAATGTTTCTGCCAAACTAATTATGAAGGAGAAAAGTGTGAAACAAAAATAGAAATAAACC

                         1601 ......1610......1620......1630......1640......1650......1660......1670......1680
CTX1_SEPES 3159 bp       1511 GAGATATTTCGAAACTCATTTCTGACCTTCAAACCGGATATAAAAACGCATTTAACGTTCCTTCGCTGACAAATATTTTG
CTX2_SEPES 3129 bp       1484 CAAATTTTGAGAAAATGGTGGTAGATCTGCAAATGGTTTACACCGATGTATTTAAAACTCCTATGATCTCAAGCGTTTTG
CTX1_DPEA 3156 bp        1508 ACGATATTGTGGAACTCGTTTCTGATCTTCAGTTGGGCTATAAAAATGCATTTAACGCTCCTTCATTGGTAAATATTTTG
CTX2_DPEA 3123 bp        1478 CAAACATTGAGAAAATGGTGTTGAATTTGCAACGCATATACAAGGATGTATTTAAAATTCCCTCGATTTCAAGCATTTTG
CTX1_SBAN 2946bp         1508 GAAATATCTTAAAACTAGTTTCTGATCTTCAAACAGGTTATAAAAATGCATTTAAAGTCCCTTCCCTAACAAATATTTTG
CTX2_SBAN 2286 bp        1484 CAAACATTGAGAAAGTGGTGGTAGATTTGCAAATGGTTTACACTGATGTATTTAAGACTCCTTCCATTTCAAACGTTTTG
CTX_EBER 3159 bp         1511 AAAACATTATGAAACTTGTTTCGGATCTACAAACTGGCTATAAAGATGCGTTTAAAGTTCCTTCGTTGGCAAATATTTTG
CTX2_SOFF 3129 bp        1484 CAAACATTGAGAAAATGGTGGTAGATTTGCAAATGGTTTACACCGATGTATTTAAAACTCCTTCGATCTCAAGCGTTTTG
CTX1_SOFF 3141bp         1493 GAAATATTTTAAAACTAATTTCTGATCTTCAAACCGGTTATAAAAATGCATTTAAGGTTCCTTCGTTGACGAATGTTTTG
CTX_ESCO 3159 bp         1511 AAAACGTTATGAAACTTGTTTCGGATCTACAAACTGGCTATAAAGATGCGTTTAAAGTTCCTTCGTTGGCAAATATTTTG
CTX2_SLES 3132 bp        1487 CAAACATTGAGAAAATGTTGTTGGATTTGCAATACGTATACAGGGATGCATTTAAAATTCCCTCAATCTCAAATGTTTTG
CTX1_SLES 3159 bp        1511 AAGATATTATCAAACTCGTTTCTGATCTTCAATTAGGCTATAAAAATGCATTTAATGCTCCCTCATTGGTAAATGTTATG
CTX_TRHO 3150 bp         1505 AAACCATCGAGAAAATGGTGTTGGACTTGCAACTCGTATACAGTGACGTTTTTAAAACTCCTTCAATATCAAGCGTTTTG
CTX_SATL 3147 bp         1508 AAAACATTATGAAGCTCGTGTCGGATCTTCAAACCGGCTATCAAAATGCGTTCAAGGTTCCGTCGTTGGCAAATGTTTTG
CTX2_SLYC 3129 bp        1484 CAAATTTTGAGAAAATGGTGGTAGATTTGCAAATGGTTTACACCGATGTATTTAAAACTCCTACAATCTCAAGCGTTTTG
CTX1_SLYC 3159 bp        1511 AAGATATTTCGAAACTCATTTCTGATCTTCAAACTGGATATAAAAATGCATTTAGCGTTCCTTCGTTGACAAATATTTTG
CTX_EPAR 3150 bp         1505 AAAGCATTATGAAACTTGTTTCAGATCTACAAACTGGCTATCAAGATGCGTTTAAAGTTCCTTCGTTGGCAAATATTTTG
CTX2_SPHAR 3129 bp       1484 CAAATTTTGAGAAAATGATGGTGGATTTGCAAATGGTTTACACTGATGTATTTAAAACCCCTACGATCTCAAGCGTTTTG
CTX1_SPHAR 3159 bp       1511 GAGATATTTCGAAACTCATTTCTGATCTTCAAAGCGGATATAAAAATGCATTTAGCGTTCCTTCGTTGACAAATGTTTTG
CTX_WSCI_PARTIAL 1653 bp  710 AGGATATCACGAAACTTGTTACAGATCTCCAATCGGGATACAAGGATGCTTTCAAAGTTCCATCGATGGCGAATATTTTG
CTX_IHAL 3156 bp         1517 ATGATATTCAAAAAATGCTCTCCGATTTGCAAAATGGCTATAAAAGTGCTTTCAAGGTTTCTTCGTTGGCCAATATTTTA
CTX2_XNOT 3183 bp        1538 CAAGCTTTGAAAAAATGATATTAGATTTGGAATCGATTTACAGCGATGTTTTTAAAACCCCTTCGATTTCAAGTGTTTTG
CTX1_XNOT2 3150 bp       1511 ATGATATCCAAAAATTGCTTTCCGATTTGCAAAACGGCTATAAAAGTGCTTTCAAGGTTTCCTCGTTGGCCAATATTTTA
CTX_OROB_PARTIAL 1431 bp 1040 ATGACATCACGAAACTCGTCTCCGATCTTCAATCGGGATATAAGAGCGCATTTAAGGCCCCTTCGTTGACAAACGTTTTG
CTX_SAFF 3147 bp         1508 AAAACATTATGAAGCTCGTGTCGGATCTTCAAACCGGCTATCAAAATGCGTTCAAGGTTCCGTCGTTGGCAAATGTTTTG
CTX_SOBS 3156bp          1508 AGAACATTATGAACCTCGTGACCGATCTGCAAAGCGGCTATAAAAACGCATTTAAGGTCCCTTCGTTGGCAAACGTTTTG
CTX_Sarc 2907 bp          965 AAGCTATTACGAAACTGCTTTCTGATC-------GGGCTATAAGTGTGCATTTAAGGTTCCTTCAATGGCAAATATTTTG
CTX2_Dopa 3123 bp        1478 CAAACATTGAGAAAATGGTGTTGAATTTGCAACGCATATACAGGGATGCATTTAAAATTCCCTTGATTTCAAGCATTTTA
CTX1_Dopa 3123 bp        1514 AAGATATTGTGGAACTCGTTTCTGATCTTCAATTGGGCTATAAAAATGCATTTAACGCTCCTTCATTGGTAAATATTTTG

                         1681 ......1690......1700......1710......1720......1730......1740......1750......1760
CTX1_SEPES 3159 bp       1591 ATCCAAGGGGAAAATCTAGCCAAACAACTTAAGAAGATGATACAAAGGATTGATAATCAGTTTGAGTTGACACACATTTT
CTX2_SEPES 3129 bp       1564 ATTCAAGCAGAAGATTTGTCGAATCAGCTGAGGAATATGCTACAAAGGATAGATCGACAATTTGAACTGACTCAAATCCT
CTX1_DPEA 3156 bp        1588 ATCCAAGGAGAAAATTTATCAAATAAATTAAAGAGAATGATGCAAAGGATCGACAGTCAATTTGAGTTGACACAAATGTT
CTX2_DPEA 3123 bp        1558 ATTCAAGGAGAAAATTTATCGAAGCAAATGAAGTATATGATGCAAAGAATAGACAAACAATTTGAATTGACACATATTCT
CTX1_SBAN 2946bp         1588 ATCCAAGGAGAAAATTTAGCAAAGCAACTTAAAAAAATGATGCAAAGGATTGACAGTCAGTTTGAGTTGACAAATATTTT
CTX2_SBAN 2286 bp        1564 ATTCAAGCAGAAGATTTGTCGAATCAGCTGAGGAACATGCTACAGAGGATAGATCGACAATTTCAACTGACCCAAATCCT
CTX_EBER 3159 bp         1591 ATTCAATCAGAGAATCTATCGAAGCAGTTAAAGAAAATGATGCGACGGATTAATAACCAGTTCGAGTTGACACAGATTTT
CTX2_SOFF 3129 bp        1564 ATACAAGCAGAAGATTTATCGAATCAACTGAAGAATATGTTGCAAAAGATAGATCGACAATTTGAACTGACACACATACT
CTX1_SOFF 3141bp         1573 ATCCAAGGGGAAAATTTGGCAAAACAACTTAAGAAGATGATGCAAAAGATTGACAATCAGTTTGAATTGACACAGATTCT
CTX_ESCO 3159 bp         1591 ATTCAATCAGAGAATCTATCGAAGCAGTTAAAAAAAATGATGCGACGGATTAATAACCAGTTCGAGTTGACACAGATTTT
CTX2_SLES 3132 bp        1567 ATCCAAGGAGAAGATTTATCGAAGCAAATGAAGTTTATGATTCAAAGAATAGACAAACAATTTGAATTAACACATATTCT
CTX1_SLES 3159 bp        1591 ATCCAAGGAGAAAATTTAGCAAAGAAATTAAAAAGAATGATGCAAAGGATCGACAATCAATTTGAGTTAACACAAATTTT
CTX_TRHO 3150 bp         1585 ATTCAAGGAGAAGATTTGTCAAAGCAGCTACGGAATTTGATTCAAAGGATAGATCAACAATTTGAAATGACGCATAGCCT
CTX_SATL 3147 bp         1588 ATTCAATCAGAGAATCTATCGAAGCAGTTAAAGAAAATGATGCGACGAATTAATAATCAGTTCGAGTTGACACAGATTTT
CTX2_SLYC 3129 bp        1564 ATTCAAGCAGAAGATTTGTCGACTCAACTGAGGAATATGCTGCAAAGGATAGATCGACAATTTGAACTGACGCAAATCCT
CTX1_SLYC 3159 bp        1591 ATCCAAGGGGAAAATCTAGCCAAACAACTTAAGAAGATGATACAAAGGATTGACAATCAGTTTGAGTTGACACATATTTT
CTX_EPAR 3150 bp         1585 ATTCAATCAGAGAATCTATCGAAACAGTTAAAGAAAATGATGCGACGTATTAATAACCAGTTCGAGTTGACACAGATTTT
CTX2_SPHAR 3129 bp       1564 ATTCAAGCAGAAGATTTGTCGACTCAACTAAGGACTATGCTGCAAAGGATAGATCGACAATTTGAACTGACGCAAATCCT
CTX1_SPHAR 3159 bp       1591 ATCCAAGGGGAAAATATAGCCAAACAACTTAAAAAGATGATACAAAGGATTGACAATCAGTTTGAGTTGACACATATTTT
CTX_WSCI_PARTIAL 1653 bp  790 ATACAGGGTGAAAATCTTGGGAAGGAATTGAAGAAAATGATGCGAAGACTAAACAACCAGTTTGAGATGACACACATTTT
CTX_IHAL 3156 bp         1597 ACCCAAGCTGAGAATTTAGCGAAACAATTAAAGAAAATGATTCAAAGGATGGACACACATTTTGAACTGACTCACATTCT
CTX2_XNOT 3183 bp        1618 ATTCAAGGCGAAGACTTATCCAAACAACTTAAGAAAATGATTCAAAAGATAAACACACAATTTGAAATGACCAATATCCT
CTX1_XNOT2 3150 bp       1591 ACCCAGGCCGAGAATTTATCGAAACAATTAAAAAAAATGATTCAAAGGATGGATACACATTTTGAATTGACTCACATTCT
CTX_OROB_PARTIAL 1431 bp 1120 ATCCTGGGAGAAAATCTATCGAGCCAATTGAAGAAAATGCTGCGAATGATCGATAATCAATTCGAGTTGACGCAGATCCT
CTX_SAFF 3147 bp         1588 ATTCAATCAGAGAATCTATCGAAGCAGTTAAAGAAAATGATGCGACGAATAAATAATCAGTTCGAGTTGACACAGATTTT
CTX_SOBS 3156bp          1588 ATTCAAACAGAGAATCTATCGAAGCAGTTGAAGAAAATGATGCGACGAATAAACAATCAGTTCGAGTTGACGCAGATTCT
CTX_Sarc 2907 bp         1038 ATACAAGGAGAACATCTCGC-----AAATGAAGAAAATGATGCGAAGAATTAACAATCAATTTGAGTTGACACAAATGTT
CTX2_Dopa 3123 bp        1558 ATTCAAGGAGAAAATTTATCGAAGCAAATGAGGTATATGATGCAAAGAATAGACAAACAATTTGAATTGACACATATTCT
CTX1_Dopa 3123 bp        1594 ATCCAAGGAGAAAATTTATCAATGAAATTAAAGAGAATGATGCAAAGGATCGACAATCAATTTGAGTTGACACAAATGTT

                         1761 ......1770......1780......1790......1800......1810......1820......1830......1840
CTX1_SEPES 3159 bp       1671 ----------------------GGTAAAATATATCAGCGATTTGCAAAAACTGGATTATATACTCAAGATCAGCTTTAAT
CTX2_SEPES 3129 bp       1644 ----------------------GGTTAAATATATCAGTCCCTTGCAAAAAATCGACTACCTACTCGATCTCAGCTTCACG
CTX1_DPEA 3156 bp        1668 ----------------------GGTCAAATATATCAGGGATATGCAAAAATTGGATTATATTCTCAAACTTGGCTTTCTT
CTX2_DPEA 3123 bp        1638 ----------------------TGTTAAATATATAAACTCGTTGCAAAAATTCGACTATCTACTTGAGCTTAGCTTTGCG
CTX1_SBAN 2946bp         1668 ----------------------GGTAAAATATATCAAAGATTTGCAAAAACTGGATTATGTACTCAAGTTAAGCTTTAAT
CTX2_SBAN 2286 bp        1644 ----------------------GGTTAAATATATTAATCCTTTGCAAAAAATCGACTACCTACTCGATCTCAGCTTCTCG
CTX_EBER 3159 bp         1671 ----------------------GATCAAATATATCCGAGATCTACAAAAGTTGGATTACATATTAAAGCTCAGCTTTAAT
CTX2_SOFF 3129 bp        1644 ----------------------GGTTAAATATATCAACCCCTTGCAAAAAATCGACTATCTACTCGATCTCAGCTTCTCG
CTX1_SOFF 3141bp         1653 ----------------------GGTCAAATATATCAACGATTTGCAAAAGTTGGACTACATACTAAAGATCAGCTTTTAT
CTX_ESCO 3159 bp         1671 ----------------------GATCAAATATATCCGAGATCTACAAAAGTTGGATTATATATTAAAGCTCAGCTTTAAT
CTX2_SLES 3132 bp        1647 ----------------------GGTTAAATATATTAACTCGTTGCAAAAATTCGACTATTTACTTGAGCTTAGTTTTGCG
CTX1_SLES 3159 bp        1671 ----------------------GGTCAAATATATAAGAGATCTGCAAAAATTGGATTATATACTCAAACTTGGCTTTAAT
CTX_TRHO 3150 bp         1665 ----------------------GGTTAAATATATTAGTTCCTTGCAAAAGATCGACTATCTACTCGAGCTAAGCTTCGCT
CTX_SATL 3147 bp         1668 ----------------------GATCAAATATATCCGAGATCTGCAAAAGTTGGATTATATATTAAAGCTTAGCTTTAAT
CTX2_SLYC 3129 bp        1644 ----------------------GGTTAAATATATCAGCCCCTTGCAAAAAATCGACTACCTGCTCGATCTCAGCTTCACG
CTX1_SLYC 3159 bp        1671 ----------------------GCTAAAATATATCAGCGATTTGCAAAAACTGGATTATATACTCAAGATCAGCTTTAGT
CTX_EPAR 3150 bp         1665 ----------------------GATCAAATATATCCGAGATCTGCAAAAGTTGGATTATATATTAAAGCTCAGCTTTAAT
CTX2_SPHAR 3129 bp       1644 ----------------------GGTTAAATATATCAGTCCCTTGCAAAAAATCGACTATCTACTCGATCTCAGCTTCACG
CTX1_SPHAR 3159 bp       1671 ----------------------GCTGAAATATATCAGCGATTTGCAAAAACTGGATTATATACTCAAGATCAGCTTTAGT
CTX_WSCI_PARTIAL 1653 bp  870 ----------------------GGTCAAGTACATCGGTGACTTGCAAAAATTGGATTATATACTGAAACTCGGTTTTGAT
CTX_IHAL 3156 bp         1677 ----------------------CGTTAAATACATTCGAGATTTACAAAAACTCGATTATATTCTCAAACTTAGCTTTAAT
CTX2_XNOT 3183 bp        1698 ----------------------CATCAAATACATCCATTCATTACAAAAGCTCCATTATTTACTGGAACTGAATTTTAAT
CTX1_XNOT2 3150 bp       1671 ----------------------TGTTAAATACATTCGAGATCTGCAAAAATTGGATTATATTCTAAAACTTAGCTTCAAT
CTX_OROB_PARTIAL 1431 bp 1200 ----------------------GATCAAATACATCGGCGATCTACAAAAGTTGGATTATATCCTGAAGCTCAGCTTTGAC
CTX_SAFF 3147 bp         1668 ----------------------GATCAAATATATCCGAGATCTGCAAAAGTTGGATTATATATTAAAGCTTAGCTTTAAT
CTX_SOBS 3156bp          1668 ----------------------GGTCAAATATATCCGAGATCTGCAAAAGTTGGACTATATATTAAAGCTGAGCTTTAAC
CTX_Sarc 2907 bp         1113 GACCAAATGTTGACCAAATGTTGACCAAATATATCAACGATCTACAAAAATTGGATTATATCCTTAAGCTCAGCTTTGCT
CTX2_Dopa 3123 bp        1638 ----------------------TGTTAAATATGTAAACTCGTTGCAAAAATTCGACTATCTACTTGAGCTGAGCTTTGCG
CTX1_Dopa 3123 bp        1674 ----------------------GGTCAAATATATCAGGGATATGCAAAAATTGGATTATATTCTCAAAATTGGCTTTCTT

                         1841 ......1850......1860......1870......1880......1890......1900......1910......1920
CTX1_SEPES 3159 bp       1729 TACAGTAAAAAGAAAATTACTGTGGACGCCTTCAGTCGAAGAATGAAGGCATTCTTGTCACTTAATCCAGTTGACTTCAT
CTX2_SEPES 3129 bp       1702 TATAGGACAAAAGCAATCACGGTTGACGCCTACAACCGAAGAATGAAATCATTCTTATCACATAATGACGTCCCCTATAT
CTX1_DPEA 3156 bp        1726 TACAGCAAAAAGAGAATCAGTGTGGATGCTTTTAGTCGAAGAATGAAGAAATTCTTGTCCCTTAATTCAATTAACTTTAT
CTX2_DPEA 3123 bp        1696 TATCAGAAAAAAGAAATCACCTTAGATGTTTATAACTTAAGAATGAAATCTTTCTTATCGCATAATGATATATATTTTAT
CTX1_SBAN 2946bp         1726 TATAGTAAAAAGAAAATCACAGTTGATGCCTTTAGTCGAAGAATGAAGGCATTTTTGTCTCTTAATCCGGCTGATTTTAT
CTX2_SBAN 2286 bp        1702 TATAGAAGAAAAGCAATCTCTGTTGACGCCTACAATCGCAGAATGAAATCTTTCTTATCTCACAACGACGTCCCGTTTAT
CTX_EBER 3159 bp         1729 TATAGTAAACAAAAGATTAGTATTGACGCTTTCAATCGGAGGATGAAGACGTTTTTGTCTCTGAATCCAATAAATTTCAT
CTX2_SOFF 3129 bp        1702 TATAGAACGAAAGCCATCACGGTTGACGCCTACAGCCAAAGAATGAAATCCTTCTTATCGCATAACGACGTCCCCTTTAT
CTX1_SOFF 3141bp         1711 TACAGTCAGAGTAAAATTTCAGTGGATGCTTTCAGTCGAAGAATGAAGAGATTTTTATCTCTTAATCCAATTGACTTCAT
CTX_ESCO 3159 bp         1729 TATAGTAAACAAAAGATTAGTATTGACGCTTTCAATCGGAGAATGAAGACATTTTTGTCTCTGAATCCAATTAATTTCAT
CTX2_SLES 3132 bp        1705 TATAAAAAAAAGGCAATCACGCTAGACGTTTATAACTTAAGAATGAAATCATTTTTATCACATAACGACATCTACTTTAT
CTX1_SLES 3159 bp        1729 TACAGCAGAAAGAGGATTAGTGTGGATGCTTTTAGTCGAAGAATGAAGAAATTCTTGTCTCTCAATCCAATAAACTTTAT
CTX_TRHO 3150 bp         1723 TACAGAACGAAAGTGATCACGATCAACGCCTACAACCGAAGAATTAAATCGTTCTTATCGCATAACGACGTACCTTTCAT
CTX_SATL 3147 bp         1726 TATAGCAAACGGAAGATCAGCATTGACTCTTTCAATCGAAGGATGAAGACGTTTTTGTCGCTCAATCCGATCAACTTCAT
CTX2_SLYC 3129 bp        1702 TATAGGACTAAAGCAATCACGGTTGACGCCTACAACCGAAGAATGAAATCATTTTTATCACATAATGACGTCCCCTTTAT
CTX1_SLYC 3159 bp        1729 TACAGTAAAAAGAAAATTACTGTGGATGCCTTTAGTCGAAGAATGAAGGCATTCTTGTCACTTAATCCAATTGACTTCAT
CTX_EPAR 3150 bp         1723 TATAGTAAACAAAAGATTAGCATTGACGCTTTCAATCGGAGGATGAAGACGTTTTTGTCTCTGAACCCAATCAATTTCAT
CTX2_SPHAR 3129 bp       1702 TATAGGACTAAAGCAATCACGGTTGATGCCTACAACCGAAGAATGAAATCATTTTTATCACATAATGATGTCCACTTTAT
CTX1_SPHAR 3159 bp       1729 TACAGTAAAAAGAAAATTACCGTGGATGCCTTCAGTCGGAGAATGAAGGCATTCTTGTCACTTAATCCGATTGACTTCAT
CTX_WSCI_PARTIAL 1653 bp  928 TATAGCAAGAAGAAGATCAACGTAGACGCGTTCAGTCGTAGAATGAAGAAATTCCTCTCTCTCAATCCGATCAATTTTGT
CTX_IHAL 3156 bp         1735 TACAGCAAACGTAAAATCTCGATTGATTCTTATAATCGAAGAATGAAAAAATTCCTGGGTTTAAATCCAATTCATTTTAT
CTX2_XNOT 3183 bp        1756 TATCAGAAAAGAACAATTTCCATTGACGTTTTTAATTTGAAAATGAAGCATTTCCTTTTGCACAACGATGTAGCATTTAT
CTX1_XNOT2 3150 bp       1729 TATAGCAAACGTAAAATTTCGGTCGATTCTTACAACCGAAGAATGAAAAAGTTCCTGGCTCTTAATCCAATTCATTTCAT
CTX_OROB_PARTIAL 1431 bp 1258 TTCAGCAAGAAGAGGATCAGCGTGGATGCGTTCAGTCGTAGAATGAAGAAGTTCCTGTCTTTGAATCCGGTGAACTTCGT
CTX_SAFF 3147 bp         1726 TATAGCAAACGGAAGATCAGCATTGACTCTTTCAATCGAAGGATGAAGACATTTTTGTCGCTCAATCCGATCAATTTCAT
CTX_SOBS 3156bp          1726 TATAGCAAACGGAAGATCAGCATTGATGGGTTCAGTCGAAGGATGAAGGCGTTTTTGTCGCTCAATCCGATCGATTTCAT
CTX_Sarc 2907 bp         1193 TACGGCAAAAAGAAAATTAGTGTGGATTCGTTCAGTCGTAGAATGAAGAAGTCCTTGGCTCTCAATCCAATTAATTCCGT
CTX2_Dopa 3123 bp        1696 TATCAGAAAAAAGAAATCACCTTAGATGTTTATAACTTAAGAATGAAATCTTTCTTATCACATAATGATATCTATTTTAT
CTX1_Dopa 3123 bp        1732 TACAGCAAAAAGAGAATCAGTGTGGATGCTTTTAGTCGAAGAATGAAGAAATTCTTGTCCCTTAATTCAATTAACTTTAT

                         1921 ......1930......1940......1950......1960......1970......1980......1990......2000
CTX1_SEPES 3159 bp       1809 CTTTCAACAATTATCAAACGCAATTCTTGCAGAAGGATTTACTGATATCCAAGGTAAAGATTTCTTTAATACCTTTAAAA
CTX2_SEPES 3129 bp       1782 ATTTAAGCAACTGTCAAATGCGCTTTTGGCGAACGGCTTTGCCGATAAAAGTGGAAACGATTTCTTCAATATATTCAAAA
CTX1_DPEA 3156 bp        1806 TTTTGAACAATTGTCGAACGCACTCTTAGCAGAAGGATTTACTGATACCAAAGGCAGAGATTTCTTCAATACGTTTAAAA
CTX2_DPEA 3123 bp        1776 ATTTAAGCAATTGACGAATGCGCTTTTAGCGAATGGTTTTGCCGATAAAAGTGGAAGCGATTTCTTTAACACGTTTAAAA
CTX1_SBAN 2946bp         1806 CTTTCAACAATTGTCAAACGCGATTCTAGCAGAGGGATTTACTGATATAGAAGGTAAGGATTTCTTTAATACCTTTAAAA
CTX2_SBAN 2286 bp        1782 ATTCAAGCAATTGTCGAATGCACTTTTGGCGAACGGTTTTGCCGATAAAAGCGGGAACGATTTCTTCAACATATTCAAAA
CTX_EBER 3159 bp         1809 GTTTCAGCAGCTGTCAAACGCCATCCTGGCCGACGGTTTTACAGACATGAAGGGTAAGGATTTCTTTAACACTTTTAAAC
CTX2_SOFF 3129 bp        1782 ATTTAAGCAATTGTCGAATGCGTTTTTAGCGGACGGTTTTGCCGATAGAAGTGGAAACGATTTCTTCAACATATTCAAAA
CTX1_SOFF 3141bp         1791 ATTTCAACAATTGTCAAACGCAATTCTAGCAGAAGGATTTACAGATATCCGAGGTAAGGATTTCTTTAATACCTTTAAAA
CTX_ESCO 3159 bp         1809 GTTTCAGCAGCTGTCAAATGCCATCCTAGCCGACGGTTTTACAGACACAAAGGGTAAGGATTTCTTTAACACTTTTAAAC
CTX2_SLES 3132 bp        1785 ATTTAAGCAATTGACAAATGCGCTTTTAGCTAACGGCTTTGCCGATAAAAGTGGAAGCGATTTCTTCAACACGTTTAAAA
CTX1_SLES 3159 bp        1809 CTTTGAACAATTATCGAACGCAATCTTAGCAGAAGGATTTACTGATACCAAAGGCAGAGATTTCTTTAATACTTTTAAAA
CTX_TRHO 3150 bp         1803 ATTTAAGCAAATGGCGAATGCGGTTTTAGCCGAGGGTTTTGCCGATAAAAGTGGAAGCGATTTCTTCAACCTGTTTAAGA
CTX_SATL 3147 bp         1806 CTTTCAGCAATTGTCAAACGCCATCATGGCCGACGGGTTTACAGACACAAAGGGGAAAGATTTCTTTAACACTTTCAAAC
CTX2_SLYC 3129 bp        1782 ATTTAAGCAATTGTCAAATGCGCTTTTGGCGAACGGCTTTGCCGATAAAAGTGGAAACGATTTCTTCAATATATTCAAAA
CTX1_SLYC 3159 bp        1809 CTTTCAACAATTATCAAACGCAATTCTTGCAGAAGGATTTAGTGATATCCGAGGTAAGGATTTCTTTAATACCTTTAAAA
CTX_EPAR 3150 bp         1803 GTTTCAGCAGCTGTCAAACGCCATCATGGCCGACGGTTTTACAGACACGAAGGGCAAGGATTTCTTTAATACTTTTAAAC
CTX2_SPHAR 3129 bp       1782 ATTTAAGCAATTGTCAAATGCGCTTTTGGCGAACGGCTTTGCCGATAAAAGTGGAAACGATTTCTTCAATATATTCAAAA
CTX1_SPHAR 3159 bp       1809 CTTTCAACAATTATCAAACGCAATTCTTGCAGAAGGATTTACTGATATCCGAGGTAAGGATTTCTTTAATACCTTTAAAA
CTX_WSCI_PARTIAL 1653 bp 1008 CTTCCAACAATTGTCAAACGCGATCCTCGCCAACGGGTTCACTGATACGAGAGGGAATGATTTCTTCAACACGTTCAAAA
CTX_IHAL 3156 bp         1815 CTTTCAACAATTATCGAATGCAATTTTAGCTGAAGGATTCACCGACGCAAAAGGTCAAGATTTTTTCAATACATTCAAAA
CTX2_XNOT 3183 bp        1836 ATTTGATCAACTATCAAACGCCATTTTAGCGGAGGGATTTGCCGATAAAAGGGGACACGACTTTTTCAACACGTTTAAAA
CTX1_XNOT2 3150 bp       1809 CTTTCAACAATTATCAAATGCAATCTTAGCAGAAGGTTTCACCGACTCAAAAGGTAAAGATTTTTTCAATACATTCAAAA
CTX_OROB_PARTIAL 1431 bp 1338 CTTTCAACAATTGTCGAACGCGATCCTTGCGGACGGCTTTACAGACGCCAAGGGCAGTGATTTCTGCAACACTTTTAAGA
CTX_SAFF 3147 bp         1806 CTTTCAGCAATTGTCAAACGCCATCATGGCCGACGGGTTTACAGACACAAAGGGGAAAGATTTCTTTAACACTTTCAAAC
CTX_SOBS 3156bp          1806 CTTCCAGCAATTGTCGAACGCCATCATGGCCGACGGGTTCACGGACACAAAGGGCAAAGATTTCTTCAACACTTTCAAGC
CTX_Sarc 2907 bp         1273 CTTCCAACAATTGTCGAACGCGATCCTTGCGAATGGATTCACAGACACCAAAGGCAAAGATTTCTTCAACACGTTCAAAA
CTX2_Dopa 3123 bp        1776 ATTTAAGCAATTGACGAATGCGCTTTTAGCGAATGGTTTTGCCGATAAAAGTGGAAGCGATTTCTTTAACACTTTTAAAA
CTX1_Dopa 3123 bp        1812 TTTTGAACAATTATCGAACGCACTCTTAGCAGAAGGATTTAGTGATACCAAAGGCAGAGATTTCTTCAATACGTTTAAAA

                         2001 ......2010......2020......2030......2040......2050......2060......2070......2080
CTX1_SEPES 3159 bp       1889 GGATGATCGCTTCTAACAGAGATGCTTGTACCGCACCATATGGAAACGAGGCAACGATATTATTAGAGCGTCTCAGCAGA
CTX2_SEPES 3129 bp       1862 AGATGATCGCTTCCGACAGAGGCGCTTGTACTAAGAAATATGGAACCGAGGCGTCTATTCTGTTCGGTAGTCTCAGTCGA
CTX1_DPEA 3156 bp        1886 GGATGATTGCCTCAAACAATGAAGCTTGTAGCGAGAATTATGGAAACCAGGCAACAATTTTATTCGATCGTATCAGTCGA
CTX2_DPEA 3123 bp        1856 AGATAATCGCATTAGACAGAGGTGCTTGTACAAATCAATATGGAAAAGAGGCAACTGCACTACTTAAACGTCTTAGTCGA
CTX1_SBAN 2946bp         1886 GGATGATCGCTTCCAACAGAGGTTCTTGTACTGAAGAATATGTAAACGAGGCAAATATATTATTCGACCGTCTCAGCAGA
CTX2_SBAN 2286 bp        1862 AGATGATCGCTTCCGAGAAAGGCGCTTGCACTAAGAAATATGGAACCGAGGCGGCTGTTCTGCTTAACAGTCTCAGTCGA
CTX_EBER 3159 bp         1889 GGATGATTGCTACCGACCGAGATGCTTGCACAGAAGAATATGGACTCGAAGCGAGTAAGTTGTTCGAACGGCTGAGTCAG
CTX2_SOFF 3129 bp        1862 AGATGATCGCTTCCGACAGAGGCTCTTGTACTAAGAAATATGGAACCGATGCGGCTATTCTGCTCGACAGCCTCAGTCGA
CTX1_SOFF 3141bp         1871 GGATGATTGCTTCGAACAGGGGCGCTTGTACTGAACAATATGGACACGAAGCATCGATCCTATTTGATCGTCTCAGCCGA
CTX_ESCO 3159 bp         1889 GGATGATTGCTACTGACCGAGATGCTTGTACGGAAGAATATGGACTCAAAGCGAGTAAGTTGTTCGAACGACTGAGTCAG
CTX2_SLES 3132 bp        1865 AGATAATCGCCTCTGACAGTGGAGCTTGTACAGAGCAATATGGAAACGAGGCTACTATACTACTTAATCGTCTTAGTCGA
CTX1_SLES 3159 bp        1889 GGATGATTGCGTCAAACAGGGAAGCTTGTAGTAAAATATATGGAAACGAAGCAACAGTATTATTCAATCGTCTCAGCCGA
CTX_TRHO 3150 bp         1883 AGATGATCGCTTCTAACAGAGGTGCTTGTACCGAGCAATATGGAAACGACTCTATCGCACTTCTTAATCGTCTCAATCGA
CTX_SATL 3147 bp         1886 GGATGATTGCTACCGACCGAGATGCTTGTACGGAAGAATACGGAATCGAGGCAAGTAGGTTGTTCGACCGGCTGAGTCAG
CTX2_SLYC 3129 bp        1862 AGATGATCGCTTCCGACAGAGGCGCTTGTACTAAGAAATATGGAACTGAGGCGACTATTCTGCTTGAAAGTCTCAGTCGA
CTX1_SLYC 3159 bp        1889 GAATGATCGCTTCTAACAGGGATGCTTGTACTACACAATATGGAAGCGAGGCAACGATATTATTAGAGCGTCTCAGCCGA
CTX_EPAR 3150 bp         1883 GGATGATTGCCACCGACCGAGATGCTTGTACAGAAGAATATGGACTCGAAGCGAGTAAGTTGTTTGATCGGCTGAGTCAG
CTX2_SPHAR 3129 bp       1862 AGATGATCGCTTCCGACAGAGGCGCTTGTACTAAGAAATATGGAACTGAGGCGACTATTCTGCTCGACAGTCTCAGTCGA
CTX1_SPHAR 3159 bp       1889 GGATGATCGCTTCTAACAGAGATGCTTGTAGTACACAATATGGAAGCGAGGCAACGATATTGTTAGAACGTCTTAGCCGA
CTX_WSCI_PARTIAL 1653 bp 1088 GAATGATCGCATCTAATAGAGGTGCTTGTACTGAAGACTATGGAAAGGACGCATCGAATTTGATGGACCGTCTGAGTCGG
CTX_IHAL 3156 bp         1895 AGATGATTGCCTCAAACCGAGGAGCCTGTAGCGCTGAATATGGAAACGAGGCTCAAGTTTTGTTTGAACGTTTAAGCGAG
CTX2_XNOT 3183 bp        1916 TGATGATCGCTTCTAGCGTAGGTGCATGTAGTGTACAATACGGGATGGAAACGACTGGCGTATTCAACCATCTCAGTCTA
CTX1_XNOT2 3150 bp       1889 AGATGATTGCCTCAAACAGAGGCGCCTGTAGCGCGGAATATGGGAATCAGGCTCAACTTCTGTTTGAACGTTTAAGCGAG
CTX_OROB_PARTIAL 1431 bp 1418 GGATGATCGCCG--------------------------------------------------------------------
CTX_SAFF 3147 bp         1886 GGATGATTGCTACCGACCGAGATGCTTGTACGGAAGAATATGGAATCGAGGCGAGTAGGTTGTTCGACCGGCTGAGTCAG
CTX_SOBS 3156bp          1886 GGATGATCGCGACCGACCGAGACGCTTGTACGGAAGAATATGGAATCGAGGCGAGTAGGTTGTTCGAGAGACTGAGTCAG
CTX_Sarc 2907 bp         1353 GGATGGTCGCTTCCGACAGAGACGCGTGTACCCAACAATACGGAAACGATGCAGCGACATTAATGGATCGTCTCAGTCGA
CTX2_Dopa 3123 bp        1856 AGATAATCGCATTAGACAGAGGTGCTTGCACAAATCAATATGGAAAAGAGGCAACTGCACTACTTAAACGTCTTAGTCGA
CTX1_Dopa 3123 bp        1892 GGATGATTGCTTCAAACAATGAAGCTTGTAGCGAGAAATATGGAAACGAGGTAACAATTTTATTCGATCGTATCAGTCGA

                         2081 ......2090......2100......2110......2120......2130......2140......2150......2160
CTX1_SEPES 3159 bp       1969 TTAGACCTCACCGCTGCCGAATCTATCCTGGCCTATTACAGCTTCGAAAGCAATTATCTCAATCCGGCAAACATGAAACG
CTX2_SEPES 3129 bp       1942 ATGGACATCACCGCTGCCGAAGCTATACTCGCTCATTACTACTTCGAAAGCTTTTATCTGAACGCCGAGAACAAGGCGAT
CTX1_DPEA 3156 bp        1966 TTAGATATCACCGCTGCTCAAGCAATCCTTGCTTATTATAATTTCGAAAGCAATTATCTCAATAAAGGAAATATGAAAAA
CTX2_DPEA 3123 bp        1936 ATGGACATTGCCGCTGCTGAAGCTATTCTTGCTCATTACCATTTTCAAAGTTTTTATCTGAACGCCGAACACAAGAAAAA
CTX1_SBAN 2946bp         1966 TTAGACATCACCGCTGCCGAAACTATCCTTGCGTATTACAGCTTTGAAAGTAATTATCTCAATGCGGAGAATATGAAGCA
CTX2_SBAN 2286 bp        1942 ATGGACATCACCGCTGCCGAAGCGATACTTGCTCATTACTACTTCGAAAGCTTTTATCTGAACGCCGAGCACAGGACAAC
CTX_EBER 3159 bp         1969 TTGGACATCATGGCGGCTGAGGCGATCCTTGCCCATTATCATTTCCAAAGCAACTACCTCACGGAGGAACATATGAAAAC
CTX2_SOFF 3129 bp        1942 ATGGACATCACCGCTGCCGAAGCGATACTTGCCCATTACCACTTCGAAAGCTTTTATCTGAACGCCGAGCACAAGACGAC
CTX1_SOFF 3141bp         1951 TTAGACATCACCGCTGCTGAATCTATACTGGCCTATTACAGTTTCGAAAGCAATTATCTCAATCCGGAAAATATGAAACG
CTX_ESCO 3159 bp         1969 TTGGACATCATGGCGGCTGAGGCGATCCTTGCCCATTATCATTTCCAAAGCAACTACCTCACGGAGGAACATATGAAAAC
CTX2_SLES 3132 bp        1945 ATGGATATCACTGCTGCCGAAGCCATTCTTGCTCATTACCATTTCGAAAGTTTTTATCTGGACGCCGAACGCAAGGCAAA
CTX1_SLES 3159 bp        1969 ATTGACATCACCGCTGCTGAAGCCATCCTTGCTTATTATAATTTCGAAAGCAATTATCTCAATACAGAAAATATGAAAAA
CTX_TRHO 3150 bp         1963 TTGGATATCACTGCTGCCGAAGCCATTCTTGCTCATTACCATTTTGAAAGCTTTTATCTGAAACCCGAACAGAAGACAAA
CTX_SATL 3147 bp         1966 TTTGACATCATGGCGGCTGAGGCGATCCTTGCCCATTATCATTTCCAAAGTAATTACCTCACGCCGGAACATATGAAAAC
CTX2_SLYC 3129 bp        1942 ATGGACATCACCGCTGCCGAAGCGATACTCGCTCATTACTACTTCGAAAGCTTTTATCTGAACGTGGAGAACAAGGCGAC
CTX1_SLYC 3159 bp        1969 TTAGACATCACTGCTGCCGAATCTATCCTGGCCTATTACAGCTTCGAAAGCAATTATCTCAATCCGGAAAACATGAAACG
CTX_EPAR 3150 bp         1963 TTGGACATCATGGCGGCTGAGGCGATCCTTGCCCATTATCATTTCCAAAGCAACTACCTCACGGAGGAACATATGAAAAC
CTX2_SPHAR 3129 bp       1942 ATGGACATCACCGCTGCCGAAGCGATACTCGCTCATTACTACTTCGAAAGCTTTTATCTGAACGCCGAGAACAAGGCGAC
CTX1_SPHAR 3159 bp       1969 TTAGACATCACTGCTGCCGAATCTATCCTGGCCTATTACAGCTTCGAAAGCAATTATCTCAATCCGGAAAACATGAAACG
CTX_WSCI_PARTIAL 1653 bp 1168 TTGGATATAACGGCAGCAGAGACCATTCTTGCCTACTACAATTTCGAGAGTAATTATCTCAACAAGAAAAATATGAGAAA
CTX_IHAL 3156 bp         1975 CTTGACATTACGGCAGCAGAAGCGATTCTTGCTCATTATGAATTTGAAAGTAATTACCTTAACTCAGAAAATATGGACTC
CTX2_XNOT 3183 bp        1996 TTGGACATGACTGCAACGGAAACTATCCTTGCTCATTACCAGTTTGAAGACCTTTACTTGAAAGACGATCATAGATCAAA
CTX1_XNOT2 3150 bp       1969 CTTGACATCACTGCGGCAGAAGCTATTCTTGCTCATTATCAATTTGAAAGTAATTACCTTAACTCAGAACATATGGCCTC
CTX_OROB_PARTIAL 1431 bp 1430 --------------------------------------------------------------------------------
CTX_SAFF 3147 bp         1966 TTGGACATCATGGCGGCTGAGGCGATCCTTGCCCATTATCATTTCCAAAGTAATTACCTCACGCCGGAACATATGAAAAC
CTX_SOBS 3156bp          1966 TTTGACATCATGGCGGCCGAGGCAATCCTTGCCCATTATCATTTTCAAAGTAACTACCTCACGACGGAACATATGAAAAC
CTX_Sarc 2907 bp         1433 TTAGATATCACCGCTGCCGAAGCCATTCTTGCTTATTACAATTTCGAAAGTAATTATCTCAAGAAGGAGCATTTGAAAGA
CTX2_Dopa 3123 bp        1936 ATGGACATTGCCGCTGCTGAAGCTATTCTCGCTCATTACCATTTTCAAAGTTTTTATCTGAACGCCGAACACAAGAAAAA
CTX1_Dopa 3123 bp        1972 TTAGATATCACCGCTGCTCAAGCAATCCTTGCTTATTATAATTTCGAAAGCAATTATCTCAATAGAGGAAATATGAAAAA

                         2161 ......2170......2180......2190......2200......2210......2220......2230......2240
CTX1_SEPES 3159 bp       2049 AATGCTCGAAAATGCTAAGCAATTAGTACGAGATTCGAAAAGACGCATGCGTAGTTATGCTAG-----------------
CTX2_SEPES 3129 bp       2022 TATGCGTTCTCGTGTCGATCAGTTGGCACAAGATTCAAAGGAAAGAATGAAAAATTACGCTAG-----------------
CTX1_DPEA 3156 bp        2046 GATGTTTAATACAGCTAAACAATTGGTAAGAGATTCGAAACGCCGCATGCGAAAATATTCCAA-----------------
CTX2_DPEA 3123 bp        2016 TATGCTTTCAAGCGCAATGCAATTAGCACAAGGTTCAACGGAAAGAATGCAAAATTATGCTAG-----------------
CTX1_SBAN 2946bp         2046 GATGCATGTAAATGCTAAGCAACTGGTACAAGATTCGAAGCGACGCATGCGAAATTATGCTAG-----------------
CTX2_SBAN 2286 bp        2022 CATGCTTTCTAGTGTCGATCAGTTGGTAAAAGATTCAAAAGAAAGAATGAAAAATTACGCTAG-----------------
CTX_EBER 3159 bp         2049 GATGCAGGCGAGCGCGAAACAATTGGTACGTGATTCAAAACGACGTATGCGAAATTATGCCCG-----------------
CTX2_SOFF 3129 bp        2022 TATGTTATCTAGTGCCGTGCAGTTGGTACAAGATTCAAAGGAAAGAATGAAAAATTATGCTAG-----------------
CTX1_SOFF 3141bp         2031 GATGCTCGAAAATGCTAAGCAATTAGTACGAGATTCGAAACGACGCATGCGAAACTATGCTAG-----------------
CTX_ESCO 3159 bp         2049 GATGCAGGCGAGCGCGAAACAATTGGTACGTGATTCAAAACGACGTATGCGAAATTATGCCCG-----------------
CTX2_SLES 3132 bp        2025 TATGCTTTCAAACGCAATGCAATTAGTACAAGGCTCAACGGAAAGAATGAAAAATTATGCAAG-----------------
CTX1_SLES 3159 bp        2049 GATGACCATCGCTGCTAAGCAATTGGTGAGAGATTCGAAACGACGAATGCGAAATTATGCCAA-----------------
CTX_TRHO 3150 bp         2043 TATGCTTTTTAATGCAAAGCAATTGGCTCAAGACTCGAAGGAAAGACTGAAAATGTATGCCAA-----------------
CTX_SATL 3147 bp         2046 GATGCAAGCGAGCGCGAAGCAGTTGGTACGCGATTCAAAGCGCCGTATGCGAAATTATGCCCG-----------------
CTX2_SLYC 3129 bp        2022 TATGCTTTCTAGTGTCGATCAGTTGGTACAAGATTCCAAGGAAAGAATGAAAAATTACGCCAG-----------------
CTX1_SLYC 3159 bp        2049 GATGCTCGAAAATGCTAAGCAATTAGTACGAGATTCGAAAAGACGCATGCGAAGTTATGCTAG-----------------
CTX_EPAR 3150 bp         2043 GATGCAGGCGAGCGCGAAACAATTGGTACGTGATTCAAAGCGGCGTATGCGAAATTATGCCCG-----------------
CTX2_SPHAR 3129 bp       2022 TATGCTTTCTAGTGTCGATCAGTTGGTACAAGATTCCAAGGAAAGAATGAAAAATTACGCCAG-----------------
CTX1_SPHAR 3159 bp       2049 GACGCTCGAAAATGCTAAACAATTAGTACGAGATTCGAAAAGACGCATGCGTAGTTATGCTAG-----------------
CTX_WSCI_PARTIAL 1653 bp 1248 GATGATAGCCGACGCAAAGCAATTGGTGAGAGATTCGAAACGTCGTATGCAGAGTTACGCCAA-----------------
CTX_IHAL 3156 bp         2055 TATGATGGAGAGCGCTAAACAATTGGTTATAGACTCAAAAAGACGCATGCGTAGTTATGCAAG-----------------
CTX2_XNOT 3183 bp        2076 CATGATTAAAGAGGCTAAACACTTCACCCAAGGTTCCAAAGAAAGAATGAAGAATTATGTAAA-----------------
CTX1_XNOT2 3150 bp       2049 TATGAAGGAGAGCGCTAAACAATTGGTCAGGGATTCAAAAAGGCGAATGCGCAGTTATGCAAA-----------------
CTX_OROB_PARTIAL 1431 bp 1430 --------------------------------------------------------------------------------
CTX_SAFF 3147 bp         2046 GATGCAAGCGAGCGCGAAGCAGTTGGTACGCGATTCAAAGCGCCGTATGCGAAATTATGCCCG-----------------
CTX_SOBS 3156bp          2046 GATGCAAACGAGCGCAAAACAGTTGGTACGCGATTCAAAGCGCCGTATGCGAAATTATGCCCG-----------------
CTX_Sarc 2907 bp         1513 GATGCACGCTAGTGCAAAGCAATTGGTACAAGATTCAAAACGACGCATGCGAAATTATGCCAAATACTGTTTGTTTGATT
CTX2_Dopa 3123 bp        2016 TATGCTTTCAAGCGCAATGCAATTAGCACAAGGTTCAACGGAAAGAATGAGAAATTATGCTAG-----------------
CTX1_Dopa 3123 bp        2052 GATGTTTAATTCTGCTAAGCAATTGGTAAGAGATTCGAAACGCCGCATGCGAAAGTATGCCAA-----------------

                         2241 ......2250......2260......2270......2280......2290......2300......2310......2320
CTX1_SEPES 3159 bp       2112 --------------------------------------------------------------------------------
CTX2_SEPES 3129 bp       2085 --------------------------------------------------------------------------------
CTX1_DPEA 3156 bp        2109 --------------------------------------------------------------------------------
CTX2_DPEA 3123 bp        2079 --------------------------------------------------------------------------------
CTX1_SBAN 2946bp         2109 --------------------------------------------------------------------------------
CTX2_SBAN 2286 bp        2085 --------------------------------------------------------------------------------
CTX_EBER 3159 bp         2112 --------------------------------------------------------------------------------
CTX2_SOFF 3129 bp        2085 --------------------------------------------------------------------------------
CTX1_SOFF 3141bp         2094 --------------------------------------------------------------------------------
CTX_ESCO 3159 bp         2112 --------------------------------------------------------------------------------
CTX2_SLES 3132 bp        2088 --------------------------------------------------------------------------------
CTX1_SLES 3159 bp        2112 --------------------------------------------------------------------------------
CTX_TRHO 3150 bp         2106 --------------------------------------------------------------------------------
CTX_SATL 3147 bp         2109 --------------------------------------------------------------------------------
CTX2_SLYC 3129 bp        2085 --------------------------------------------------------------------------------
CTX1_SLYC 3159 bp        2112 --------------------------------------------------------------------------------
CTX_EPAR 3150 bp         2106 --------------------------------------------------------------------------------
CTX2_SPHAR 3129 bp       2085 --------------------------------------------------------------------------------
CTX1_SPHAR 3159 bp       2112 --------------------------------------------------------------------------------
CTX_WSCI_PARTIAL 1653 bp 1311 --------------------------------------------------------------------------------
CTX_IHAL 3156 bp         2118 --------------------------------------------------------------------------------
CTX2_XNOT 3183 bp        2139 --------------------------------------------------------------------------------
CTX1_XNOT2 3150 bp       2112 --------------------------------------------------------------------------------
CTX_OROB_PARTIAL 1431 bp 1430 --------------------------------------------------------------------------------
CTX_SAFF 3147 bp         2109 --------------------------------------------------------------------------------
CTX_SOBS 3156bp          2109 --------------------------------------------------------------------------------
CTX_Sarc 2907 bp         1593 GATTTTACAGCATACCAACCCAATGGGTCATTTAATGCTGAAACATTTTTAGAGTCTTGCGACTTTGGTAAAAAAGGTGT
CTX2_Dopa 3123 bp        2079 --------------------------------------------------------------------------------
CTX1_Dopa 3123 bp        2115 --------------------------------------------------------------------------------

                         2321 ......2330......2340......2350......2360......2370......2380......2390......2400
CTX1_SEPES 3159 bp       2112 --------------------------------------------------------------------------------
CTX2_SEPES 3129 bp       2085 --------------------------------------------------------------------------------
CTX1_DPEA 3156 bp        2109 --------------------------------------------------------------------------------
CTX2_DPEA 3123 bp        2079 --------------------------------------------------------------------------------
CTX1_SBAN 2946bp         2109 --------------------------------------------------------------------------------
CTX2_SBAN 2286 bp        2085 --------------------------------------------------------------------------------
CTX_EBER 3159 bp         2112 --------------------------------------------------------------------------------
CTX2_SOFF 3129 bp        2085 --------------------------------------------------------------------------------
CTX1_SOFF 3141bp         2094 --------------------------------------------------------------------------------
CTX_ESCO 3159 bp         2112 --------------------------------------------------------------------------------
CTX2_SLES 3132 bp        2088 --------------------------------------------------------------------------------
CTX1_SLES 3159 bp        2112 --------------------------------------------------------------------------------
CTX_TRHO 3150 bp         2106 --------------------------------------------------------------------------------
CTX_SATL 3147 bp         2109 --------------------------------------------------------------------------------
CTX2_SLYC 3129 bp        2085 --------------------------------------------------------------------------------
CTX1_SLYC 3159 bp        2112 --------------------------------------------------------------------------------
CTX_EPAR 3150 bp         2106 --------------------------------------------------------------------------------
CTX2_SPHAR 3129 bp       2085 --------------------------------------------------------------------------------
CTX1_SPHAR 3159 bp       2112 --------------------------------------------------------------------------------
CTX_WSCI_PARTIAL 1653 bp 1311 --------------------------------------------------------------------------------
CTX_IHAL 3156 bp         2118 --------------------------------------------------------------------------------
CTX2_XNOT 3183 bp        2139 --------------------------------------------------------------------------------
CTX1_XNOT2 3150 bp       2112 --------------------------------------------------------------------------------
CTX_OROB_PARTIAL 1431 bp 1430 --------------------------------------------------------------------------------
CTX_SAFF 3147 bp         2109 --------------------------------------------------------------------------------
CTX_SOBS 3156bp          2109 --------------------------------------------------------------------------------
CTX_Sarc 2907 bp         1673 GTTTTGTTTCAAATAGGTGTCTTGCGACTTTGGTAAACAAATCAGGGGGTGGGGATGAATAAGGTTTGGTCTAGATCCTT
CTX2_Dopa 3123 bp        2079 --------------------------------------------------------------------------------
CTX1_Dopa 3123 bp        2115 --------------------------------------------------------------------------------

                         2401 ......2410......2420......2430......2440......2450......2460......2470......2480
CTX1_SEPES 3159 bp       2112 --------------------------------------------------------------------------------
CTX2_SEPES 3129 bp       2085 --------------------------------------------------------------------------------
CTX1_DPEA 3156 bp        2109 --------------------------------------------------------------------------------
CTX2_DPEA 3123 bp        2079 --------------------------------------------------------------------------------
CTX1_SBAN 2946bp         2109 --------------------------------------------------------------------------------
CTX2_SBAN 2286 bp        2085 --------------------------------------------------------------------------------
CTX_EBER 3159 bp         2112 --------------------------------------------------------------------------------
CTX2_SOFF 3129 bp        2085 --------------------------------------------------------------------------------
CTX1_SOFF 3141bp         2094 --------------------------------------------------------------------------------
CTX_ESCO 3159 bp         2112 --------------------------------------------------------------------------------
CTX2_SLES 3132 bp        2088 --------------------------------------------------------------------------------
CTX1_SLES 3159 bp        2112 --------------------------------------------------------------------------------
CTX_TRHO 3150 bp         2106 --------------------------------------------------------------------------------
CTX_SATL 3147 bp         2109 --------------------------------------------------------------------------------
CTX2_SLYC 3129 bp        2085 --------------------------------------------------------------------------------
CTX1_SLYC 3159 bp        2112 --------------------------------------------------------------------------------
CTX_EPAR 3150 bp         2106 --------------------------------------------------------------------------------
CTX2_SPHAR 3129 bp       2085 --------------------------------------------------------------------------------
CTX1_SPHAR 3159 bp       2112 --------------------------------------------------------------------------------
CTX_WSCI_PARTIAL 1653 bp 1311 --------------------------------------------------------------------------------
CTX_IHAL 3156 bp         2118 --------------------------------------------------------------------------------
CTX2_XNOT 3183 bp        2139 --------------------------------------------------------------------------------
CTX1_XNOT2 3150 bp       2112 --------------------------------------------------------------------------------
CTX_OROB_PARTIAL 1431 bp 1430 --------------------------------------------------------------------------------
CTX_SAFF 3147 bp         2109 --------------------------------------------------------------------------------
CTX_SOBS 3156bp          2109 --------------------------------------------------------------------------------
CTX_Sarc 2907 bp         1753 TCCTTTGACCTGTCTGGCTTGGTTGGACCTACCAGGAATTTAATTCCGGCCAGCGTAGCTCTCCGGGTCATTGGAACATT
CTX2_Dopa 3123 bp        2079 --------------------------------------------------------------------------------
CTX1_Dopa 3123 bp        2115 --------------------------------------------------------------------------------

                         2481 ......2490......2500......2510......2520......2530......2540......2550......2560
CTX1_SEPES 3159 bp       2112 ---------------------------------------------ATACTGGGAACGCACGAGTTGCCCGCCCCTGAATG
CTX2_SEPES 3129 bp       2085 ---------------------------------------------CTACTGGGAACGCACGAGTTGTCCGCCACTTAATA
CTX1_DPEA 3156 bp        2109 ---------------------------------------------TTACTGGGAAAACACAAGTTGTCCACCGTTGAATG
CTX2_DPEA 3123 bp        2079 ---------------------------------------------GTACTGGGAAAGAACGAGTTGTCCCCCCTTAAATG
CTX1_SBAN 2946bp         2109 ---------------------------------------------ATACTGGGAACGCACGAGTTGCCCGCCCTTGAATG
CTX2_SBAN 2286 bp        2085 ---------------------------------------------ATACTGGGAACGCACGAGTTGTCCGCCACTTAATG
CTX_EBER 3159 bp         2112 ---------------------------------------------ATACTGGGAACGCACAAGTTGCCCGCCGTTAAACG
CTX2_SOFF 3129 bp        2085 ---------------------------------------------ATACTGGGAACGCACGAGTTGTCCGCCCTTGAATA
CTX1_SOFF 3141bp         2094 ---------------------------------------------ATACTGGGAACGCACGAGTTGTCCGCCCTTGAATA
CTX_ESCO 3159 bp         2112 ---------------------------------------------ATACTGGGAACGCACAAGTTGCCCGCCATTAAACG
CTX2_SLES 3132 bp        2088 ---------------------------------------------GTACTGGGAACGCACGAGTTGTCCGCCCTTAAATG
CTX1_SLES 3159 bp        2112 ---------------------------------------------ATACTGGGAAGGCACAAGTTGTCCGCCTCTGAACG
CTX_TRHO 3150 bp         2106 ---------------------------------------------CTACTGGGAACGCACGAGTTGCCCACCCCTGAATG
CTX_SATL 3147 bp         2109 ---------------------------------------------CTATTGGGAACGCACAAGTTGCCCGGCGTTAAATG
CTX2_SLYC 3129 bp        2085 ---------------------------------------------ATACTGGGAACGCACGAGTTGTCCGCCTCTTAATA
CTX1_SLYC 3159 bp        2112 ---------------------------------------------ATACTGGGAACACACGAGTTGCCCGCCCCTGAATG
CTX_EPAR 3150 bp         2106 ---------------------------------------------ATACTGGGAACGCACAAGTTGCCCGGCGTTAAACG
CTX2_SPHAR 3129 bp       2085 ---------------------------------------------CTACTGGGAACGCACGAGTTGTCCGCCTCTTAATA
CTX1_SPHAR 3159 bp       2112 ---------------------------------------------ATACTGGGAACACACAAGTTGCCCGCCCCTGAATG
CTX_WSCI_PARTIAL 1653 bp 1311 ---------------------------------------------ATACTGGGAACGTAATAGTTGTCCCCCTCTGGATG
CTX_IHAL 3156 bp         2118 ---------------------------------------------ATATTGGGAACGTACCAGTTGTCCGCCCCTGAATG
CTX2_XNOT 3183 bp        2139 ---------------------------------------------ATATTGGCACAAGACGAGTTGTCCGCCCTTGAATG
CTX1_XNOT2 3150 bp       2112 ---------------------------------------------ATATTGGGAACGTACCAGTTGCCCGCCACTGAATG
CTX_OROB_PARTIAL 1431 bp 1430 --------------------------------------------------------------------------------
CTX_SAFF 3147 bp         2109 ---------------------------------------------CTATTGGGAACGCACAAGTTGCCCGGCGTTAAATG
CTX_SOBS 3156bp          2109 ---------------------------------------------ATATTGGGAACGCACAAGTTGCCCGCCGTTAAATG
CTX_Sarc 2907 bp         1833 CAAGCCACTCCACCACGCCAAGGTACTTATTCGTGGAGTGGCCAAATACTGGGAACGCACGAGTTGCCCGACCTTGAATG
CTX2_Dopa 3123 bp        2079 ---------------------------------------------GTACTGGGAAAAAACGAGTTGTCCCCCCTTAAATG
CTX1_Dopa 3123 bp        2115 ---------------------------------------------TTACTGGGAAAACACAAGTTGTCCACCCTTGAATG

                         2561 ......2570......2580......2590......2600......2610......2620......2630......2640
CTX1_SEPES 3159 bp       2147 TCACTCACCTTACCCAAACCGGATGTGGGGCCTTATTGAGTTTCGAAGGCATGAAAGTGAAATTGTCTTGCGATGGCGGT
CTX2_SEPES 3129 bp       2120 TTCCTTACCTCAACCAGACCGGATGTGGCGAATTGTTGAGTTTTGAAGGCATGAAAGTAAAGCTGTCTTGTGATGGAGGC
CTX1_DPEA 3156 bp        2144 TCACTTACGTTAGGCAAAATGGATGCGGTGCCATGTTGAGCTTCGAGGGTATGGAAGTAAGTTTGTCTTGCGACGGAAAC
CTX2_DPEA 3123 bp        2114 TCACTCACCTCACTCAGACTGGATGTGGTACCATGTTGAGTTTCGAAGGTATGAAAGTGAAATTGTTTTGTGATGGAGGC
CTX1_SBAN 2946bp         2144 TCACTCATCTTACGCAAGCCGGATGTGGGGCCTTATTGAGCTTCGAAGGCATGAAAGTAAAATTGTCCTGCGATGGTGGA
CTX2_SBAN 2286 bp        2120 TCACTTACCTCAACGAGACCGGATGTGGCGAACTGTTGAGTTTCGAAGGCATGAAAGTGAAGCTGTCTTGTGTTGAAGGC
CTX_EBER 3159 bp         2147 TCACTCACCTCAGGCAAAACGGATGTAGCAACATGTTAAGCTTTGAAGGTATGACAGTGAAATTGTCCTGCGCTGAGGGA
CTX2_SOFF 3129 bp        2120 TCACTTACCTCAATCAGACTGGGTGTGGCGAATTACTGAGCTTCGAAGGCATGAAAGTGAAGCTGTCTTGTGATGGCGGC
CTX1_SOFF 3141bp         2129 TCACTCAACTTACCCAAACCGGATGTGGGGACCTATTAAGTTTCGAAGGCATGAAAGTGAAACTGTCTTGCGATGGCGGC
CTX_ESCO 3159 bp         2147 TCACTCACCTCAGGCAAAACGGATGTGGCAACATGTTAAGCTTTGAGGGTATGAGAGTGAAATTGTCCTGCGTTGGGGGA
CTX2_SLES 3132 bp        2123 TCACTTACCTCACTCAGATAGGATGTGGCGCTATGTTGAGTTTCCAGGGCATGAAAGTGAAATTGTTTTGTGATGGCGGC
CTX1_SLES 3159 bp        2147 TCACTTACCTTACCCAAAGTGGATGCGGTGCCATGTTGAGCTTCGAAGGCATGAACGTAAGTTTGTCTTGCGATGGAGAT
CTX_TRHO 3150 bp         2141 TCACCTACCTTACCCAGACCGGATGTGGGGCCATGTTGAGTTACGAAGGCATGAAAGTGAAATTGAATTGTGATGGTGGC
CTX_SATL 3147 bp         2144 TCACTCACCTCAGGCAAAACGGATGTGGCGACATGCTAAGCTTCGAGGGGATGAAGGTGAAATTGTCCTGCGTTGACGGT
CTX2_SLYC 3129 bp        2120 TCCCTTACCTCAACCAGACGGGATGTGGCGAACTGTTGAGTTTCGAAGGCATGAAAGTAAAGCTATCTTGTAATGGAGGC
CTX1_SLYC 3159 bp        2147 TCACTAATCTTGCCCAAACCGGGTGTGGGGCTTTTCTGAGTTTCGAAGGCATGAAAGTGAAATTATCTTGCGATGGCGGC
CTX_EPAR 3150 bp         2141 TCACTCACCTCCGACAAAACGGATGTGGCAACATGTTAAGCTTTGAGGGTATGAAAGTGAAATTGTCCTGCGCTGGGGGA
CTX2_SPHAR 3129 bp       2120 TCCCTTACCTCAACCAGGCCGGATGTGGCGAACTGTTGAGTTTCGAAGGCATGAAAGTAAAGCTGTCTTGTGATGGAGGC
CTX1_SPHAR 3159 bp       2147 TCACTAATCTTTCCCAAACCGGATGTGGGGCTTTTCTGAGTTTCGAAGGCATGAAAGTGAAACTGTCTTGCGATGGTGGC
CTX_WSCI_PARTIAL 1653 bp 1346 TCCCTGGCCTCGAGCAGACTGGATGTGGATCCATGTTGAGTTTCGAAGGCATGAAACTGAAAGTATCTTGCGATGGAGGT
CTX_IHAL 3156 bp         2153 TTAGTTACCTTGTTCAAAATGGCTGCGGCTATTTGTTGAGCTTTGAAGGCATGAAAGTTGATTTGGCTTGCGAGGACGGA
CTX2_XNOT 3183 bp        2174 CTACTCACCTAATACAAGTCGGATGTTCAGCTATGTTGAGTTACGAAGGCATGAAAGTGGAATTATCTTGTGAGGGAGGC
CTX1_XNOT2 3150 bp       2147 CTAGTTTTCTTGTTCAAAATGGTTGCGGCGATTTGTTGAGTTTTGAAGGCATGAAAGTGGATTTAGCTTGCGAGGGCGGA
CTX_OROB_PARTIAL 1431 bp 1430 --------------------------------------------------------------------------------
CTX_SAFF 3147 bp         2144 TCACTCACCTCAGGCAAAACGGATGTGGCGACATGCTAAGCTTCGAGGGGATGAAGGTGAAATTGTCCTGCGTTGACGGC
CTX_SOBS 3156bp          2144 TCACTTACCTCAGACAAAACGGATGCGGCAGCATGCTAAGCTTCGAGGGGATGAAAGTGAAATTGTCCTGCGTTGACGGT
CTX_Sarc 2907 bp         1913 TTACTGGCCTCACACAGACTGGGTGCGGCGCCATGTTGAGCTTCGAAGGCATGAAAGCGACATTGTCTTGCAATGGAGAC
CTX2_Dopa 3123 bp        2114 TCACTCACCTGACTCAGACTGGATGTGGTACCATGTTAAGTTTCGAAGGCATGAAAGTGAAATTGTTTTGTGATGGAGGC
CTX1_Dopa 3123 bp        2150 TCACGTACCTTAACCAAAGTGGATGCGGTGCCAAATTGAGTTTCAAGGGCATGGAAGTAAGTTTGTCTTGCGATGGAGAC

                         2641 ......2650......2660......2670......2680......2690......2700......2710......2720
CTX1_SEPES 3159 bp       2227 AG----GGCAGCAGTGCCACAGAACATTGAATGTGTCAATGTGAATGGTAACTTGCAATGGAGCGCCACACCCAAGTGTG
CTX2_SEPES 3129 bp       2200 AG----GGAAGCCGTCCATACAGCGATCGAGTGCAGCATATCGGAAGGCAAACTCAAATGGAACGCTACTGCAAAGTGTG
CTX1_DPEA 3156 bp        2224 AG----GGCAGTAGAACCACAAAATGTTAAATGTGTCAAGTTGAATGGTAAACTGCAATGGAGTGCTACTCCCAGGTGTC
CTX2_DPEA 3123 bp        2194 AG----AGAAGCCGTTCCCCCAGTAATTGAATGCAATCTATCAAAAGGCAAACTCAAATGGAATGCTACTGCCAAGTGCG
CTX1_SBAN 2946bp         2224 AG----AGCAGCAGTGCCGCGGACAATTGAATGCGTCAGTGTGAACGGCAAGTTGCAATGGAGCGCCACGCCTAAGTGTG
CTX2_SBAN 2286 bp        2200 AT----GGAAGTCATTCCTAAAGAGATCGAGTGCAGGTTATCGGAAGGCAAACTCGAATGGAACGCTACTGTAAAGTGTG
CTX_EBER 3159 bp         2227 CG----AGTGGCCGAACCGCGAAATATCGAATGCGTCAATTCGCGTGGTAAACTGCAATGGAGTGCCACACCAAGATGCC
CTX2_SOFF 3129 bp        2200 AG----GGAAGCAGTTCCGACAGCGATTGAGTGCAGGTTATCGGGAGGGAAACTTGAATGGAATGCTACTGCAAAGTGTG
CTX1_SOFF 3141bp         2209 AG----AGCGGCAGTGCCGAAGACCATTGAATGTGTTAATGTGAATGGTAACTTGCAATGGAGCGCTACACCCAAATGTG
CTX_ESCO 3159 bp         2227 CG----AGTAGCCGAACCGCAGAATATCGAATGTGTCAATTCGCGTGGTAAGCTGCAATGGAGTGCCACACCAAGATGCC
CTX2_SLES 3132 bp        2203 AG----AGAAGCAGTTCCTCCTGTGATTGAGTGCAATCTATCGGAAGACAAACTCGAATGGAACGCTACTGCCAAGTGCG
CTX1_SLES 3159 bp        2227 AG----AGCAGTAGAACCGCAAAATATTGAATGTATCAAGTTGAATAATAAGCTGCAGTGGAGCGCCACACCAAGGTGTG
CTX_TRHO 3150 bp         2221 AG----AAAGGCTATTCCTCCAGTAATTGAGTGCAGTTTAACGGAAGGCAAACTCGAATGGAATGCAACTGCCAAGTGCG
CTX_SATL 3147 bp         2224 CG----AGTAGCCGAGCCGCGAAATATTGAATGCGTCAACTCGCATGGTAAATTGCAATGGAGTGCCACACCAAGATGTC
CTX2_SLYC 3129 bp        2200 AG----GCAAGCCGTCCCTACAGCGATTGAGTGCAGCTTATCGGAAGGCAAACTCAAATGGAACGCTACTGCAAAGTGCG
CTX1_SLYC 3159 bp        2227 AG----GGCAGCAGTGCCGCAGAACATTGAATGTGTCAATGTGAATGGTAACTTGCAATGGAGCGCCACACCCAAGTGTG
CTX_EPAR 3150 bp         2221 CG----AGTAGCCGAGCCGCGAAATATCGAATGCGTCAATTCGCACGGTAAACTGCAATGGAGTGCCACACCAAGATGCC
CTX2_SPHAR 3129 bp       2200 AG----GGAAGTCGTCCCTACAGCGATCGAGTGCATCTTATCGGAAGGCAAACTCAAATGGAACGCTACTGCAAAGTGCG
CTX1_SPHAR 3159 bp       2227 AG----GGCAGCAGTGCCGAAGAGCATTGAATGCGTCAATGTGAATGGTAACTTGCAATGGAGCGCCACACCCAAGTGTG
CTX_WSCI_PARTIAL 1653 bp 1426 AA----AGTAGCGAAACAGCGAAATATCGAATGCGTCAAGAGGAAAGCTGGGTTACAATGGAGYGCCACGCCCAGTTGTT
CTX_IHAL 3156 bp         2233 AG----ACGGGCAGCACCGCAAAGGATAGAGTGCGTGAAAGTAGGCGGGAAATTACAATGGAGTGCAGAAGCCGAATGTA
CTX2_XNOT 3183 bp        2254 AG----ACAGGCCGTTCCACCCGTGATTGAGTGTCGGTCATTTGGAAATAAACTTAATTGGAACTCGTCCATTCACTGTG
CTX1_XNOT2 3150 bp       2227 CG----TCGAGCAGCACCACAAACGATTGAGTGCGTTAAAGTGAATAGAAAATTACAATGGAGTGCAATGCCCGAATGTA
CTX_OROB_PARTIAL 1431 bp 1430 --------------------------------------------------------------------------------
CTX_SAFF 3147 bp         2224 CG----AGTAGCCGAGCCGCGAAATATTGAATGCGTCAACTCGCATGGTAAATTGCAATGGAGTGCCACACCAAGATGTC
CTX_SOBS 3156bp          2224 CG----AGTAGCAGAGCCGCAAAATATTGAATGCGTCAGTTCGCGTGGTAAATTGCAATGGAGTGCCACACCAAAATGTC
CTX_Sarc 2907 bp         1993 AGAGAGAGCAGCAGAGCCGCAAACTATTGAATGTGTTAAGATGAATGGTAAGTTGCAATGGACTGTCACACCCACGTGTG
CTX2_Dopa 3123 bp        2194 AG----AGAAGCCGTTCCTCCAGTGATTGAGTGCAATCTATCAAAAGGCAAACTCGAATGGAATGCTACTGCCAAGTGCG
CTX1_Dopa 3123 bp        2230 AG----AGCAGTAGAACCACAAAATGTTAAATGTGTCAAGTTGAATGGTAAACTGCAATGGAGTGCTACTCCCAGGTGTC

                         2721 ......2730......2740......2750......2760......2770......2780......2790......2800
CTX1_SEPES 3159 bp       2303 AATCGAGTTGGTCCAGGTGGTCTAAGTGGAGTGCTTGCGCTTCCACTTGTGGCAATGCTACCCAGTCTCGAAGGCGGAGA
CTX2_SEPES 3129 bp       2276 TACCGGCCTGGTCGGCATGGGGAGAATGGAGTTCTTGCAGTTCAACTTGCGGGTCAGGTACCCAGACACGAACACGAGTA
CTX1_DPEA 3156 bp        2300 TAGCAAGTTGGTCTCAGTGGTCTCAATGGACTCCTTGTACTTCCACGTGTGGAAATGGCACTCAGTCCCGACAGCGAATA
CTX2_DPEA 3123 bp        2270 TACCGGGTTGGTCAACATGGGGAGAATGGAGTTCTTGTAGCTCAACTTGCGGCCAAGGTGTGCAGACTCGAAGCAGAGAA
CTX1_SBAN 2946bp         2300 CAGCGGGCTGGTCCAAGTGGTCTAAATGGACTGCTTGCGCTTCCACTTGTGGTAATGCTACCCAGTCTCGAAGGAGAAGA
CTX2_SBAN 2286 bp        2276 TACCGGGT------------------------------------------------------------------------
CTX_EBER 3159 bp         2303 TAGCGAAGTGGTCTCAGTGGACTCAATGGGAACCGTGTTCTGCCACTTGTGGCGATGGTATTCAATTACGAAGACGAACC
CTX2_SOFF 3129 bp        2276 TACCCGGTTGGTCGGCATGGGGTGAATGGGGTTCTTGTAACTCAACTTGCGGGTTAGGTATTCAAACACGAAAACGAGAA
CTX1_SOFF 3141bp         2285 CGGCGAGTTGGTCCAGGTGGTCTCAGTGGACTGCTTGTGCTTCCACATGTGGTAACGCTACCAAGTCTCGAAGGCGGAGA
CTX_ESCO 3159 bp         2303 TAGCGAAGTGGTCTCCGTGGTCTCAATGGGAACCGTGTTCTGCCACTTGTGGCGATGGTATTCAATTACGAAGACGAACC
CTX2_SLES 3132 bp        2279 TACCGGGTTGGTCGACATGGAGAGAATGGAGTTCTTGCAGCTCAACTTGCGGACCAGGTGTGCAAACTCGAAGCAGAGAA
CTX1_SLES 3159 bp        2303 TAGCAGGTTGGTCTAATTGGTCTCAATGGACTCCTTGTACTTCAACATGTGGAAATGGCACACAGTCTCGAAGGCGAATA
CTX_TRHO 3150 bp         2297 TACCGGGTTGGTCCGCGTGGGGGGAATGGGGTTCTTGTAGCTCTACTTGCGGGTCAGGAATTCAGATTCGAAACAGAGAA
CTX_SATL 3147 bp         2300 TGGCGAAGTGGTCTGAGTGGTCTCAGTGGGGCTCCTGTTCTGCCACTTGTGGCGGTGGCGTTCAATCGCGAAGACGGTTC
CTX2_SLYC 3129 bp        2276 TACCCGGCTGGTCGGCATGGGGAGAATGGAGTTCTTGCAGCTCAACTTGCGGGCCAGGTACCCAGACACGAACACGAGAA
CTX1_SLYC 3159 bp        2303 AAGCGAGTTGGTCCAGGTGGTCTAAGTGGAGTGCTTGCGCTTCCACTTGTGGTAATGCTACCCAGACCCGAAGGCGGAGA
CTX_EPAR 3150 bp         2297 TAGCGAAGTGGTCTCAATGGTCTCAATGGGAACCGTGTTCTGCCACTTGCGGTGATGGTGTTCAATCGCGAAGACGAACC
CTX2_SPHAR 3129 bp       2276 TACCGGGCTGGTCGGCATGGGGAGAATGGAGTTCTTGCAGCTCAACTTGCGGGCCAGGTACCCAGACACGAACACGAGAA
CTX1_SPHAR 3159 bp       2303 AAGCGAGTTGGTCCAGGTGGTCTAAGTGGAGTGCTTGCGCTTCCACTTGTGGTAATGCTACCCAGTCTCGAAGACGGAGA
CTX_WSCI_PARTIAL 1653 bp 1502 CAGCGAGTTGGTCTAAATGGTCGTCTTGGAGTCAATGCTCTTCAAGCTGTGGAAATGCAACTCAGACTCGTCGTCGGGTT
CTX_IHAL 3156 bp         2309 AATCAAGTTGGTCTGAATGGTCGGAGTGGTCGAGTTGCCCCGAAATGTGTGACTCTTCAACCACCACTCGAAGTCGGGTG
CTX2_XNOT 3183 bp        2330 CACCCAACTGGTCAATATGGGGTGAATGGAGTTCATGCTCTTCTACTTGTGGCCGAGGTAGCCAAATACGAAGCCGAAAA
CTX1_XNOT2 3150 bp       2303 AATCAAGTTGGTCTGAATGGTCGGAGTGGTCTAGTTGCCCTGAAACCTGCGACACTTCGATCAGCACTCGAAGTCGTGTA
CTX_OROB_PARTIAL 1431 bp 1430 --------------------------------------------------------------------------------
CTX_SAFF 3147 bp         2300 TGGCGAAGTGGTCCGAGTGGTCTCAGTGGGGCCCCTGTTCTGTCACTTGTGGCGGTGGCGTTCAATCGCGAAGACGGTTC
CTX_SOBS 3156bp          2300 TGTCGACTTGGTCTCGATGGTCCAAGTGGACGCCCTGTTCTGCCACTTGTGGCGATGGCGTTCAATCGCGAAGACGGACC
CTX_Sarc 2907 bp         2073 TCGCGAGTTGGTCCGCGTGGTCTGCGTGGACTCCTTGTGCTTCCACTTGTGGAAATGCTACCCAGTCTCGTAGTCGGGTA
CTX2_Dopa 3123 bp        2270 TACCGGGTTGGTCAACATGGGGAGAATGGAGTTCTTGTAGCTCAACTTGCGGCCAAGGTGTGCAGACTCGAAGCAGAAAA
CTX1_Dopa 3123 bp        2306 TAGCAAGTTGGTCTCAGTGGTCTCAATGGACTCCTTGCACTTCCACGTGTGGAAATGGCACTCAGTCCCGAAAGAGAATA

                         2801 ......2810......2820......2830......2840......2850......2860......2870......2880
CTX1_SEPES 3159 bp       2383 TGTCTGGGACAATCTGAGAGTGAAAAATGCATAGGGCCGTCAAAACAGGTACGAAAATGTTTCGTCGAAGATTGTTGCCA
CTX2_SEPES 3129 bp       2356 TGTCGGGGAGAAACCCAAGATGAACAGTGTAAAGGGTCATCGACAGACAGGCGTTCTTGTAGTAGCGAAGATTGTTGCCA
CTX1_DPEA 3156 bp        2380 TGCAACGGAGAATCTGAGAGTGAAAAATGTGAAGGGTCGGCGACTGATGTGCGCAATTGTTTTCTCACAGATTGCTGTCA
CTX2_DPEA 3123 bp        2350 TGTTTGGGAGAAACTAACGACGAACACTGTAATGGTTCATCAAAAGACACACGCACCTGTAATAAAGAAGATTGTTGCCA
CTX1_SBAN 2946bp         2380 TGTAACGGTCAATCTGAGAGTGAAAGATGCAAAGGGCCGTCAACTCAGGTACGAAAATGTTCCGTCGAAGATTGTTGTCA
CTX2_SBAN 2286 bp        2284 --------------------------------------------------------------------------------
CTX_EBER 3159 bp         2383 TGCCAGGGAGACCCAAGCGGTGATCAATGCCAAGGTCAGGCCGTTGTCAAGCGAAAATGTTTCGTGCAAGATTGTTGTCA
CTX2_SOFF 3129 bp        2356 TGTAGGGGAGAAGACCAAGACCAACAATGCAAAGGCTCATCGACAGACAAACGTAGCTGTAGTAGCGAAGATTGTTGCCA
CTX1_SOFF 3141bp         2365 TGTATCGGAGAATCTGACACTGACAAATGCAAAGGGCAATCGTCTCAAGTGCGGAAATGTTTCGTCGAAGACTGTTGCCA
CTX_ESCO 3159 bp         2383 TGCCAGGGAGACCCAAGCGGTGGTCAATGCCAAGGCCAGGCCGTTGTCAAGCGAAAATGCTTCGTGCAAGATTGTTGTCA
CTX2_SLES 3132 bp        2359 TGTTTAGGAGAAACTAAAAGCGAACATTGTAAAGGTTCATCAAAAGACACGCGCTCTTGTAGTACAGAAGATTGTTGTCA
CTX1_SLES 3159 bp        2383 TGCAACGGAGAATCTGAGGGCGAAAAATGCAAAGGGTTGACAACTGATGTGCGCAACTGTTTTCTCTCAGATTGCTGTCA
CTX_TRHO 3150 bp         2377 TGTCTGGGAGAAACTGAGGAAGAAAAATGTCAAGGCACACCGAAAGACACTCGCACTTGTAGTAATGAAGATTGTTGTCA
CTX_SATL 3147 bp         2380 TGCCAGGGAGAC---------GATCAATGCCAAGGTCGGGCTGTTGCCACGCGAAAATGTTTCATGCAAGATTGCTGTCA
CTX2_SLYC 3129 bp        2356 TGTCTGGGAGAAACCCAAAATGAACAGTGTAAAGGGTCATCGACAGACAGGCGTTCTTGTAGTAACGAAGATTGTTGCCA
CTX1_SLYC 3159 bp        2383 TGTCTGGGACAATCTGAGAGTGAAAAATGCAAAGGGCCGTCAAAACAAGTACGAAAATGTTTCGTCGAAGATTGTTGCCA
CTX_EPAR 3150 bp         2377 TGCCAGGGAGACCCAGGCGGTGATCAATGCCAAGGCCAGGCCGTTGTCAGTCGAAAATGCTTCACGCAAGATTGTTGTCA
CTX2_SPHAR 3129 bp       2356 TGTCTGGGAGAAACCCAAGATGAACAGTGTAAAGGGTCATCGATAGACAGGCGTTCTTGTAGTAGCGAAGATTGTTGCCA
CTX1_SPHAR 3159 bp       2383 TGTCTGGGACAATCTGACAGTGAAAAATGCAAAGGGCCGTCAAAACAGGTACGAAAATGTTTCGTCGAAGATTGTTGCCA
CTX_WSCI_PARTIAL 1653 bp 1582 TGTAACGGGGGCAC------------GTGTGGAGGGGCGTCGACTGATGTTCGTAGTTGTCCATTCAAAGA---------
CTX_IHAL 3156 bp         2389 TGCAATGGAAGC---CCCGGTGAAACATGTGTAGGGCAAGCCAGTCAGGAAAAGAAATGTTTCCCAGAAGATTGTTGTCA
CTX2_XNOT 3183 bp        2410 TGTTTGGGGGAGACTCACACGGAACATTGCAAAGGAGAGCAAGATGAAACACGATCTTGTTCAACAGAAGACTGTTGCCA
CTX1_XNOT2 3150 bp       2383 TGCAATGGAAAT---GAAGGTGAAATATGTGAAGGACCACCTGCACAGAAAAAGAAATGTTTCCCGGATGATTGTTGCCA
CTX_OROB_PARTIAL 1431 bp 1430 --------------------------------------------------------------------------------
CTX_SAFF 3147 bp         2380 TGCCAGGGAGAC---------GATCAATGCCAAGGTCGGCCTGTTGCCACGCGAAAATGTTTCATGCAAGATTGCTGTCA
CTX_SOBS 3156bp          2380 TGCCAGGGAGAATCGGGCGGCGAGCAATGCCAAGGTCAGGCTGTTGCCACGCGAAAATGTTCCATGCAAGATTGCTGTCA
CTX_Sarc 2907 bp         2153 CGCAACGGAG------------GGAAATGCAAAGGGTCGTCAA---------------CTTTTCTCGAAGATTGCTGTCA
CTX2_Dopa 3123 bp        2350 TGTTTGGGAGAAACTAACGACGAACACTGTAATGGTTCATCAAAAGACACACGCACCTGTAATAAAGAAGATTGTTGCCA
CTX1_Dopa 3123 bp        2386 TGCAACGGAGAATCTGAGAGCGAAAAATGTGAAGGGTCGGCCACTGATGGGCGCAATTGTTTTCTCACAGATTGCTGTCA

                         2881 ......2890......2900......2910......2920......2930......2940......2950......2960
CTX1_SEPES 3159 bp       2463 GGAGAAATACGGCAAATTTAAATGCGATAACAATAAATGCATTTCCCTTTCTCGGGTTTGCGATGGAAATGACGACTGTC
CTX2_SEPES 3129 bp       2436 AGCTAAATTTGGCAAGTTTAAGTGCCCAGTTGGTTGGTGTATCGACCTCTCGAAGGTTTGCGACGGGACTACCGATTGTG
CTX1_DPEA 3156 bp        2460 AAAAATCTATGGAAAATTTAAATGCGACGATACTAAATGTCTTGATATTTCACAGTTATGCGATGGATTTGATGACTGTC
CTX2_DPEA 3123 bp        2430 GGAAAAATACGGCAAATTCAAATGTCCAGTTGGTTGGTGTATTGATCTTTCGAGGGTTTGTGACGGGACAGCTGATTGTG
CTX1_SBAN 2946bp         2460 AGAGAAATACGGCAAATTTAAATGCGACAGTGACAAATGTATTCCTCTTTCTCAGGTTTGCGATGGAACTGATAACTGCC
CTX2_SBAN 2286 bp        2284 --------------------------------------------------------------------------------
CTX_EBER 3159 bp         2463 CACGAAATTCGGTAAATTTAAATGCGATAATAGCACATGTCTAAGTATTTCTCAGGTGTGCGATGGAATTGATGATTGTC
CTX2_SOFF 3129 bp        2436 AGCTAAATTTGGCAAGTTTAAGTGTCCGGTTGGTTGGTGTATCGATCTTTCGAGGGTTTGCGACGGGACAACCGATTGTG
CTX1_SOFF 3141bp         2445 GGAGAAATACGGCAAATTTAAATGCGGCAACGACAAATGCATTTCCATTTCTCAGGTTTGCGATGGAATCGATGACTGTC
CTX_ESCO 3159 bp         2463 CGTGAAATTCGGTAAATTTAAATGCGATAATAGCACATGTCTAAGTATTTCTCAGGTGTGCGATGGAATTGACGATTGTC
CTX2_SLES 3132 bp        2439 GGAAAAATTTGGCAAATTCAAATGCCCAATTGGTTGGTGTATTGATCTTTCAAGAGTTTGCGACGGAACAGCCGACTGTG
CTX1_SLES 3159 bp        2463 AGAAATGTACGGCAAATTTAAATGCGACAGTACTAAGTGCCTTGATCTTTCTCAGTTATGCGATGGATTCGACGACTGTC
CTX_TRHO 3150 bp         2457 GGCTAAATTTGGTAAGTTCAAATGCCCGGTAGGTTGGTGTATTGATCTTTTGAGGGTTTGTGACGGGACAACGGACTGTG
CTX_SATL 3147 bp         2451 CGCGAAATTCGGTAAATTTAAATGCGACAATAGCACATGTCTACGCATTTCTCAGGTCTGCGATGGAATTGATGACTGTT
CTX2_SLYC 3129 bp        2436 AGCTAAATTTGGAAAGTTTAAGTGTCCAGTTGGTTGGTGTATCGATCTTTCGAAGGTTTGCGACGGGACTACCGATTGTG
CTX1_SLYC 3159 bp        2463 GGAAAAATACGGCAAATTTAAATGCAACAACAATAAATGCATTTCCCTTTCTCGGGTTTGCGATGGAAATGATGATTGTC
CTX_EPAR 3150 bp         2457 CGCGAAATTCGGCAAATTTAAATGCGATAATAGTACATGTCTGAGTATTTCTCAGGTGTGCGATGGAATTGACGATTGTC
CTX2_SPHAR 3129 bp       2436 AGCTAAATTTGGCAAGTTTAAGTGCCCAGTTGGTTGGTGTATCAATCTCTCGAAGGTTTGCGACGGGACTACCGATTGTG
CTX1_SPHAR 3159 bp       2463 GGAGAAATACGGCAAATTTAAATGTAACAACAATAAATGCATTTCCCTTTCTCGGGTTTGCGATGGAAATGATGACTGTC
CTX_WSCI_PARTIAL 1653 bp 1641 --------------------------------------------------------------------------------
CTX_IHAL 3156 bp         2466 AGCTAAATATGGTAAGTTTCGATGCGACGCCGAAACTTGTTTGGATGTGTCGGATATTTGTAATGGCGTCGAACAATGTT
CTX2_XNOT 3183 bp        2490 AGGGAAATATGGAAAATTTAAATGTCCTGTTGGTTGGTGCCTTGACCTTTCTCAAGTCTGTGATGGAATTACTCATTGTG
CTX1_XNOT2 3150 bp       2460 AGCAAAATACGGTAAATTTAAATGCGATACTGATACTTGTTTGAATCTATCGGACATTTGTAATGGAATTGAACAATGTT
CTX_OROB_PARTIAL 1431 bp 1430 --------------------------------------------------------------------------------
CTX_SAFF 3147 bp         2451 CGCGAAATTCGGTAAATTTAAATGCGACAATAGCACATGTCTACGCATTTCTCAGGTCTGCGATGGAATTGATGACTGTT
CTX_SOBS 3156bp          2460 GGTGAAATTCGGTAAATTTAGATGCGACAACAGCACATGTCTACGTCGTTCTCAGGTCTGCGATGGAATCGATGACTGTT
CTX_Sarc 2907 bp         2206 AGAGA-ATACGGCAAATTTAAATGCGACGATAGTAAATGCCTCGATATTTCCCAGGTATGCGACGGCATCGATCATTGTG
CTX2_Dopa 3123 bp        2430 GGAAAAATACGGCAAATTCAAATGTCCAGTTGGTTGGTGTATTGATCTTTCGAGGGTTTGTGACGGAACAGCCGATTGTG
CTX1_Dopa 3123 bp        2466 AGAAATCTATGGCAAATTTAAATGCGACGATACTAAATGTCTTGATATTTCACAGTTATGCGATGGATTTGATGATTGTC

                         2961 ......2970......2980......2990......3000......3010......3020......3030......3040
CTX1_SEPES 3159 bp       2543 GTAATGCAGAAGACGAGTCGAAAAGTCGGTGCAAGTATCTTCGCTCCGGTGACAGAATCGCCCTACGCAACATGGCCTAC
CTX2_SEPES 3129 bp       2516 CCATTGATGCAGACGAGGCGAAAGGAAGATGTAATTATCTTCGATCCGGTGACAGAATTGCCCTGCGCAATTTAGCTTCA
CTX1_DPEA 3156 bp        2540 ATGATGGAACAGATGAGTCCAAGGACAAATGCAAATATCTTCGGTCAGGTGACAGAATCGCTCTCCGCAACATGGCCTCT
CTX2_DPEA 3123 bp        2510 CCCTTGATGAAGACGAGGCAAAAACAAGGTGTAATTATCTTCGGTCCGGTAACAGAATTGCACTGCGTAATTTGGCTACC
CTX1_SBAN 2946bp         2540 TTGATAAAACAGACGAATTGAAAAGTCGATGCAACTACCTCCGTTCGGGTGACAGAATCGCCCTTCGCAACATGGCCTTC
CTX2_SBAN 2286 bp        2284 --------------------------------------------------------------------------------
CTX_EBER 3159 bp         2543 TAGATGGAACAGACGAGTCGTCTACGAAATGTAAATATTTGCGTTCTGGTAACAGAATCGCCCTTCGTAATATGGACTAT
CTX2_SOFF 3129 bp        2516 CCACTGACGCAGATGAGTCGAAAGGAAGATGCAATTATCTTCGATCCGGTGACAGAATTGCCCTGCGCAATTTAGCTTCC
CTX1_SOFF 3141bp         2525 TTGATGAAAGTGACGAATCGAAAAGTCGTTGCAAATATCTTCGCTCTGGTGACAGAATCGCTCTCCGCAACATGGGCTTC
CTX_ESCO 3159 bp         2543 TAGATGGAACCGACGAGTCGTCTACGAAATGTAAATATTTGCGCTCTGGTAACAGAATCGCCCTTCGTAATATGGACTAT
CTX2_SLES 3132 bp        2519 CCCTTGATGAAGACGAAGCAAAAGCAAGGTGTAATTATCTTCGGTCTGGTAACAGAATTGCACTACGAAATTTGGCAACC
CTX1_SLES 3159 bp        2543 TTGACGGAATAGACGAGTCAAAGGAGAGATGCAAATATCTTCGCTCAGGTGACAGAATCGCCCTCCGCAACATGGCCTTT
CTX_TRHO 3150 bp         2537 CCCTTGATGCAGACGAGGCAAGAGGGAGGTGTAATTATCTCCGGTCCGGTGATAGAATCGCTCTACGTCACTTGGCTTCT
CTX_SATL 3147 bp         2531 CGGATGGAAGCGACGAATCGTCTACGAAATGTAAATATTTGCGCTCCGGTGACAGAATCGCTCTTCGTAACATGGACTAT
CTX2_SLYC 3129 bp        2516 CCATTGATGCAGACGAGGCGAAAGGAAGATGTAATTATCTTCGATCCGGTGACAGAATTGCCCTGCGCAATTTAGCTTCA
CTX1_SLYC 3159 bp        2543 GTGATGCAGCAGACGAGTCGAAAAGTCGGTGCAAATATCTTCGCTCCGGTGACAGAATCGCCCTACGCAATATGGCTTAC
CTX_EPAR 3150 bp         2537 TAGATGGAACCGACGAATCGTCTACGACATGTAAATATTTGCGCTCTGGTGACAGAATCGCCCTTCGTAATATGGACTCT
CTX2_SPHAR 3129 bp       2516 CCATTGATGCAGACGAGGCGAAAGGAAGATGTAATTATCTTCGATCCGGTGACAGAATTGCCCTGCGCAGTTTAGCTTCA
CTX1_SPHAR 3159 bp       2543 TTGATGCAGCAGACGAGTCGAAAAGTCGTTGCAAATATCTTCGCTCCGGTGACAGAATCGCCCTACGCAACATGGCTTAC
CTX_WSCI_PARTIAL 1653 bp 1641 --------------------------------------------------------------------------------
CTX_IHAL 3156 bp         2546 CTGATGGATCGGACGAATCGA------AATGTGATTATCTTCGATCCGGAGATAGAATTGCCCTCCGAAATATGCGTTAC
CTX2_XNOT 3183 bp        2570 CAAGCGATGCAGATGAATCTACAGCCACATGTAATTATTTAAGATCTGGAAAAAGAATAGCACTTCGAAGTTTGGGTTCA
CTX1_XNOT2 3150 bp       2540 CCGATGGATCAGATGAATCAC------AATGTGATTATGTTCGATCTGGGGACCGGATTGCTCTCCGCAACATGCGTTTC
CTX_OROB_PARTIAL 1431 bp 1430 --------------------------------------------------------------------------------
CTX_SAFF 3147 bp         2531 CGGATGGAAGCGACGAATCGTCTACGAAATGTAAATATTTGCGCTCTGGTGACAGAATCGCTCTTCGTAACATGGACTAT
CTX_SOBS 3156bp          2540 CGGACGGAAGCGACGAATCGTCTACGAAATGTAAATATCTGCGCTCTGGTGACAGAATCGCCCTTCGTAACATGGACTCT
CTX_Sarc 2907 bp         2285 TCGATAGGACCGACGAGTCAAAGGCAAGATGCAAATATCTCCGATCAAGAGACAAAATTGCTCTCCGCAATGTTGCCTAT
CTX2_Dopa 3123 bp        2510 CCCTTGATGAAGACGAGGCAAAAACAAGGTGTAATTATCTTCGGTCCGGTAACAGAATTGCACTGCGTAATTTGGCTACC
CTX1_Dopa 3123 bp        2546 ATGATGGTACAGATGAGTCAAAGGACAAATGCAAATATCTTCGCTCAGGTGACAGAATCGCTCTCCGCAACATGGCCTCT

                         3041 ......3050......3060......3070......3080......3090......3100......3110......3120
CTX1_SEPES 3159 bp       2623 TCTCAGGAGTGGCTCAGCGTGCAGTACACTGACGCCGTTCA---AGCCGATTTATATTACGGCCGGGCTTACCTGAACCA
CTX2_SEPES 3129 bp       2596 TCTTTCGACTGGCTCAGCGTTCGTAATACAGACTACCTTAGTACGCCTCAGCTGCGATACGGAAGGGCTTTTCTTGATCG
CTX1_DPEA 3156 bp        2620 TCTCAGGATTGGCTTAGTGTAAAGTACACAGATGCCGTTCA---GGCTGATTTATATTACGGTCGGGCCTACCTGGATCG
CTX2_DPEA 3123 bp        2590 TCTTCTGATTGGCTCAGCGTCCTTTATACAGATTACGTTGTTACTCCTCATTTGCGATATGGAAGAGCTTATCTTAATCG
CTX1_SBAN 2946bp         2620 TCTCAGGAGTGGCTAAGCGTGCAGTACACGGACTACGTTCA---AGCCAATTTATATTACGGCCGGGCTTACTTGGATCA
CTX2_SBAN 2286 bp        2284 --------------------------------------------------------------------------------
CTX_EBER 3159 bp         2623 TCGGCGGATTGGCTCAGCGTCCAGTATACAGACTATGTTCA---AGCCAATTTATATTATGGTCGGGCGTATTTAGACCG
CTX2_SOFF 3129 bp        2596 TCTGTCGACTGGCTCAGCGTTCTTAGAACAGACTACCTTAACAGCCCTCAATTGCAATACGGAAGAGCTTTTCTCGATCG
CTX1_SOFF 3141bp         2605 TCCCAGGAGTGGCTCAGCGTGCAGTACACAGACGCAGTTCA---GGCCAATCTATATTACGGTCGGGCCTACCTGAACCA
CTX_ESCO 3159 bp         2623 TCGATGGATTGGCTCAGCGTCCAGTATACAGACTACGTTCA---AGCCAATTTATATTATGGTCGGGCGTATTTAGACCG
CTX2_SLES 3132 bp        2599 TCTTCTGACTGGCTCAGTGTTCGTTATACAGATTACGTTATTCCTCCTCATTTGCGATATGGAAGAGCTTATCTTAATAG
CTX1_SLES 3159 bp        2623 TCCCAAGAGTGGCTTAGTGTACAGTACACAGATACCGTCCA---AGCTGATTTATATTACGGTCGAGCCTACCTGGACCA
CTX_TRHO 3150 bp         2617 TCTTTCGATTGGCTCAGCCTTCGTCATACAGATCGCATTATCACCCCTTCTTTACGATATGGAAGAGCTCATCTTGATCG
CTX_SATL 3147 bp         2611 TCGCAGGATTGGATCAGTGTTCAGTATACGGACTACGTTCA---AGCCAATTTATATTATGGCCGGGCGTATTTAGACCG
CTX2_SLYC 3129 bp        2596 TCTTTCGACTGGCTCAGCGTTCGCAATACAGATTACCTTAACACTCCTCATCTGCGATATGGAAGAGCCTTTCTTGATCG
CTX1_SLYC 3159 bp        2623 TCTCAGGAGTGGCTCAGCGTACAGTACACTGACGCTGTTCA---AGCCAATTTATATTACGGCCGGGCTTACCTGAACCA
CTX_EPAR 3150 bp         2617 TCGATGGATTGGCTTAGTGTCCAGTATACAGACTACGTTCA---AGCGAATTTATATTATGGTCGAGCGTATTTAGACCG
CTX2_SPHAR 3129 bp       2596 TCTTTCGACTGGCTCAGCGTTCGTCATACAGATTACCTTAACAGTCCTCAGTTGCGATACGGAAGAGCTTTTCTTGATCG
CTX1_SPHAR 3159 bp       2623 TCTCAGGAGTGGCTCAGCGTGCAGTACACTGACGCTGTTCA---AGCCAGTTTATATTACGGCCGGGCTTACCTGAACCA
CTX_WSCI_PARTIAL 1653 bp 1641 --------------------------------------------------------------------------------
CTX_IHAL 3156 bp         2620 TCACAAGACTGGCTCAGCGTCCAATATACAGACGCCGTACA---AGCTGATCTTCATTACGGGCGCGCCTATCTCGACGA
CTX2_XNOT 3183 bp        2650 TCAAAGAATTGGCTTAGCGTATTGTATTCTGATTACGTGAGCCGGGGCGACCTTCGTTATGGAAGGGCACACGTGACTCA
CTX1_XNOT2 3150 bp       2614 TCTCAGGATTGGCTCAGCGTCAAAAATACAGACGCTGTACA---AGCTAGTCTTTATTACGGACGCGCCTACCTTGACGA
CTX_OROB_PARTIAL 1431 bp 1430 --------------------------------------------------------------------------------
CTX_SAFF 3147 bp         2611 TCGCAGGATTGGATCAGTGTTCAGTATACAGACTACGTTCA---AGCTAATTTATATTATGGTCGGGCGTATTTAGACCG
CTX_SOBS 3156bp          2620 TCGCAGGATTGGATCAGTGTCCAGTACACAGACTACGTTCA---GGCCAATCTATATTATGGCCGCGCCTATTTAGACCG
CTX_Sarc 2907 bp         2365 TCACAGGAGTGGCTCAGCGTGCAATATACAGATTACGTACA---GTCCAATTTATATTACGGTCGGGCCTACCCGGACCA
CTX2_Dopa 3123 bp        2590 TCTTCTGATTGGCTTAGCGTCCTTTATACAGATTACGTTGTTTCTCCTCATTTGCGATATGGAAGAGCTTATCTTAATAG
CTX1_Dopa 3123 bp        2626 TCCCAGGATTGGCTCAGTGTACAGTACACAGATGCCGTTCA---GGCTGATTTATATTACGGTCGGGCCTACCTGAATCG

                         3121 ......3130......3140......3150......3160......3170......3180......3190......3200
CTX1_SEPES 3159 bp       2700 TTGTATTAAAGGAGACCACGTAACTTCCAGTGAGTGGAATTCTTGTGCTGGTCAGTCATTGCTTATTTATGGAAATTACG
CTX2_SEPES 3129 bp       2676 CTGTATTAAAACTAATCGGGTCATCGAATCCGAATGGCGAGAATGTGTCGGTAACTCGATGCTTATTTATGGTAGTCATA
CTX1_DPEA 3156 bp        2697 TTGCATTAAGGGCGATCAGGTCACTGATTACGAATGGAAGAACTGTGCCGGTCAATCAATGCTTATTTATGGAAATTATA
CTX2_DPEA 3123 bp        2670 ATGCATTAAGTCAGAGAGGGTTGCTGCATCAGAATGGAAGCAGTGTGAAGGTAATTCGATGCTCGTCTACGGAAGTCACA
CTX1_SBAN 2946bp         2697 TTGCATTAAAGGCGACCGCGTGACCTCTAGTGAATGGGATTCTTGCGCTGGTCAGTCAATGCTAATTTATGGAAATTACG
CTX2_SBAN 2286 bp        2284 --------------------------------------------------------------------------------
CTX_EBER 3159 bp         2700 CTGCATTAAAGGAGACGATGTTACTGATTACGAATGGAAAAACTGCCCTGGGCAGTCGATGCGTGTTTACGGAAATTACC
CTX2_SOFF 3129 bp        2676 CTGTATTAAAACTCATCATATCACCGAATCCGAATGGCGACAATGCAAAGGTAACTCGATGCTTATTTATGGTAGTCACA
CTX1_SOFF 3141bp         2682 TTGCATTAAAGGAACCCAAGTGACGTCAAGTGAATGGAAATCTTGTCCAGGCCAGTCAATGCTTATTTATGGAAATTACA
CTX_ESCO 3159 bp         2700 CTGCATTAAAGGAGACGATGTTACTGATTACGAATGGAAAAACTGCCCTGGGCAGTCGATGCTTGTTTACGGAAATTACC
CTX2_SLES 3132 bp        2679 ATGTATTAAGACCGAAAAGGTTACTGCATCAGAATGGAAGCAATGTGAAGGTAATTCGATGCTTGTCTATGGAAGTCACA
CTX1_SLES 3159 bp        2700 TTGCATTAAGGGGGATGACGTCACTGATAATGAATGGAAGAATTGTGCTGGTCAGTCAATGCTTATTTATGGAAATTACA
CTX_TRHO 3150 bp         2697 ATGTATTAAAACTGATCAGGTCACTGCACCGGAATGGAAACAGTGTATGGACAATTCGATGCTTGTCTATGGGAGTCACA
CTX_SATL 3147 bp         2688 CTGCATTAAAGGTGACGAGGTTACTGATAACGAATGGAAGAACTGCCCTGGACAGTCGATGCTTGCTTACGGAAATTACG
CTX2_SLYC 3129 bp        2676 TTGTATTAAAACTAATCAGGTCCTTGAATCCGAATGGCGAGAATGTAACGGTAACTCCATGCTTATTTATGGTAGTCATA
CTX1_SLYC 3159 bp        2700 TTGCATTAAAGGAGACCGCGTGACTTCCAGTGAATGGAATTCTTGTGCTGGTCAGTCATTGCTTATTTATGGAAATTACC
CTX_EPAR 3150 bp         2694 CTGCATTAAAGGAGACGACGTTGCTAGTGACGAATGGAAAAACTGCCCTGGGCAGTCGATGCTTGTTTACGGAAATTACC
CTX2_SPHAR 3129 bp       2676 TTGTATTAAAACTAATCAGGTCCTCGAATCCGAATGGCGAAAATGTGACGGCAACTCGATGCTTATTTATGGTAGTCATA
CTX1_SPHAR 3159 bp       2700 TTGCATTAAAGGAGACCACGTGACTTCCAGTGAATGGAATTCTTGTGCCGGTCAGTCATTGCTTATTTATGGAAATTACC
CTX_WSCI_PARTIAL 1653 bp 1641 --------------------------------------------------------------------------------
CTX_IHAL 3156 bp         2697 TTGCATCCAAGATGATGACGTGACCGACGATGAATGGAAGTACTGTGCTGGCCAGTCGTTACGTATTTACGGAAATTACC
CTX2_XNOT 3183 bp        2730 GTGCATAAAAGGTGATTTTGTTGCAAGTTCCGAATGGACAGACTGTCCTGGTAATTCAATGCTTATCTACGGCAGCTACA
CTX1_XNOT2 3150 bp       2691 CTGCATCCATGATGATGATGTAACCGATGATGAATGGAAGTACTGTGATGGTCAGTCTTTACGTATTTACGGAAATTATC
CTX_OROB_PARTIAL 1431 bp 1430 --------------------------------------------------------------------------------
CTX_SAFF 3147 bp         2688 CTGCATTAAAGGTGACGAGGTCACTGATAACGAATGGAAGAACTGCCCTGGACAGTCGATGCTTGCTTACGGAAATTACG
CTX_SOBS 3156bp          2697 CTGTATTAAAGGTGACGACGTCACTGAGAACGAATGGAAGAACTGTCCTGGACAATCGATGCTTGCTTACGGAAATTACG
CTX_Sarc 2907 bp         2442 TTGCATTAAGGATGATCACGTTACTGATAGTGAATGGAAGAATTGTGTCGGTCAATTGATGAATGTTTACGGAAGTTACA
CTX2_Dopa 3123 bp        2670 ATGTATTAAGTCAGAGAGGGTTGCTGCATCAGAATGGAAGCAGTGTGAAGGTAATTCGATGCTCGTCTACGGAAGTCACA
CTX1_Dopa 3123 bp        2703 TTGCATTAAGGGCGATGAGGTCACTGATTACGAATGGAAGAACTGTGCGGGTCAATCGATGCTTATTTATGGAAATTATA

                         3201 ......3210......3220......3230......3240......3250......3260......3270......3280
CTX1_SEPES 3159 bp       2780 AAAACGGGAAAATAGGAAAAGCCATCAGATTCGGTGATAAAATCGCTATGTATTACCGTAAAACGAACTATCATTATCGC
CTX2_SEPES 3129 bp       2756 ATAACGCAAGAATAGGCCAATCGATTAGGTATGGCGACCGTATTGCCATGTATTACCGAAAAACTCATTCCCATTTTCGT
CTX1_DPEA 3156 bp        2777 AGAACGGTGAAACAGGAAAGGCCATCAAATATGGCGATAAAATCGCCATTTATTATCGCAAAACAAACTTTCATTATCGC
CTX2_DPEA 3123 bp        2750 ACCAAGGAAGAGATGGTCAATCAATTAGATATGGCGACCGCATTGCTTTGTATTACCGAAAAACGTATTCCGAATATCGT
CTX1_SBAN 2946bp         2777 AAAATGGGAAAACAGGAAAAGCCATCAGATTCGGTGATAAAATCGCCATGTATTACCGTAAAACGGAGTTTCATTATCGC
CTX2_SBAN 2286 bp        2284 --------------------------------------------------------------------------------
CTX_EBER 3159 bp         2780 AGAACGGTAAATCTGGCAAGGCTATTAGGTATGGCGATAAGATCGCCCTCTACTATCGCAAAACCAATTATCATTATCGC
CTX2_SOFF 3129 bp        2756 ATAACGCAAGAGTAGGTCAATCGATTAGGTATGGCGATCGTATTGCCATGTATTACCGAAAAACCTATTCCCAATTTCGT
CTX1_SOFF 3141bp         2762 AAAACGGCAAAGCAGGAAAAGCGATCAGATTCGGTGATAAAATCGCTATGTATTACCGTAAAACTGCGTATCATTATCGA
CTX_ESCO 3159 bp         2780 AGAACGGTAAATCTGGCAAGGCTATTAGGTATGGCGATAAGATCGCCCTCTACTATCGCAAAACCAATTATCATTATCGC
CTX2_SLES 3132 bp        2759 ACCAAAGAAGAGAAGGTCAGTCAATTAGATATGGCGATCGCATTGCTTTGTACTACCGAAAAACGTATTCCAAATATCGC
CTX1_SLES 3159 bp        2780 AGAACGGCCAAATAGGAAAGGCCATCAAATATGGCGATAAAATCGCCATGTATTATCGCAAAACAAATTATCATTATCGC
CTX_TRHO 3150 bp         2777 ATAACGGAAGAGAAGGCCAATCAATCCGATACGGTGATCGTATTGCTATGTACTACCGAAAAACGTATTCCCACCATCGC
CTX_SATL 3147 bp         2768 AGAACGGTAAAACTGGCACGGCTATAAGGTATGGTGATAAGATCGCTCTCTACTACCGCAAAACCAATTATCATTATCGC
CTX2_SLYC 3129 bp        2756 ATAACGCAAGAATTGGCCAATCTATTAGGTATGGCGACCGTATTGCCATGTATTACCGAAAAACTCATTCCCAGTTTCGT
CTX1_SLYC 3159 bp        2780 AAAACGGGAAAAAAGGAAAAGCCATCAGATTCGGTGATAAAATAGCTATGTATTACCGTAAAACGAACTATCATTATCGC
CTX_EPAR 3150 bp         2774 AGAACGGTAAATCTGGTAAGGCTATTAGGTATGGCGATAAGATCGCCCTCTACTACCGCAAAACCAATTATCATTATCGC
CTX2_SPHAR 3129 bp       2756 ATAACGCAAGAATTGGCCAATCGATTAGGTATGGCGACCGTATTGCCATGTATTACCGAAAAACTCACTCCCATTTTCGT
CTX1_SPHAR 3159 bp       2780 AAAACGGGAAAAAAGGAAAAGCCATCAGATTCGGTGATAAAATCGCTATGTATTACCGTAAAACGAACTATCATTATCGC
CTX_WSCI_PARTIAL 1653 bp 1641 --------------------------------------------------------------------------------
CTX_IHAL 3156 bp         2777 AGAACGGCAAAACTGGAAAAGCCATCAGATATGGAGATCGAGTCGCACTCTATTACAGAAAAAGTAATTATCACTATCGA
CTX2_XNOT 3183 bp        2810 ATAATGCAAGGAAAGGTCAGGCAATCAGATATGGGGATCGAATTGCAATGTACTATACAAAGACAAATGCAAAATATCGA
CTX1_XNOT2 3150 bp       2771 AAAACGGGAAAACTGGAAAACCTATCAGATTCGGAGATCGAATCGCTTTGTATTACCGAAAAACTAATTATCACTACAAG
CTX_OROB_PARTIAL 1431 bp 1430 --------------------------------------------------------------------------------
CTX_SAFF 3147 bp         2768 AGAACGGTAAAACTGGCAAGGCTATTAGGTATGGTGATAAGATCGCTCTCTATTACCGCAAAACCAATTATCATTATCGC
CTX_SOBS 3156bp          2777 AAAACGGTAAGACCGGCAAGGCTATTAGGTATGGTGATAAGATCGCCCTCTACTACCGCAAGACCAATTACCATTATCGG
CTX_Sarc 2907 bp         2522 AGAACGGAAAACAAGGGACGGCCATCAGATACGGTGAAAGAATCGTTATGTATTACCGCAAAACGAATTATCATTGGAGA
CTX2_Dopa 3123 bp        2750 ACCAAGGAAGAGATGGTCAATCAATTAGATATGGCGACCGCATTGCTTTGTATTACCGAAAAACGTATTCCGAATATCGT
CTX1_Dopa 3123 bp        2783 AGAACGGTGAAAAAGGAAAGGCCATCAAATATGGCGATAAAATCGCCATGTATTATCGCAAAACAAACTATCATTACCGC

                         3281 ......3290......3300......3310......3320......3330......3340......3350......3360
CTX1_SEPES 3159 bp       2860 TGGTTTATTTGTTACCCAACTTACTGTATGACTTATACTTGTCCTAAAAAAGCTGGATCATTTACTTTTGGTCCT-AACG
CTX2_SEPES 3129 bp       2836 TGGTTCAGGTGTTACTCTGATTATTGTATAACGTACACTTGCGATAAAGCTCCAGGTCAGTTCGATTTTAGCGAC-AGAG
CTX1_DPEA 3156 bp        2857 TGGCTCCGTTGTTATTCGTCTTACTGCATGACATATACCTGTGACAAAAAACCGGGAACTTTTGATTTTGGTTCC-AACG
CTX2_DPEA 3123 bp        2830 TGGTTCAGATGTTATTCTGATTATTGTATATCGTATATCTGCTCAAAAGCTCCGGGCCAGTTCGATTTTAGTGAC-ACAG
CTX1_SBAN 2946bp         2857 TGGTTTATTTGCTATCCAACTTATTGTATGACTTACACTTGCGATAAAAAGTGGGGATCATTTGATTTTGGTTCCCAACG
CTX2_SBAN 2286 bp        2284 --------------------------------------------------------------------------------
CTX_EBER 3159 bp         2860 TGGTTCCGGTGTTATACGAAATATTGTATGACTTACACTTGCGAGGTGAAATATGGAAGTTTTGATTTCAGTAGT-AAAG
CTX2_SOFF 3129 bp        2836 TGGTTCAGATGTTACTCTGATTATTGTATAACGTACACTTGCGATAAAGCTCCGGGAAAGTTCGATTTTAGCGAC-ATAG
CTX1_SOFF 3141bp         2842 TGGTTTATTTGTTACCCAACTTACTGTATGACATACACTTGTCCTAAAAAGCCGGGATCGTTTACTTTTGGCCCC-GGCG
CTX_ESCO 3159 bp         2860 TGGTTCCGGTGTTATACGAAATACTGTATGACTTACACTTGCGAGGTGAGATATGGAAGTTTTGATTTCAGTAGT-AAAG
CTX2_SLES 3132 bp        2839 TGGTTCAGATGTTACGCTGATTATTGTATAACGTACATTTGCGCAAAAGCTCCGGGTCAGTTCGATTTTAGTGAT-AGAG
CTX1_SLES 3159 bp        2860 TGGTTCATTTGTTATCCGGATAACTGCATGACGTATATCTGTGACAAAAGCCCGGGAACTTTTGACTTTACTTCC-AAAG
CTX_TRHO 3150 bp         2857 TGGTTCAGATGTTACTCTAATTATTGTATGACGTACACTTGCACGAAATCTCCAGATCAGTTCGATTTTAGTGAC-GTTG
CTX_SATL 3147 bp         2848 TGGTTCCGTTGTTACACGAAATACTGTATGACTTACACCTGCGAGGTAAGATATGGAAGTTTTGATTTCAGCAGC-AAAG
CTX2_SLYC 3129 bp        2836 TGGTTCAGATGTTACTCTGATTATTGCATAACGTACACTTGCGATAAAGCTCCAGGTCAGTTCGATTTTAGCGAC-AGAG
CTX1_SLYC 3159 bp        2860 TGGTTTATTTGTTACCCAACTTACTGTATGACATATACATGCCCTAAAACTGCTGGATCATTTACTTTTGGTCCC-AACG
CTX_EPAR 3150 bp         2854 TGGTTCCGGTGTTATACGAAATACTGTATGACTTACACTTGCGAGGTGAGACATGGAGATTTTGATTTCAGTAGT-AAAG
CTX2_SPHAR 3129 bp       2836 TGGTTCAGATGTTACTCTGATTATTGTATAACGTACACTTGCGATAAAGCTCCAGGTCAGTTCGATTTTAGCGAC-AGAG
CTX1_SPHAR 3159 bp       2860 TGGTTCATTTGTTACCCAACTTACTGTATGACTTATACTTGTCCTAAAAAAGCTGGATCATTTACTTTTGGTCCC-AACG
CTX_WSCI_PARTIAL 1653 bp 1641 --------------------------------------------------------------------------------
CTX_IHAL 3156 bp         2857 TGGTTCATTTGTTATCCGACTTATTGTATGACTTACACTTGTGATAAAAGATACGGTGAGTTTGATTTTAACCCG-AGAG
CTX2_XNOT 3183 bp        2890 TGGTTCACGTGTTACCAGTCGTATTGTATGACGTATATTTGTAACAAGGCACCGGGAGAATTTGATTTTGGTGAA-TTCG
CTX1_XNOT2 3150 bp       2851 TGGTTCATTTGTTATCCTACTTATTGTATGACTTTTACTTGTGATAAAAGATATGGTTCGTTCGATTTTAACCCG-CAGG
CTX_OROB_PARTIAL 1431 bp 1430 --------------------------------------------------------------------------------
CTX_SAFF 3147 bp         2848 TGGTTCCGTTGTTACACGAAATACTGTATGACTTACACCTGCGAGGTAAGATATGGAAGTTTTGATTTCAGCAGC-AAAG
CTX_SOBS 3156bp          2857 TGGTTCCGGTGTTATACGAAATACTGTATGACTTACACTTGCGAGGTAAGACACGGAAGTTTTGATTTCAGTAGT-AAAG
CTX_Sarc 2907 bp         2602 TGGTTCATCTGTTACCCAAAATACTGTATGACTTATACCTGCGGGAAAAAAGCGGGGACTTTTGATTTTGGTCAT-GAAG
CTX2_Dopa 3123 bp        2830 TGGTTCAGATGTTATTCTGATTATTGTATATCGTACATTTGCTCAAAAGCTCCGGGTCAGTTCGATTTTAGTGAC-ACAG
CTX1_Dopa 3123 bp        2863 TGGTTCCGTTGTTACTCGTCTTACTGCATGACATATACCTGTGACAAAAAATGGGGAACTTTTGATTTTCGTTCC-AACG

                         3361 ......3370......3380......3390......3400......3410......3420......3430......3440
CTX1_SEPES 3159 bp       2939 GAGGCTGTGACGAGTACGAATTTTACATTATTAACTATAACGATAAGCTATCGAGGGACCCAGTAAAGCCTGGAGATGTG
CTX2_SEPES 3129 bp       2915 GTGGATGCAAATCATATGAATTCGTTATTCAAAACTATGAGCATCCCAACGCAACGACTCCAATTCAAAATGGCGATATC
CTX1_DPEA 3156 bp        2936 GAGGCTGTCAAGAATACGAATTTTACATTACAAAATATGACGATCCCCTAGCAAATGGCCCGGTAAAAGCTGGAGATTTA
CTX2_DPEA 3123 bp        2909 GTGGGTGCAAATCAAATGAATTCATCATTCAAAACTATGAACATCCCAATTCGACAACCCCCATTCGAAATGGAGACATT
CTX1_SBAN 2946bp         2937 GAGGTT--------------------------------------------------------------------------
CTX2_SBAN 2286 bp        2284 --------------------------------------------------------------------------------
CTX_EBER 3159 bp         2939 GGGGGTGTAAAGAGTACGAATTCGACATTACAAACTACGACGATCCTCAAGCTAGGGGTCCAGTAAAGGCGGGAGATGTG
CTX2_SOFF 3129 bp        2915 GTGGATGTAAACAATACGAATTCGTTATTCAAAACTATGAATATCCTAATGCGACGACGCCAATTCGAAATGGTGACATC
CTX1_SOFF 3141bp         2921 GAGGCTGTGACGAGTATGAATTTTATATTAGCGACTATAATGATAAACTATCGAGGAAACCAGTAAAGGCTGGAGATATA
CTX_ESCO 3159 bp         2939 GGGGATGTAAAGAGTACGAATTCGACATTATAAACTACGACGATCCTCAAGCTAGGGGTCCAGTAAAGGCGGGAGATGTG
CTX2_SLES 3132 bp        2918 GTGGATGCAAGTCAAACGAATTCATCATTCAAAACTATGAACATCCTAACATGACGACGCCAATTCGAAATGGAGACATT
CTX1_SLES 3159 bp        2939 GAGGCTGTCAAGAATACGAATTTTACATTACAAAATATGATGATCGTCAGGCGATTGGTCCAGTAAAATCTGGAGATATG
CTX_TRHO 3150 bp         2936 GTGGGTGCAAACAGTACGAATTCGTGATTCAAAACTATGAACATCCCAACTCAACGGCGCCTGTTCGAAATGGCGACATT
CTX_SATL 3147 bp         2927 GAGATTGTAAAGAGTACGAATTCGACATCACAAACTACGACGATCCCCAGGCTAGGGGTCCAGTAAAGGCGGGAGATGTG
CTX2_SLYC 3129 bp        2915 GTGGATGCAACTCATACGAATTCATTATTCAAAACTATGAGCATCCCAACGCGACGACTCCAGTTCAAAATGGCGATATC
CTX1_SLYC 3159 bp        2939 GAGGCTGTGACGAGTACGAATTTTACATTATTAACTATAACGATAATCTATCGAGGGACCCAGTAAAGGCTGGAGATGTG
CTX_EPAR 3150 bp         2933 GGGGGTGTAAAGAGTATGAATTCGACATTAAAAACTACGACGATCCTCAGGCTAGGGGTCCAGTAAAGGCGGGAGATGTG
CTX2_SPHAR 3129 bp       2915 GTGGATGCAAATCATACGAATTCGTTATTCAAAACTATGAGCATCCCAACGCGACGACTCCAATTCGTAATGGCGATATC
CTX1_SPHAR 3159 bp       2939 GAGGCTGTGACGAGTACGAATTTTACATTATTAACTATAACGATAAACTATCGAGGGAACCAGTAAAGGCTGGAGATGTA
CTX_WSCI_PARTIAL 1653 bp 1641 --------------------------------------------------------------------------------
CTX_IHAL 3156 bp         2936 GAGAGTGCAAAGAATACCAGTTCTATCTTGCTGACTATAATAATCGAGGATCGAGAGCCCCTGTTAAGCCTGGGGATATT
CTX2_XNOT 3183 bp        2969 GCGGTTGTAAAGAAAACGAGTTTATAATCCGGAATTTTCAGAATCCGAATGCGACGAATCCAATTCGAAACGGGGACATT
CTX1_XNOT2 3150 bp       2930 GAGAGTGCAAGGATTACCAGTTTTATGTTGCTGATTATAACAATCCAGCATCGCGGGCCCCTGTTAAGCCTGGAGATGTT
CTX_OROB_PARTIAL 1431 bp 1430 --------------------------------------------------------------------------------
CTX_SAFF 3147 bp         2927 GAGATTGTAAAGAGTACGAATTCGACATTACAAACTACGACGATCCCCAGGCTAGGGGCCCAGTAAAGGCGGGAGATGTG
CTX_SOBS 3156bp          2936 GCGATTGTAAAGAGTACGAATTCGACATTACAAACTACGACGATCCCCAGGCTCGGGGACCGGTAATGGCGGGAGATGTG
CTX_Sarc 2907 bp         2681 GAAGATGTGAGGAGTACGAATTTTGCATTGAGAAATATGACGATCTCCAAGCACGCGGCCCCGTGAAGGCCGGAGATGTG
CTX2_Dopa 3123 bp        2909 GTGGGTGCAAATCAAATGAATTCATCATTCAAAACTATGAACATCCCAATTCGACGACCCCAATTCGAAATGGAGACATT
CTX1_Dopa 3123 bp        2942 GAGGCTGTAAAGAATACGAATTTCACATTACAAAATATGACGATCGTCAAGCAAATGGCCCGGTAAAAGCTGGAGATTTA

                         3441 ......3450......3460......3470......3480......3490......3500......3510......3520
CTX1_SEPES 3159 bp       3019 ATCACCCTTGCTAACAACAGAGGATCTGTAAAAGGCAACGGCTATAACAGAAATATTAATATAAATGATTGTACAGTGAA
CTX2_SEPES 3129 bp       2995 GTGTTTATACACAACAGCAACGGTGCTCTGAGAGGAAACGGTTACTGGAATAAAATCAATCAAAGGAAATATTCATCTG-
CTX1_DPEA 3156 bp        3016 GTCATAATTTCTGACGGAAGAGGCGCTTTGAAAGGAAATGGATATTACAAAAATATATATCAAGATGGTTGTATTGTCAA
CTX2_DPEA 3123 bp        2989 GTGATCATTTCTGCTGATAAAGGAGCTTTAAGGGGAAATGGTTACAATCATAAAATCACTCTTAAGAAATGTTCCTTTG-
CTX1_SBAN 2946bp         2943 --------------------------------------------------------------------------------
CTX2_SBAN 2286 bp        2284 --------------------------------------------------------------------------------
CTX_EBER 3159 bp         3019 GTGGTCGTTTCAAACGGAAACGGTGCCATAAGAGGCAAGGGGTATTACAAAGAGATAAGTCAAAGTGATTGCGTTAAAAA
CTX2_SOFF 3129 bp        2995 GTGTTTATACACAATAGTAACGGTGCTTTGAAAGGTAACGGTTACTGGAATAAAATCACGCAAAAGAAATGTTCATATG-
CTX1_SOFF 3141bp         3001 ATCACTATTGCTAACAACAGAGGATCCATAAAAGGCAACGGCTATAATAGAAATATCAATATAAATGACTGTACCGTAAA
CTX_ESCO 3159 bp         3019 GTGGTCGTTTCAAACGGTAACGGTGCCATAAGAGGCAAGGGGTATTACAAAGAGATAAATCAAAGTGATTGCTTTAAAAA
CTX2_SLES 3132 bp        2998 GTGGTTATTTCCAATGATAACGGTGCTTTGAAGGGAAATGGTTACAATAATAAAATCACCACAAAGAAATGTTCATCTG-
CTX1_SLES 3159 bp        3019 GTCTTCATTTCTGACGGTAAAGGAGCTATAAAAGGAAATGGCTATTATAAAAATATAAATCAAGATCTTTGCGCTATCAA
CTX_TRHO 3150 bp         3016 GTGTTCATCTCTAATAATGACGGTGCTTTGAAGGGCAACGGATACTGGAATCAAATCAGCCAAAAGAAATGTTCATATG-
CTX_SATL 3147 bp         3007 GTCGTCATTTCAGACGGAAAGGGTGCCGTGAGAGGCAACGGCTATTACAAAGAGATAAGTCAGAGTGATTGCGTTCAAAA
CTX2_SLYC 3129 bp        2995 GTGTTTATACACAATAGTAATGGTGCTCTGAGAGGTAACGGTTACTGGAATAAAATCAATCAAAAGAAATGTTCATCTG-
CTX1_SLYC 3159 bp        3019 ATCACCATTGCTAACAACAGAGGATCTGTTAAAGGCAACGGCTATAATAGAAATATTAATATGAATGATTGTACCGTGAA
CTX_EPAR 3150 bp         3013 GTGGTCGTTTCAAACGGAAATGGTGCCGTAAAAGGCAACGGGTATTACAAAGAGGTAAGTCAAAGTGATTGCGTTAAAAA
CTX2_SPHAR 3129 bp       2995 GTGTTTATACACAATAGTAATGGTGCTCTGAGAGGTAACGGTTACTGGAATAAAATCAATCAAAAGAAATGTTCATCTA-
CTX1_SPHAR 3159 bp       3019 ATCACCATTGCTAACAACAGAGGATCTATAAAAGGCAACGGCTATAATAGAAATATTAATACAGATGATTGTACCGTGAA
CTX_WSCI_PARTIAL 1653 bp 1641 --------------------------------------------------------------------------------
CTX_IHAL 3156 bp         3016 ATTACCATATCGAATGGCAGAGGGTCAATCAAAGGCAATGGACATTCGAAAACAATTAATCAAGACGACTGCACTGTAAG
CTX2_XNOT 3183 bp        3049 GTCTTTATTTCAAACAAAGATGGATCGTTGAAAGCAGGAAGTTTGTGGGACGATATAACTCTTAAGAAATGCTCTTCAG-
CTX1_XNOT2 3150 bp       3010 ATTACCATATCGAATGGAAGAGGAGCCATAAAGGGTAATGGTCATTCCAAAACCATAAATCAAGATGATTGCACTGTAAG
CTX_OROB_PARTIAL 1431 bp 1430 --------------------------------------------------------------------------------
CTX_SAFF 3147 bp         3007 GTCGTCATTTCAGACGGAAAGGGTGCCGTGAGAGGCAACGGCTATAACAAAGAGATAAGTCAAAGTGATTGCGTTCAAAA
CTX_SOBS 3156bp          3016 GTTGTCATTTCAAACGGAAACGGTGCAGTGAGAGGCAACGGCTATTACAAAGAGATAAGTCAAAGCGATTGCGTGAAGAA
CTX_Sarc 2907 bp         2761 GTCGCCATCTCTAACAGTAGAGGAGCTTTAAAAGGTAACGGATATTACAAAGATATAGGTCAGGATGTTTGTATTCATTT
CTX2_Dopa 3123 bp        2989 GTGATCATTTCTGCTGATAACGGGGCTTTAAAGGGAAATGGTTACAATCGCAAAATCACTCTTAAGAAATGTTCCTTTG-
CTX1_Dopa 3123 bp        3022 GTCATCATTTCTGACAGAAGAGGCGCTTTGAAAGGAAATGGATATTACAAAAGTATAAATCAAGATGGTTGTATTACCAA

                         3521 ......3530......3540......3550......3560......3570......3580.......
CTX1_SEPES 3159 bp       3099 ACGAGCACAGGA------CGACAGAATCGAATGCAATGCCAACGCTTGGCAAATCTTTATCAAATAG
CTX2_SEPES 3129 bp       3074 -CAGA----AAT------TGAATCTGCAGAGTGCAAAGAAATTGCATGGCAAATTTTTATTCATTAA
CTX1_DPEA 3156 bp        3096 ACGAGTCTTGGA------CAATAATATCAATTGTAGTGCTAATGCTTGGCAAATCTTTATAAAGTAA
CTX2_DPEA 3123 bp        3068 -AGGA----ACT------TGCCTCTTCAAAGTGTAAAGATATTGCTTGGCAAATATTTATTCAATAA
CTX1_SBAN 2946bp         2943 ---------------------------------------------------------------GTAA
CTX2_SBAN 2286 bp        2284 ----------------------------------------------------------------TGA
CTX_EBER 3159 bp         3099 GCGAGAAAGGAA------TAGAAGTATAAATTGTAAAGCGAGCAGTTGGCAGATTTATATTCAATAG
CTX2_SOFF 3129 bp        3074 -CAGA----ACT------TGATTCTTCAGAGTGCAAAGGAATTACATGGCAAATCTTTATTCAGTAA
CTX1_SOFF 3141bp         3081 TCGCGCGCTGGA------CAACAGAATTGAATGCAATGCCAACGCTTGGCAAATCTTTATGCAATAG
CTX_ESCO 3159 bp         3099 GCGAGAGAGGAA------TAGAAGTATAAATTGTAAAGCGAGCAGTTGGCAGATTTATATTCAATAG
CTX2_SLES 3132 bp        3077 -CGGA----AAT------TGTTTCTTCAAAATGCAAAGATATTGCTTGGCAAATCTTTATTCAGTAA
CTX1_SLES 3159 bp        3099 AAGAGTTCAAAA------CAAAAATCTAACTTGTCATGCTAACACTTGGCAAATCTTCATAAAATAG
CTX_TRHO 3150 bp         3095 -CGAC----AAT------TGTCTCATCACAGTGCAAAGAAATTGCTTGGCAAATTTTTATTCAGTAA
CTX_SATL 3147 bp         3087 ACGACAACGGAA------TAGAAATATGAATTGTAAAGCGAGCAGTTGGCAGATTCATATCAAATAA
CTX2_SLYC 3129 bp        3074 -CAGA----AAT------TGAATCTGCAGAGTGCAAAGAAATTGCATGGCAAATCTTTATTCAGTAA
CTX1_SLYC 3159 bp        3099 ACGAGCGCAGGA------CGATAGAATCGAATGCAATGCCAACGCTTGGCAAATCTTTATCAAATAG
CTX_EPAR 3150 bp         3093 GCGACAAAGGAA------TAGAAGTCTGAATTGTAAAGCGAGCAGTTGGCAGATTTATATTCAATAG
CTX2_SPHAR 3129 bp       3074 -CAGA----AAT------TGAATCTGCAGAGTGCAAAGATATTGCATGGCAAATCTTTATTCAGTAA
CTX1_SPHAR 3159 bp       3099 TCGAGCGCTGGA------CGATAGAATCGAATGCAATGCCAACGCTTGGCAAATCTTTATCAAATAG
CTX_WSCI_PARTIAL 1653 bp 1641 ------------------------------------------------------TTGTTGTCAAGAG
CTX_IHAL 3156 bp         3096 GCGTTCACAAGA------CAGTTCGATAGATTGCAAAGCAAACGCATGGCAGATTTATGTGAAATAA
CTX2_XNOT 3183 bp        3128 -AGAA----AAT------GGACTCTTTTGAGTGCAAGTCAATTGCGTGGCAATTGTTTATTCAATGA
CTX1_XNOT2 3150 bp       3090 GAGAGTACACGA------CAAAAGAGTTGATTGTAAAGCAAATGCATGGCAGATTTACGTGAAATAA
CTX_OROB_PARTIAL 1431 bp 1430 -----------------------------------------------------------------CA
CTX_SAFF 3147 bp         3087 ACGACAACGGAA------TAGAAATATGAATTGTAAAGCGAGCAGTTGGCAGATTCATATCAAATAA
CTX_SOBS 3156bp          3096 ACGAGAAAGGAA------TAGAAGTATGACTTGTAAAGCGAGCAGTTGGCAGATTTATATCAAATAG
CTX_Sarc 2907 bp         2841 ACGAGAACAAAAGATTGCCAGTCATCTTAATTGTAATGCTAACGCATGGCAAATCTTTATACAATAG
CTX2_Dopa 3123 bp        3068 -AGGA----ACT------TGCCTCTTCAAAGTGTAAAGATATTGCTTGGCAAATATTCGTTCAATAA
CTX1_Dopa 3123 bp        3102 ACGAGTCTCGGA------CAATAATATCAATTGTAGTGCTAATGCTTGGCAAATCTTTATAAAGTAG


```

**B**

```
                            1 ........10........20........30........40........50........60........70........80
CTX1_AESCU 3159 bp          1 M--M-GTSRCVILLFALLLWAANAAPPEIHTTRP--------N------VPEEIKRPNSTEIE-TPAVKQL-----ETPS
CTX2_AESCU 3129 bp          1 M--V--LWQ-VLFLVPLLWQCVQGFSPDDNATS-----------------GDHPVAMDETKKDNQTENQTP-----QFPT
CTX1_DPEA 3156 bp           1 M--S-GTWWHVPFLFPLLLLGANGAPSEITT-KF--------P------LLEENIHFNSTEPKPTAALKDL-----DPPL
CTX2_DPEA 3123 bp           1 M--V--LWQ-VLFLFPLLWLTIRCFPLEDNAAV----------------VCDQAQVMNQTGCK-PSESNTP-----DFPM
CTX1_ABAN 2946bp            1 M--M-AEWRYTVFLFPLLLLAANAAPPESHTTRP--------K------VPE-EYQTNSTEII-TPAVKQL-----ETPT
CTX2_ABAN 2286 bp           1 M--F--LWQ-VLFLVPLLWQHAQGILPDDGATS-----------------HDQLVAMNETKKDDQTKNQTP-----QIPT
CTX_EBER 3159 bp            1 M--MGSSWPRVLLLFVLPLLGVSGANVMVNNTEL--------D------IIQMMNTTSSTEVE-TPLVKQF-----ETPS
CTX2_SOFF 3129 bp           1 M--V--LWQ-RLFFVPLLWQCVQGVLPDDGATS-----------------RDHPVAMNETKKEEVTEKQTP-----QFPT
CTX1_SOFF 3141bp            1 M--K-EAWRYVILFLAVLLLAANGAPLESTTSE--------------------PEETNSTVIA-TPAVKII-----ETPS
CTX_ESCO 3159 bp            1 M--MGSSWPWVLFLFALPLLGISGATVMVNNTEL--------D------IIQMMNTTSSTEVE-TPLVKQF-----ETPS
CTX2_SLES 3132 bp           1 M--V--LWQ-VWFLFSLLWLAVRGFPPEENTAI----------------VCNQLEAMNQTGCK-SPEHNNS-----NLPM
CTX1_SLES 3159 bp           1 M--I-EAKWHVSLLLLLLLLGANEASSDIIATKF--------P------FLEKNTHLNSTETKPIPKVKDL-----EPPL
CTX_TRHO 3150 bp            1 M--V--LWQ-ALFLLPMLWLSVHGSSSDDSTAAG---------------VSSQMVAMNETEKVPGTESDRPANRSSELPS
CTX_SATL 3147 bp            1 M--G-SSWPRVLLFFALLSLGIGGATVAVDNSQV--------D------ILGMINTTGPTEVE-TPVVKQM-----ETPS
CTX2_ALYC 3129 bp           1 M--V--LWQ-VLFLVPLLCQCVQGFSPDDIAIS-----------------RDHPVAIDETKKDNQTETQTP-----QFPT
CTX1_ALYC 3159 bp           1 M--M-GTSLCVILLFALLLWAANAAPPESHTTRP--------K------VPEEIKRPNSTEIV-TPAVKTL-----ETPT
CTX_EPAR 3150 bp            1 M--MGSSWPRVLLLCALPLLVVSGATVL--DTEL--------D------IFQLMNTTSSTEVE-TPLVKQF-----ETPS
CTX2_APHAR 3129 bp          1 M--V--QWQ-VLFLVPLLWQCVQGFSPDDITTS-----------------RDHPVAMDETKKDNQTETQTP-----QFPT
CTX1_APHAR 3159 bp          1 M--M-GTSRCVILLFTLLLWAANAAPPESHTTRP--------K------IPEEIKRPNSTEIV-TPAVKQL-----ETPT
CTX_WSCI_PARTIAL 1653 bp    1 --------------------------------------------------------------------------------
CTX_IHAL 3156 bp            1 M--A-TEWQLIFFTVALLIQGNNGAPTDTNVSQV--------I------VPNVPMDNNSTEIK-TPVVKEM-----ESPT
CTX2_XNOT 3183 bp           1 M--T--LWQ-ALFLHLFLLICATVVSSQDCDAPNSYTSDIDPSINHSDHQLNGTDVGNTTLCK-ETEIKSS-----GLPT
CTX1_XNOT2 3150 bp          1 M--V-AEWQLIFVTVALVFQGTNGAPENTNVSQI--------V------VPNVPTEFNSTEIK-TPVVKEM-----ESPT
CTX_OROB_PARTIAL 1431 bp    1 --------------------------------------------------------------------------------
CTX_SAFF 3147 bp            1 M--G-SSWPRVLLFFALLSLGIGGATVAVDNSQV--------D------ILGMINTTGPTEVE-TPIVKQM-----ETPS
CTX_SOBS 3156bp             1 M--G-SSWPRLLLLFVLLSLGISGETAEIDNSQV--------H------IFGMTNTTGPTEVE-TPTVKQL-----ETPS
CTX_Sarc 2907 bp            1 --------------------------------------------------------------------------------
CTX2_Dopa 3123 bp           1 M--V--LWQ-VLFVFPLLWLTIRCFPLEDNAAV----------------VYDQAQVMNQTGYK-PPERNTP-----EFPM
CTX1_Dopa 3123 bp           1 MKMR-GTWWHVPLLFPLLLLGANGAPSEITT-KF--------P------LLEENIYLNSTEPKPTAALKDL-----EPPL

                           81 ........90.......100.......110.......120.......130.......140.......150.......160
CTX1_AESCU 3159 bp         58 IFLLTTLEVAEADVDSTLETMKDR--NKKNSAKLSKIGNNMKSLLSVFSVFGGFLSLLSVVTTTSDLQVISDMFTGVNRK
CTX2_AESCU 3129 bp         54 VMIMTTLEVVKSEIEDVLNYEGEA--GNKFFSIKKKHANSLKIFNSLFNALGGFLTVLSAFTQTSDLEVITAMFKEVNKK
CTX1_DPEA 3156 bp          58 IFLQTSLEVAEADLDRNIETIKEK--SKNPAAKVSKFGHGVKKICSMFNVFGGFFSLLATVTSSSDLKVITGMFDEVNKK
CTX2_DPEA 3123 bp          54 TMIMTTLEMVQTEIGELQNFETEQ--LTNDFG---KKSKTLKVFSSVFNALGGFLTLLSVFTGTDDLEVISSMFKQVNKK
CTX1_ABAN 2946bp           57 IFLLTTMEVIEADVDTMLEAMKDK--NKKKAAKLTKFGNDMKSLLGVFSVMSGFLSFLSVITTTSDLQVISDMFTGVNRK
CTX2_ABAN 2286 bp          54 AMIMTTLEVVKSEIDDALNYEGEA--GTKIFRITSKHAKTLKIFNSFLNAFGGYLSLLSAFTQTSDLEAITSMFKEVNKK
CTX_EBER 3159 bp           59 IFLLTTLEEVDADFDKRLETIKDK--NKKPPAKMSSIGNSMKKFCSMFSVMSGFLSLLSVVSTTSDLQAITDMFDGVNKK
CTX2_SOFF 3129 bp          54 AMIMTTLEVVRSEIEDVLNYEGEA--VNNFLSISKKHAKTLKIFNSLFTAFGGFLSVLSTFTQTSDLEVITGMFKEVNKK
CTX1_SOFF 3141bp           52 IFLLTTLEVAEAEVDKSLESMKSR--NKKNAAQFDKIGKNVKSLLSVFSVFSGFLSLLSVVTTTSDLQVISDMFTGVNKK
CTX_ESCO 3159 bp           59 IFLLTTLEEVNTDFDKRLETIKDK--NKKSPAKISSIGNSMKKFCSMFSVMSGFLSLLSVVSTTSDLQAITDMFDGVNKK
CTX2_SLES 3132 bp          54 VMIMTTLEVVQSEIGELQNEETEK--ESNSLSKKTKKSKILKIFSSLFNAFGGFLNLLSVFTGTDDLEVISSMFKEVNKR
CTX1_SLES 3159 bp          59 IFLQTGLEVAEADLDKNLETIKEK--KKNPIAKISRFGNGVRLFCSIFSITSGFFSLLATVTSSSDLQVISDMFTEVNKK
CTX_TRHO 3150 bp           61 IIIMTTLEVMQSEIEKTQNDGTEK--RINGLSKGIKHAKILKVFCSLFNAFGGFLTVLSALTGTSDLEVISSMFEEVNKK
CTX_SATL 3147 bp           58 IFLLTTLEEVDTDFDKRLETIKDK--NKKTPAKMTSFGNSMKKFCSMFSVMSGFLSLLSVVSTTSDLQAITDMFDGVNKK
CTX2_ALYC 3129 bp          54 LMIMTTLEVVKSEIEDFLNYEGEA--GNSFFSVKKKHAKTLKIFNSLFNALGGFLTVLSAFTQTSDLEVITAMFKEVNKK
CTX1_ALYC 3159 bp          58 IFLLTTLEVTEADVDSTLEAMKDK--NKKNSAKLSKIGNNMKSILSVFSVMSGFLSLLSVITTTSDLQVISDMFTGVNRK
CTX_EPAR 3150 bp           57 IFLLTALEEVNTDFDKRLDTIKDK--NKKPPAKMSSIGNTMKKFCSMFSVMSGFLSLLSVVTTTSDLQAITDMFDGVNKK
CTX2_APHAR 3129 bp         54 VMIMTTLEVVKSEIEDVLNYEGEA--GNKFFSVKKKHAKTLKIFNSLFNALGGFLTVLSAFTQTSDLEVITAMFKEVNKK
CTX1_APHAR 3159 bp         58 IFLLTTLEVAEADVDSTLESMKEK--NKKNSAKLSKLGNNMKSLLSVFSVMSGFLSLLSVITTTSDLQVISDMFTGVNRK
CTX_WSCI_PARTIAL 1653 bp    1 --------------------------------------------------------------------------------
CTX_IHAL 3156 bp           58 IFLMTSLEVAESDFEKKMDLASNK--VKKNPMKLTKIGNSLRRFTSMFSMLGGFLSMLAVVTTTTDLQVISDMFGEVNKK
CTX2_XNOT 3183 bp          70 VVIITSLEVIQVEIDSYQDEDNNKDSKKNKPKRTPILSKSVKMFSSFLNSMGGFLSLLSVFTDTSDLEVISGMFKEVHKK
CTX1_XNOT2 3150 bp         58 IFLMTSLEVAESDFEKKMDSASNK--VKKNPVKLTKFGNSLRRFTSMFSMLGGFLSMLAVMTTTTDLQVISDMFGEVNKK
CTX_OROB_PARTIAL 1431 bp    1 --------------------------------------------------------------------------------
CTX_SAFF 3147 bp           58 IFLLTTLEEVDTDFDKRLETIKDK--NKKTPAKMTSFGNSMKKFCSMFSVMSGFLSLLSVVSTTSDLQAITDMFDGVNKK
CTX_SOBS 3156bp            58 IFLLTSLEEIDTDFDKRLETLKDK--TKKTSSKMTSLGNGLKKFCSMFSVMSGFLSLLSVVSTTSDLQAITDMFHGVNKK
CTX_Sarc 2907 bp            1 --------------------------------------------------------------------------------
CTX2_Dopa 3123 bp          54 TMIMTTLEMVQTEIGELQNFETEQ--LTNDFG---KKSKTLTVFSSVFNALGGFLTLLSVFTETDDLEVISSMFKQVNKK
CTX1_Dopa 3123 bp          60 IFLQTSLEVTEADLDKNIEAIKEK--SKNPAAKVSKFGHGVKKICSMFSVFGGFFSLLAAVTSSSDLTVITGMFDKVNKK

                          161 .......170.......180.......190.......200.......210.......220.......230.......240
CTX1_AESCU 3159 bp        136 LDQINDKLDKLDNSVELQGLLTNYIPWQYSVKNGIEKLIETYKKMVEETNMNKRRLMAENFILFFENNQIESNINNLIKL
CTX2_AESCU 3129 bp        132 LDRITRRIDNLENSIELQRLLSNYIPWHFSVINGMEKLTETYTSMAQEADIKKRRIQAERFIKFFEDNNIESHINNLIRI
CTX1_DPEA 3156 bp         136 LNKITDQLKKLDNSVQLEGLLSNYIPWQYSVTNGFEKLVETYTAMVKETDINKRSLMSENYILYFENNQIESNINNLIKL
CTX2_DPEA 3123 bp         129 LDKISQQIDNVENTVELQGLLSNYIPWHFSVINGIEKLSETFSLMAIEADIRKRRIQAEDFIKFFEDNQIESHVNNLIQV
CTX1_ABAN 2946bp          135 LDQINDKLDKLDNSVELHSLLSNFIPWQYSVKNGIEKLVETYSKMVKETNINKRRLLAENFISFFENNQIEANINNLIKL
CTX2_ABAN 2286 bp         132 LDRITQRIDNLGNTVKLQGLLSNYIPWHFSVINGIEKLTETYTSMAKEADIRKRRIQAENFIKFFEDNNIESHINNLIRI
CTX_EBER 3159 bp          137 LDDISDRLGRLDDKVELQTLLSNYMPWQYSVSNGIEKLVETYSAMSKVSEINRRRIIAENFILFFENNQIEANINNLVRI
CTX2_SOFF 3129 bp         132 LDRINRRIDSLGNTVELQRHLSHYIPWHFSVINGMEKLTETYTSMAKEADIRKRRIQAESFIKYFEDNNIESNVNNLIRL
CTX1_SOFF 3141bp          130 LDQINDKLDKLDNSVELQGLLSNYIPWQYSVKNGIEKLIETYKKMVRETNINKRRLIAGNFILFFENNQIESNVNNLVKL
CTX_ESCO 3159 bp          137 LDDISDRLGRLDDKVELQALLSNYMPWQYSVSNGIEKLIETYSAMSKVSEINRRRIIAENFILFFENNQIEANINNLMRI
CTX2_SLES 3132 bp         132 LDKIALQIDNVENTVELQGLLSNYIPWYFSIINGIEKLAETYSVMAKEPDIRKRRNQAEDFIKFFEDNQIESHFDNLMKL
CTX1_SLES 3159 bp         137 LDKITDRLDKLDNSIELHGLLSNYIPWQYSVTNGIEKLVETYTAMVKEPDFNKRRLIAENFILYFENNQIELNINNLIKL
CTX_TRHO 3150 bp          139 LDNIARRIDNVESAVEMQGLLSNYIPWHFSVINGIERLTETYSAMAKEPEIRKRRIQAEHFIKFFEDNQIESNVNNLIRL
CTX_SATL 3147 bp          136 LDDISDRLGRLDDSVELQGLLSSYIPWQYSVTNGVEKLTETYSAMSKVSEINRRRIIAENFILFFENNHIEANINNLVRI
CTX2_ALYC 3129 bp         132 LDRITRRIDNLENSVELQRLLSNYIPWHFSVINGMEKLTESYTSMAQEADVRKRRIQAERFIKFFEDNNIESHINNLIRI
CTX1_ALYC 3159 bp         136 LDQINDKLDKLDNSVELQGLLSNYIPWQYAVKNGIEKLIETYKTMVEETDINKRRLIAENFILFFENNQIESNINNLIKL
CTX_EPAR 3150 bp          135 LDDISDRLGRLDDKVELQTLLSNYMPWQYSVSNGIEKLIETYSAMSKVSEINRRRIIAENFILFFENNQIEANINNLMRI
CTX2_APHAR 3129 bp        132 LDRITRRIDNLENSIELQRLLSNYIPWHFSVINGMEKLTETYTSMAQEADVRKRRIQAERFIKFFEENNIESHINNLIRI
CTX1_APHAR 3159 bp        136 LDQINDKLDKLDNSVELQGLLSNYIPWQYSVKNGIEKLIETYKAMVEETDINKRRLIAENFILFFENNQIESNINNLIKL
CTX_WSCI_PARTIAL 1653 bp    1 --------------------------------------------------------------------------------
CTX_IHAL 3156 bp          136 LDKINDRLVKLDNSVELQSLLSNYIPWEYSITNGIHKLVETYTSMAKESNMHQRRAIAGNFINFFENNQIESNINNLIYI
CTX2_XNOT 3183 bp         150 LDKIYNRIDNLKGSIELQGLLSNYIPWYFTVINGVEMLTETYMSMAKEQDLRKRRILVENYISYFESNQVGSNVNNLLRM
CTX1_XNOT2 3150 bp        136 LDKINDQLIKLDDSVALQSLLSNYIPWEYSITNGINKLIETYTAMAKESNMHERRVIAGDFINFFENNQIESNINNLIYI
CTX_OROB_PARTIAL 1431 bp    1 -------------------LLSNYIPWQYSVKHGMEKLVETYTAMAKEGDINKRRLLAEHFIIFFENNQIESNIYNLIKL
CTX_SAFF 3147 bp          136 LDDISDRLGRLDDSVELQGLLSSYIPWQYSVTNGVEKLTETYSAMSKVSEINRRRILAENFILFFENNQIEAHINNLVRI
CTX_SOBS 3156bp           136 LDDISDRLGRLDDSVELQGLLSSYIPWQYSVANGVEKLIETYSAMSKVSEINRRRILAENFILFFENNQIEANVNNLMRI
CTX_Sarc 2907 bp            1 --------------------------------------------MAKEEDINKRRIIAEGFIIFFESNQIESNINNLIKL
CTX2_Dopa 3123 bp         129 LDKISQQIDNVENTVELQGLLSNYIPWHFSVINGIEKLSETFSHMAIEEDIRKRRIQAEDFIKFFEDNQIESHVNNLIQL
CTX1_Dopa 3123 bp         138 LDKITHQLEKLDNSVQLQGLLSNYIPWQYSVTNGFEKLVETYTAMVKETDINKRSLMSENFILYFENNQIESNINNLIKL

                          241 .......250.......260.......270.......280.......290.......300.......310.......320
CTX1_AESCU 3159 bp        216 TTTTDAV-HQNMLFNELLDE-AGCDIIRLTRIYMHVRRIFYQGTQLVLAYNSFKQMDPPEMKKYLNALIFIRNMYQSRVW
CTX2_AESCU 3129 bp        212 TTTSDSA-FYKNIYRLLINK-AGCNLPKLKVIYERVTQIVTSGSQLILAYRSFMQIEIPKLKKYLDALLLFRQIYEKSIW
CTX1_DPEA 3156 bp         216 TLTTDAV-HQNQLFDKLIDE-AGCDIVRLTRLHFQIQQIFTQGCQLVLAYNSFKRMESPSIQKYIDALIYIRNIYESRVW
CTX2_DPEA 3123 bp         209 TTKSDSV-FYENIYQSLFNK-ANCKMNKLNLIHEKVTGIITSGSQLVLAYRSLKQLEKPKMTKYLNALFSLRKMFDAQTW
CTX1_ABAN 2946bp          215 TTTADAV-HRNVLFNELIDE-AGCDYIRLTRIYMHIRRIFFQGCQLVLSYHSFKQMDPPETKKYINALVFIRNMYQSRVW
CTX2_ABAN 2286 bp         212 TTTSDSA-FYENVYRLLINK-AGCNLLKLKVIYERVTQIVTSGSQLILAYRSFMHMQVPKLKKYFDSLFIFRQIYEKSIW
CTX_EBER 3159 bp          217 TTTSDAI-QQTNLFIELIEE-SKCNVIRLSNVYLQVKRIFMQGSQLVQAYHAFKQMEMPNIQKYLDALVYIRTAYESRLW
CTX2_SOFF 3129 bp         212 TTTSDSA-FYKNIYRLLLNK-AGCNLLRLKVIYERVSQIVTSGSQLILAYRSFMQMQIPKAKKYFDVLFLFRQIYDKRIW
CTX1_SOFF 3141bp          210 TTTTDAV-HQNMLFEELIDE-AGCDIVRLTRIYMHVRRIFYQGCQLVLAYNSFKQMEPPEIKKYLNALIFIRNMYQARIW
CTX_ESCO 3159 bp          217 TTTSDSI-QQTNLFQELIKE-SECDVIRLSNVYLHVKRIFMQGSQLVQAYHAFKQMEMPNIQKYLDALLYIRTAYESRLW
CTX2_SLES 3132 bp         212 TTVSDSV-FYKNTFQLLLKK-AGCKMPKLNLIYERVTRILTSGSQLVLAYRSFKQLEKPKMTKYMNALFLLRQIFEAHDW
CTX1_SLES 3159 bp         217 TVTSNII-QQNQLFNELIDE-AGCNIIRLTRLHIYIKRIFTQGGLLIVAYNLFKQMEPPNIQKFLNALIYIRKIYESRVW
CTX_TRHO 3150 bp          219 TTVSDSV-FYKNIYSSLINK-AGCNMQKLNSIYERVTRIITSGSQLILAYRSFKQMDKPKLKKYLNALFSLRQIFEARVW
CTX_SATL 3147 bp          216 TTTSDTI-QQTNLFKELIEK-SDCDIIRLTNVYLHVKRIFMQGSQLVQAYHAFKQMEMPNIQKYLDALVYIRTVYESRVW
CTX2_ALYC 3129 bp         212 TTTSDSA-FYKNIYILLINK-AGCNLPKLKVIYERVTQIVTSGAQLILAYRSFMQIQIPKLKKYLDALFLFRQIYEKRIW
CTX1_ALYC 3159 bp         216 TTTTDAV-HQNMLFNELIDE-AGCDIVRLTRIYMHVRRIFYQGTQLVLAYNSFKQMDPPEIKKYLNALIFIRNMYQSRVW
CTX_EPAR 3150 bp          215 TTTSDTI-QQTNLFQELIEK-SECDVIRLSNVYLHVKRTFMQGSQLVQAYHTFKQMEMPNIQKYLDALVYIRTVYESRLW
CTX2_APHAR 3129 bp        212 TTTSDSA-FYKNIYRLLINK-AGCNLPKLKVIYERVTQIVTSGSQLILAYRSFMQIQMPKLKTYLDALFLFRQIYEKRIW
CTX1_APHAR 3159 bp        216 TTTTDAV-HQNMLFNELIDE-AGCDFVRLTRIYMHVRRIFYQGTQLVLAYNSFKQMDPPEIKKYLNALIFIRNMYQSRVW
CTX_WSCI_PARTIAL 1653 bp    1 -----------------------------------------------------KQMEPPRIEQFTEALLVIRKLYGNRIW
CTX_IHAL 3156 bp          216 TTTSSAVLHKKKLFDELIDK-AGCNFVRLTGIYTHMKRIFTQGSQLILAYYTFKDGKVPLVQTYIDALTSIRNTYDYRIW
CTX2_XNOT 3183 bp         230 TTTTDSG-IHKNIYKLMLKK-AGCNLSKLKEVHEKVARIIMSGSELVLAYRIFKRMEKPNQKKYINALLFYRRLYENQVW
CTX1_XNOT2 3150 bp        216 TITSSASLGKKKLFDELIDE-AGCNFIRLTGIYTHMRRIFTQGSQLVLAYYSFKDGKVPLVQTYIDALTSIRNTYDYRIW
CTX_OROB_PARTIAL 1431 bp   62 TTTSDIV-QQKNLFNELIEE-AGCDAIRLTQVYLHIKRIFTEGCQLILAYRSFKQMEAPRIEGFLDAVAFIRKMYDKRVW
CTX_SAFF 3147 bp          216 TTTSDTI-QQTNLFKELIEK-SDCDIIRLTNVYLHVKRIFMQGSQLVQAYHAFKQMEMPNIQKYLDALVYIRTVYESRVW
CTX_SOBS 3156bp           216 TTTSDTI-QQTNLFKELIEK-SECNIIRLTNVYLHMKRIFMQGSQLVQAYHAFKQMEMPSIQKYLDALVYIRTLYESRVW
CTX_Sarc 2907 bp           37 TTESDTV-QKKNCLTN*LTKRTATSSG-------*HKFICISNEYLPKVLN*F*PITP*SIWN-LRAFRSSLRLYY-SLG
CTX2_Dopa 3123 bp         209 TTKSDSV-FHENIYQSLFNK-ANCKMNELNLIHEKVTGIVTSGSQLVLAYRSFKQLEKPKLTKYLNALFSFRKMFDAQTW
CTX1_Dopa 3123 bp         218 TLTTDTI-HQNQLFNKLIDE-AGCDIVRLTRLHFQIQQLFTKGCQLVLAYNSFKQMESPSIQKYIDALIKIRNIYESRVW

                          321 .......330.......340.......350.......360.......370.......380.......390.......400
CTX1_AESCU 3159 bp        294 HCKETTIAQSKKD---IKDI--V-KTNAK--FGITTVLRKINSELSRKYPWYSWSIVTVK-KMLANQRNSTLGNQFYEME
CTX2_AESCU 3129 bp        290 HCKETAIDRSKKT---IYRM--L-IGKKS-----RVPLKRIASMLSRNFPWYSWSLGLTR-KWIGSRENVVSGNQYYELK
CTX1_DPEA 3156 bp         294 YCKETTIVQSKKD---VVKI--V-TANSH--LSITPLLKNINDDLSRNYPWYSWSIVNIR-RMLRQEKNSPIGNQYYELE
CTX2_DPEA 3123 bp         287 YCKETAIDRSKKA---IFQM--I-NGTKK-----RFPLKRIGKMLSINYPWYSWSLGLTKPKSPHFRDNFYSGNQFYELK
CTX1_ABAN 2946bp          293 HCKENTIAHSKKD---IKDV--V-KKDAK--LGITTVIKNINSELSKKYPWYSWSIVTVR-KMLSDQRNSTIGNQYYEME
CTX2_ABAN 2286 bp         290 HCKETAISRSKNA---VYRM--L-YGKKS-----RVSLKVIANMLSYNYPWYSWSLGLTR-KRIGSRENVVSGNQYYELK
CTX_EBER 3159 bp          295 RCKETAILRSKNT---IVEI--V-KRDPH--LGIIKLLVNIKEELSKTYPWYSWSVVNLQ-RMIGAEVNATTGNQFYEME
CTX2_SOFF 3129 bp         290 RCKETAIDRSKRA---VYRM--L-IGEKS-----RSSLKRIANMLSSNYPWYSWSLGLTT-KWTDSRENVVSGNQYYELK
CTX1_SOFF 3141bp          288 HCKETTIANSKKD---IKDI--A-KTHAK--FGISTLLTNINKELSKKYPWYSWSIVTIR-KQLASQRNSTSGNQYYEML
CTX_ESCO 3159 bp          295 RCKETSILRSKNS---IIEI--A-KRDSH--LGIIKLLVNIKKELSKTYPWYSWSVVNLQ-RMIGAEVNATSGNQFYEME
CTX2_SLES 3132 bp         290 HCKETAIARSKKA---VLQM--I-SGNKK-----RVGLKSIAKMLSINYPWYSWSLGLTKKRSLHSRQNFVSGNQFYELK
CTX1_SLES 3159 bp         295 HCKETTIIRSKND---VVKI--I-TKNPH--LEITTLLKKIIDNLSRNYPWYSWSIVNVR-RMLSEERNSTLGNQYYELE
CTX_TRHO 3150 bp          297 YCKETAIPRSKRN---ILRM--L-GGRKS-----RPALKRIGNVLSKNYPWYTWSLGLTR-NQLNFNRESASGNQFYELK
CTX_SATL 3147 bp          294 RCKETSIVRSKND---IIKI--A-KRDSN--VGIIKLLGNINRDLSKTHPWYSWSIVNLR-RMIGAEANATSGNQFYEME
CTX2_ALYC 3129 bp         290 HCKETAIDRSKKT---IYRM--L-NGKKS-----RVPLKRIASMLSRNFPWYSWSLGLTR-KWIGSRENIVSGNQYYELK
CTX1_ALYC 3159 bp         294 HCKETTIAHSKKD---ITDI--V-KTNAK--FGITTVLRKINSELSRKYPWYSWSIVTVK-KLLANQRNSTLGNQFYEME
CTX_EPAR 3150 bp          293 RCKETSILRSKNE---IMKI--V-KRDSH--LGIIKLLVNIKKELSKTHPWYSWSVVNLR-RMIGAEVNATSGNQFYEME
CTX2_APHAR 3129 bp        290 HCKETAIDRSKKT---IYRM--L-NGKKS-----RVPLKRIASMLSRNFPWYSWSLGLTR-KWIGSRENVVSGNQYYELK
CTX1_APHAR 3159 bp        294 HCKETTIAHSKKD---ITDI--V-KANAK--FGITTVLTKINSELSRKYPWYSWSIVTVK-KLLASQRNSTLGNQFYEME
CTX_WSCI_PARTIAL 1653 bp   28 HCKETTIARSKTD---IEEI--V-KNNTG--NGITSSLEKINAELSKKYPWYSWSVVNVR-RLLEAEQDSVIGNQYYQME
CTX_IHAL 3156 bp          295 HCKETTITRSKSD---VDKL--V-KKNSNEGRGATVTVKNVNKALSKKYPWYSWSIINNK-RWVGSKKSATSGNQYYEML
CTX2_XNOT 3183 bp         308 YCKEMAIIRSKSA---VKKM--L-SGVKA-----IPSLKRIAKMLSANFPWYSWSLGITK-KWNSARHNYVSGNQFYEME
CTX1_XNOT2 3150 bp        295 HCKETTITRSKSD---VDKV--V-TKSSEEERGATVTLNNVNKALSQKYPWYSWSIVNIK-RLTSS--TSASGNQYYEMI
CTX_OROB_PARTIAL 1431 bp  140 HCKETTIVRSKDD---IAKI--A-KKYSN--FGIPAQLKNINVALSRKYPWYSWSIVNVG-GLLE--ADSAMGNQYYEMT
CTX_SAFF 3147 bp          294 RCKETSIVRSKND---IIKI--A-KRDSN--VGIIKLLGNINRDLSKTHPWYSWSIVNLR-RMIGAEANATSGNQFYEME
CTX_SOBS 3156bp           294 RCKETSITRSKND---ITKI--A-KRDSR--LGIIKSLGNINKELSKTHPWYSWSIVNLR-RLIGAEVDSTSGNQYYEIQ
CTX_Sarc 2907 bp          102 KCTGTALGTVKKTRSPSPRMTL*KKNDSD--LGITTLLKNVNGALSKKYPWYSWSIVNVR-RLLDAKSGSTMGNQYYQME
CTX2_Dopa 3123 bp         287 YCKETAIDRSKKA---VFQM--I-NGTKK-----RFPLKRIGKMLSINYPWYSWSLGLTKQKSPHFRDNFYSGNQFYELK
CTX1_Dopa 3123 bp         296 YCKETTIVNSKKD---VVKI--V-TTNSH--LSITPLLKNINYGLSRKYPWYSWSIVNVR-KLLRNEKNSPTGNQYYELE

                          401 .......410.......420.......430.......440.......450.......460.......470.......480
CTX1_AESCU 3159 bp        365 AVGPHGSNFVVIWQGFKEHSQCEDIQKANTVAVLTICKSCHQSHVFTPSNMLNKNTCPNNQYPQVKAFIDRREPFRDEIQ
CTX2_AESCU 3129 bp        358 DLWPLGIYLVVIWQGVTETSQCNQMPKANTVVFLDICKRCPKTYMYSPKNSLSSIKCPDEKYPRLKKFIDKRFPDGTES-
CTX1_DPEA 3156 bp         365 KVGSNGVNLVVTWQGSNEKSQCREIGKANTFLFLSICKSCERSHVLISENMQSKNKCPKENYPQVKAFIDQRGPEVDKNE
CTX2_DPEA 3123 bp         356 GLWPLETSLLITWQGVNETSRCNRMLKANTIIFLDMCKRCLKTYLYAPENMLTSIKCPKERNPDIKNYIDKTFPGGTEI-
CTX1_ABAN 2946bp          364 KVGPYGSNFVVTWQGSKENSQCEDIYKANTVVVLTICKSCHNSHVFTPSNMLDKNKCPKDRYPQVKALIDQHEPQRNERE
CTX2_ABAN 2286 bp         358 DLWPFGIHLMVIWQGVTETSHCNQMPNANTVVFLDLCKKCPKTYMYSSKNMLSSIKCPKEKYPNLKKFIDKQFPDRTES-
CTX_EBER 3159 bp          366 KVGSYGWNLAVTWQGTDEKSQCQDIGKAQTVVFLSICKRCHTSRVAISGDMLSMNKCPAKTYPEVKAFINQRGSNLVKNE
CTX2_SOFF 3129 bp         358 DLWPFRTYLVVIWQGVTETSQCNRMPKANTVVFFDICKRCTETYIYSPKNTLSSIKCPSERYPKLRKFINKRFPDGTES-
CTX1_SOFF 3141bp          359 SVGEYGSNLVVTWQGSKEEAQCEDIQKANTVVVVSICKSCHNSHVFTPSNMLDKNSCPQNNYPQVKAIINPREPQFEESD
CTX_ESCO 3159 bp          366 KVGSYGWNLAVTWQGTDEKSQCQDIGKAKTVVFLSICKRCHTSRVAISGDMLSKNKCPAKTYPEVKAFINQRGSNLVKNE
CTX2_SLES 3132 bp         359 GLWPFEMTLIITWQGVNEISRCNHMLKANTIIFLDICKRCVKTYLYVPQNMLSSMKCPQERYPIIKDFIDKSFPDGTES-
CTX1_SLES 3159 bp         366 KIGPKGVNLVVTWQSSNEKSSCQAIGKVNTFVFLSICPSCQDSHVFISDNMLSKNKCPKENYPQVKDFIDQRGPELDKNG
CTX_TRHO 3150 bp          365 RLWPLGISLVITWQGVNEVSRCNQMLKANTIVFIDICKSCQKSYLYASRNMLASIKCPKERYPKIKKFIDKKFPNDTRS-
CTX_SATL 3147 bp          365 KVGSYGWNLAVAWQGTDEKSQCQDIGKANTVVFLSICTACHTSRVSISKDMLSKNKCPAETYPEVKAFINQRGSHLVKNE
CTX2_ALYC 3129 bp         358 DLWPLGIYLVVIWQGVTETSQCSHMPKANTVVFLDMCKRCPQTYMYSPKKSLSSIKCPNEKYPKLKKFIDKRFPDGTES-
CTX1_ALYC 3159 bp         365 AVGPHGSNLVVIWQGSKENSQCEDIQKANTVTVLTICKSCHQSHVFTPSNMLNKNKCPNDQYPQVKAFIDRREPMRRERE
CTX_EPAR 3150 bp          364 KVGSYGWNLAVTWQGTDEKSQCQDIGKANTVLFLSICKACHTSRVAISKDMLSKNKCPAETYPEVKAFINQRGSHLVKNE
CTX2_APHAR 3129 bp        358 DLWPLGIYLVVIWQDVTQTSQCNQMPKANTVVFLDMCKRCPQTYMYSPKKSLSSIKCPNEKYPKLKKFIDKRFPDGTES-
CTX1_APHAR 3159 bp        365 AVGPHGSNLVVIWQGSKENSQCEDIQKANTVAVLTICKSCHQSHVFTPNNMLNKNKCRNDQYPQVKAFIDRREPMREEIE
CTX_WSCI_PARTIAL 1653 bp   99 KVGKHGVNLVVAWQGSDEKPQCSDVREANTFLFLSICKSCKNSHTFVPKNMISKTKCPKKRYPEVKALIDERGPHLDENG
CTX_IHAL 3156 bp          368 KVGKYGMNIVVAWQGVDEKAECYDIDRSKTFLFIGICRGCQNSHVSINEDMLSKNKCPKNRYPRVKAFIDRQGPELDRNS
CTX2_XNOT 3183 bp         376 GYLPLGLSVIVVWQGVNEKSNCKLMPKANTIVLIETCKRCKNSYLYGSKTMLTSSKCPTNTFPNIKSLIDARTPHGTRS-
CTX1_XNOT2 3150 bp        366 KVGKYGMNLVVAWQSADEKPECYDIDQSKTFLFLSICKRCQNSHVSINENMLNKNKCPKNLYPRVKAYINREGPELDKDG
CTX_OROB_PARTIAL 1431 bp  209 KVGSHGMNLVVSWQGSDENPQCQDIRKANTLVFLSICKSCQNSRTYIPSNMLSNNKCPPKRNPQVKAFIDQKGSHLYRNM
CTX_SAFF 3147 bp          365 KVGSYGWNLAVAWQGTDEKSQCQDIGKANTVVFLSICTACHTSRVSISKDMLSKNKCPAETYPEVKAFINQRGSHLVKNE
CTX_SOBS 3156bp           365 NVGSHGWNLAVTWQGTDEKSQCQDIGKANTVVFLSICKACHTSRVSIAKDMLSKNKCPAGTYPEVKAFINQRGSHLVKNE
CTX_Sarc 2907 bp          178 KVGYYGWNLIVTWQGSAEKPQCHNIREANTFVFLSIGKSCQISRTYILKNMLSKNNCPKNVNHQVKALIDERGPHPDKNR
CTX2_Dopa 3123 bp         356 GLWSLETSLLITWQGVNETSRCNRMLKANTIIFLDMCKRCLKTHLYAPENMLTSIKCPKERNPDIKNYIDKSFPGGTEI-
CTX1_Dopa 3123 bp         367 KVGSNGVNLVVTWQGSNERSQCREIGKANTFLFLSICQSCEQSHVLISENMQSKNKCPKENYSQVKAFIDQRGPEVDKNR

                          481 .......490.......500.......510.......520.......530.......540.......550.......560
CTX1_AESCU 3159 bp        445 RKKSDVFWVAAGFKAPGNPCNHGCNGHGECKVVPYTDQFQCFCHGNYEGKMCQKKIQMKRDISKLISDLQTGYKNAFNVP
CTX2_AESCU 3129 bp        437 -PGRSLHWIAAGFKPKKEPCNNACNNHGQCKIIPYTDQIQCFCYANYVGHNCETEISGDTNFEKMVVDLQMVYTDVFKTP
CTX1_DPEA 3156 bp         445 -KKHDAFWVAAGFKSSGDACSHRCNNHGECRMVPYTDQFQCFCQTNYEGEKCETKIEINHDIVELVSDLQLGYKNAFNAP
CTX2_DPEA 3123 bp         435 -PGRRIHWVASGFKQQRNPCRNACNNQGQCKVIPYTSQIQCFCYANYIGKNCETEIVEETNIEKMVLNLQRIYKDVFKIP
CTX1_ABAN 2946bp          444 RKGVDSFWVAAGFKSYGNPCNHRCNDHGECKVVPYTDQFQCFCHDNYEGKMCDRKLQMKRNILKLVSDLQTGYKNAFKVP
CTX2_ABAN 2286 bp         437 -PDRSLHWIVAGFKSRKNPCENACNNHGQCKVIPHTDQIQCFCDANYVGHNCETEIVADTNIEKVVVDLQMVYTDVFKTP
CTX_EBER 3159 bp          446 -RKHDAFWVAAGFKSPGNPCDHRCNNHGECKIIPYTDQFQCFCQSSYEGKTCEEKIQMNKNIMKLVSDLQTGYKDAFKVP
CTX2_SOFF 3129 bp         437 -PGRITHWIAAGFKGRKQPCKNACNNHGQCKVIPYTDQIQCFCYANYVGDNCETEIAVDTNIEKMVVDLQMVYTDVFKTP
CTX1_SOFF 3141bp          439 RKKLDVFWVAAGFKAPGNPCSHRCNGHGECKVVPYTDQFQCFCHDSYEGKMCDQKIQMKRNILKLISDLQTGYKNAFKVP
CTX_ESCO 3159 bp          446 -RKHDTFWVAAGFKSPGNPCDHRCNNHGECKIIPYTDQFQCFCQSGYEGKTCEEKIQMNKNVMKLVSDLQTGYKDAFKVP
CTX2_SLES 3132 bp         438 -PGRRIYWIAAGFKRQRNPCKNACNKNGQCKVIPYTHQSQCFCYANYMGENCETEIVQETNIEKMLLDLQYVYRDAFKIP
CTX1_SLES 3159 bp         446 -KKYDAFWVAAGFKSLGNSCNHHCNNHGECRVVPYTDQFQCFCQADYEGEKCDKKIEINQDIIKLVSDLQLGYKNAFNAP
CTX_TRHO 3150 bp          444 -PSGKIHWIAAGFKPRGDPCKNACNNNGQCKIIPYTDQVQCFCYASYAGENCETEIVEDKTIEKMVLDLQLVYSDVFKTP
CTX_SATL 3147 bp          445 -KEYDTFWVAAGFKSPGNPCDHRCNDHGECKIIPYTDQFQCFCQSNYEGEKCERKIQMNKNIMKLVSDLQTGYQNAFKVP
CTX2_ALYC 3129 bp         437 -PGRSIHWIAAGYKAEKEPCKNACNNHGQCKVIPYTDQIQCFCYANYVGYNCETEISGDTNFEKMVVDLQMVYTDVFKTP
CTX1_ALYC 3159 bp         445 RKKTDIFWVAAGFKAPGNPCNHRCNDHGECKVVPYTDEFQCFCHNNYEGKMCHKKIQMKQDISKLISDLQTGYKNAFSVP
CTX_EPAR 3150 bp          444 -KEHDAFWVAAGFKSPGNPCDHRCNNHGECKIIPYTDQFQCFCQSSYEGKKCERKIQMNKSIMKLVSDLQTGYQDAFKVP
CTX2_APHAR 3129 bp        437 -PGRSIQWIAAGFKAEKEPCKNACNNHGQCKVIPYTDQIQCFCYANYVGYNCETEISGETNFEKMMVDLQMVYTDVFKTP
CTX1_APHAR 3159 bp        445 KKKTDIFWVAAGFKASGNPCNHRCNGHGECKVVPYTDQFQCFCHNNYEGKMCDKKIQMKRDISKLISDLQSGYKNAFSVP
CTX_WSCI_PARTIAL 1653 bp  179 -EGHDAFWIAAGFKVSRNPCSHRCNNHGQCKVVPYTDQFQCFCYADYEGQMCEREIKMNQDITKLVTDLQSGYKDAFKVP
CTX_IHAL 3156 bp          448 -KGYDAFWLAAGFSSETNPCSHRCNNRGECKVVPYTNQFQCFCRNNYEGEKCESKIEIKNDIQKMLSDLQNGYKSAFKVS
CTX2_XNOT 3183 bp         455 -LNDEIFWIAAGFKPRKNPCKNACNNNGRCKAIPFTNEIQCFCFANFDGEQCEIEIVENTSFEKMILDLESIYSDVFKTP
CTX1_XNOT2 3150 bp        446 -KKYDAFWLAAGISNERNPCSHRCNNRGECKVVPYTNQFQCFCQNNYEGEKCESKIEVKNDIQKLLSDLQNGYKSAFKVS
CTX_OROB_PARTIAL 1431 bp  289 -KKKDVFWVAAGFKASGNPCSHRCTNRGKCKVVPYTNQFQCFCQASYEGEKCESQIEMNDDITKLVSDLQSGYKSAFKAP
CTX_SAFF 3147 bp          445 -KEYDTFWVAAGFKSPGNPCDHRCNDHGECKIIPYTDQFQCFCQSNYEGEKCERKIQMNKNIMKLVSDLQTGYQNAFKVP
CTX_SOBS 3156bp           445 -KEYDTFWVAAGFKSPGNPCDHRCNNHGECKIIPYTDQFQCFCQPRYEGEKCERKIQLNKNIMNLVTDLQSGYKNAFKVP
CTX_Sarc 2907 bp          258 -KEHDVFWVAAGFKSSGNPCDHRCNNHGKCKVVPYTDQFECFCQASNEGEMCESKIEMNQAITKLLSDRAISVHLRFLQ-
CTX2_Dopa 3123 bp         435 -PGRRIHWIASGFKQKMKPCRNACNNQGQCKVIPYTSQIQCFCYANYIGKNCETEIVEETNIEKMVLNLQRIYRDAFKIP
CTX1_Dopa 3123 bp         447 -EKHDAFWVAAGFKSSGDACGHRCNNHGECRLVPYTDQFQCFCQTNYEGEKCETKIEINQDIVELVSDLQLGYKNAFNAP

                          561 .......570.......580.......590.......600.......610.......620.......630.......640
CTX1_AESCU 3159 bp        525 SLTNILIQGENLAKQLKKMIQRIDNQFELTHILVKYISDLQKLDY-ILKISF-------------------NYSKKKITV
CTX2_AESCU 3129 bp        516 MISSVLIQAEDLSNQLRNMLQRIDRQFELTQILVKYISPLQKIDY-LLDLSF-------------------TYRTKAITV
CTX1_DPEA 3156 bp         524 SLVNILIQGENLSNKLKRMMQRIDSQFELTQMLVKYIRDMQKLDY-ILKLGF-------------------LYSKKRISV
CTX2_DPEA 3123 bp         514 SISSILIQGENLSKQMKYMMQRIDKQFELTHILVKYINSLQKFDY-LLELSF-------------------AYQKKEITL
CTX1_ABAN 2946bp          524 SLTNILIQGENLAKQLKKMMQRIDSQFELTNILVKYIKDLQKLDY-VLKLSF-------------------NYSKKKITV
CTX2_ABAN 2286 bp         516 SISNVLIQAEDLSNQLRNMLQRIDRQFQLTQILVKYINPLQKIDY-LLDLSF-------------------SYRRKAISV
CTX_EBER 3159 bp          525 SLANILIQSENLSKQLKKMMRRINNQFELTQILIKYIRDLQKLDY-ILKLSF-------------------NYSKQKISI
CTX2_SOFF 3129 bp         516 SISSVLIQAEDLSNQLKNMLQKIDRQFELTHILVKYINPLQKIDY-LLDLSF-------------------SYRTKAITV
CTX1_SOFF 3141bp          519 SLTNVLIQGENLAKQLKKMMQKIDNQFELTQILVKYINDLQKLDY-ILKISF-------------------YYSQSKISV
CTX_ESCO 3159 bp          525 SLANILIQSENLSKQLKKMMRRINNQFELTQILIKYIRDLQKLDY-ILKLSF-------------------NYSKQKISI
CTX2_SLES 3132 bp         517 SISNVLIQGEDLSKQMKFMIQRIDKQFELTHILVKYINSLQKFDY-LLELSF-------------------AYKKKAITL
CTX1_SLES 3159 bp         525 SLVNVMIQGENLAKKLKRMMQRIDNQFELTQILVKYIRDLQKLDY-ILKLGF-------------------NYSRKRISV
CTX_TRHO 3150 bp          523 SISSVLIQGEDLSKQLRNLIQRIDQQFEMTHSLVKYISSLQKIDY-LLELSF-------------------AYRTKVITI
CTX_SATL 3147 bp          524 SLANVLIQSENLSKQLKKMMRRINNQFELTQILIKYIRDLQKLDY-ILKLSF-------------------NYSKRKISI
CTX2_ALYC 3129 bp         516 TISSVLIQAEDLSTQLRNMLQRIDRQFELTQILVKYISPLQKIDY-LLDLSF-------------------TYRTKAITV
CTX1_ALYC 3159 bp         525 SLTNILIQGENLAKQLKKMIQRIDNQFELTHILLKYISDLQKLDY-ILKISF-------------------SYSKKKITV
CTX_EPAR 3150 bp          523 SLANILIQSENLSKQLKKMMRRINNQFELTQILIKYIRDLQKLDY-ILKLSF-------------------NYSKQKISI
CTX2_APHAR 3129 bp        516 TISSVLIQAEDLSTQLRTMLQRIDRQFELTQILVKYISPLQKIDY-LLDLSF-------------------TYRTKAITV
CTX1_APHAR 3159 bp        525 SLTNVLIQGENIAKQLKKMIQRIDNQFELTHILLKYISDLQKLDY-ILKISF-------------------SYSKKKITV
CTX_WSCI_PARTIAL 1653 bp  258 SMANILIQGENLGKELKKMMRRLNNQFEMTHILVKYIGDLQKLDY-ILKLGF-------------------DYSKKKINV
CTX_IHAL 3156 bp          527 SLANILTQAENLAKQLKKMIQRMDTHFELTHILVKYIRDLQKLDY-ILKLSF-------------------NYSKRKISI
CTX2_XNOT 3183 bp         534 SISSVLIQGEDLSKQLKKMIQKINTQFEMTNILIKYIHSLQKLHY-LLELNF-------------------NYQKRTISI
CTX1_XNOT2 3150 bp        525 SLANILTQAENLSKQLKKMIQRMDTHFELTHILVKYIRDLQKLDY-ILKLSF-------------------NYSKRKISV
CTX_OROB_PARTIAL 1431 bp  368 SLTNVLILGENLSSQLKKMLRMIDNQFELTQILIKYIGDLQKLDY-ILKLSF-------------------DFSKKRISV
CTX_SAFF 3147 bp          524 SLANVLIQSENLSKQLKKMMRRINNQFELTQILIKYIRDLQKLDY-ILKLSF-------------------NYSKRKISI
CTX_SOBS 3156bp           524 SLANVLIQTENLSKQLKKMMRRINNQFELTQILVKYIRDLQKLDY-ILKLSF-------------------NYSKRKISI
CTX_Sarc 2907 bp          336 --WQIF*YKENIS-QMKKMMRRINNQFELTQMLTKC*PNVDQIYQRSTKIGLYP*AQLCLRQKEN*CGFVQS*NEEVLGS
CTX2_Dopa 3123 bp         514 LISSILIQGENLSKQMRYMMQRIDKQFELTHILVKYVNSLQKFDY-LLELSF-------------------AYQKKEITL
CTX1_Dopa 3123 bp         526 SLVNILIQGENLSMKLKRMMQRIDNQFELTQMLVKYIRDMQKLDY-ILKIGF-------------------LYSKKRISV

                          641 .......650.......660.......670.......680.......690.......700.......710.......720
CTX1_AESCU 3159 bp        585 DAFSRRMKAFLSLNPVDFIFQQLSNAILAEGFTDIQGKDFFNTFKRMIASNRDACTAPYGNEATILLERLSR--------
CTX2_AESCU 3129 bp        576 DAYNRRMKSFLSHNDVPYIFKQLSNALLANGFADKSGNDFFNIFKKMIASDRGACTKKYGTEASILFGSLSR--------
CTX1_DPEA 3156 bp         584 DAFSRRMKKFLSLNSINFIFEQLSNALLAEGFTDTKGRDFFNTFKRMIASNNEACSENYGNQATILFDRISR--------
CTX2_DPEA 3123 bp         574 DVYNLRMKSFLSHNDIYFIFKQLTNALLANGFADKSGSDFFNTFKKIIALDRGACTNQYGKEATALLKRLSR--------
CTX1_ABAN 2946bp          584 DAFSRRMKAFLSLNPADFIFQQLSNAILAEGFTDIEGKDFFNTFKRMIASNRGSCTEEYVNEANILFDRLSR--------
CTX2_ABAN 2286 bp         576 DAYNRRMKSFLSHNDVPFIFKQLSNALLANGFADKSGNDFFNIFKKMIASEKGACTKKYGTEAAVLLNSLSR--------
CTX_EBER 3159 bp          585 DAFNRRMKTFLSLNPINFMFQQLSNAILADGFTDMKGKDFFNTFKRMIATDRDACTEEYGLEASKLFERLSQ--------
CTX2_SOFF 3129 bp         576 DAYSQRMKSFLSHNDVPFIFKQLSNAFLADGFADRSGNDFFNIFKKMIASDRGSCTKKYGTDAAILLDSLSR--------
CTX1_SOFF 3141bp          579 DAFSRRMKRFLSLNPIDFIFQQLSNAILAEGFTDIRGKDFFNTFKRMIASNRGACTEQYGHEASILFDRLSR--------
CTX_ESCO 3159 bp          585 DAFNRRMKTFLSLNPINFMFQQLSNAILADGFTDTKGKDFFNTFKRMIATDRDACTEEYGLKASKLFERLSQ--------
CTX2_SLES 3132 bp         577 DVYNLRMKSFLSHNDIYFIFKQLTNALLANGFADKSGSDFFNTFKKIIASDSGACTEQYGNEATILLNRLSR--------
CTX1_SLES 3159 bp         585 DAFSRRMKKFLSLNPINFIFEQLSNAILAEGFTDTKGRDFFNTFKRMIASNREACSKIYGNEATVLFNRLSR--------
CTX_TRHO 3150 bp          583 NAYNRRIKSFLSHNDVPFIFKQMANAVLAEGFADKSGSDFFNLFKKMIASNRGACTEQYGNDSIALLNRLNR--------
CTX_SATL 3147 bp          584 DSFNRRMKTFLSLNPINFIFQQLSNAIMADGFTDTKGKDFFNTFKRMIATDRDACTEEYGIEASRLFDRLSQ--------
CTX2_ALYC 3129 bp         576 DAYNRRMKSFLSHNDVPFIFKQLSNALLANGFADKSGNDFFNIFKKMIASDRGACTKKYGTEATILLESLSR--------
CTX1_ALYC 3159 bp         585 DAFSRRMKAFLSLNPIDFIFQQLSNAILAEGFSDIRGKDFFNTFKRMIASNRDACTTQYGSEATILLERLSR--------
CTX_EPAR 3150 bp          583 DAFNRRMKTFLSLNPINFMFQQLSNAIMADGFTDTKGKDFFNTFKRMIATDRDACTEEYGLEASKLFDRLSQ--------
CTX2_APHAR 3129 bp        576 DAYNRRMKSFLSHNDVHFIFKQLSNALLANGFADKSGNDFFNIFKKMIASDRGACTKKYGTEATILLDSLSR--------
CTX1_APHAR 3159 bp        585 DAFSRRMKAFLSLNPIDFIFQQLSNAILAEGFTDIRGKDFFNTFKRMIASNRDACSTQYGSEATILLERLSR--------
CTX_WSCI_PARTIAL 1653 bp  318 DAFSRRMKKFLSLNPINFVFQQLSNAILANGFTDTRGNDFFNTFKRMIASNRGACTEDYGKDASNLMDRLSR--------
CTX_IHAL 3156 bp          587 DSYNRRMKKFLGLNPIHFIFQQLSNAILAEGFTDAKGQDFFNTFKKMIASNRGACSAEYGNEAQVLFERLSE--------
CTX2_XNOT 3183 bp         594 DVFNLKMKHFLLHNDVAFIFDQLSNAILAEGFADKRGHDFFNTFKMMIASSVGACSVQYGMETTGVFNHLSL--------
CTX1_XNOT2 3150 bp        585 DSYNRRMKKFLALNPIHFIFQQLSNAILAEGFTDSKGKDFFNTFKKMIASNRGACSAEYGNQAQLLFERLSE--------
CTX_OROB_PARTIAL 1431 bp  428 DAFSRRMKKFLSLNPVNFVFQQLSNAILADGFTDAKGSDFCNTFKRMIAA------------------------------
CTX_SAFF 3147 bp          584 DSFNRRMKTFLSLNPINFIFQQLSNAIMADGFTDTKGKDFFNTFKRMIATDRDACTEEYGIEASRLFDRLSQ--------
CTX_SOBS 3156bp           584 DGFSRRMKAFLSLNPIDFIFQQLSNAIMADGFTDTKGKDFFNTFKRMIATDRDACTEEYGIEASRLFERLSQ--------
CTX_Sarc 2907 bp          408 QSN*FRLPTIVERDPCEWIHRHQRQRFLQHVQKDGR-----FRQRRVYPTIRKRCSDINGSSQSIRYHRCRSHSCLLQFR
CTX2_Dopa 3123 bp         574 DVYNLRMKSFLSHNDIYFIFKQLTNALLANGFADKSGSDFFNTFKKIIALDRGACTNQYGKEATALLKRLSR--------
CTX1_Dopa 3123 bp         586 DAFSRRMKKFLSLNSINFIFEQLSNALLAEGFSDTKGRDFFNTFKRMIASNNEACSEKYGNEVTILFDRISR--------

                          721 .......730.......740.......750.......760.......770.......780.......790.......800
CTX1_AESCU 3159 bp        657 ------------------------------LDLTAAESILAYYSFESNYLNPANMKRMLENAKQ----------------
CTX2_AESCU 3129 bp        648 ------------------------------MDITAAEAILAHYYFESFYLNAENKAIMRSRVDQ----------------
CTX1_DPEA 3156 bp         656 ------------------------------LDITAAQAILAYYNFESNYLNKGNMKKMFNTAKQ----------------
CTX2_DPEA 3123 bp         646 ------------------------------MDIAAAEAILAHYHFQSFYLNAEHKKNMLSSAMQ----------------
CTX1_ABAN 2946bp          656 ------------------------------LDITAAETILAYYSFESNYLNAENMKQMHVNAKQ----------------
CTX2_ABAN 2286 bp         648 ------------------------------MDITAAEAILAHYYFESFYLNAEHRTTMLSSVDQ----------------
CTX_EBER 3159 bp          657 ------------------------------LDIMAAEAILAHYHFQSNYLTEEHMKTMQASAKQ----------------
CTX2_SOFF 3129 bp         648 ------------------------------MDITAAEAILAHYHFESFYLNAEHKTTMLSSAVQ----------------
CTX1_SOFF 3141bp          651 ------------------------------LDITAAESILAYYSFESNYLNPENMKRMLENAKQ----------------
CTX_ESCO 3159 bp          657 ------------------------------LDIMAAEAILAHYHFQSNYLTEEHMKTMQASAKQ----------------
CTX2_SLES 3132 bp         649 ------------------------------MDITAAEAILAHYHFESFYLDAERKANMLSNAMQ----------------
CTX1_SLES 3159 bp         657 ------------------------------IDITAAEAILAYYNFESNYLNTENMKKMTIAAKQ----------------
CTX_TRHO 3150 bp          655 ------------------------------LDITAAEAILAHYHFESFYLKPEQKTNMLFNAKQ----------------
CTX_SATL 3147 bp          656 ------------------------------FDIMAAEAILAHYHFQSNYLTPEHMKTMQASAKQ----------------
CTX2_ALYC 3129 bp         648 ------------------------------MDITAAEAILAHYYFESFYLNVENKATMLSSVDQ----------------
CTX1_ALYC 3159 bp         657 ------------------------------LDITAAESILAYYSFESNYLNPENMKRMLENAKQ----------------
CTX_EPAR 3150 bp          655 ------------------------------LDIMAAEAILAHYHFQSNYLTEEHMKTMQASAKQ----------------
CTX2_APHAR 3129 bp        648 ------------------------------MDITAAEAILAHYYFESFYLNAENKATMLSSVDQ----------------
CTX1_APHAR 3159 bp        657 ------------------------------LDITAAESILAYYSFESNYLNPENMKRTLENAKQ----------------
CTX_WSCI_PARTIAL 1653 bp  390 ------------------------------LDITAAETILAYYNFESNYLNKKNMRKMIADAKQ----------------
CTX_IHAL 3156 bp          659 ------------------------------LDITAAEAILAHYEFESNYLNSENMDSMMESAKQ----------------
CTX2_XNOT 3183 bp         666 ------------------------------LDMTATETILAHYQFEDLYLKDDHRSNMIKEAKH----------------
CTX1_XNOT2 3150 bp        657 ------------------------------LDITAAEAILAHYQFESNYLNSEHMASMKESAKQ----------------
CTX_OROB_PARTIAL 1431 bp      --------------------------------------------------------------------------------
CTX_SAFF 3147 bp          656 ------------------------------LDIMAAEAILAHYHFQSNYLTPEHMKTMQASAKQ----------------
CTX_SOBS 3156bp           656 ------------------------------FDIMAAEAILAHYHFQSNYLTTEHMKTMQTSAKQ----------------
CTX_Sarc 2907 bp          482 K*LSQEGAFERDAR*CKAIGTRFKTTHAKLCQILFV*LILQHTNPM-GHLMLKHF*SLATLVKKVCFVSNRCLATLVNKS
CTX2_Dopa 3123 bp         646 ------------------------------MDIAAAEAILAHYHFQSFYLNAEHKKNMLSSAMQ----------------
CTX1_Dopa 3123 bp         658 ------------------------------LDITAAQAILAYYNFESNYLNRGNMKKMFNSAKQ----------------

                          801 .......810.......820.......830.......840.......850.......860.......870.......880
CTX1_AESCU 3159 bp        691 ----------------------------------------LVRDSKRRMRSYARYWERTSCPPLNVTHLTQTGCGALLSF
CTX2_AESCU 3129 bp        682 ----------------------------------------LAQDSKERMKNYASYWERTSCPPLNIPYLNQTGCGELLSF
CTX1_DPEA 3156 bp         690 ----------------------------------------LVRDSKRRMRKYSNYWENTSCPPLNVTYVRQNGCGAMLSF
CTX2_DPEA 3123 bp         680 ----------------------------------------LAQGSTERMQNYARYWERTSCPPLNVTHLTQTGCGTMLSF
CTX1_ABAN 2946bp          690 ----------------------------------------LVQDSKRRMRNYARYWERTSCPPLNVTHLTQAGCGALLSF
CTX2_ABAN 2286 bp         682 ----------------------------------------LVKDSKERMKNYARYWERTSCPPLNVTYLNETGCGELLSF
CTX_EBER 3159 bp          691 ----------------------------------------LVRDSKRRMRNYARYWERTSCPPLNVTHLRQNGCSNMLSF
CTX2_SOFF 3129 bp         682 ----------------------------------------LVQDSKERMKNYARYWERTSCPPLNITYLNQTGCGELLSF
CTX1_SOFF 3141bp          685 ----------------------------------------LVRDSKRRMRNYARYWERTSCPPLNITQLTQTGCGDLLSF
CTX_ESCO 3159 bp          691 ----------------------------------------LVRDSKRRMRNYARYWERTSCPPLNVTHLRQNGCGNMLSF
CTX2_SLES 3132 bp         683 ----------------------------------------LVQGSTERMKNYARYWERTSCPPLNVTYLTQIGCGAMLSF
CTX1_SLES 3159 bp         691 ----------------------------------------LVRDSKRRMRNYAKYWEGTSCPPLNVTYLTQSGCGAMLSF
CTX_TRHO 3150 bp          689 ----------------------------------------LAQDSKERLKMYANYWERTSCPPLNVTYLTQTGCGAMLSY
CTX_SATL 3147 bp          690 ----------------------------------------LVRDSKRRMRNYARYWERTSCPALNVTHLRQNGCGDMLSF
CTX2_ALYC 3129 bp         682 ----------------------------------------LVQDSKERMKNYARYWERTSCPPLNIPYLNQTGCGELLSF
CTX1_ALYC 3159 bp         691 ----------------------------------------LVRDSKRRMRSYARYWEHTSCPPLNVTNLAQTGCGAFLSF
CTX_EPAR 3150 bp          689 ----------------------------------------LVRDSKRRMRNYARYWERTSCPALNVTHLRQNGCGNMLSF
CTX2_APHAR 3129 bp        682 ----------------------------------------LVQDSKERMKNYASYWERTSCPPLNIPYLNQAGCGELLSF
CTX1_APHAR 3159 bp        691 ----------------------------------------LVRDSKRRMRSYARYWEHTSCPPLNVTNLSQTGCGAFLSF
CTX_WSCI_PARTIAL 1653 bp  424 ----------------------------------------LVRDSKRRMQSYAKYWERNSCPPLDVPGLEQTGCGSMLSF
CTX_IHAL 3156 bp          693 ----------------------------------------LVIDSKRRMRSYARYWERTSCPPLNVSYLVQNGCGYLLSF
CTX2_XNOT 3183 bp         700 ----------------------------------------FTQGSKERMKNYVKYWHKTSCPPLNATHLIQVGCSAMLSY
CTX1_XNOT2 3150 bp        691 ----------------------------------------LVRDSKRRMRSYAKYWERTSCPPLNASFLVQNGCGDLLSF
CTX_OROB_PARTIAL 1431 bp      --------------------------------------------------------------------------------
CTX_SAFF 3147 bp          690 ----------------------------------------LVRDSKRRMRNYARYWERTSCPALNVTHLRQNGCGDMLSF
CTX_SOBS 3156bp           690 ----------------------------------------LVRDSKRRMRNYARYWERTSCPPLNVTYLRQNGCGSMLSF
CTX_Sarc 2907 bp          557 GGGDE*GLV*ILSFDLSGLVGPTRNLIPASVALRVIGTFKPLHHAKVLIRGVAKYWERTSCPTLNVTGLTQTGCGAMLSF
CTX2_Dopa 3123 bp         680 ----------------------------------------LAQGSTERMRNYARYWEKTSCPPLNVTHLTQTGCGTMLSF
CTX1_Dopa 3123 bp         692 ----------------------------------------LVRDSKRRMRKYANYWENTSCPPLNVTYLNQSGCGAKLSF

                          881 .......890.......900.......910.......920.......930.......940.......950.......960
CTX1_AESCU 3159 bp        731 EGMKVKLSCDGGRAAVPQNIECVNVNGNLQWSATPKCESSWSR------------WSKWSACASTCGNATQSRRRRCLGQ
CTX2_AESCU 3129 bp        722 EGMKVKLSCDGGREAVHTAIECSISEGKLKWNATAKCVPAWSA------------WGEWSSCSSTCGSGTQTRTRVCRGE
CTX1_DPEA 3156 bp         730 EGMEVSLSCDGNRAVEPQNVKCVKLNGKLQWSATPRCLASWSQ------------WSQWTPCTSTCGNGTQSRQRICNGE
CTX2_DPEA 3123 bp         720 EGMKVKLFCDGGREAVPPVIECNLSKGKLKWNATAKCVPGWST------------WGEWSSCSSTCGQGVQTRSRECLGE
CTX1_ABAN 2946bp          730 EGMKVKLSCDGGRAAVPRTIECVSVNGKLQWSATPKCAAGWSK------------WSKWTACASTCGNATQSRRRRCNGQ
CTX2_ABAN 2286 bp         722 EGMKVKLSCVEGMEVIPKEIECRLSEGKLEWNATVKCVPG*---------------------------------------
CTX_EBER 3159 bp          731 EGMTVKLSCAEGRVAEPRNIECVNSRGKLQWSATPRCLAKWSQ------------WTQWEPCSATCGDGIQLRRRTCQGD
CTX2_SOFF 3129 bp         722 EGMKVKLSCDGGREAVPTAIECRLSGGKLEWNATAKCVPGWSA------------WGEWGSCNSTCGLGIQTRKRECRGE
CTX1_SOFF 3141bp          725 EGMKVKLSCDGGRAAVPKTIECVNVNGNLQWSATPKCAASWSR------------WSQWTACASTCGNATKSRRRRCIGE
CTX_ESCO 3159 bp          731 EGMRVKLSCVGGRVAEPQNIECVNSRGKLQWSATPRCLAKWSP------------WSQWEPCSATCGDGIQLRRRTCQGD
CTX2_SLES 3132 bp         723 QGMKVKLFCDGGREAVPPVIECNLSEDKLEWNATAKCVPGWST------------WREWSSCSSTCGPGVQTRSRECLGE
CTX1_SLES 3159 bp         731 EGMNVSLSCDGDRAVEPQNIECIKLNNKLQWSATPRCVAGWSN------------WSQWTPCTSTCGNGTQSRRRICNGE
CTX_TRHO 3150 bp          729 EGMKVKLNCDGGRKAIPPVIECSLTEGKLEWNATAKCVPGWSA------------WGEWGSCSSTCGSGIQIRNRECLGE
CTX_SATL 3147 bp          730 EGMKVKLSCVDGRVAEPRNIECVNSHGKLQWSATPRCLAKWSE------------WSQWGSCSATCGGGVQSRRRFCQGD
CTX2_ALYC 3129 bp         722 EGMKVKLSCNGGRQAVPTAIECSLSEGKLKWNATAKCVPGWSA------------WGEWSSCSSTCGPGTQTRTRECLGE
CTX1_ALYC 3159 bp         731 EGMKVKLSCDGGRAAVPQNIECVNVNGNLQWSATPKCEASWSR------------WSKWSACASTCGNATQTRRRRCLGQ
CTX_EPAR 3150 bp          729 EGMKVKLSCAGGRVAEPRNIECVNSHGKLQWSATPRCLAKWSQ------------WSQWEPCSATCGDGVQSRRRTCQGD
CTX2_APHAR 3129 bp        722 EGMKVKLSCDGGREVVPTAIECILSEGKLKWNATAKCVPGWSA------------WGEWSSCSSTCGPGTQTRTRECLGE
CTX1_APHAR 3159 bp        731 EGMKVKLSCDGGRAAVPKSIECVNVNGNLQWSATPKCEASWSR------------WSKWSACASTCGNATQSRRRRCLGQ
CTX_WSCI_PARTIAL 1653 bp  464 EGMKLKVSCDGGKVAKQRNIECVKRKAGLQWSATPSCSASWSK------------WSSWSQCSSSCGNATQTRRRVCNGG
CTX_IHAL 3156 bp          733 EGMKVDLACEDGRRAAPQRIECVKVGGKLQWSAEAECKSSWSE------------WSEWSSCPEMCDSSTTTRSRVCNGS
CTX2_XNOT 3183 bp         740 EGMKVELSCEGGRQAVPPVIECRSFGNKLNWNSSIHCAPNWSI------------WGEWSSCSSTCGRGSQIRSRKCLGE
CTX1_XNOT2 3150 bp        731 EGMKVDLACEGGRRAAPQTIECVKVNRKLQWSAMPECKSSWSE------------WSEWSSCPETCDTSISTRSRVCNGN
CTX_OROB_PARTIAL 1431 bp      --------------------------------------------------------------------------------
CTX_SAFF 3147 bp          730 EGMKVKLSCVDGRVAEPRNIECVNSHGKLQWSATPRCLAKWSE------------WSQWGPCSVTCGGGVQSRRRFCQGD
CTX_SOBS 3156bp           730 EGMKVKLSCVDGRVAEPQNIECVSSRGKLQWSATPKCLSTWSR------------WSKWTPCSATCGDGVQSRRRTCQGE
CTX_Sarc 2907 bp          635 EGMKATLSCNGDRESSRAANY*M---C*DEW*VAMDCHTHVCRELVRVVCVDSLCFHLW-KCYPVS*SGTQRRE------
CTX2_Dopa 3123 bp         720 EGMKVKLFCDGGREAVPPVIECNLSKGKLEWNATAKCVPGWST------------WGEWSSCSSTCGQGVQTRSRKCLGE
CTX1_Dopa 3123 bp         732 KGMEVSLSCDGDRAVEPQNVKCVKLNGKLQWSATPRCLASWSQ------------WSQWTPCTSTCGNGTQSRKRICNGE

                          961 .......970.......980.......990......1000......1010......1020......1030......1040
CTX1_AESCU 3159 bp        799 SESEKCIGPSKQVRKCFVEDCCQEKYGKFKCDNNKCISLSRVCDGNDDCRNAEDESKSRCKYLRSGDRIALRNMAYSQEW
CTX2_AESCU 3129 bp        790 TQDEQCKGSSTDRRSCSSEDCCQAKFGKFKCPVGWCIDLSKVCDGTTDCAIDADEAKGRCNYLRSGDRIALRNLASSFDW
CTX1_DPEA 3156 bp         798 SESEKCEGSATDVRNCFLTDCCQKIYGKFKCDDTKCLDISQLCDGFDDCHDGTDESKDKCKYLRSGDRIALRNMASSQDW
CTX2_DPEA 3123 bp         788 TNDEHCNGSSKDTRTCNKEDCCQEKYGKFKCPVGWCIDLSRVCDGTADCALDEDEAKTRCNYLRSGNRIALRNLATSSDW
CTX1_ABAN 2946bp          798 SESERCKGPSTQVRKCSVEDCCQEKYGKFKCDSDKCIPLSQVCDGTDNCLDKTDELKSRCNYLRSGDRIALRNMAFSQEW
CTX2_ABAN 2286 bp             --------------------------------------------------------------------------------
CTX_EBER 3159 bp          799 PSGDQCQGQAVVKRKCFVQDCCHTKFGKFKCDNSTCLSISQVCDGIDDCLDGTDESSTKCKYLRSGNRIALRNMDYSADW
CTX2_SOFF 3129 bp         790 DQDQQCKGSSTDKRSCSSEDCCQAKFGKFKCPVGWCIDLSRVCDGTTDCATDADESKGRCNYLRSGDRIALRNLASSVDW
CTX1_SOFF 3141bp          793 SDTDKCKGQSSQVRKCFVEDCCQEKYGKFKCGNDKCISISQVCDGIDDCLDESDESKSRCKYLRSGDRIALRNMGFSQEW
CTX_ESCO 3159 bp          799 PSGGQCQGQAVVKRKCFVQDCCHVKFGKFKCDNSTCLSISQVCDGIDDCLDGTDESSTKCKYLRSGNRIALRNMDYSMDW
CTX2_SLES 3132 bp         791 TKSEHCKGSSKDTRSCSTEDCCQEKFGKFKCPIGWCIDLSRVCDGTADCALDEDEAKARCNYLRSGNRIALRNLATSSDW
CTX1_SLES 3159 bp         799 SEGEKCKGLTTDVRNCFLSDCCQEMYGKFKCDSTKCLDLSQLCDGFDDCLDGIDESKERCKYLRSGDRIALRNMAFSQEW
CTX_TRHO 3150 bp          797 TEEEKCQGTPKDTRTCSNEDCCQAKFGKFKCPVGWCIDLLRVCDGTTDCALDADEARGRCNYLRSGDRIALRHLASSFDW
CTX_SATL 3147 bp          798 ---DQCQGRAVATRKCFMQDCCHAKFGKFKCDNSTCLRISQVCDGIDDCSDGSDESSTKCKYLRSGDRIALRNMDYSQDW
CTX2_ALYC 3129 bp         790 TQNEQCKGSSTDRRSCSNEDCCQAKFGKFKCPVGWCIDLSKVCDGTTDCAIDADEAKGRCNYLRSGDRIALRNLASSFDW
CTX1_ALYC 3159 bp         799 SESEKCKGPSKQVRKCFVEDCCQEKYGKFKCNNNKCISLSRVCDGNDDCRDAADESKSRCKYLRSGDRIALRNMAYSQEW
CTX_EPAR 3150 bp          797 PGGDQCQGQAVVSRKCFTQDCCHAKFGKFKCDNSTCLSISQVCDGIDDCLDGTDESSTTCKYLRSGDRIALRNMDSSMDW
CTX2_APHAR 3129 bp        790 TQDEQCKGSSIDRRSCSSEDCCQAKFGKFKCPVGWCINLSKVCDGTTDCAIDADEAKGRCNYLRSGDRIALRSLASSFDW
CTX1_APHAR 3159 bp        799 SDSEKCKGPSKQVRKCFVEDCCQEKYGKFKCNNNKCISLSRVCDGNDDCLDAADESKSRCKYLRSGDRIALRNMAYSQEW
CTX_WSCI_PARTIAL 1653 bp  532 ----TCGGASTDVRSCPFKDCCQE--------------------------------------------------------
CTX_IHAL 3156 bp          801 -PGETCVGQASQEKKCFPEDCCQAKYGKFRCDAETCLDVSDICNGVEQCSDGSDES--KCDYLRSGDRIALRNMRYSQDW
CTX2_XNOT 3183 bp         808 THTEHCKGEQDETRSCSTEDCCQGKYGKFKCPVGWCLDLSQVCDGITHCASDADESTATCNYLRSGKRIALRSLGSSKNW
CTX1_XNOT2 3150 bp        799 -EGEICEGPPAQKKKCFPDDCCQAKYGKFKCDTDTCLNLSDICNGIEQCSDGSDES--QCDYVRSGDRIALRNMRFSQDW
CTX_OROB_PARTIAL 1431 bp      --------------------------------------------------------------------------------
CTX_SAFF 3147 bp          798 ---DQCQGRPVATRKCFMQDCCHAKFGKFKCDNSTCLRISQVCDGIDDCSDGSDESSTKCKYLRSGDRIALRNMDYSQDW
CTX_SOBS 3156bp           798 SGGEQCQGQAVATRKCSMQDCCQVKFGKFRCDNSTCLRRSQVCDGIDDCSDGSDESSTKCKYLRSGDRIALRNMDSSQDW
CTX_Sarc 2907 bp          701 ----------MQRVVNFSRRLLSREYGKFKCDDSKCLDISQVCDGIDHCVDRTDESKARCKYLRSRDKIALRNVAYSQEW
CTX2_Dopa 3123 bp         788 TNDEHCNGSSKDTRTCNKEDCCQEKYGKFKCPVGWCIDLSRVCDGTADCALDEDEAKTRCNYLRSGNRIALRNLATSSDW
CTX1_Dopa 3123 bp         800 SESEKCEGSATDGRNCFLTDCCQEIYGKFKCDDTKCLDISQLCDGFDDCHDGTDESKDKCKYLRSGDRIALRNMASSQDW

                         1041 ......1050......1060......1070......1080......1090......1100......1110......1120
CTX1_AESCU 3159 bp        879 LSVQYTDAVQ-ADLYYGRAYLNHCIKGDHVTSSEWNSCAGQSLLIYGNYENGKIGKAIRFGDKIAMYYRKTNYHYRWFIC
CTX2_AESCU 3129 bp        870 LSVRNTDYLSTPQLRYGRAFLDRCIKTNRVIESEWRECVGNSMLIYGSHNNARIGQSIRYGDRIAMYYRKTHSHFRWFRC
CTX1_DPEA 3156 bp         878 LSVKYTDAVQ-ADLYYGRAYLDRCIKGDQVTDYEWKNCAGQSMLIYGNYKNGETGKAIKYGDKIAIYYRKTNFHYRWLRC
CTX2_DPEA 3123 bp         868 LSVLYTDYVVTPHLRYGRAYLNRCIKSERVAASEWKQCEGNSMLVYGSHNQGRDGQSIRYGDRIALYYRKTYSEYRWFRC
CTX1_ABAN 2946bp          878 LSVQYTDYVQ-ANLYYGRAYLDHCIKGDRVTSSEWDSCAGQSMLIYGNYENGKTGKAIRFGDKIAMYYRKTEFHYRWFIC
CTX2_ABAN 2286 bp             --------------------------------------------------------------------------------
CTX_EBER 3159 bp          879 LSVQYTDYVQ-ANLYYGRAYLDRCIKGDDVTDYEWKNCPGQSMRVYGNYQNGKSGKAIRYGDKIALYYRKTNYHYRWFRC
CTX2_SOFF 3129 bp         870 LSVLRTDYLNSPQLQYGRAFLDRCIKTHHITESEWRQCKGNSMLIYGSHNNARVGQSIRYGDRIAMYYRKTYSQFRWFRC
CTX1_SOFF 3141bp          873 LSVQYTDAVQ-ANLYYGRAYLNHCIKGTQVTSSEWKSCPGQSMLIYGNYKNGKAGKAIRFGDKIAMYYRKTAYHYRWFIC
CTX_ESCO 3159 bp          879 LSVQYTDYVQ-ANLYYGRAYLDRCIKGDDVTDYEWKNCPGQSMLVYGNYQNGKSGKAIRYGDKIALYYRKTNYHYRWFRC
CTX2_SLES 3132 bp         871 LSVRYTDYVIPPHLRYGRAYLNRCIKTEKVTASEWKQCEGNSMLVYGSHNQRREGQSIRYGDRIALYYRKTYSKYRWFRC
CTX1_SLES 3159 bp         879 LSVQYTDTVQ-ADLYYGRAYLDHCIKGDDVTDNEWKNCAGQSMLIYGNYKNGQIGKAIKYGDKIAMYYRKTNYHYRWFIC
CTX_TRHO 3150 bp          877 LSLRHTDRIITPSLRYGRAHLDRCIKTDQVTAPEWKQCMDNSMLVYGSHNNGREGQSIRYGDRIAMYYRKTYSHHRWFRC
CTX_SATL 3147 bp          875 ISVQYTDYVQ-ANLYYGRAYLDRCIKGDEVTDNEWKNCPGQSMLAYGNYENGKTGTAIRYGDKIALYYRKTNYHYRWFRC
CTX2_ALYC 3129 bp         870 LSVRNTDYLNTPHLRYGRAFLDRCIKTNQVLESEWRECNGNSMLIYGSHNNARIGQSIRYGDRIAMYYRKTHSQFRWFRC
CTX1_ALYC 3159 bp         879 LSVQYTDAVQ-ANLYYGRAYLNHCIKGDRVTSSEWNSCAGQSLLIYGNYQNGKKGKAIRFGDKIAMYYRKTNYHYRWFIC
CTX_EPAR 3150 bp          877 LSVQYTDYVQ-ANLYYGRAYLDRCIKGDDVASDEWKNCPGQSMLVYGNYQNGKSGKAIRYGDKIALYYRKTNYHYRWFRC
CTX2_APHAR 3129 bp        870 LSVRHTDYLNSPQLRYGRAFLDRCIKTNQVLESEWRKCDGNSMLIYGSHNNARIGQSIRYGDRIAMYYRKTHSHFRWFRC
CTX1_APHAR 3159 bp        879 LSVQYTDAVQ-ASLYYGRAYLNHCIKGDHVTSSEWNSCAGQSLLIYGNYQNGKKGKAIRFGDKIAMYYRKTNYHYRWFIC
CTX_WSCI_PARTIAL 1653 bp      --------------------------------------------------------------------------------
CTX_IHAL 3156 bp          878 LSVQYTDAVQ-ADLHYGRAYLDDCIQDDDVTDDEWKYCAGQSLRIYGNYQNGKTGKAIRYGDRVALYYRKSNYHYRWFIC
CTX2_XNOT 3183 bp         888 LSVLYSDYVSRGDLRYGRAHVTQCIKGDFVASSEWTDCPGNSMLIYGSYNNARKGQAIRYGDRIAMYYTKTNAKYRWFTC
CTX1_XNOT2 3150 bp        876 LSVKNTDAVQ-ASLYYGRAYLDDCIHDDDVTDDEWKYCDGQSLRIYGNYQNGKTGKPIRFGDRIALYYRKTNYHYKWFIC
CTX_OROB_PARTIAL 1431 bp      --------------------------------------------------------------------------------
CTX_SAFF 3147 bp          875 ISVQYTDYVQ-ANLYYGRAYLDRCIKGDEVTDNEWKNCPGQSMLAYGNYENGKTGKAIRYGDKIALYYRKTNYHYRWFRC
CTX_SOBS 3156bp           878 ISVQYTDYVQ-ANLYYGRAYLDRCIKGDDVTENEWKNCPGQSMLAYGNYENGKTGKAIRYGDKIALYYRKTNYHYRWFRC
CTX_Sarc 2907 bp          771 LSVQYTDYVQ-SNLYYGRAYPDHCIKDDHVTDSEWKNCVGQLMNVYGSYKNGKQGTAIRYGERIVMYYRKTNYHWRWFIC
CTX2_Dopa 3123 bp         868 LSVLYTDYVVSPHLRYGRAYLNRCIKSERVAASEWKQCEGNSMLVYGSHNQGRDGQSIRYGDRIALYYRKTYSEYRWFRC
CTX1_Dopa 3123 bp         880 LSVQYTDAVQ-ADLYYGRAYLNRCIKGDEVTDYEWKNCAGQSMLIYGNYKNGEKGKAIKYGDKIAMYYRKTNYHYRWFRC

                         1121 ......1130......1140......1150......1160......1170......1180......1190......1200
CTX1_AESCU 3159 bp        958 YPTYCMTYTCPKKAGSFTFGPNGGCDEYEFYIINYNDKLSRDPVKPGDVITLANNRGSVKGNGYNRNININDCTVKRAQD
CTX2_AESCU 3129 bp        950 YSDYCITYTCDKAPGQFDFSDRGGCKSYEFVIQNYEHPNATTPIQNGDIVFIHNSNGALRGNGYWNKINQRKYSSAEI--
CTX1_DPEA 3156 bp         957 YSSYCMTYTCDKKPGTFDFGSNGGCQEYEFYITKYDDPLANGPVKAGDLVIISDGRGALKGNGYYKNIYQDGCIVKRVLD
CTX2_DPEA 3123 bp         948 YSDYCISYICSKAPGQFDFSDTGGCKSNEFIIQNYEHPNSTTPIRNGDIVIISADKGALRGNGYNHKITLKKCSFEEL--
CTX1_ABAN 2946bp          957 YPTYCMTYTCDKKWGSFDFGSQRR--------------------------------------------------------
CTX2_ABAN 2286 bp             --------------------------------------------------------------------------------
CTX_EBER 3159 bp          958 YTKYCMTYTCEVKYGSFDFSSKGGCKEYEFDITNYDDPQARGPVKAGDVVVVSNGNGAIRGKGYYKEISQSDCVKKRERN
CTX2_SOFF 3129 bp         950 YSDYCITYTCDKAPGKFDFSDIGGCKQYEFVIQNYEYPNATTPIRNGDIVFIHNSNGALKGNGYWNKITQKKCSYAEL--
CTX1_SOFF 3141bp          952 YPTYCMTYTCPKKPGSFTFGPGGGCDEYEFYISDYNDKLSRKPVKAGDIITIANNRGSIKGNGYNRNININDCTVNRALD
CTX_ESCO 3159 bp          958 YTKYCMTYTCEVRYGSFDFSSKGGCKEYEFDIINYDDPQARGPVKAGDVVVVSNGNGAIRGKGYYKEINQSDCFKKRERN
CTX2_SLES 3132 bp         951 YADYCITYICAKAPGQFDFSDRGGCKSNEFIIQNYEHPNMTTPIRNGDIVVISNDNGALKGNGYNNKITTKKCSSAEI--
CTX1_SLES 3159 bp         958 YPDNCMTYICDKSPGTFDFTSKGGCQEYEFYITKYDDRQAIGPVKSGDMVFISDGKGAIKGNGYYKNINQDLCAIKRVQN
CTX_TRHO 3150 bp          957 YSNYCMTYTCTKSPDQFDFSDVGGCKQYEFVIQNYEHPNSTAPVRNGDIVFISNNDGALKGNGYWNQISQKKCSYATI--
CTX_SATL 3147 bp          954 YTKYCMTYTCEVRYGSFDFSSKGDCKEYEFDITNYDDPQARGPVKAGDVVVISDGKGAVRGNGYYKEISQSDCVQKRQRN
CTX2_ALYC 3129 bp         950 YSDYCITYTCDKAPGQFDFSDRGGCNSYEFIIQNYEHPNATTPVQNGDIVFIHNSNGALRGNGYWNKINQKKCSSAEI--
CTX1_ALYC 3159 bp         958 YPTYCMTYTCPKTAGSFTFGPNGGCDEYEFYIINYNDNLSRDPVKAGDVITIANNRGSVKGNGYNRNINMNDCTVKRAQD
CTX_EPAR 3150 bp          956 YTKYCMTYTCEVRHGDFDFSSKGGCKEYEFDIKNYDDPQARGPVKAGDVVVVSNGNGAVKGNGYYKEVSQSDCVKKRQRN
CTX2_APHAR 3129 bp        950 YSDYCITYTCDKAPGQFDFSDRGGCKSYEFVIQNYEHPNATTPIRNGDIVFIHNSNGALRGNGYWNKINQKKCSSTEI--
CTX1_APHAR 3159 bp        958 YPTYCMTYTCPKKAGSFTFGPNGGCDEYEFYIINYNDKLSREPVKAGDVITIANNRGSIKGNGYNRNINTDDCTVNRALD
CTX_WSCI_PARTIAL 1653 bp      --------------------------------------------------------------------------------
CTX_IHAL 3156 bp          957 YPTYCMTYTCDKRYGEFDFNPRGECKEYQFYLADYNNRGSRAPVKPGDIITISNGRGSIKGNGHSKTINQDDCTVRRSQD
CTX2_XNOT 3183 bp         968 YQSYCMTYICNKAPGEFDFGEFGGCKENEFIIRNFQNPNATNPIRNGDIVFISNKDGSLKAGSLWDDITLKKCSSEKM--
CTX1_XNOT2 3150 bp        955 YPTYCMTFTCDKRYGSFDFNPQGECKDYQFYVADYNNPASRAPVKPGDVITISNGRGAIKGNGHSKTINQDDCTVRRVHD
CTX_OROB_PARTIAL 1431 bp      --------------------------------------------------------------------------------
CTX_SAFF 3147 bp          954 YTKYCMTYTCEVRYGSFDFSSKGDCKEYEFDITNYDDPQARGPVKAGDVVVISDGKGAVRGNGYNKEISQSDCVQKRQRN
CTX_SOBS 3156bp           957 YTKYCMTYTCEVRHGSFDFSSKGDCKEYEFDITNYDDPQARGPVMAGDVVVISNGNGAVRGNGYYKEISQSDCVKKRERN
CTX_Sarc 2907 bp          850 YPKYCMTYTCGKKAGTFDFGHEGRCEEYEFCIEKYDDLQARGPVKAGDVVAISNSRGALKGNGYYKDIGQDVCIHLREQK
CTX2_Dopa 3123 bp         948 YSDYCISYICSKAPGQFDFSDTGGCKSNEFIIQNYEHPNSTTPIRNGDIVIISADNGALKGNGYNRKITLKKCSFEEL--
CTX1_Dopa 3123 bp         959 YSSYCMTYTCDKKWGTFDFRSNGGCKEYEFHITKYDDRQANGPVKAGDLVIISDRRGALKGNGYYKSINQDGCITKRVSD

                         1201 ......1210........
CTX1_AESCU 3159 bp       1038 --DRIECNANAWQIFIK*
CTX2_AESCU 3129 bp       1028 --ESAECKEIAWQIFIH*
CTX1_DPEA 3156 bp        1037 --NNINCSANAWQIFIK*
CTX2_DPEA 3123 bp        1026 --ASSKCKDIAWQIFIQ*
CTX1_ABAN 2946bp          981 ----------------L*
CTX2_ABAN 2286 bp             ------------------
CTX_EBER 3159 bp         1038 --RSINCKASSWQIYIQ*
CTX2_SOFF 3129 bp        1028 --DSSECKGITWQIFIQ*
CTX1_SOFF 3141bp         1032 --NRIECNANAWQIFMQ*
CTX_ESCO 3159 bp         1038 --RSINCKASSWQIYIQ*
CTX2_SLES 3132 bp        1029 --VSSKCKDIAWQIFIQ*
CTX1_SLES 3159 bp        1038 --KNLTCHANTWQIFIK*
CTX_TRHO 3150 bp         1035 --VSSQCKEIAWQIFIQ*
CTX_SATL 3147 bp         1034 --RNMNCKASSWQIHIK*
CTX2_ALYC 3129 bp        1028 --ESAECKEIAWQIFIQ*
CTX1_ALYC 3159 bp        1038 --DRIECNANAWQIFIK*
CTX_EPAR 3150 bp         1036 --RSLNCKASSWQIYIQ*
CTX2_APHAR 3129 bp       1028 --ESAECKDIAWQIFIQ*
CTX1_APHAR 3159 bp       1038 --DRIECNANAWQIFIK*
CTX_WSCI_PARTIAL 1653 bp      ------------------
CTX_IHAL 3156 bp         1037 --SSIDCKANAWQIYVK*
CTX2_XNOT 3183 bp        1046 --DSFECKSIAWQLFIQ*
CTX1_XNOT2 3150 bp       1035 --KRVDCKANAWQIYVK*
CTX_OROB_PARTIAL 1431 bp      ------------------
CTX_SAFF 3147 bp         1034 --RNMNCKASSWQIHIK*
CTX_SOBS 3156bp          1037 --RSMTCKASSWQIYIK*
CTX_Sarc 2907 bp          930 IASHLNCNANAWQIFIQ*
CTX2_Dopa 3123 bp        1026 --ASSKCKDIAWQIFVQ*
CTX1_Dopa 3123 bp        1039 --NNINCSANAWQIFIK*

```

**Figure S1: Nucleotide and Protein alignment of deca-ctx found in 18 squid and cuttlefish species. (A)** Nucleotide Sequence alignment of *deca-ctx* Coding Sequence. **(B)** Protein Sequence alignment of DECA-CTX protein.

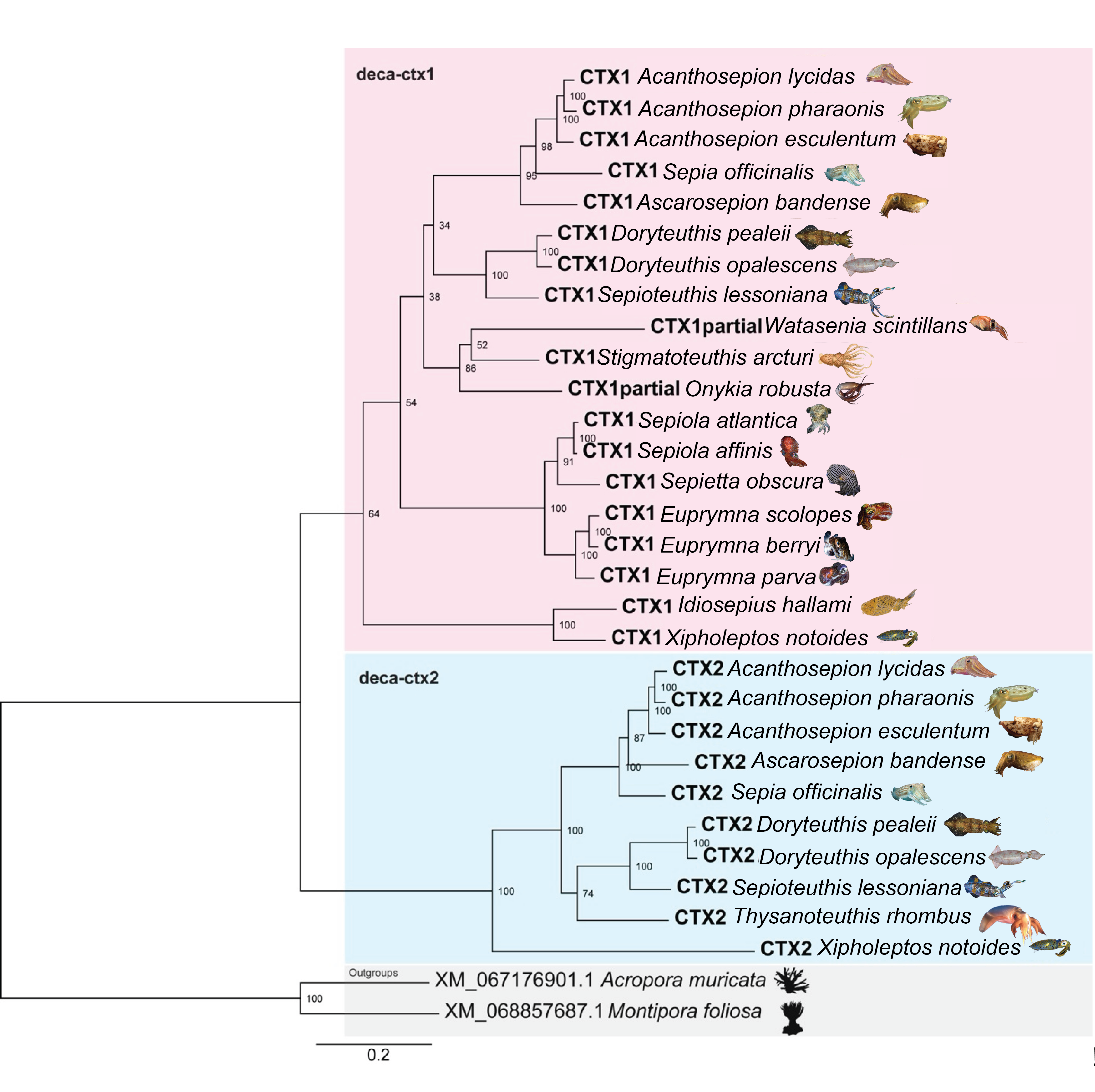

**Figure S2: Rooted tree of Duplication of cephalotoxin (*se-ctx*) present in squid and cuttlefish lineages**. Maximum likelihood phylogentic tree rooted with “*se-ctx*-like“ sequences found in cnidarians *Acropora muricata* and *Montipora foliosa* that were previously reported is shown (30% similarity). We present the rooted tree as these sequences are routinely cited as “*se-ctx like*,” and it has been reported in the proteome of the cnidarian *Bunodactis verrucosa* with a potential venomous function in predation of molluscs. The branching of the trees indicates a duplication event with two distinct clades distinguished by the presence of two Cephalopoda paralogous se-ctx genes, *deca-ctx1* and *deca-ctx2,* highlighted in pink and blue respectively. The grey area highlights the non-cephalopods with “se-*ctx* like" homologs. The values represent the node support in percentage (1000 bootstrap). The GenBank accession numbers are given for outgroup species.

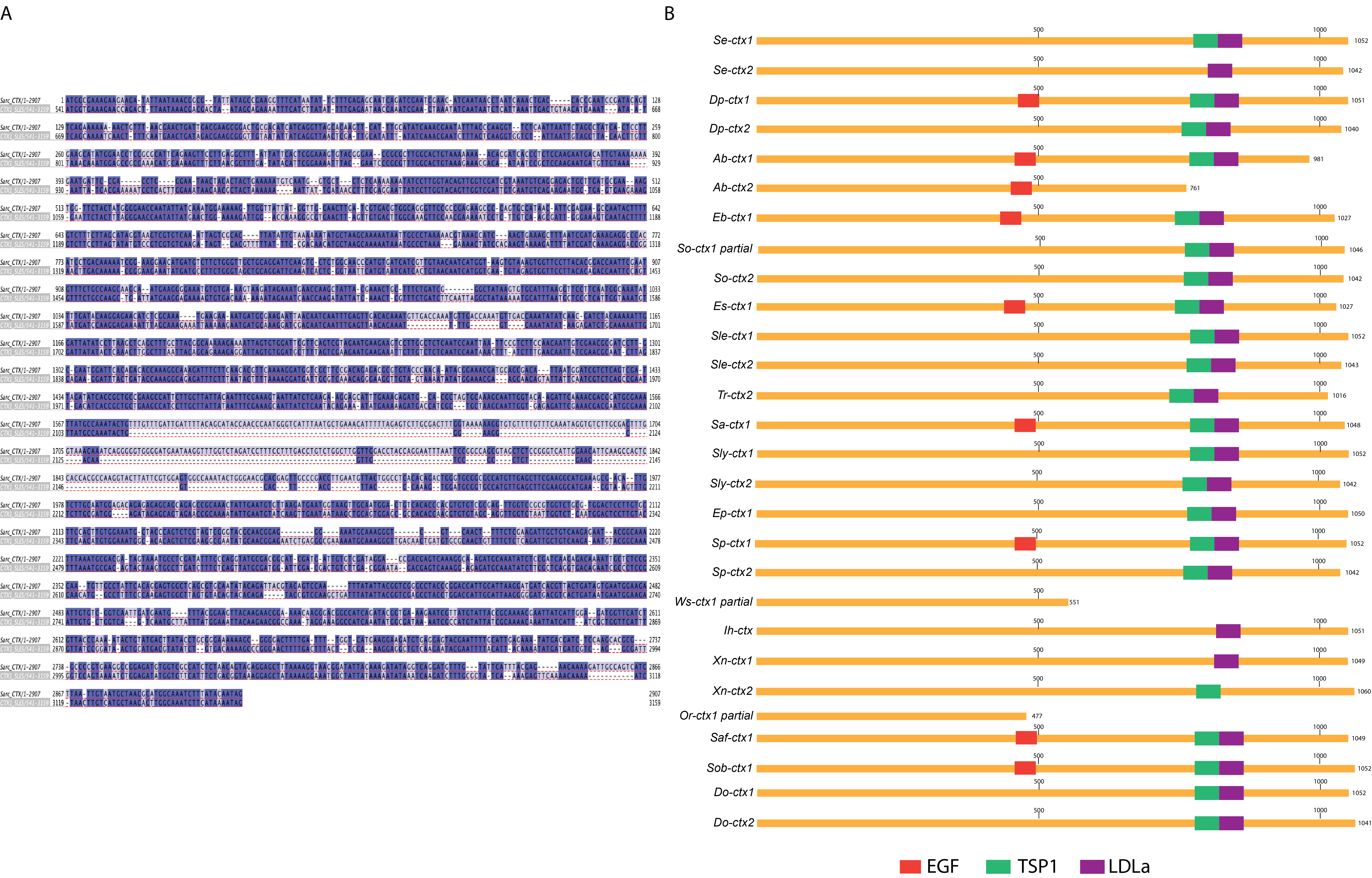

**Supplementary Figure 3. Pairwise alignment and Conserved domains among the 28 putative deca-ctx paralogs**. **(A)** Pairwise genomic sequence comparison between *Sarc_ctx* and *sly_ctx1*. *sarc_ctx* appears to code for a heavily truncated protein due to an apparent large deletion in the middle of the gene. Whether this represents a true deletion or sequencing error is unknown. **(B)** NCBI conserved domain search was used to identify conserved domains present in each of the predicted protein sequences. Predicted domains include EGF(red), TSP1(green) and LDLa(purple).

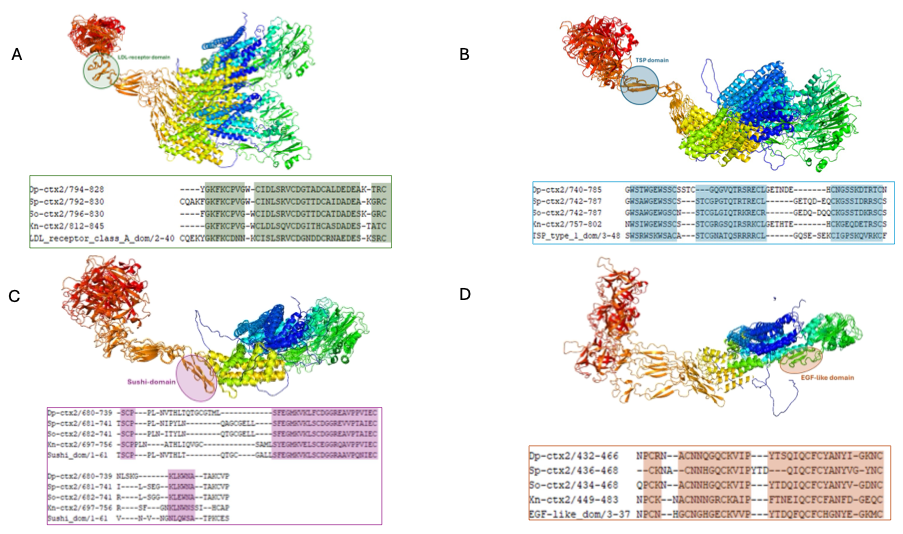

**Supplementary Figure 4: Structural and sequence domain alignment to the CTX homologs.** Regions of alignment to LDL-receptor-class-A **(A)**, TSP-type-1 **(B)**, Sushi**(C)**, and EGF-like **(D)** domains to Cluster 1 are highlighted within colored circles corresponding to the sequence alignment underneath each image. The sequence alignment only highlights regions of alignment between the cluster and domains with the highest conservation. The entire length of the sequence is not depicted.

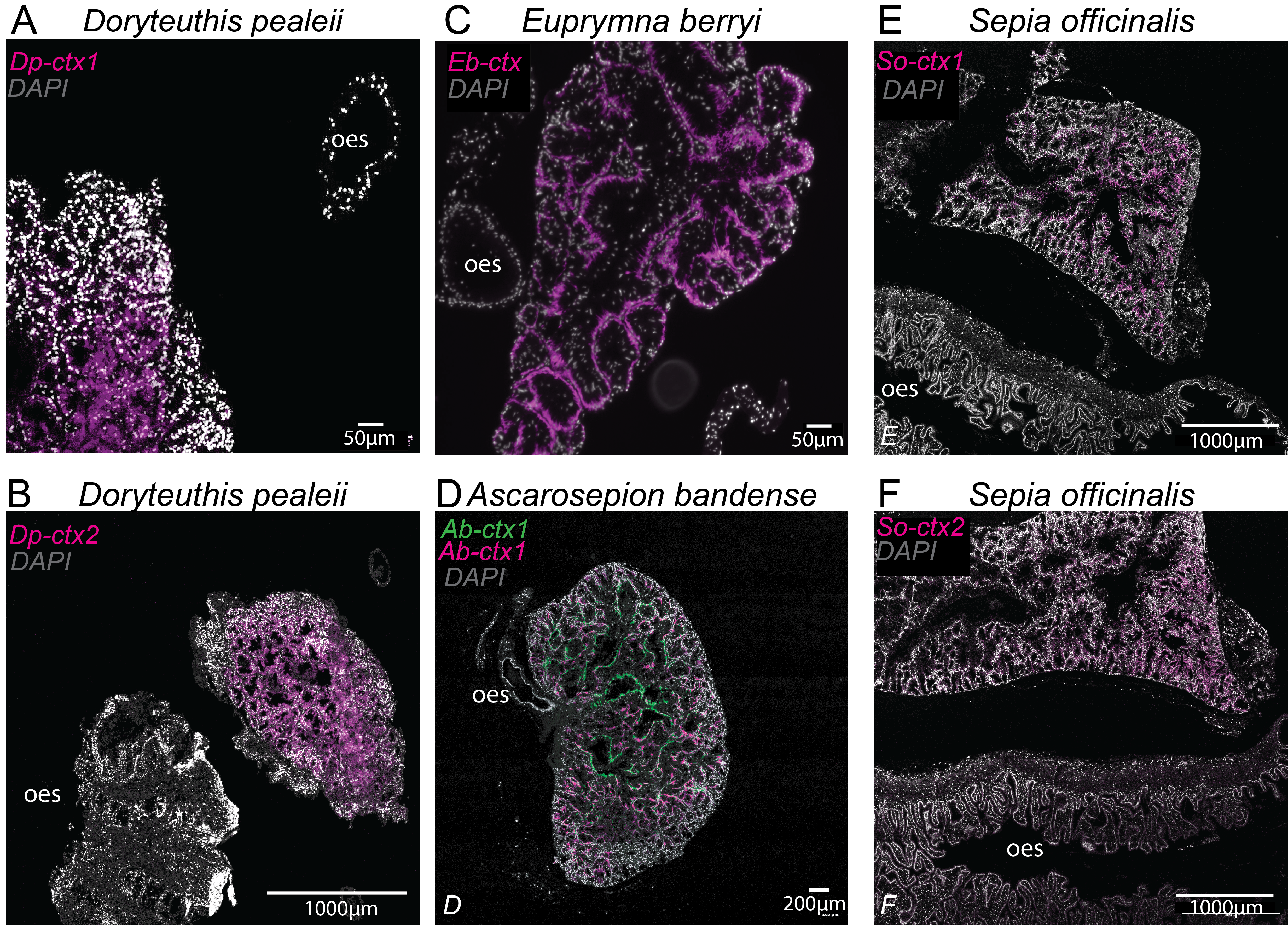

**Supplementary Figure 5: HCR supplementary figures.** HCR staining of whole adult PSG tissue sections from **(A,B,G,H)** *D.pealeii* **(C)** *E.berryi* and **(D)** *A.bandense and* ***(E,F)*** *Sepia officinalis* showing HCR staining is PSG tissue specific. Oes = oesophagus.

**A**

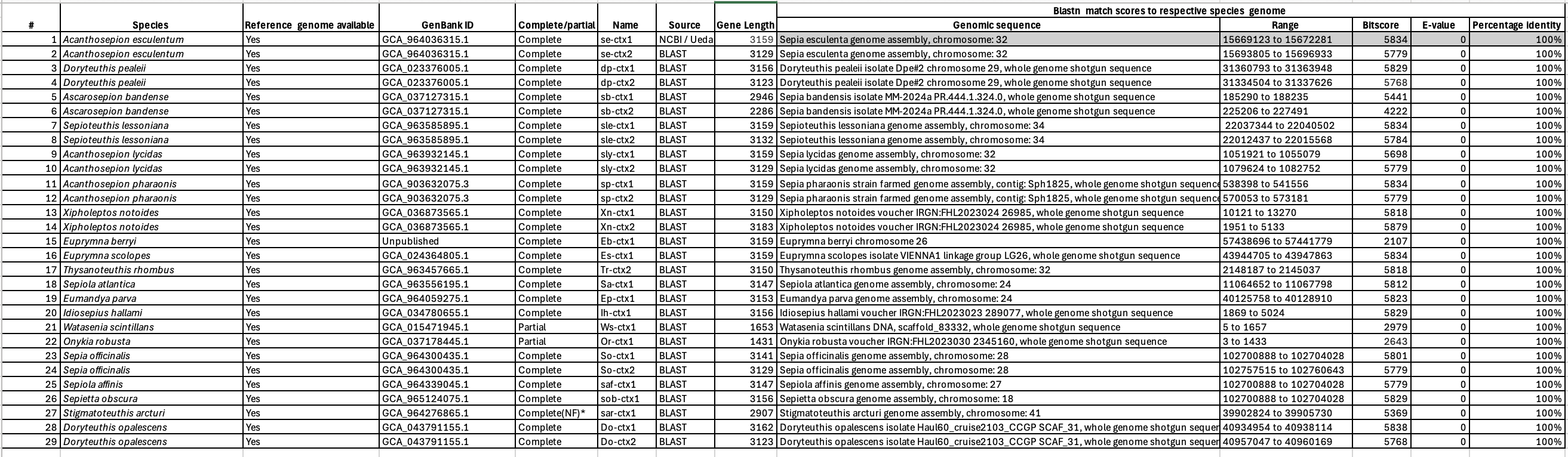

**B**

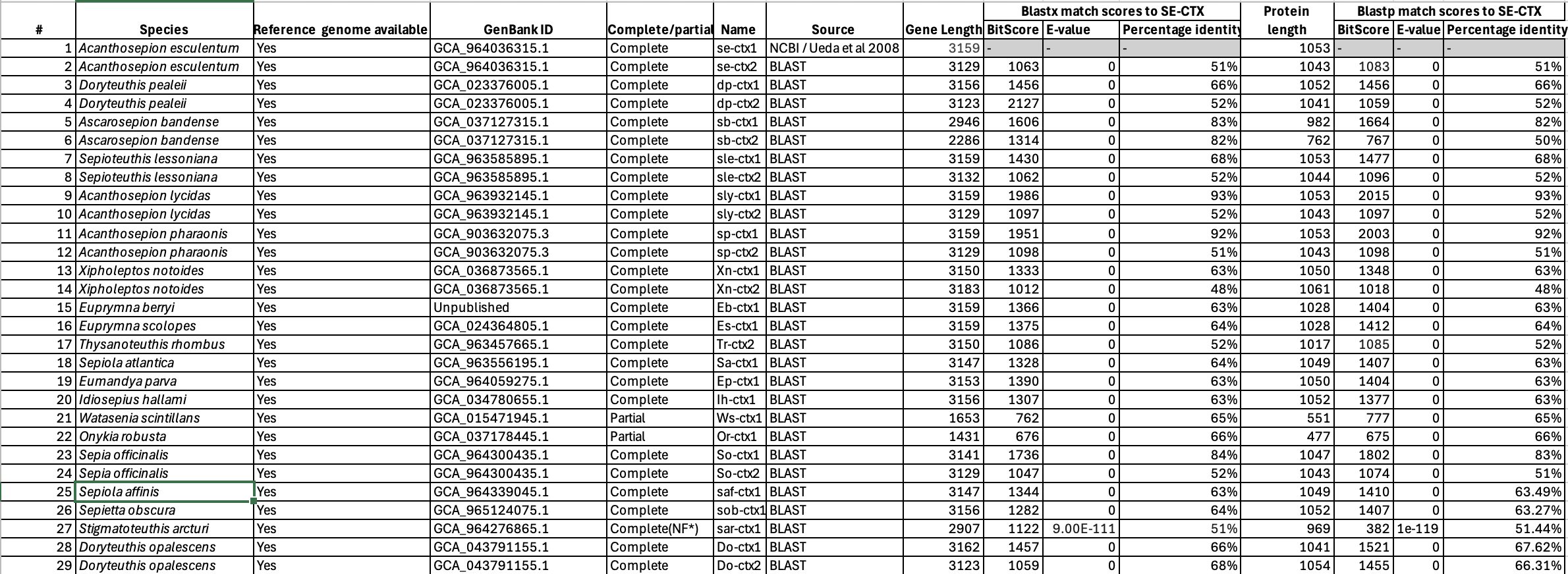
 **Supplementary Table 1. Putative *se-ctx* homologs identified from 16 species. (A)** Genomic locations of deca-*ctx* genes and TBLASTN results confirming presence of putative deca-*ctx* genes in their respective genomes. **(B)** BLASTX and BLASTP match scores to the original reference *se-ctx* sequence

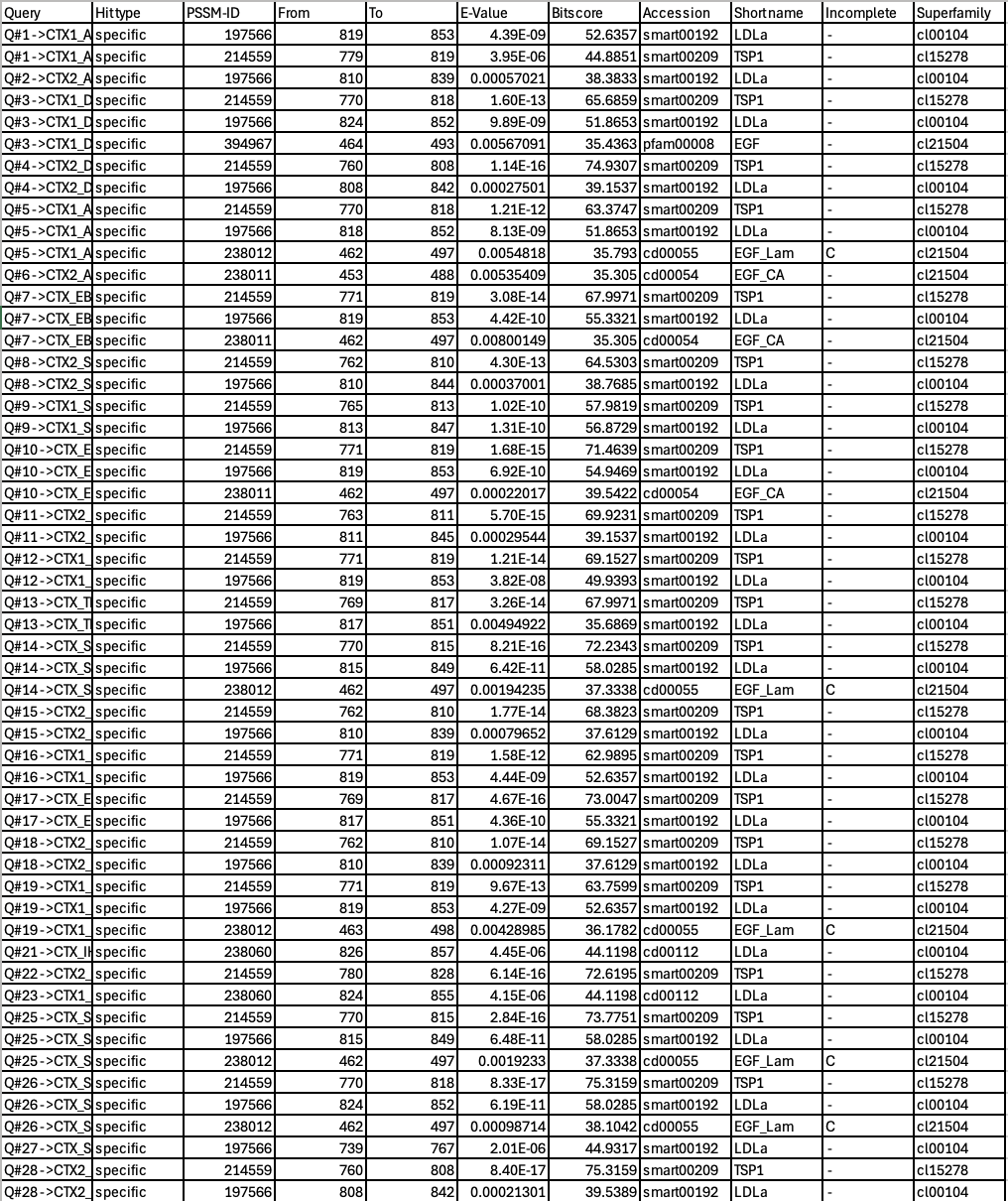

**Supplementary Table 2.** Detailed results of NCBI conserved domains search, showing bitscore and E-values for individual domain results for each putative *ctx* gene.
