## Supplementary material for "Lineage-Specific Venom Gene Expression Shapes Chemical Diversity in Cephalopods": Main Figures

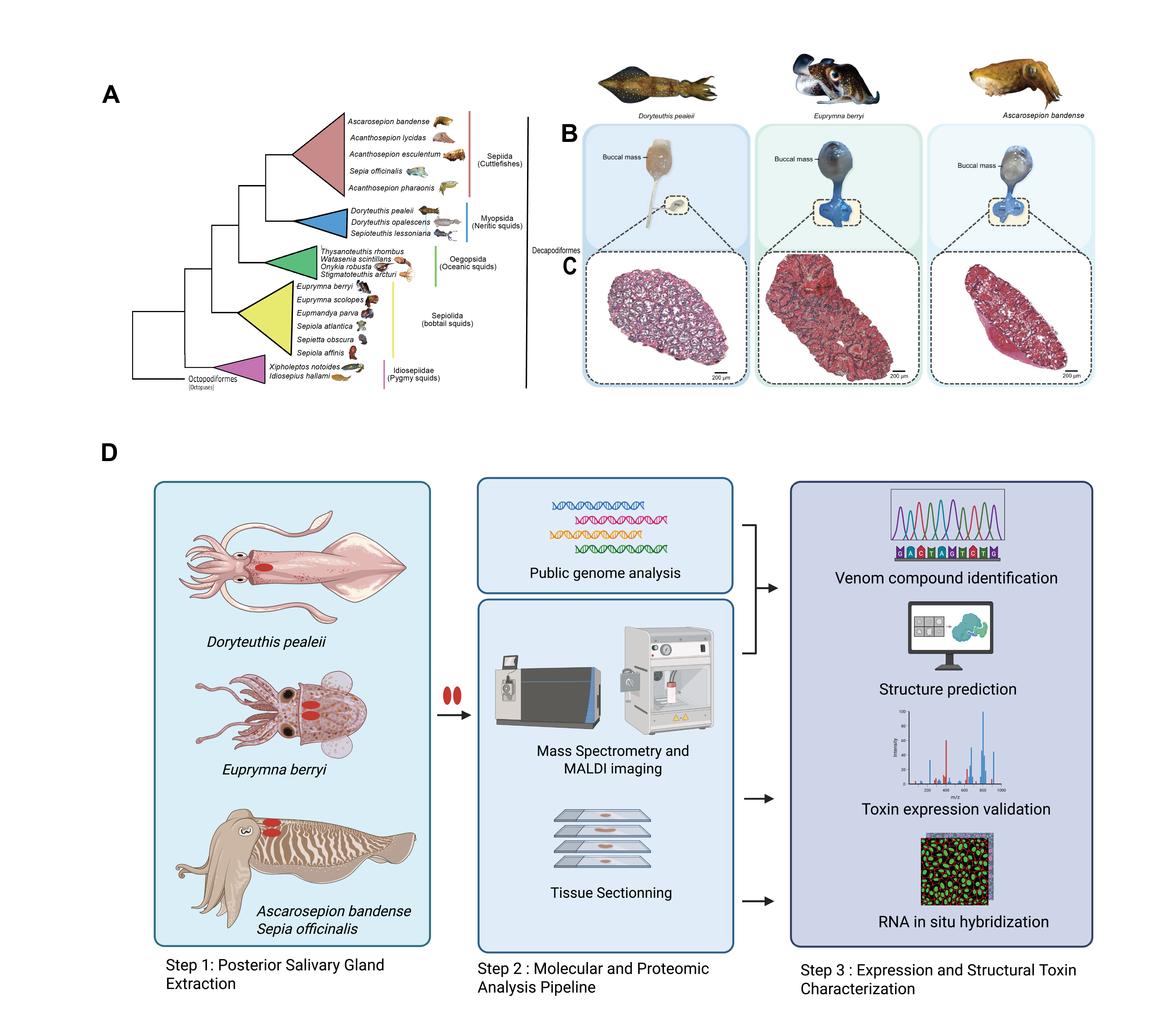
**Figure 1. Decapodiformes are squids and cuttlefish with single or paired and posterior salivary glands. (A)** Coleoid cephalopods can be broadly divided into decapodiformes (squids, cuttlefish) and octopodiformes. (**B)** Posterior salivary glands (PSG) were extracted from adult D*oryteuthis pealeii, Euprymna berryi* and *Ascarosepion bandense.* Positions of the intact salivary gland relative to esophagus and buccal mass for each of the three species are shown in cartoons and as actual extracted tissue**. (C)** H&E staining of paraffin-embeddings of these PSG sections reveal tissues with similar tubular morphologies. *E. berryi* and *A. bandense* appear to have more similar tubular structures in their PSG as compared to D*. pealeii.* **(D)** Pipeline used to identify, visualize and characterize putative cephalotoxin homologs across squid and cuttlefish species. In Step 1: Posterior salivary glands were extracted from *Doryteuthis pealeii*, *Euprymna berryi*, *Ascarosepion bandense* and *Sepia officinalis* species. In Step 2: Extracted posterior salivary glands were processed for RNA sequencing, Mass spectrometry and tissue sectioning. In Step 3: Publicly available genomic data along with generated transcriptomic data was used to identify the sequences of putative homologs to the previously identified *se-ctx* gene for available species. Mass spectrometry was used to validate venom gene expression and RNA-hybridization chain reaction was used to visualized expression of venom gene transcripts in adult and hatchling tissue sections. Protein structure prediction of the putative sequences was performed using AlphaFold2.0.

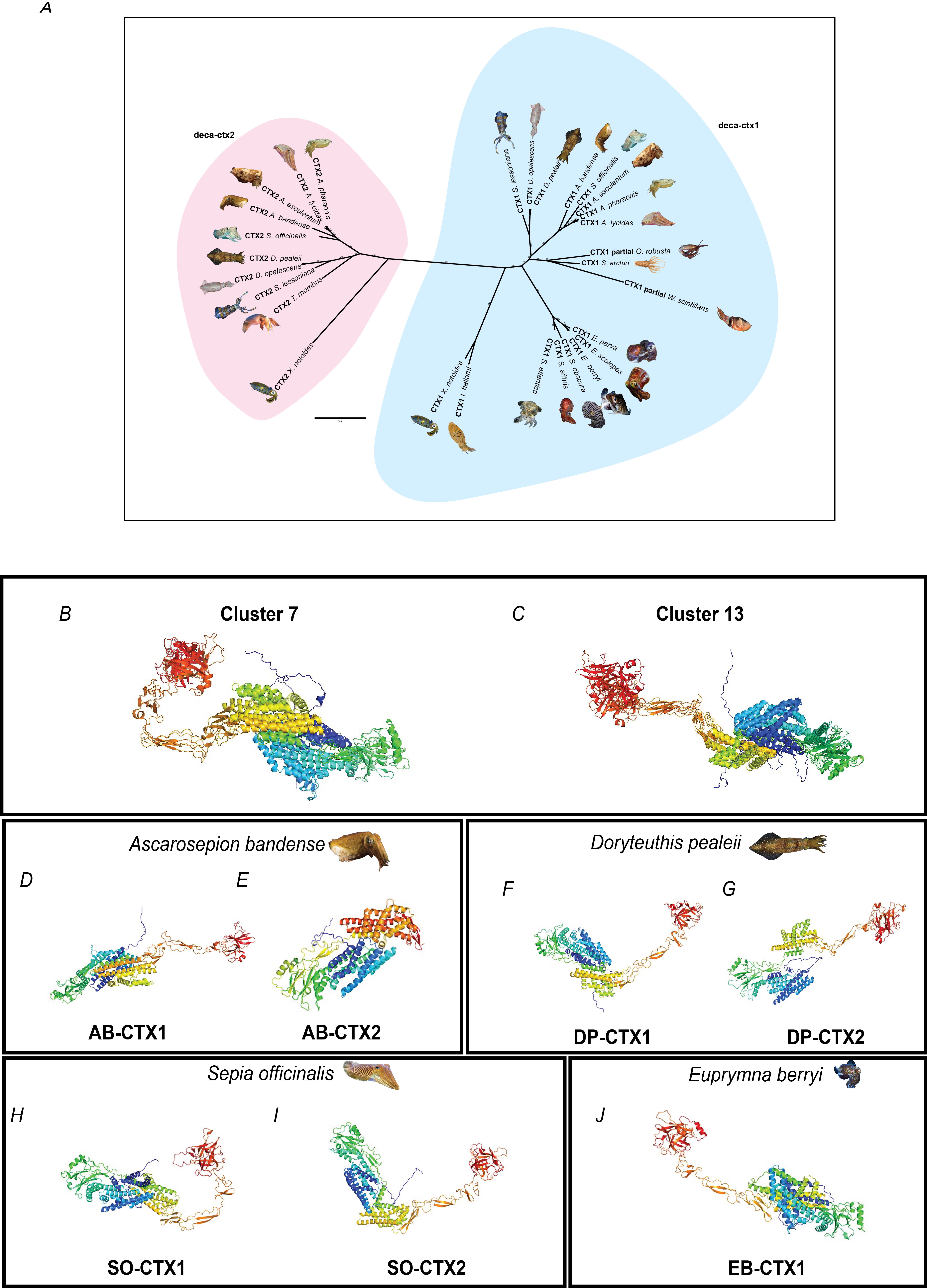

**Figure 2: Duplication of cephalotoxin *se-ctx* gene present in squid and cuttlefish lineages display diversity of chemical structures. (A)** Unrooted Maximum likelihood inference of the *se*-*ctx* homolog sequences of cephalopods. Two distinct paralogous se-*ctx genes, deca-ctx1*and *deca-ctx2*, are revealed in the tree reconstruction. The blue clade indicates *deca-ctx1* and includes genus *Euprymna, Eumandya, Sepiola, and Idiosepius genera* species exclusively. The pink clade indicates *deca-ctx2* and includes genera *Ascarosepion, Acanthosepion, Sepia, Doryteuthis, and Sepioteuthis genera,* but no Bobtail squid (*Euprymna)* species. The values represent the node support in percentage (1000 bootstrap). Cephalopoda species curated in this database version (Accessed July 2025) are included in the present tree. **(B-J)** AlphaFold2.0 predicted structures and clustering of Deca-CTX homologs. All predicted structures were clustered with qTMclust from TM-align and fell into two major clusters: (B) Cluster7 made up of 2 structures, and (C) Cluster13, also made up 3 structures. Shown also are representative singleton predicted structures of species-specific Deca-CTX from (D,E) *Ascarosepion bandense,* (F,G) *Doryteuthis pealeii,* (H,I) *Sepia officinalis*, and *(J)* *Euprymna berryi.* All structures are colored by chainbow using PyMol.

**Figure 3: Visualisation of *dual deca-ctx* expression in adult posterior salivary gland tissue indicate heterologous tissue localization.** Hybridization Chain Reaction fluorescence microscopy images of PSG sections in adult specimens showing the expression of **A)** *Doryteuthis pealeii* dp-*ctx1* mRNA (magenta) at **(i)** 20x and **(ii)** 63x magnification; *Doryteuthis pealeii dp- ctx2* mRNA(yellow) at **(iii)** 20x and **(iv)** 63x magnification; dp-*ctx1* mRNA (magenta) and *dp- ctx2* mRNA(yellow) at **(v)** 20x and **(vi)** 63x magnification. **B)** *Ascarosepion bandense* ab-*ctx1* mRNA (green) at **(i)** 20x and **(ii)** 63x magnification; *Ascarosepion bandense ab-ctx2* mRNA(magenta) at **(iii)** 20x and **(iv)** 63x magnification; ab-*ctx1* mRNA (green) and *ab-ctx2* mRNA(magenta) at **(v)** 20x and **(vi)** 63x magnification. **C)** *Sepia officinalis so-ctx1* mRNA (green) at **(i)** 20x and **(ii)** 63x magnification; *Sepia officinalis so-ctx2* mRNA(magenta) at **(iii)** 20x and **(iv)** 63x magnification; *so-ctx1* mRNA (green) and *so-ctx2* mRNA(magenta) at **(v)** 20x and **(vi)** 63x magnification. **D)** *Euprymna berryi* *eb-ctx1* mRNA (magenta) at **(i)** 20x and **(ii)** 63x magnification. Sections are counterstained with DAPI (grey) to visualize nuclei.

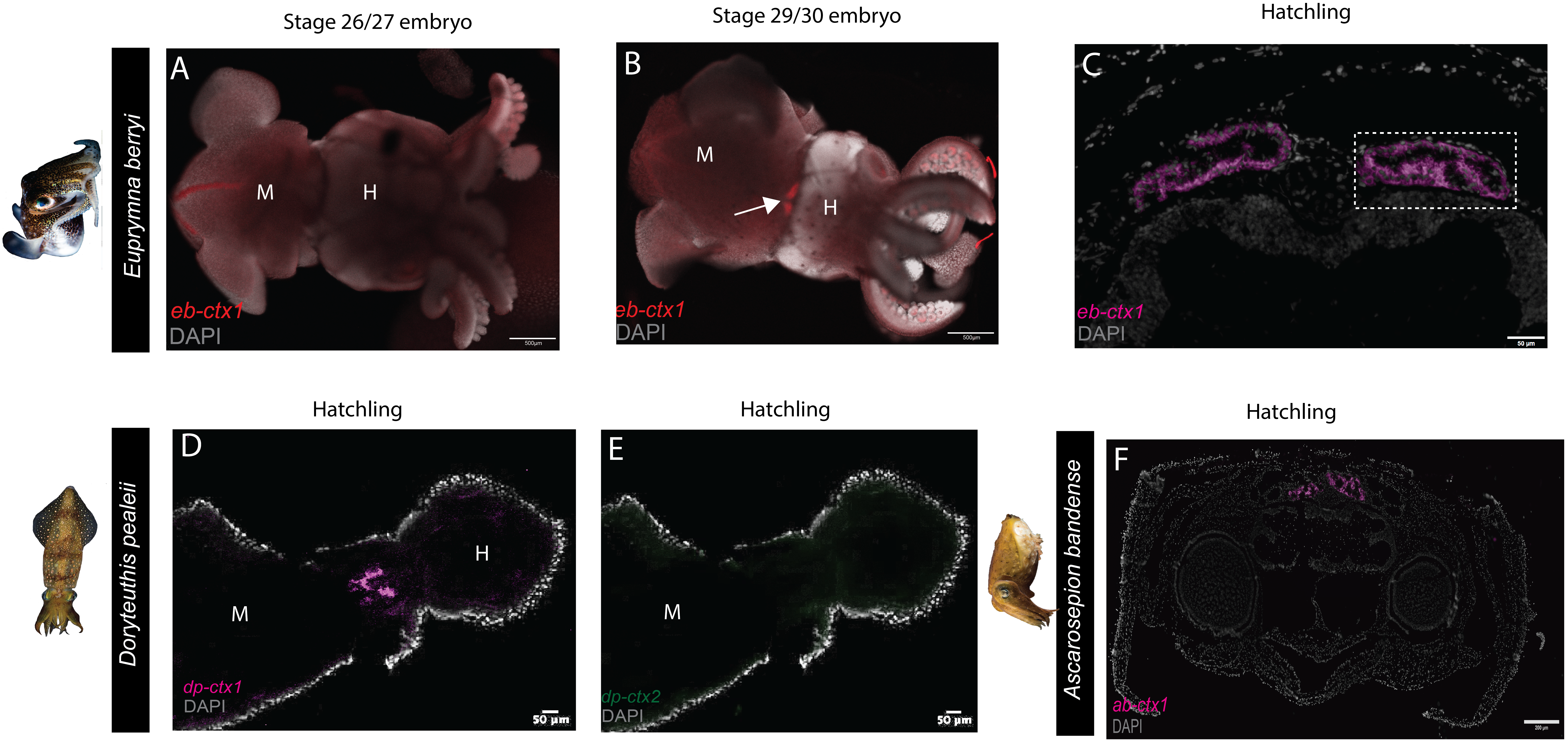

**Figure 4: Differential developmental expression of deca*-ctx* homologs in squid and cuttlefish hatchlings and embryos.** Hybridization Chain Reaction fluorescence microscopy images. **(A)** Whole mounts of *E.berryi* embryos were stained with *eb-ctx1*-specific (red) probe and DAPI revealed no staining in day 26/27 embryos. The redline shown on the mantle opposite the tenetacles is distinctive nonspecific staining in *E. berryi* embryos. (**B) C**lear staining was seen in day 29/30 embryos/hatchlings. White arrow indicates PSG-specific expression. (**C)** PSG-specific expression of *eb-ctx1* mRNA in *E.berryi* hatchling sections. The two E. berryi PSGs both express *eb-ctx1*. Enlarged view highlighted in the white hashed-box. Whole mounts of *D.pealeii* hatchlings were stained with *dp-ctx1-*specific (magenta) probe, *dp-ctx2-*specific (green) probe and DAPI, **(D)** Indicates *dp-ctx1* staining in the region of the two PSGs. **(E)** No *dp-ctx2* staining is present in the same whole mount.. **(F)** Similar to *E. berryi,,* we observe PSG-specific expression in both glands of *Ab-ctx1* mRNA in *A .bandense* hatchling sections. Sections are counterstained with DAPI (grey) to visualize nuclei.

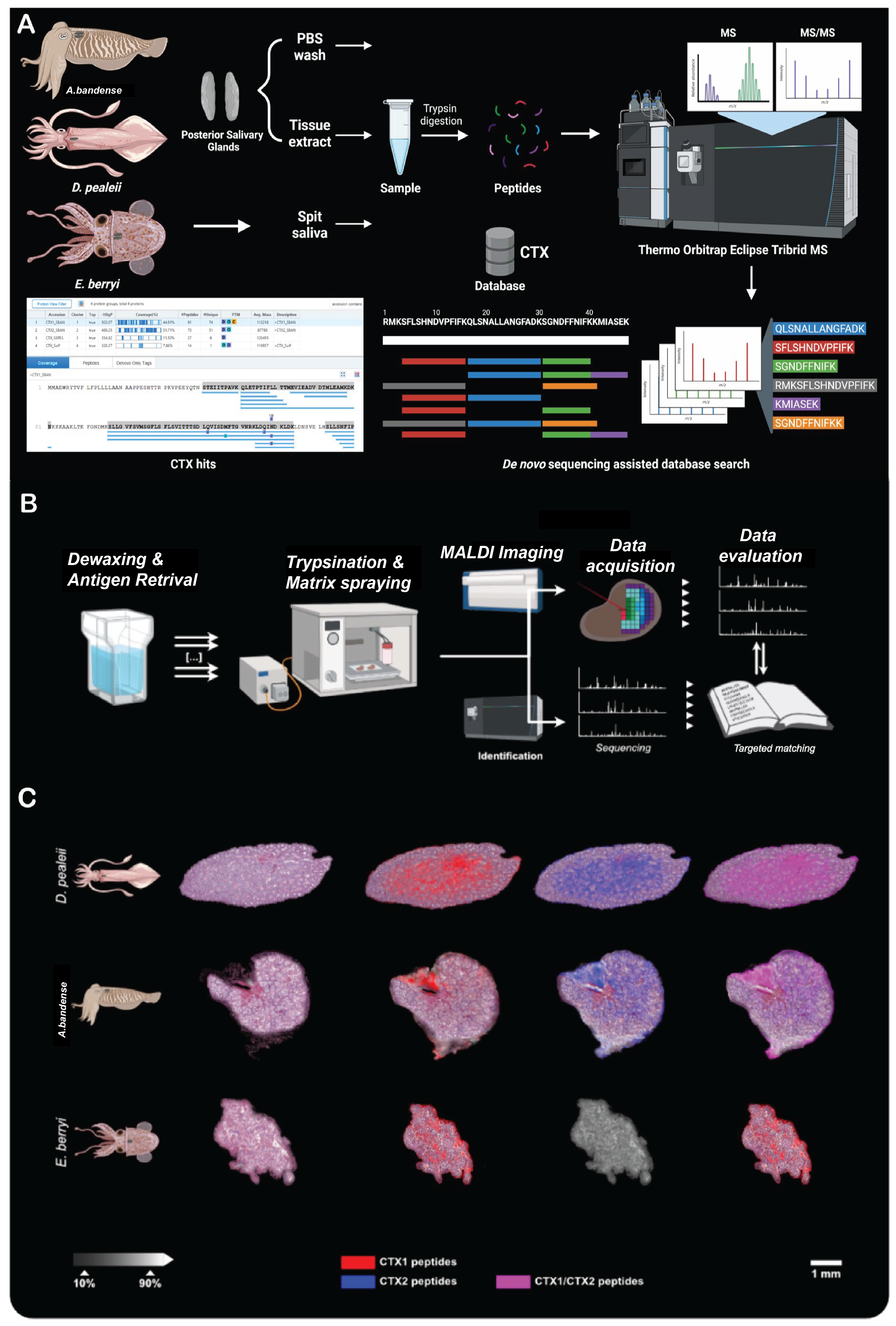

**Figure 5. Proteomic confirmation of DECA-CTX protein expressions validates presence of transcripts in squid and cuttlefish venom glands.** Shown is the bottom-up and spatial venom proteomic analyses of posterior salivary gland tissue sections for three different cephalopod venom systems. **(A)** Schematic of the mass spectrometry (LC-MS/MS) workflow used to identify cephalotoxins (CTXs) from the posterior salivary glands (PSGs) of *A.bandense, D. pealeii*, and *E. berryi*. PSGs from *A.bandense*, *D. pealeii*, and *E. berryi* were dissected, homogenized in PBS, lyophilized, and processed for proteomic analysis. **(B)** Preparation and analysis of spatial venomics using matrix-assisted laser desorption/ionization (MALDI) mass spectrometry imaging (MSI). Tissue sections of venom glands from *D. paeleii*, *E. berryi*, and *A. bandense* were deparaffinized, rehydrated, tryptic digested and loaded on to a ultrafleXtreme MALDI-ToF/ToF mass spectrometer. Raw data were processed using baseline correction, TIC normalization, and peak alignment. Peptide matches between MALDI-MSI and nLC-MS peptide library were identified using an in-house script, matching m/z values within 0.2 Da and selecting peptides based on the highest confidence score. **(C)** Visualization of DECA-CTX1, red), DECA-CTX2, blue) and both DECA-CTX1 and DECA-CTX2, purple) by characteristic m/z values (mass features) within the venom gland system (20 μm spatial resolution). The spatial venom compound distribution and internal relative intensities (white scale bar) are shown for a set of three mass features per compound and species Top row: *D. pealeii* composed of m/z 1376.79, 1697.91, 2458.13 (DECA-CTX1) and m/z 1287.74, 1725.86, 2082.08 (DECA-CTX2). Middle row: *A.bandense* composed of m/z 1123.60, 1564.65, 1709.61 (DECA-CTX1) and m/z 1621.85, 1974.82, 2226.06 (DECA-CTX2). Bottom row: *E. berryi* composed of m/z 1584.82, 1798.66, 2229.96 (DECA-CTX). The histological image was prepared post-MALDI-MSI by hematoxylin/eosin (H/E) staining and is shown for tissue section orientation. The spatial segmentation of the venom gland systems resulted in different numbers extracted peak ions (±0.47 Da) from normalized spectra within an m/z 600−3200 mass range (see Supplementary Tables 2).
